# AquIRE reveals multiple mechanisms of clinically induced RNA damage and the conservation and dynamics of glycoRNAs

**DOI:** 10.1101/2025.04.01.646554

**Authors:** Zijian Zhang, Zornitsa Vasileva Kotopanova, Kexin Dang, Xiangxu Kong, Nicole Simms, Tin Wai Yuen, Aynur Sönmez, Lauren Forbes Beadle, Emma Hilton, Taqdees Qureshi, Marianna Coppola, Callum David Holmes, Kwan Ting Kan, Mark Ashe, Patrick Gallois, Hilary Ashe, Denis L.J. Lafontaine, Michael Braun, Mark Saunders, Paul Sutton, David Thornton, John R. P. Knight

## Abstract

RNA is subject to many modifications, from small chemical changes such as methylation through to conjugation of biomolecules such as glycans. As well as these endogenously written modifications, RNA is also exposed to damage induced by its environment. Certain clinical compounds are known to drive covalent modifications of RNA with a growing appreciation for how these affect function. To understand the regulation of these modifications we need a reliable, sensitive and rapid methodology for their quantification. Thus, we developed AquIRE and applied it to the analysis of drug-induced RNA damage, showing this to be widespread with intricate temporal dynamics. Using the same methodology we identify RNA:protein crosslinking and the rewriting of the epitranscriptome as a consequence of clinical RNA damage. We also demonstrate how liquid-liquid phase separation increases RNA damage and expand the horizons of the glycoRNA world across the kingdoms of life and into cell-free glycoRNA.

## Introduction

Antimetabolites or alkylating agents are administered to over 2 million people worldwide each year to manage diseases including cancer, autoimmunity and viral infections^1,2^. These agents are multifaceted in their mechanisms of action, leading to detrimental side effects along with clinical benefits. The advent of targeted therapies has meant a sharp decline in approvals for new antimetabolite or alkylating agents yet for many diseases they remain the best or only clinical options. In fact, globally the usage of these drugs continues to grow. Mechanistically, these compounds cause damage to multiple target biomolecules, which occurs in a specific manner related to the chemical structure of the drug. Surprisingly, given their extensive clinical use, their exact mechanisms of action remain poorly understood.

Our knowledge of chemotherapies has traditionally been DNA-centric with their effects on other biomolecules considered unimportant compared to DNA damage. Recent reports have challenged this dogma and presented RNA as an important target for clinical compounds^1,3^. For instance, it is known that incorporation of the antimetabolite 5FU into RNA correlates with its cytotoxicity^4–7^, while the pseudo-alkylating agent oxaliplatin impacts RNA-related functions unlike its related compounds cisplatin and carboplatin^8^. Highlighting the biological importance of RNA damage, intricate molecular mechanisms to detect and manage its impact have recently been described^9–15^. This work has primarily used tool compounds and physical stresses such as irradiation, raising the question of when these pathways become engaged in the physiological setting. Addressing this, roles for RNA damage pathways have recently been shown to impact inflammation, stem cell function and aging^16–19^. With some RNAs known to be as long lived as DNA in non-mitotic cells^20^, understanding the physiological causes and dynamics of RNA damage is a pressing need.

To understand RNA as a direct drug target, the field requires an easy to implement method to directly measure specific RNA damage from multiple compounds, ideally from low input samples. With this in mind, we developed AquIRE – Aqueous Identification of RNA Elements – and used this to quantify relative levels of direct RNA damage from the antimetabolite 5FU and the (pseudo)alkylating agents oxaliplatin and temozolomide. Our methodology is remarkably sensitive and has intrinsic flexibility for the desired target of interest. As such, we further applied AquIRE to three different RNA elements, endogenously written chemical modifications (m6A and pseudouridine), RNA:protein crosslinks driven by proximity and reactive oxygen species (ROS) and cell-bound and cell-free glycoRNAs. Throughout we reveal the pervasive effects of drug-induced RNA damage on RNA biology; showing the relationship between drugs, the epitranscriptome and specific RNA:protein crosslinking events.

Using AquIRE to detect glycoRNAs allows their analysis from as little as 10ng of RNA using an easily accessible and highly reproducible protocol. We vastly expand the known breadth of glycoRNA expression to include organisms from *Xenopus* to plants to single-celled microbes and prokaryotes. In parallel, we reveal glycoRNA expression dynamics during the earliest stages of development and in senescence and that glycoRNAs are present in liquid, cell-free, samples from 7 different organisms. Finally, we show that multiple RNA-damaging clinical compounds elevate glycoRNA expression and that glycoRNAs are required for optimal cytotoxicity from 5FU. Altogether this work provides a new method to identify and study RNA elements that will address key questions about epitransciptomics, RNA-binding protein biology, glycoRNAs and the effects of clinical RNA damage.

## Results

### AquIRE detects 5FU:RNA localisation and dynamics

We have previously used an anti-BrdU antibody to detect 5FU in RNA at single cell resolution^21^. Here, we used this same antibody in an RNA dot blot to quantify 5FU incorporation into RNA after treatment of colorectal cancer cells (HCT116). In parallel, we analysed IVT RNA made in the presence of 5FUTP, as a positive control. Unexpectedly, we observed no signal from cellular RNA (**Supplemental Figure 1A**) under the same conditions where we have previously observed 5FU in RNA by immunofluorescence^21^. We were able to detect a signal from IVT RNA with 100% 5FUTP incorporation but this signal dropped off sharply at 75% and was barely visible at 25%, resulting in a non-significant correlation between 5FU content and signal (**Supplemental Figure 1B**). We hypothesised that immobilisation of purified RNA on membranes occludes or destroys our epitope. Consistently, the fluorine epitope of 5FU sits on the Hoogsteen base edge, while membranes are designed to display the Watson-Crick edge for hybridisation^1^. Furthermore, adducts of 5FU that occur when crosslinking RNA to membranes, result from defluorination^22^, which would destroy the epitope we want to measure.

Thus, we sought to develop an alternative membrane-independent approach to biochemically detect 5FU incorporation in purified RNA. We opted for aqueous detection to maximise the surface area of RNA, and number of epitopes available. We took advantage of the interaction between RNA polyA tails and commercially available oligodT beads to tether RNA through their 3’ ends, while leaving the full surface area of the remaining sequence accessible in aqueous solution. Crucially we developed this method using buffers amenable to coincubation with protein-based detection reagents and coupled this with the enzymatic addition of polyA tails to total RNA using commercially available polyA polymerase. We termed this method AquIRE for Aqueous Identification of RNA Elements (**Figure 1A**).

**Figure 1:**
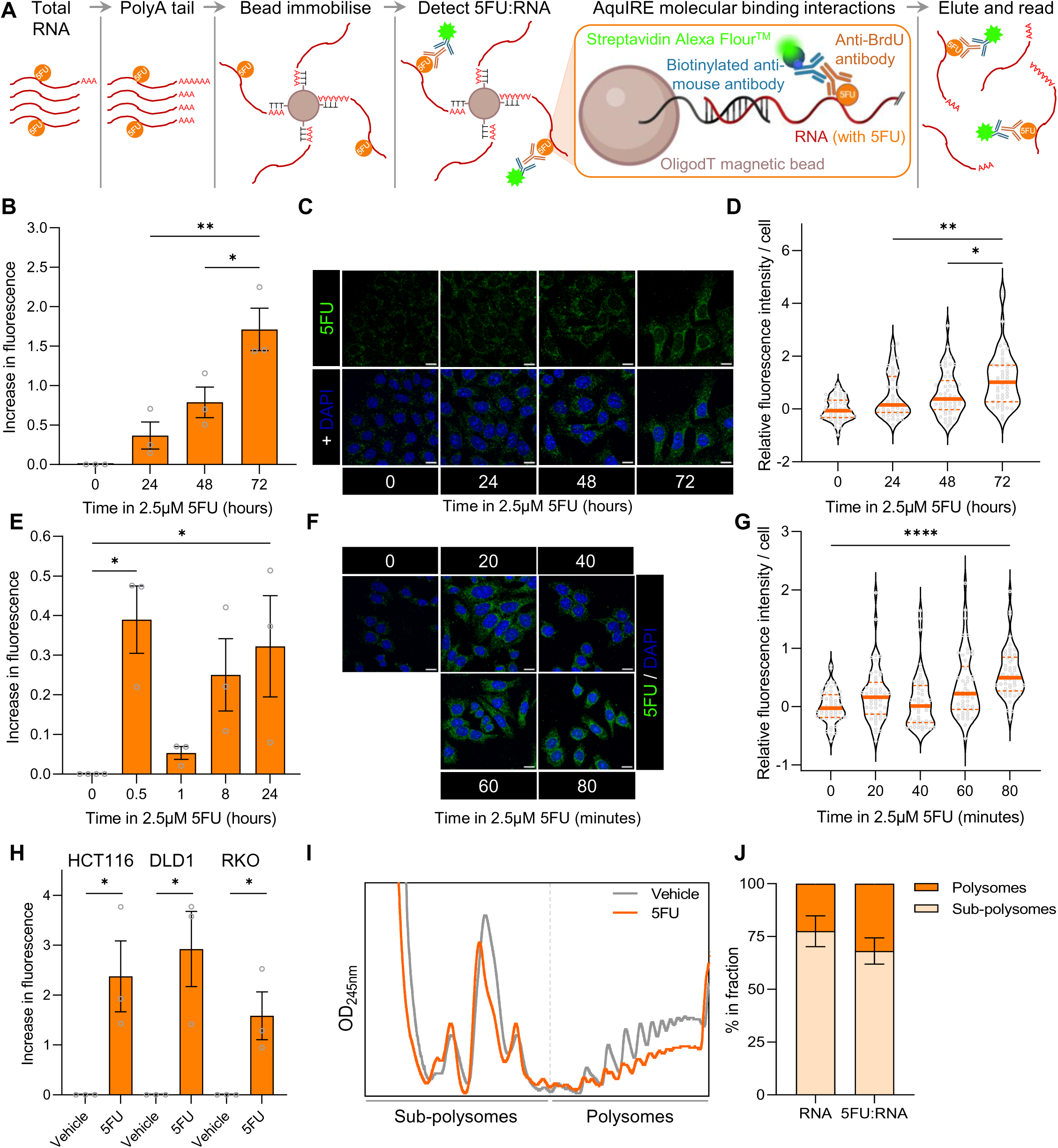
Detecting RNA damage by 5FU using AquIRE. A) Schematic of the experimental protocol for AquIRE. Total RNA, containing modifications (in this example 5FU) is polyA tailed then immobilised on oligodT beads. RNA is sequentially exposed to a primary antibody to detect 5FU, or other RNA elements, a biotin-tagged secondary antibody then Alexa Fluor^TM^-tagged streptavidin. Finally, water is used to elute a fluorescent signal proportional to the amount of 5FU in the RNA. B) HCT116 cells were treated with 2.5µM 5FU for the indicated times, RNA extracted, the equal amounts of RNA analysed for RNA content. The graph shows AquIRE fluorescent measurements normalised against vehicle treatment set to 0. Bars represent the average of 3 biological replicates, each shown as grey circles and the error bars are SEM. Significance was calculated by ANOVA using Šídák multiple comparison testing. C) HCT116 cells were treated as in A then stained for 5FU incorporation into RNA and counterstained with DAPI. Scale bar 50µm. D) Violin plots of the quantification of the 5FU incorporation in cells as shown in C. Each grey circle represents one of 50 individual cells analysed per timepoint in this biological replicate. The thick orange lines are the mean and dashed lines are quartiles. Data are presented relative to vehicle treatment (0 hours), which is set to 0. Significance was determined by Kruskal Wallis analysis with Dunn’s multiple comparison testing. Significant differences between drug treated groups are annotated. All treatments were significantly different to 0 hours. E) HCT116 cells were treated with 5FU at 2.5µM for the times shown and analysis performed as in A. Bars represent the average of at least 3 biological replicates for each timepoint, shown as grey circles, and the error bars are SEM. Significance was analysed by ANOVA. F) HCT116 were treated with 2.5µM 5FU for the indicated times and presented as in C. Scale bar 50µm. G) 5FU intensity per cell from F was calculated for 50 cells per indicated timepoint. Data are presented in violin plots as described in D. Significance was calculated using a Kruskal Wallis test. H) RNA was analysed from HCT116, DLD1 or RKO cells were treated with vehicle or 10µM 5FU for 72 hours. Graph shows the levels of 5FU incorporation relative to the vehicle set to 0. Data are n=3 biological replicates, with error bars showing SEM. Significance was determined by unpaired t-test. I) HCT116 cells were treated with 10µM 5FU for 24 hours then their cytoplasmic fraction separated by sucrose density gradient. From these gradients, OD_254nm_ polysome traces were obtained and overlaid here. Data are representative of two biological replicates. J) RNA was extracted from the sub-polysome and polysome fractions of the 5FU treated sample shown in I, with total RNA distribution between the fractions (left) and 5FU:RNA content determined by AquIRE (right) plotted ±SEM. * *P*<0.05, ** *P*<0.01, **** *P*<0.0001

First, we used IVT 5FU-containing RNAs as detailed above and saw a strong correlation between 5FU content and fluorescent signal (**Supplemental Figure 1C**). Next, we asked if AquIRE could detect 5FU incorporation into total RNA extracted from HCT116 colorectal cancer cells treated with clinically achievable doses of 5FU over a 72-hour time course. This revealed a steady increase in 5FU in RNA with increased exposure time (**Figure 1B**), correlating perfectly with 5FU incorporation measured using our previously published immunofluorescent technique (**Figures 1C-D and Supplemental Figure 1D**). We then analysed shorter, more clinically relevant timepoints. The method proved remarkably sensitive, able to detect 5FU incorporation using as little as 100ng of RNA. Importantly, we were able to recover >100% of the RNA input at the end of the experiment, showing that the sample RNA is well retained on the oligodT beads, while the increase in RNA recovery compared to input is likely due to the addition of polyA tails (**Supplemental Figure 1E**).

The short time course revealed previously unknown dynamics in the incorporation of 5FU into RNA. We observed a consistent peak at 30 minutes, followed by a marked reduction at 1 hour, preceding a steady increase back to levels seen at 30 minutes by 24 hours (**Figure 1E**). Again, this result was recapitulated by immunofluorescence, identifying a biphasic pattern in 5FU incorporation into RNA (**Figures 1F-G and Supplemental Figure 1F**). Expanding this, we confirmed the incorporation of 5FU into RNA in multiple colorectal cancer cell lines (**Figure 1H**), and that 5FU is present in translating polysomes (**Figures 1I-J**). The latter observation is consistent with our previous work that 5FU induces a robust ribosome quality control response^21^, with this work consistent with 5FU:RNA participating in translation.

### RNA damage is induced by multiple clinical agents

Having shown that the antimetabolite 5FU causes direct RNA damage we asked whether further cytotoxic chemotherapies do likewise. First, we analysed the pseudo-alkylating drug oxaliplatin and its sister compounds cisplatin and carboplatin. We adapted our AquIRE method using a different antibody that has previously been used to detect platinum adducts on DNA (**Figure 2A**). First, we detected oxaliplatin covalently bound to RNAs after a 6-hour dose of 50µM (**Figure 2B**). A lower, clinically achievable dose of 2.5µM, did not induce detectable adducts at this time, but did result in adducts after 24-hours of treatment (**Figure 2C**). In contrast neither cisplatin nor carboplatin were able to induce consistent detectable adducts on RNA either at high concentrations or after prolonged exposure (**Figure 2C**).

**Figure 2:**
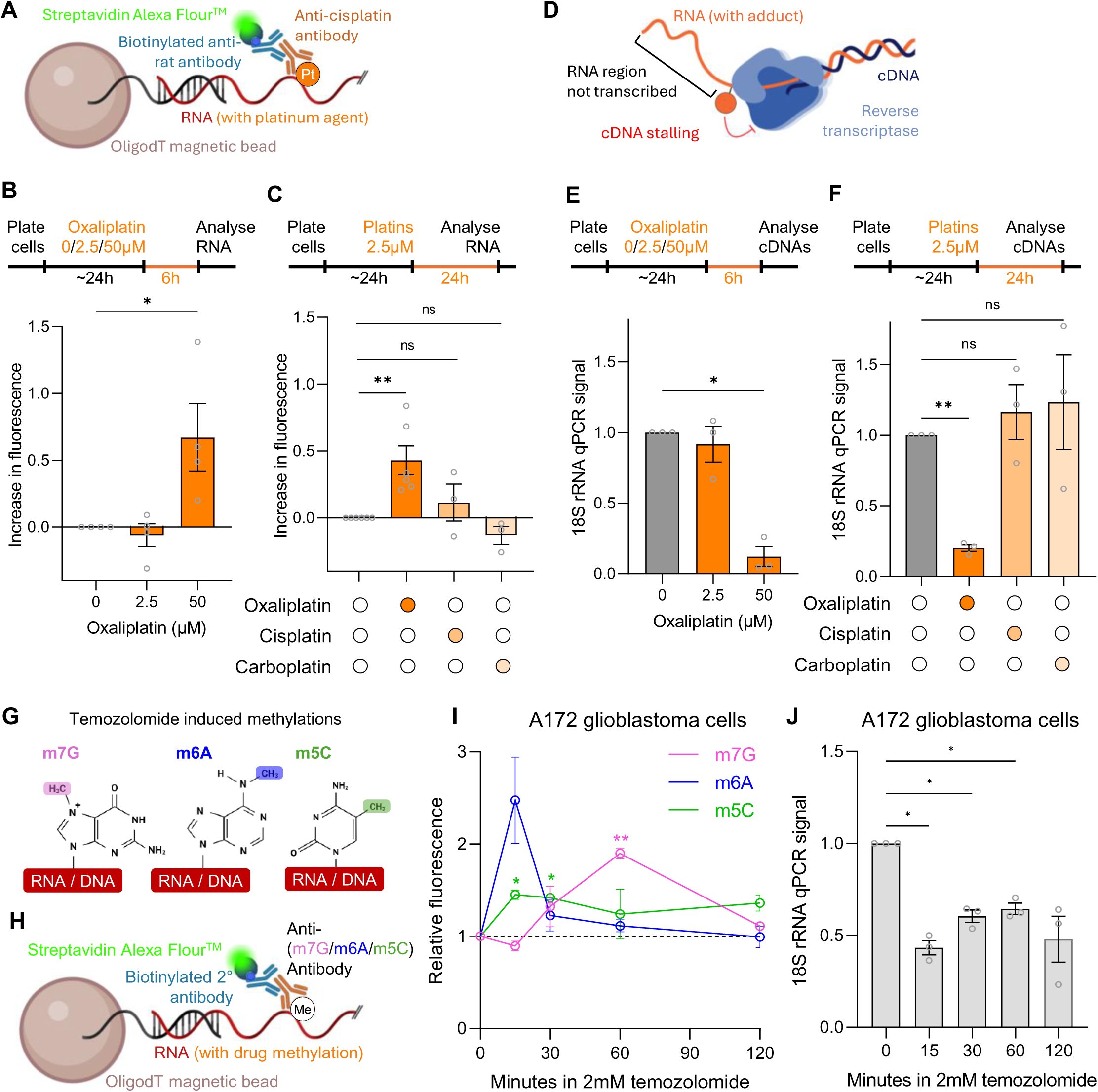
Oxaliplatin and temozolomide cause detectable RNA damage. A) Schematic of oxaliplatin:RNA detection by AquIRE. B) Top, timeline of drug treatments outlining that HCT116 cells were treated with increasing doses of oxaliplatin for 6 hours prior to extraction and analysis of total RNA. Bottom, graph plotting the average of 4 biological replicates detecting oxaliplatin by AquIRE in RNA after 0, 2.5 and 50µM treatments. Bar represents the mean ±SEM, normalised against 0µM which is set to 0. Significance was calculated by ANOVA using Šídák multiple comparison testing. C) Top, timeline of drug treatments of HCT116 cells, which were treated with 2.5µM of each platinum agent for 24 hours prior to analysis of RNA. Bottom, bar graph shows the levels of platinum agent detected in RNA, shown as the average of ≥3 biological replicates. Statistical significance, or lack thereof, was determined by ANOVA using Šídák multiple comparison testing. D) Schematic of the cDNA reverse transcriptase stalling assay. Adducts, shown as an orange ball, stop the movement of reverse transcriptase resulting in RNA that is not reverse transcribed. This can be detected by qPCR of the resulting cDNA template; lower signal signifies more adducts. E) Top, timeline of the drug treatments of HCT116 cells for this analysis, which are the same as in B. Bottom, graph depicting the levels of a qPCR amplicon in the 18S rRNA after oxaliplatin treatment compared to vehicle treatment set to 1. Significance was calculated by ANOVA using Šídák multiple comparison testing. F) Top, schematic of platinum agent treatment of HCT116 cells and analysis, which is as in C. Bottom, analysis of adduct formation after oxaliplatin, cisplatin or carboplatin treatment. Statistical significance, or lack thereof, was determined by ANOVA using Šídák multiple comparison testing. G) Representation of the effects of temozolomide on DNA and its hypothesised effect on RNA – the methylation of m7G, m6A or m5C. H) Schematic of the AquIRE detection method for temozolomide-induced RNA damage. I) A172 cells were treated with 2mM temozolomide for the indicated durations, RNA extracted and the relative abundance of m7G, m6A and m5C plotted compared to vehicle treatment set to 1. Data show the average of 3 biological replicates ±SEM for each methylation. Significance compared to vehicle was calculated by ANOVA with Šídák multiple comparison testing. J) RNA extracted from A172 cells after the indicated times in 2mM temozolomide was reverse transcribed and analysed by the cDNA stalling analysis. qPCR of the 18S rRNA amplicon are plotted for 3 biological replicates ±SEM. Significance compared to vehicle was calculated by ANOVA with Šídák multiple comparison testing. * *P*<0.05, ** *P*<0.01

To confirm drug-induced RNA damage with an alternative, transcript specific method, we adapted a previously published cDNA stalling methodology^23^ (**Figure 1D**). Here, adducts on RNA block reverse transcriptase, resulting in loss of cDNA that is detected by qPCR. Following reverse transcription with random hexamers, we designed qPCR primers to amplify known solvent exposed regions of the abundant 18S rRNA. Consistent with the AquIRE results for total RNA damage, the levels of 18S cDNA were dramatically reduced by oxaliplatin treatment (**Figure 2E**). This reduction was not due to reduced 18S rRNA levels, which remained intact (**Supplemental Figure 2A**). Further corroborating the specificity of RNA damage caused by oxaliplatin, the reverse transcription stalling method showed that oxaliplatin, but not cisplatin or carboplatin, reduced the 18S rRNA signal in the cDNA stalling technique (**Figure 2F**).

Temozolomide is used for the treatment of glioblastoma due to its ability to cross the blood-brain-barrier. Within the body, temozolomide activation results in a highly reactive methyldiazonium ion that directly methylates biomolecules^24^. In DNA this occurs primarily at the N7-position of guanosines (m7G) with very little evidence for C5-position methylation of cytosine (m5C) or the N6-position of adenosine (m6A)^25^. However, whether and in what proportion these positions are methylated in RNA is unknown. Thus, using antibodies specific to m7G, m6A and m5C we adapted our AquIRE platform to detect temozolomide induced RNA damage (**Figure 2G-H**). Using RNA extracted from A172 glioblastoma cells with and without treatment with temozolomide, we saw induction of each of these methylation events with remarkable temporal specificity (**Figure 2I**). m6A occurs rapidly then drops sharply, m5C occurs rapidly but is retained and m7G is the latest to occur and is transient (**Figure 2I**). Methylation of RNA also impairs reverse transcription, allowing us to read out temozolomide-induced RNA damage using the cDNA stalling assay. qPCR for the 18S rRNA shows rapid and sustained RNA damage, illustrated by a reduction in signal (**Figure 2J**). This is consistent with the methylation changes seen by AquIRE, where at each timepoint at least one of the analysed methylations is elevated compared to control.

### AquIRE detects endogenously written RNA modifications

We have shown that AquIRE can detect covalent RNA damage caused by three different chemotherapies. The chemical modifications of RNA directed by writer enzymes expand the coding and functional capability of RNA, being termed the epitransciprome^26^. We expected that our AquIRE methodology would detect biological differences in the levels of these endogenous changes in RNA. We tested this using m6A and pseudouridine.

Using a commercially available m6A antibody (**Supplemental Figure 3A**), we found that the AquIRE platform gave exceptional sensitivity over a range of m6A levels (**Supplemental Figure 3B**). Using this sensitivity, we asked whether consistent differences in m6A between biological samples could be detected, doing this for total RNA extracted from 5 different colorectal cancer cell lines (**Supplemental Figure 3C**) and RNA from 4 tissues from 3 mice (**Supplemental Figure 3D**). Furthermore, we were able to recapitulate the modulation of m6A levels associated with the switch from maternal to zygotic transcription in *Drosophila* embryos (**Supplemental Figure 3E**)^27^. Finally, we saw a reduction in m6A in both total and polyA+ RNA following inhibition of the m6A writer enzyme METTL3 using the STM2457 inhibitor (**Supplemental Figure 3F**). Omitting the *in vitro* polyA tailing step from our protocol allowed analysis of endogenous polyA RNA, where an even greater reduction was seen (**Supplemental Figure 3F**).

To test pseudouridylation we used a commercially available antibody in our AquIRE protocol (**Supplemental Figure 3G**), finding a highly sensitive correlation of signal to pseudouridine content (**Supplemental Figure 3H**). Using the same panel of colorectal cancer cell lines and 4 tissues from 3 mice we were able to measure relative pseudouridine levels with sufficient accuracy to infer significant differences. (**Supplemental Figure 3I-J**). Finally, 5FU incorporation into RNA has previously been shown to reduce pseudouridine levels, due to the inability of pseudouridylating enzymes to chemically modify a fluorinated substrate^28^. An AquIRE analysis of RNA extracted from two colorectal cancer cell lines treated with 5FU supports these previous findings (**Supplemental Figure 3L**).

### AquIRE can detect specific and global RNA:protein crosslinks

RNAs are bound by RNA-binding proteins (RBPs), which change localisation, conformations or functions on transcripts. We reasoned that our antibody-based AquIRE method could be modified to detect proteins covalently bound to RNA to better understand RBP function (**Figure 3A**). We first used UV crosslinking to produce RNA:protein crosslinks *in situ* in HCT116 cells and confirmed that AquIRE can detect these crosslinks for two canonical RNA binding proteins, fibrillarin and NPM1 (**Figure 3B**). Next, we used a protein-binding dye, called NanoOrange, that fluoresces only in the presence of protein in the AquIRE protocol to quantify global protein crosslinking to RNA (**Figure 3C**). For both the specific and global protein approaches we observe a significant increase in crosslinking following UV irradiation (**Figures 3B-C**). To confirm that this signal was due to protein binding we carried out an on-bead protein digest with Proteinase K prior to detection of fibrillarin and NPM1 (**Figure 3D-E and Supplemental Fig 4A**). Proteinase K significantly reduced the signal detected after UV crosslinking for fibrillarin, with a trend towards a reduction for NPM1. Thus, AquIRE can readily detect covalent RNA:protein crosslinks that are retained through the RNA isolation and the analysis process.

**Figure 3:**
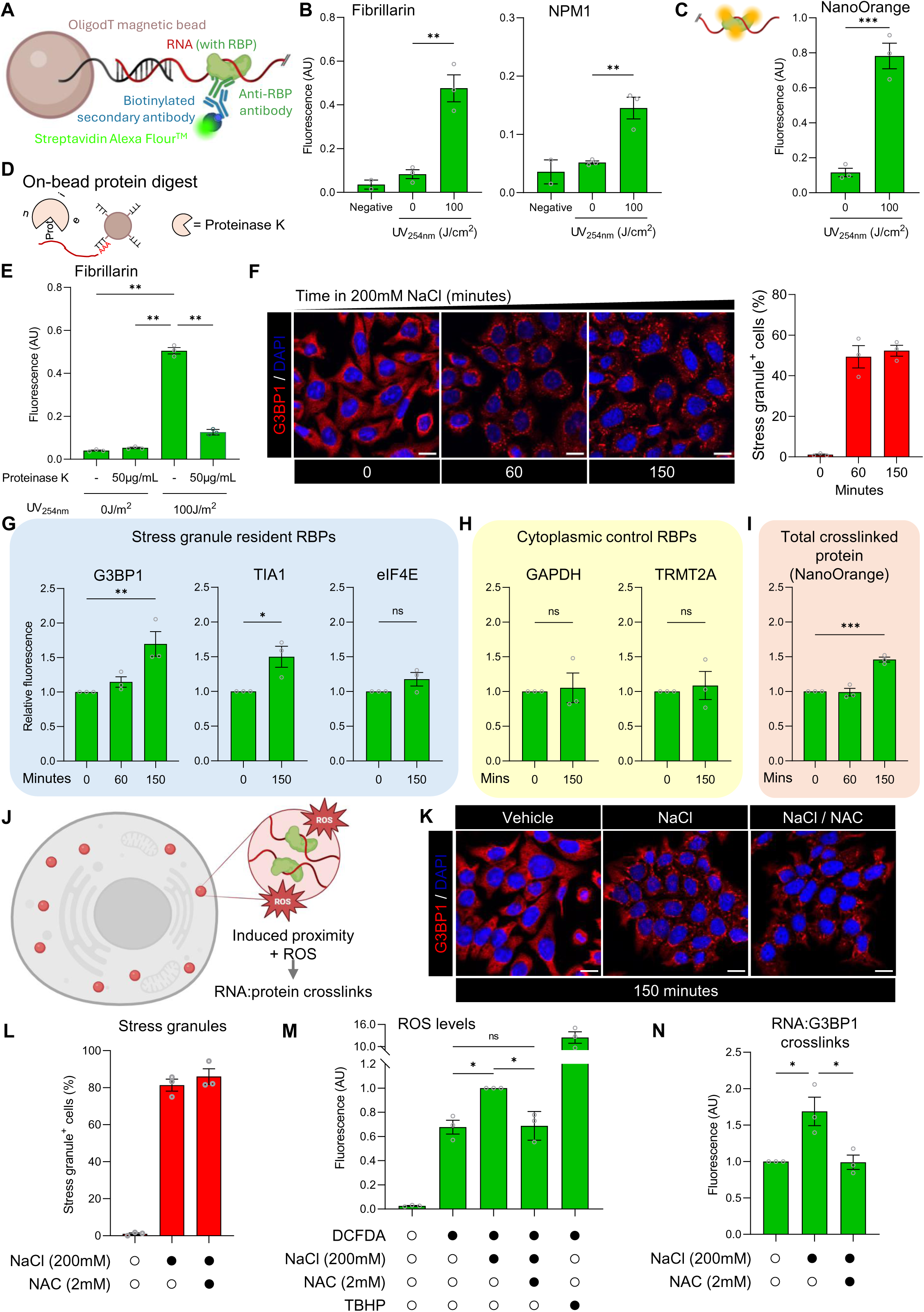
AquIRE detects proximity-driven RBP crosslinking to RNA. A) Schematic of the AquIRE detection approach to determine covalent RNA binding protein (RBP) crosslinking to RNA. B) HCT116 cells were crosslinked with UV at 254nm at the shown intensities directly prior to lysis and RNA extraction. Fibrillarin and NPM1 crosslinking to total RNA was detected by AquIRE compared to the fluorescence of the elution buffer (water) which acts as the experimental negative control. Values are plotted as raw fluorescence values. Bars represent the mean of 3 biological replicates ±SEM. Significance was determined by unpaired t test. C) Total RNA from HCT116 cells extracted after the indicated UV crosslinking protocols was analysed for NanoOrange binding. This version of AquIRE allows total protein crosslinking to be detected. Significance was determined by unpaired t test. D) Schematic of the additional step for AquIRE to include an on-bead protein digest using proteinase K. This is added after bead immobilisation but before primary antibody incubation. E) RNA as in B was analysed with the additional protein digest step or a parallel incubation without enzyme. Detection of fibrillarin crosslinking to RNA was performed by AquIRE. Values shown are the raw fluorescence reads from 3 biological replicates ±SEM. Statistical significance compared to column 3 was determined by ANOVA using Šídák multiple comparison testing. F) Left, HCT116 cells were treated with 200mM salt for the indicated times and G3BP1 localisation determined by immunofluorescence in parallel to DAPI staining. Scale bar 50µm.Right, quantification of the percentage of cells exhibiting at least 3 stress granules, defined as being a cytoplasmic G3BP1 foci. Data represent the average of 3 biological replicates ±SEM. G) HCT116 cells were treated as in F and RNA extracted. AquIRE was used to detect three stress granule resident proteins in the absence of UV induced crosslinking. The mean fluorescence is shown relative to no salt treatment for 3 biological replicates ±SEM. For G3BP1, statistical comparison to the 0 timepoint used an ANOVA with Šídák multiple comparison testing. For TIA1 and eIF4E, unpaired t tests were used. H) The crosslinking of two cytoplasmic proteins was analysed from the same RNA as analysed in G. Data are the mean fluorescence presented relative to 0 minutes salt for three biological replicates ±SEM. Significance was analysed by unpaired t test. I) NanoOrange AquIRE was used on the same RNA as in G and H. Mean fluorescence is shown relative to 0 minutes for 3 biological replicates ±SEM. Statistical comparison to the 0 timepoint was performed using an ANOVA with Šídák multiple comparison testing. J) Representation of a cell with cytoplasmic stress granules in red. Zoomed image shows the proximity of protein and RNA in these granules, as well as the presence of ROS. K) HCT116 cells were analysed as in F for after 150 minute exposure to 200mM NaCl alone or in combination with 2mM NAC. Scale bar 50µm. L) Stress granule positive cell percentages as in F, plotted as the average of 3 technical replicates ±SEM. M) HCT116 cells were pretreated with the DCFDA detection reagent then NaCl, NAC or the positive control TBHP for 150 minutes. Mean fluorescence reads, proportional to ROS levels, are plotted for 3 biological replicates relative to NaCl treated cells set to 1. Significance was tested by ANOVA with Šídák multiple comparison testing, not including columns 1 or 5, with significant values plotted between the three experimental groups of interest. N) HCT116 cells were treated with NaCl and/or NAC as in K, then isolated total RNA analysed by AquIRE for G3BP1 crosslinking. The mean fluorescence from three biological replicates is plotted ±SEM relative to RNA from untreated cells. Statistical comparison was performed using an ANOVA with Šídák multiple comparison testing. Only significant differences are shown. * *P*<0.05, ** *P*<0.01, *** *P*<0.001

To test the sensitivity and applicability of this method we used a situation where RBP binding is dynamically changed. To do this, we induced the liquid-liquid phase separation (LLPS) of stress granules through incubation of cells with 200mM NaCl (**Figure 3F**)^29^. Stress granules forming within 60 minutes of incubation in hyperosmotic conditions were imaged as G3BP1^+^ cytoplasmic foci (**Figure 3F**). In the absence of stress, G3BP1 is distributed diffusely throughout the cytoplasm, while upon stress biomolecular condensates phase-separate. These contain RNAs and RBPs, such as G3BP1, which are associated by multivalent weak interactions. We therefore asked whether AquIRE could detect these dynamic changes in the proximity of RNA and proteins for the canonical stress granule marker G3BP1. Surprisingly, we found that G3BP1 crosslinking to RNA was unchanged by stress granule formation following UV exposure, showing consistently high RNA linkage (**Supplemental Figure 4B**). However, in the absence of UV crosslinking we noticed a consistent increase in G3BP1 crosslinked to RNA after 150 minutes of hyperosmotic treatments (**Figure 3G and Supplemental Figure 4B**). To investigate this further we analysed 4 additional cytoplasmic RBPs, TIA1 and eIF4E that are present within stress granules and GAPDH and TRMT2A that are not. TIA1 showed a similar increase in RNA crosslinking after 150 minutes of salt treatment to G3BP1, while eIF4E showed a trend towards an increase (**Figure 3G**). In contrast, neither GAPDH nor TRMT2A crosslinked with RNA to a greater extent after salt treatment showing that crosslinking showed specificty for stress granule-resident proteins (**Figure 3H**). Total protein crosslinking to RNA, detected using NanoOrange, showed a significant increase after 150 minutes of salt treatment (**Figure 3I**). These data suggest that the formation of stress granules drives the covalent crosslinking of proteins to RNA because of their induced proximity (**Figure 3J**).

To preclude the possibility that non-crosslinked proteins could copurify with isolated RNA through guanidine:ethanol precipitation and then remain associated with RNA through the 2% SDS washes at the start of the AquIRE protocol, we performed an analysis using recombinant protein. Recombinant G3BP1 was preincubated with RNA from either vehicle or salt-treated cells then processed through the AquIRE protocol, using G3BP1 antibodies (**Supplemental Figure 4C**). Recombinant G3BP1 was not detectable by this method, indicating that the protein detected on RNA extracted from cells is covalently to RNA bound *in situ* only upon stress granule formation (**Supplemental Figure 4D**).

We hypothesised that RNA:protein crosslinks of the type observed with G3BP1 form due to proximity as well as the presence of intracellular ROS. To test this, we induced stress granule formation using 200mM salt in the absence or presence of the ROS scavenger N-acetyl cysteine (NAC). NAC treatment did not alter the formation of stress granules, either on its own or in combination with salt (**Figure 3K-L and Supplemental Figure 4E**), but did effectively reduce ROS levels (**Figure 3M**). When ROS levels are restrained by NAC, the RNA:G3BP1 crosslinks measured by AquIRE were restored to the amount seen in the absence of stress granule formation (**Figure 3N**). Thus, the sensitivity of AquIRE allows us to show that the formation of stress granules involving LLPS drives the ROS-dependent covalent crosslinking of RNAs and proteins.

### 5FU within RNA traps and inhibits specific proteins

We previously confirmed that 5FU in RNA cannot be converted to pseudouridine^28^ (**Supplemental Figure 3L**). Concomitant with the inability to convert 5FU to pseudouridine, the writers for this modification become covalently linked to RNA when attempting to convert it to pseudouridine (**Figure 4A**)^28^. Indeed, this mechanism is conserved in other 5-position pyrimidine modifying enzymes, such as writers of dihydrouridine or 5-methyluridine (**Figure 4A**)^30,31^. These previous reports have used high doses of 5FU or its metabolites in yeast or non-clinically relevant human cells as chemical reporters to identify these 5-position pyrimidine modifiers. However, whether 5FU induces RNA:protein crosslinks in clinically relevant models was unknown. To test this, we treated colorectal cancer cell lines with 5FU and used AquIRE to detect RNA:protein crosslinking of specific writers for pseudouridine (DKC) dihydrouridine (DUS3L), and 5-methyluridine (TRMT2A). Each of these enzymes became covalently linked to RNA following 5FU treatment in HCT116 cells (**Figure 4B**), while DUS3L and TRMT2A (but not DKC) also crosslinked to RNA in a second CRC cell line, DLD1 (**Supplemental Figure 5A**). The 5FU-dependent conjugation of each of these proteins to RNA was similar to the conjugation seen following UV irradiation (**Supplemental Figure 5B**).

**Figure 4:**
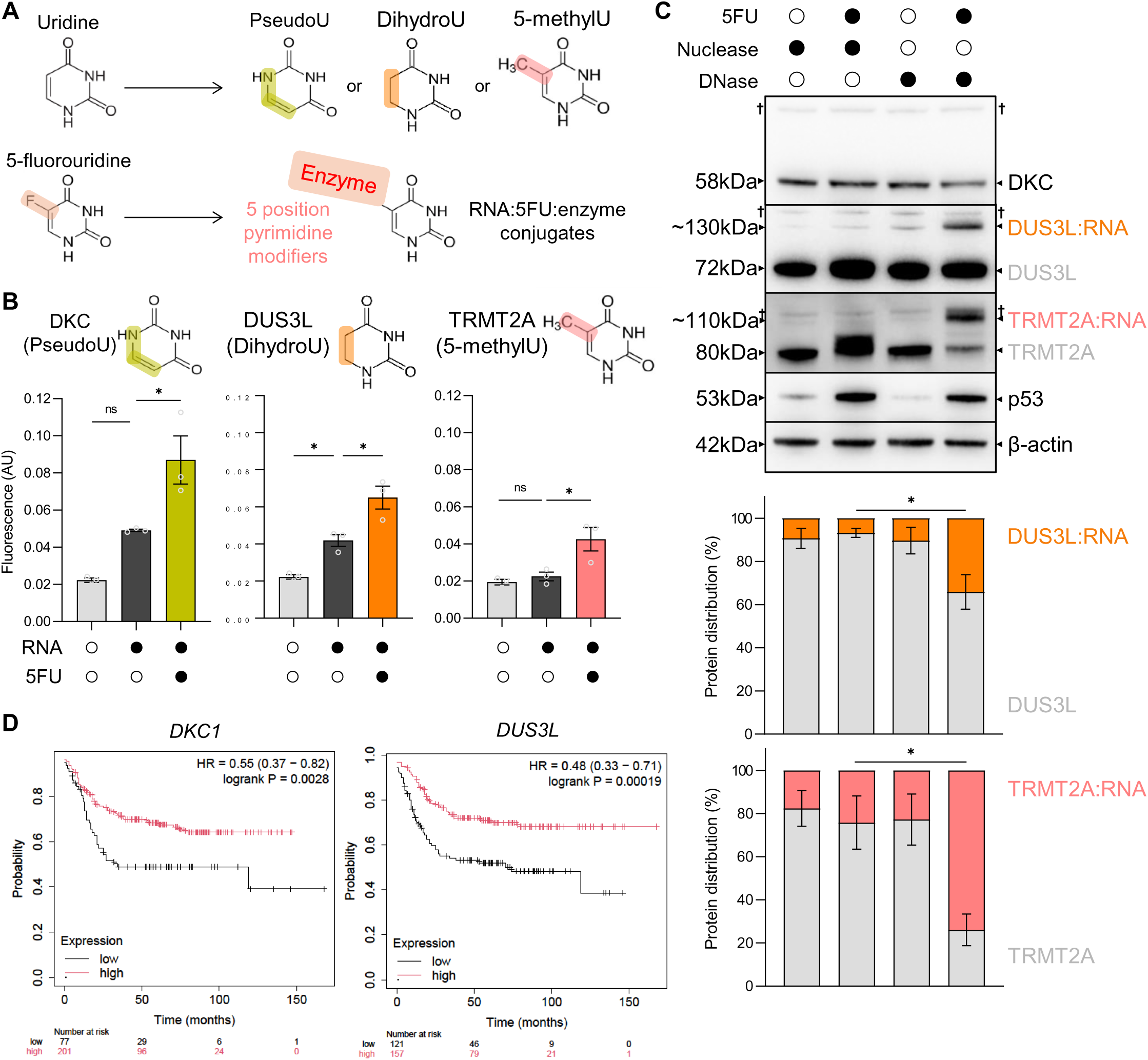
5FU crosslinks specific proteins to RNA. A) Top, representation of 5-position modifying enzyme activity upon uridine bases. Highlighted areas denote the difference in the modified base compared to canonical uridine. Below, indication of the modified base product of 5FU, highlighting the location of the divergent fluorine. 5FU negates conversion to the modified uridine bases shown above, instead covalently trapping the modifying enzymes. B) HCT116 cells were treated with 10µM 5FU for 72 hours, total RNA extracted and crosslinking of three 5-position pyrimidine modifying enzymes analysed by AquIRE. The top images show the modifying enzyme and the name and structure of the modification. Below, the graphs plot the mean fluorescence values for enzyme crosslinking from 3 biological replicates ±SEM. Grey bars indicate the fluorescence from the water elution as a negative control. Statistical comparisons were calculated against the vehicle treatment (middle column) using an ANOVA with Šídák multiple comparison testing. C) Top, HCT116 cells were treated with 2.5µM for 72 hours and protein extracted in the presence of nuclease or DNase. After digest, lysates were separated by SDS-PAGE and analysed by western blotting for expression of 5-position modifying enzymes. P53 expression serves as a control for 5FU efficacy and β-actin as a loading control. † indicates non-specific bands. Blots are representative of n=3 biological replicates. Quantification of these replicates are represented in the graphs below, showing the relative amount of DUS3L and TRMT2A present as protein or as protein crosslinked to RNA. Significance was tested using a 2-way ANOVA with Šídák multiple comparison testing. D) Kaplan-Meier plots produced using kmplot.com show the effects of tumour expression of *DKC1* or *DUS3L* on colon cancer patient survival. For *DKC1* data are plotted for 278 adjuvant therapy treated colon cancer patients split into low in black (n=77) and high in red (n=201). For *DUS3L* the numbers were 121 for low expression and 157 for high expression. Numbers below the graph indicate the number of surviving patients at each time. Marks on the graph indicate censored patients. The hazard ration (HR) and exact log rank *P* value are shown for each transcript. * *P*<0.05

To confirm these RNA:protein crosslinks we used western blotting to analyse protein lysates following selective digestion with nuclease or DNase. Nuclease digests both DNA and RNA while DNase leaves RNA intact. Depending on the treatment, we observe two bands, the protein band and, migrating more slowly, an RNA:protein band. Nuclease digestion resulted in the majority of the signal for DKC, DUS3L and TRMT2A being in the protein band, although 5FU treatment slowed the migration of this band for TRMT2A (**Figure 4C**). Detection of p53 induction was used as a control for 5FU treatment. In contrast, DNase digest after 5FU treatment reduced the abundance of the protein band for the three proteins and gave a clear induction of an RNA:protein band for DUS3L and TRMT2A (**Figure 4C**). These data confirm the covalent crosslinking of RNAs to specific 5-position pyrimidine modifying enzymes, which we observe at two separate timepoints (**Supplemental Figure 5C**).

Next, we analysed the importance of these enzymes in the clinical response to 5FU by assessing the impact of expression of *DKC1*, *TRMT2A* and *DUS3L* on patient outcome. Using publicly available data for adjuvant treated colon cancer patient survival we found that tumours with high expression of either *DKC1* or *DUS3L* had significantly longer survival compared to patients with tumours with low expression (**Figure 4D and Supplemental Table 1**). Although *TRMT2A* expression made no significant impact (**Supplemental Figure 5D and Supplemental Table 1**), high DKC1 expression more than doubled and high *DUS3L* nearly tripled survival (**Figure 4D**). Therefore 5FU, ubiquitously used for colon cancer adjuvant therapy, appears more cytotoxic to tumours expressing higher levels of DKC1 and DUS3L, proteins that it covalently conjugates to RNA.

### Oxaliplatin is a bifunctional agent that conjugates RNAs to proteins

We next asked whether other chemotherapies are capable of RNA:protein crosslinking, focusing on oxaliplatin. Once activated, the Pt(II) atom in platinating agents, such as oxaliplatin, covalently binds nucleophilic positions on biomolecules^32^. The Pt(II) atom has two coordinate positions through which it can form covalent bonds, allowing bifunctional conjugation. For example, cisplatin can crosslink different DNA strands or drive the formation of DNA:protein crosslinks^33^. Given that oxaliplatin is able to damage RNA we asked whether it drives bifunctional RNA:protein crosslink formation. This would create similar covalent complexes as outlined in **Figure 4** for 5FU, but via a distinct transcription-independent mechanism. Because oxaliplatin shows tropism for nucleoli^35,40^, we first asked whether nucleolar proteins are covalently bound to RNA following oxaliplatin treatment. We found that significantly more fibrillarin, DKC and RPS6 become covalently bound to RNA upon high doses of oxaliplatin (**Figure 5A**), which is conserved for fibrillarin and RPS6 after a clinically achievable 24-hour treatment (**Supplemental Figure 6A**). The AquIRE signal was reduced by on-bead proteinase K digestion, confirming that the signal is due to covalent protein conjugation with RNA (**Supplemental Figure 6B**).

**Figure 5:**
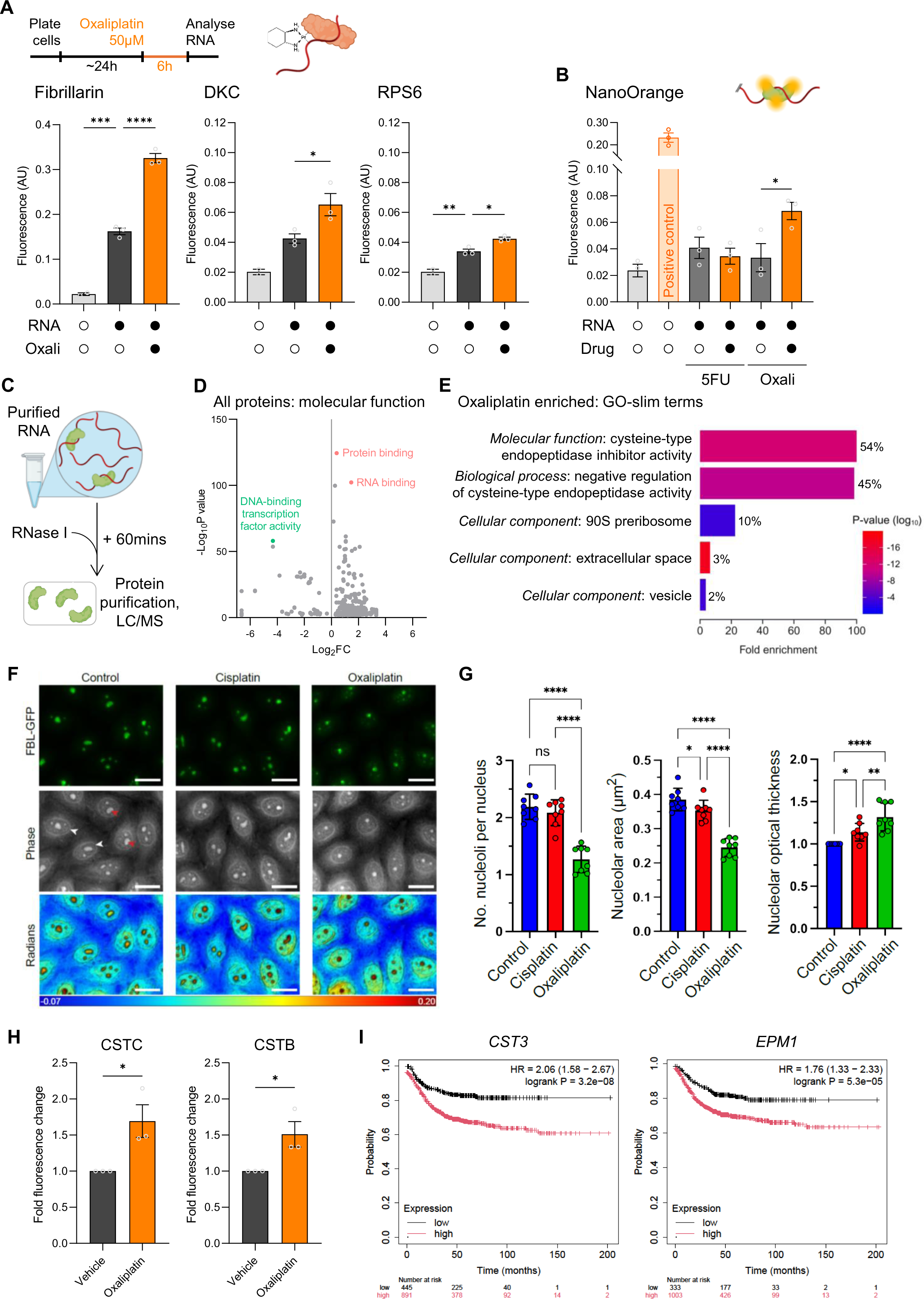
Oxaliplatin crosslinks specific proteins to RNA. A) Top, timeline of oxaliplatin treatment of HCT116 cells and pictorial representation of oxaliplatin crosslinking RNA to protein. Bottom, RNA from this analysis was analysed for crosslinking of 3 different proteins. Graphs show the raw fluorescence values for RNA from cells with and without oxaliplatin, compared to the elution buffer, water, in column 1. Statistical comparison was performed using an ANOVA with Dunnett multiple comparison testing. B) RNA was extracted from HCT116 cells treated with 10µM 5FU or vehicle for 72 hours or 50µM oxaliplatin for 6 hours and analysed by NanoOrange AquIRE in biological triplicate. The first two columns are the negative control water elution and positive control, neither with RNA. These columns plot the mean of technical replicates ±SEM. Significance is shown for oxaliplatin compared to its vehicle, which was calculated using an unpaired t test. C) Schematic of experimental approach to identify proteins crosslinked to RNA by digesting RNA and performing LC/MS proteomics. D) Volcano plot showing the molecular function GO terms for all protein identified by LC/MS. Selected GO terms are highlighted with their names. E) The 98 proteins enriched in oxaliplatin-treated RNA samples were analysed for GO-slim term enrichment. Terms are listed with their class in italics then full-term name. Bars plot the fold enrichment of proteins in our list compared to expected, while the colour of each bar corresponds to each term’s *P* value. Numbers next to each bar define the percentage of proteins in each term that are present in our dataset. F) HeLa-FBL-GFP cells treated or not with cisplatin or oxaliplatin (10 µM, 4 h) were imaged by fluorescence-DHM correlative microscopy. The GFP (green) and DHM (phase, greyscale) channels are shown. The change in optical path length (radians) is represented in pseudo-colouring. The phase shifts are expressed in radians (from-0.07 to 0.20). Scale bar, 10µm. G) Graphs depicting the mean number of nucleoli per cell nucleus, mean area of individual nucleoli (µm^2^) and mean nucleolar optical thickness (fold change) from 8 biological replicates ± standard deviation. H) RNA extracted from cells treated with 50µM oxaliplatin for 6 hours or vehicle was analysed for crosslinked CST3 or CSTB using AquIRE. Data are from 3 biological replicates plotted as the fold change in signal from the vehicle ±SEM. Significance was tested by unpaired t test. I) Kaplan-Meier plots produced using kmplot.com stratifying the tumour expression of *CST3* and *EPM1* (the gene encoding CSTB) on colon cancer patient survival. For *CST3* data are plotted for 1336 colon cancer patients split into low in black (n=445) and high in red (n=891). For *EPM1* the numbers were 333 for low expression and 1003 for high expression. Numbers below the graph indicate the surviving patients at each time. Marks on the graph indicate censored patients. The hazard ration (HR) and exact log rank *P* value are shown for both transcripts. * *P*<0.05, ** *P*<0.01, *** *P*<0.001, **** *P*<0.001

Given the striking concordance for RNA:protein crosslink formation for both 5FU and oxaliplatin, we leveraged NanoOrange global protein AquIRE to compare the conjugation activity of both drugs. RNA:protein crosslinking by 5FU was below the detection threshold of the method, but the protein conjugation caused by oxaliplatin was easily detected (**Figure 5B**). To explore this further, we performed an unbiased analysis to determine the proteins that are covalently linked to RNA following oxaliplatin treatment. Column-purified RNA from vehicle or oxaliplatin treated HCT116 cells was incubated with RNase I, followed by protein precipitation and LC/MS (**Figure 5C**). We detected a total of 1983 proteins in at least one of the six samples analysed (Supplemental Table 2). Consistent with the experimental design, Gene Ontology analysis of the molecular function of these proteins revealed an enrichment for proteins known to participate in RNA or protein binding, and a lack of proteins that participate in DNA-binding and transcription (**Figure 5D**).

A total of 98 proteins that exhibited significantly increased covalent binding to RNA following oxaliplatin treatment. Gene Ontology analysis revealed that, notably, 10% of the annotated ‘90S pre-ribosome’ proteins significantly crosslinked with RNA following oxaliplatin treatment (**Figure 5E**). The 90S pre-ribosome a multi-megadalton ribonucleoprotein particle complex involved in the earliest stages of ribosomal subunit assembly on nascent rRNA within the nucleolus. This aligns with previous findings that oxaliplatin suppresses nucleolar function^34,35^. Our data further supports this by providing direct evidence that oxaliplatin-induced RNA:protein crosslinks form within nucleoli. The pre-rRNA species present in the 90S pre-ribosome consist of 40.7% guanosine (**Supplemental Figure 6C**), and the N7 position of guanosine is the most reactive nucleotide position to oxaliplatin^36^. Thus, the nucleolar preference for oxaliplatin may simply reflect the high concentration of its preferred molecular target there – guanosine. Within nucleoli some oxaliplatin molecules form bifunctional crosslinks between the guanosine-rich RNA and 90S-resident proteins. To further investigate this interaction, we performed the cDNA stalling assay for the pre-rRNA species present within the 90S pre-ribosome, finding extensive RNA damage at lower concentrations than the 18S rRNA analysis performed previously (**Supplemental Figure 6D and Figures 2E-F**). This confirms the tropism of oxaliplatin for RNAs found in the 90S pre-ribosome and is a clear indication of oxaliplatin having transcript specificity, likely based simply on guanosine content.

The nucleolus is a biomolecular condensate, formed by LLPS and other phase transitions^37^, whose material state impacts its capacity to make ribosomes^38,39^. The extensive RNA:protein crosslinks formed by oxaliplatin within the 90S pre-ribosome will likely have a profound effect on the normal LLPS behaviour of the nucleolus, consistent with previous reports^35^. In contrast to oxaliplatin, cisplatin does not have nucleolar-specific activity^40^. To test this further we asked whether cisplatin could crosslink RNA and protein, observing no detectable conjugation of fibrillarin or DKC (**Supplemental Figure 6E**). With this knowledge, we applied the recently developed digital holographic microscopy (DHM) technique which is based on optical path length (OPL) variation to assess the material state of the nucleolus^38^. We treated HeLa cells with the same concentration of cisplatin or oxaliplatin (10µM) for 4 hours and imaged them by DHM. As DHM captures OPL variation, it does not require any staining of the samples to visualize refringent internal cell structures. Typically, the cell nucleus’ contour can easily be detected by DHM (see white arrowheads in the phase channel of **Figure 5F**), as do the prominent masses inside corresponding to nucleoli (red arrowheads). The presence of nucleoli was confirmed by detection of GFP-tagged fibrillarin, stably expressed in these cells (**Figure 5F**).

Five readouts were inspected: the number of nucleoli per cell nucleus, mean area of individual nucleoli, circularity, roundness, and nucleolar optical thickness (**Figure 6G and Supplemental Table 3**). Visual inspection of the fluorescence and phase (DHM) channels revealed distinct behaviours upon exposure to the two drugs: in cells treated with oxaliplatin, the nucleoli were less numerous, smaller and rounder (**Figure 5F**). This contrasts with cells treated with cisplatin in which nucleoli appeared more like those of control cells. These observations were confirmed by quantification (**Figure 5G and Supplemental Figure 6F**). The nucleolar optical thickness was computed and found to increase by 30% upon treatment with oxaliplatin (**Figure 5G**); cisplatin treatment also led to an increase in nucleolar thickness, but never to this extent. Thus, oxaliplatin causes the crosslinking of RNAs to nucleolar proteins, concomitantly disrupting the number, size, roundness, and material state of nucleoli. In contrast, cisplatin neither crosslinks proteins to RNA nor causes dramatic changes to the morphological features of physical properties of the nucleolus.

**Figure 6:**
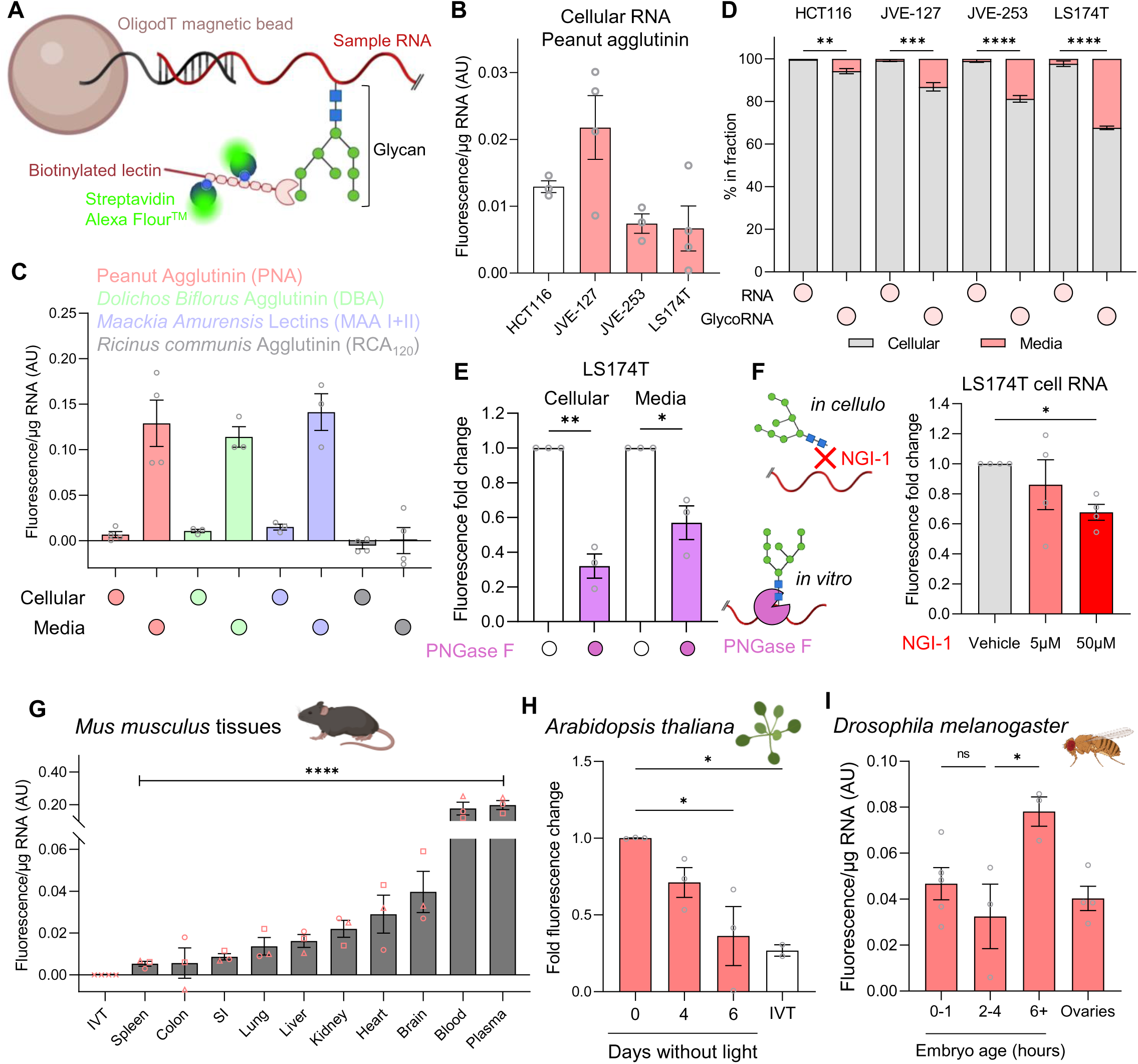
AquIRE detects glycoRNAs across the domains of life. A) Schematic of the AquIRE methodology to detect glycoRNAs using biotinylated lectins. B) RNA extracted from four colorectal cancer cell lines was analysed for glycoRNA levels using the Peanut agglutinin (PNA) lectin. Values are plotted as the fluorescence per µg of RNA from 3 biological replicated ±SEM after normalisation against *in vitro* transcribed (IVT) RNA. HCT116 cells (white bar) are non-mucinous in origin, while the other cells lines (red bars) are mucinous. C) RNA from LS174T cells and growth media was isolated and analysed for glycoRNA expression using 4 different lectins. Values are presented as the fluorescence per µg of RNA normalised against IVT RNA from at least 3 biological replicates. D) RNA was extracted and quantified from four different cell lines and their growth media, then the distribution of RNA plotted for a 3 biological replicates (left bar per cell line). The glycoRNA content of the same RNA samples was quantified using PNA and the distribution of signal normalised against RNA content and plotted as a percentage. Significance was determined independently for each cell line by 2-way ANOVA with Šídák multiple comparison testing. E) RNA from LS174T cells or growth media was digested with PNGase F to remove N-glycans, followed by AquIRE using PNA to detect glycoRNAs. Graphs show the fold change in fluorescence after normalisation against IVT RNA for 3 biological replicates ±SEM. Significance was tested by paired t test. F) LS174T cells were treated with the N-glycosylation inhibitor NGI-1 for 24 hours at the indicated concentrations and RNA isolated from cells. PNA glycoRNA levels are expressed as the fold change in fluorescence normalised against IVT RNA compared to vehicle treatment. Significance compared to vehicle was tested using an ANOVA with Šídák multiple comparison testing. G) RNA was extracted from the indicated tissues from 3 different mice (each a different symbol) and analysed by AquIRE from PNA binding to glycoRNA. Graph shows the mean fluorescence per µg of RNA for each tissue, which was normalised against IVT RNA set to 0 (also shown). Significance was determined by ANOVA analysis. H) RNA was extracted from *Arabidopsis thaliana* Col-0 ecotype leaves that had been grown in the absence of light for the indicated times. PNA AquIRE detected glycoRNA content in biological triplicate of these RNA samples compared to an equivalent mass of IVT RNA. The fold change in raw fluorescence reads is plotted compared to the 0 days timepoint. Significance was determined by ANOVA with Šídák multiple comparison testing. I) RNA was extracted from *Drosophila* (*y*^1^ *w*^67c^^23^) embryos at the indicated timepoints or adult ovary tissue and glycoRNA content determined by AquIRE. Graph plots the mean fluorescence per µg for each sample from at least 3 biological replicates. Within the embryo samples, significance compared to 2-4 hour timepoint was tested using an ANOVA with Šídák multiple comparison testing. * *P*<0.05, ** *P*<0.01, *** *P*<0.001, **** *P*<0.001

Our proteomic analysis also revealed a striking enrichment of proteins with molecular functions and biological processes related to cysteine-type endopeptidase activity (**Figure 5E**). These enrichments were due to the oxaliplatin-dependent crosslinking of cystatin proteins with RNA. Of the 9 single domain cystatin proteins, 6 significantly crosslink with RNA after oxaliplatin treatment. The cystatins are small intrinsically disordered proteins that are cytosolic or extracellular inhibitors of specific cysteine-proteases. While the cystatin family have not been studied as RBPs, intrinsic disorder is common among RBPs and cystatin B (CSTB) was identified as an RBP in multiple interactome capture analyses^41^, and shows nucleolar localisation by immunofluorescence^42^. Consistently, we confirmed that CSTB, and cystatin C (CSTC) are RNA binding proteins by analysing RNA:protein crosslinks by AquIRE following UV irradiation (**Supplemental Figure 6G**). Furthermore, AquIRE confirmed RNA crosslinking of CSTB and CSTC was increased by oxaliplatin treatment (**Figures 5H**), which occurs at clinically achievable dose of oxaliplatin (**Supplemental Figure 6H**). If these cystatins directly bind to oxaliplatin and RNA we questioned whether their expression influences colorectal cancer patient survival. Indeed, we saw that high expression of either *CST3* or *EPM1* (the genes encoding CSTC and CSTB proteins, respectively) significantly correlated with reduced survival of colorectal cancer patients, either as a whole population (**Figure 5I and Supplemental Table 1**) or in patients who received adjuvant therapy (**Supplemental Figure 6I and Supplemental Table 1**). This implies that higher cystatin expression, and likely higher RNA:cystain crosslinking, limits the cytotoxicity of oxaliplatin.

### AquIRE sensitively detects endogenous glycoRNAs

GlycoRNAs are a class of heterogeneous RNAs that are covalently modified with glycan moieties and expressed on mammalian cell surfaces^43^. They have known functions in neutrophil recruitment and cell attachment^44,45^, while upon cell surfaces they enable peptide entry via interaction with specific RBPs^46^. However, simple questions regarding glycoRNA conservation among species and non-cell surface localisation remain to be answered. We reasoned that the AquIRE platform could address this without the need of orthogonal labelling reagents^43^, or covalent modification of glycoRNA moieties^47^ through the use of glycan-sensitive lectins in place of the antibody approach used thus far. Therefore, we modified the AquIRE protocol using biotin-tagged lectins (**Figure 6A**) and were able to detect a consistent and significant fluorescent signal, using RNA isolated from 4 different colorectal cancer cell lines (**Figure 6B**). We used the lectin peanut agglutinin (PNA) for this initial detection and gained similar results with two alternative lectins, *Dolichos Biflorus* agglutinin (DBA) and the *Maackia Amurensis* lectins (MAA I+II) (**Figure 6C**). This was lectin-specific as *Ricinus communis* agglutinin (RCA_120_) gave no increase in fluorescence compared to glycan-free IVT RNA. MAA II was previously shown to bind to glycoRNAs^43,44^, while this is the first time that glycoRNAs have been shown to have sugar-structures bound by either PNA or DBA. Both of these lectins bind to terminal glycans commonly found on O-glycosylated proteins across multiple species (**Supplemental Figure 7A**)^48^, while to date glycoRNAs have been shown as substrates for N-glycosylation. Supporting our observation, two studies have independently demonstrated that N-glycans in glycoRNAs contain the T antigen bound by PNA^49^ and that O-glycosylation occurs on RNA^50^.

Given the conservation in their glycan biogenesis^43^, we asked whether glycoRNAs share the role of glycoproteins in shaping the microenvironment. To address this, we analysed the RNA and glycoRNA content of the growth media from cultures of four colorectal cancer cell lines. Three of these lines, JVE-127, JVE-253 and LS174T originated from the mucinous subtype of colorectal cancer characterised by copious levels of cancer-cell derived mucus. This mucus is made of O-glycosylated mucin proteins, with elevated production of these glycoproteins retained in culture. In comparison, the HCT116 cell line is not mucinous and produces orders of magnitude less glycosylated mucins. Quantifying the glycoRNA and total RNA distribution we found that in each case a greater fraction of glycoRNAs was found in the cell-free fraction than the distribution of total RNA (**Figure 6D**). This is indicative of an active mechanism of glycoRNA delivery into the extracellular environment by these cell lines. In line with this, the fraction of extracellular glycoRNA in the mucinous cell lines are consistently higher than the fraction in the non-mucinous cell line. Indeed, in LS174T cells 32% of glycoRNAs are extracellular, compared to only 2.5% of total RNA. Furthermore, the glycoRNA fluorescence per µg of RNA is almost 20-fold higher in cell-free RNA than cellular RNA, an observation consistent for multiple lectins for LS174T cells (**Figure 6C**) and with the PNA lectin across the panel of CRC cell lines (**Figures 6B and Supplemental Figure 6B**).

Having established that AquIRE can detect unlabelled glycoRNAs from multiple sources we next sought to use this technology to confirm previous observations regarding glycoRNA conjugation and synthesis. To do so we chose the LS174T cell line, which has the highest level of RNA per mL of media (**Supplemental Figure 7C**) and detected glycoRNAs using PNA. First, we incubated purified cellular and extracellular RNA with the N-glycosylase PNGase F, which was previously shown to remove glycoconjugates from RNA^43^. In agreement, we observed a reduction in the fluorescence signal from both cellular and cell-free RNA following incubation (**Figure 6E**). In parallel, we treated cells with the oligosaccharyltransferase (OST) inhibitor NGI-1, which is known to inhibit the N-glycosylation of RNAs^43^. Again, our detection method saw a reduction in glycoRNA expression in cellular RNA following NGI-1 treatment (**Figure 6F**). However, there was no reduction in the glycoRNAs detected in the cell-free fraction (**Supplemental Figure 7D**). It is unclear whether this is due to the cell-free glycoRNAs having a longer half-life or an extended biogenesis pathway. Nevertheless, these data confirm that the glycoRNAs we detect display highly similar molecular biology to those described previously and present for the first time the observation of cell-free glycoRNAs.

### GlycoRNAs are widely expressed across the kingdoms of life

Next, we leveraged the ability to measure glycoRNAs in any cell, tissue or liquid sample to identify which species express glycoRNA. First, we sampled 9 tissues and blood plasma from wild-type mice and quantified the relative glycoRNA expression normalised to IVT RNA and expressed per µg of input RNA. GlycoRNAs could be detected in all tissues and within blood plasma, with more than an order of magnitude between the highest expressing tissue, blood, and the lowest, spleen (**Figure 6G**). This tissue-specificity likely indicates distinct functions that are yet to be revealed. The RNA recovery at the end of these AquIRE experiments varied by tissue, from over 100% down to 40% (**Supplemental Figure 7E**). The reasons for this are unclear, but there was not a positive correlation between fluorescent signal and RNA recovery (**Supplemental Figure 7F**).

To date, glycoRNAs have only been described in mammals. We therefore sought to test the conservation of these molecules by analysing their expression in diverse organisms and models of specific biology, from embryo development to senescence. Throughout, we maintain a focus on the expression of glycoRNAs in cellular and cell-free samples. First, we found that the model plant organism, *Arabidopsis*, expresses glycoRNAs, both as a seedling and in leaves (**Supplemental Figure 7G-H**). Using an *Arabidopsis* model of leaf senescence, we found that glycoRNA levels were significantly reduced during senescence and essentially absent in leaves deprived of light for 6 days (**Figure 6H**). Next, we used *Drosophila* embryos to model the levels of glycoRNAs during embryonic development and in the ovary. The transition from maternal to zygotic transcription occurs in two waves between 1 and 3 hours of development^52^ and has been correlated with changes in RNA modifications such as m6A^27^. GlycoRNAs were present at all timepoints in our analysis, as well as in adult fly ovaries (**Figure 6I**). The presence of glycoRNAs in the 0-1 hour embryos, prior to the onset of zygotic transcription, shows that they are maternally deposited. In addition, there is a significant increase in glycoRNAs in older stage embryos, showing that glycoRNAs are abundant during embryogenesis (**Figure 6I**). These fundamental observations dramatically expand the horizons of what is known about where and when glycoRNAs are expressed.

To test the species conservation of glycoRNAs further we asked if the single-celled eukaryote model, *Saccharomyces cerevisiae* expressed glycoRNAs. We found expression of glycoRNAs in both the *Saccharomyces cerevisiae* cells and in cell-free media extracts (**Supplemental Figure 7I**). Significantly more glycoRNA was detected in the growth media than total RNA, leading to the conclusion that *Saccharomyces cerevisiae* actively release glycoRNAs into their environment. In a similar analysis, we asked if the prokaryotic model organism, *Escherichia coli,* expresses glycoRNAs within cells and in its environment. We found glycoRNAs present in both sample types, with an enrichment of glycoRNAs in cell-free RNA compared to cellular RNA (**Figure 7A**). Indeed, we observe that nearly 75% of glycoRNAs are cell-free in this *E. coli* model.

**Figure 7:**
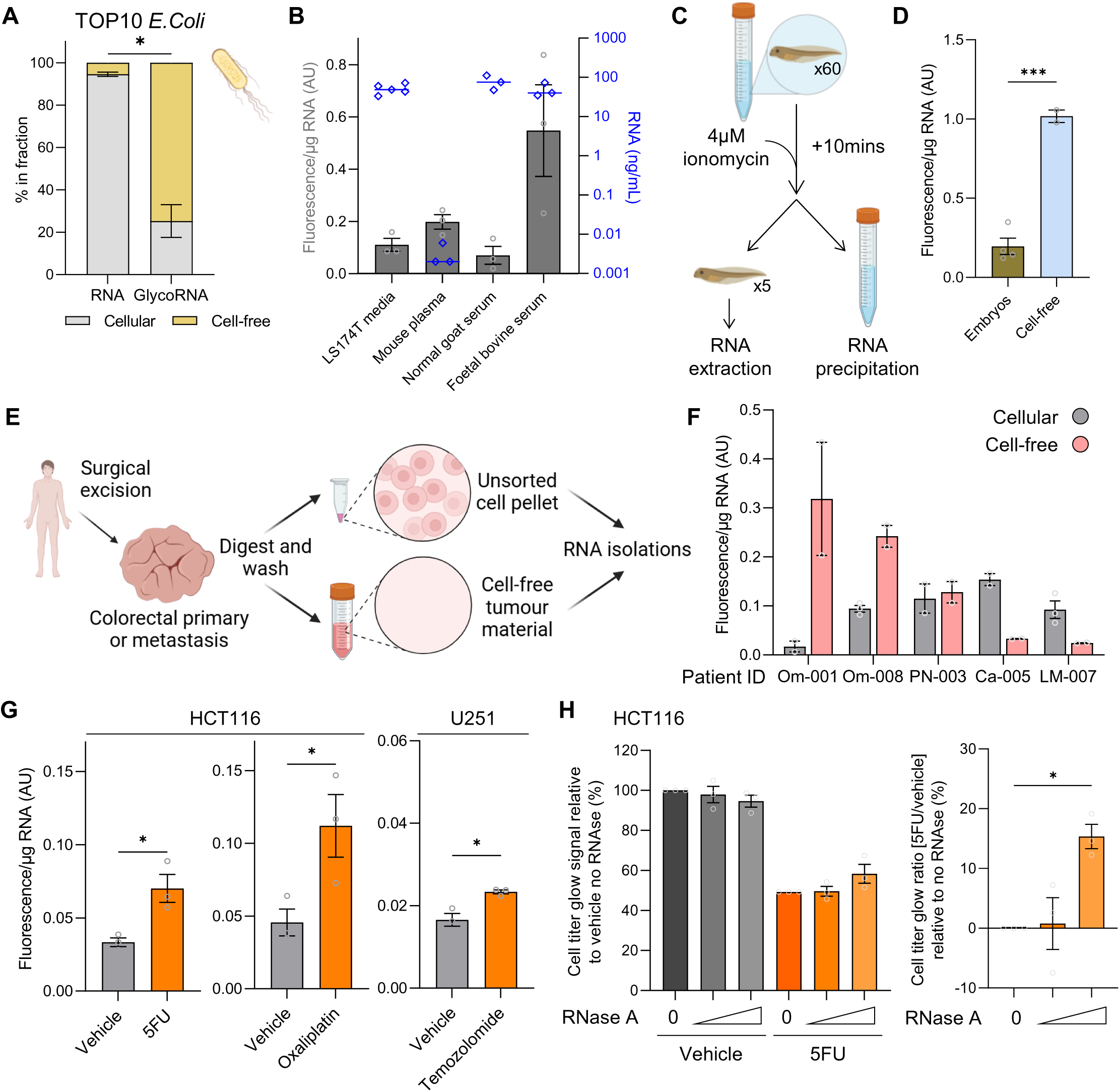
Detecting cell-free glycoRNAs and their influence on 5FU response. A) RNA from *Escherichia coli* (TOP10) cells and growth media was isolated and quantified then the distribution of RNA plotted for 4 biological replicates (left). The glycoRNA content of the same RNA samples was quantified using PNA and the distribution of signal normalised against RNA content and plotted as a percentage (right). Significance was determined independently for each cell line by 2-way ANOVA with Šídák multiple comparison testing. B) RNA was extracted from the media of LS174T cells, mouse plasma and serum from goat and cow. The grey bars plot the mean glycoRNA fluorescent signal per µg of RNA, while the blue bars plot the RNA content of each liquid sample in ng/mL. C) Ionomycin secretagogue protocol for isolation of whole organism and cell-free *Xenopus laevis* RNA. Embryos were treated for 10 minutes with ionomycin then RNA extracted from pooled whole embryos or their growth media. D) RNA samples as in C were analysed for glycoRNA content by AquIRE. Graph plots the mean fluorescence per µg of RNA from 4 pooled embryo samples and 2 cell-free samples. Significance was determined by unpaired t test. E) Schematic of colorectal tumour processing to generate unsorted cell pellets and cell-free tumour material. RNA was extracted from each fraction for analysis. F) RNA from E was analysed by AquIRE and plotted as the fluorescence per µg of RNA for 5 biologically independent tumours, showing the mean of at least two technical replicates. The x axis lists the unique tumour name. G) HCT116 colorectal cancer cells were treated with 10µM 5FU for 72 hours, 50µM oxaliplatin for 6 hours or their vehicles and U251 glioblastoma cells were treated with 2mM temozolomide or vehicle for 2 hours. GlycoRNA content was determined in RNA extracted after these treatments, with the graphs plotting the mean fluorescence per µg from 3 biological replicates per drug ±SEM. Significance was determined by unpaired t test. H) HCT116 cells were treated with 10µM 5FU for 72 hours in the presence of 100 or 200µg/mL RNase A or no enzyme. Right, the change in Cell Titer Glo viability analysis signal was plotted relative to vehicle with no enzyme set to 100%. Due to variability in the efficacy of 5FU between biological replicates (5% SEM) values after drug treatment were normalised against the average effect of 5FU in the absence of enzyme (49.3%). Right, the ratio of the cell viability signal of 5FU / vehicle for the three different enzyme conditions was plotted as a percentage change in viability compared to no enzyme set to 0. Data represent the mean of 3 biological replicates ±SEM. Significance compared to no enzyme condition was tested using an ANOVA with Šídák multiple comparison testing. * *P*<0.05, *** *P*<0.001

### GlycoRNAs are present in cell-free samples and determine chemotherapy responses

Having observed glycoRNAs present in the blood plasma of mice (**Figure 6G**) we analysed the expression of RNA and glycoRNAs in commercially available mammalian serum products from *Capra aegagrus hircus* (normal goat serum) and *Bos taurus* (foetal bovine serum). RNA was present in both cell-free liquids, consistent both with previous reports^51^ and with the similar or higher glycoRNA fluorescence values per µg of RNA seen in media conditioned by LS174T cells (**Figure 7B**). Next, we used *Xenopus tropicalis* embryos treated with the secretagogue ionomycin to detect glycoRNAs in whole embryos or their cell-free environment (**Figure 7C**). We were able to detect high expression of glycoRNAs in both samples, with the cell-free samples showing a significantly higher signal per µg of RNA than embryos (**Figure 7D**). Of note, *Xenopus* embryos showed the highest expression of glycoRNAs per µg of RNA across the >20 samples from organisms or cells, while the media in which *Xenopus* were treated with ionomycin showed the highest glycoRNA/µg RNA of any sample analysed, including >10 cell-free samples (**Supplemental Figure 7J**).

We leveraged our AquIRE methodology to detect glycoRNAs from clinical samples, focusing on the glycoRNAs expression in the tumours of colorectal cancer patients. Following surgical excision, five tumours were processed to yield a paired unsorted cell pellet and cell-free tumour material (**Figure 7E**). RNA was extracted from both sources and analysed for glycoRNA content expressed as fluorescence/µg RNA. GlycoRNAs were detected at varying levels across all 10 samples (**Figure 7F**). GlycoRNA expression displayed intra-and inter-heterogeneity across patient samples. When comparing the same sample type between different patients, glycoRNA levels differed by an order of magnitude. However, between sample types for individual patients, some displayed similar expression while others had large differences. These data clearly show that glycoRNAs are likely ubiquitously expressed in and released by tumours in patients. Our next analyses asked whether these glycoRNAs are functional or dynamic.

Linking back to our analysis of drug-induced RNA damage, we asked whether specific chemotherapies could modulate the levels of glycoRNAs in disease specific cell line models. Thus, we treated the HCT116 cell line with 5FU or oxaliplatin and observed a consistent increase in glycoRNA expression (**Figure 7G**). Similarly, treating the glioblastoma cell line U251, with temozolomide also resulted in an increase in glycoRNA expression (**Figure 7G**). RNA from HCT116 cells is damaged by either 5FU or oxaliplatin (**Figures 1-4**), while U251 RNA demonstrates m7G accumulation after temozolomide treatment (**Supplemental Figure 7K**).

Finally, to functionally test the role of glycoRNAs in the response to chemotherapy we removed glycoRNAs from cell culture models using RNase A, similar to performed previously^43–45^. RNase A digests all cell-free and cell surface RNAs from HCT116 cells, which we treated in parallel with and without 5FU. Removing RNA had little effect on HCT116 cell viability (**Figure 7H**). However, in the presence of 5FU, HCT116 cells lacking cell-free and cell surface RNAs were significantly more viable than undigested controls (**Figure 7H**). Thus, cell-free or cell surface RNA is required for the maximum cytotoxicity of 5FU in HCT116 tumour cells. This reveals a previously unknown disease modulatory mechanism, likely attributable to glycoRNAs, in determining cell responses to RNA damaging agents.

## Discussion

The cellular RNA damage responses are well documented and are beginning to be linked to normal processes and patho-physiology^3,9,10,13,16,18^. Based on previous observations, we hypothesised that common chemotherapies cause RNA damage and set out to directly measure this biology. To do so, we developed our AquIRE research platform, measuring drug induced RNA damage for 5FU, oxaliplatin and temozolomide. This allowed us to document their previously unknown temporal dynamics and multifaceted mechanisms of action. We present three examples of the many antimetabolite and alkylating compounds, prompting the question of RNA damage by these related compounds.

The long-accepted mechanism of action for 5FU is the inhibition of thymidylate synthase (TS) via the formation of a covalent complex with a 5FU metabolite^53^. Challenging this assumption, the importance of 5FU incorporation into RNA altering its function and RNA damage in determining how cells respond to 5FU was recently published^4,7^. Consistent with previous reports linking 5FU incorporation into RNA to cytotoxity from the 1980s^5,6^. In truth, 5FU administration has both effects at the same time (affecting DNA synthesis and RNA metabolism and function) and both will contribute to cytotoxicity. We show here that the crosslinking mechanism that inhibits TS is conserved for 5FU incorporated into RNA which traps 5-position pyrimidine modifying enzymes. This adds an additional layer to our understanding of how 5FU functions, with incorporation into RNA presenting opportunities to impact the epitranscriptome by trapping and inhibiting writer enzymes. Indeed a recent study demonstrates a sustained inhibition of both pseudouridine and methyluridine even after removal of 5FU^54^. Our data indicates that this is likely due to inactivation of the writer enzymes by 5FU within RNA, requiring these to be synthesised in the absence of drug to reactivate the writer pathway.

In our study, 5FU shares an RNA:protein crosslinking mechanism with oxaliplatin and an epitranscriptomic modifying mechanism with temozolomide. We see that oxaliplatin is a more effective RNA:protein crosslinker than 5FU and that temozolomide has a wider effect on the epitranscriptome. Thus, our work indicates that direct targeting of RNA or its epitranscriptome, both focuses of drug development programs of the 21^st^ century^55–57^, have been contributing to clinical benefit for decades. Learning from classic chemotherapies may identify where to apply technological advances for specific RNA targeting in the future and thereby reduce collateral effects.

Liquid-liquid phase separation has been implicated in ribosome biogenesis, mRNA synthesis and fate, chromatin biology, and protein transport^58^. Here we show that the proximity of RNAs and proteins defined by LLPS in stress granules results in RNA:protein crosslinking due to ROS. Stress granules are known to harbour ROS^59^ and function to promote cell viability under a range of stress conditions. We reveal that the condensation of biomolecules in stress granules in fact directly damages them to some extent. ROS are found throughout cells^60^, implying similar proximity-induced damage mechanism could impact other LLPS bodies. Does the benefit of proximity for increased function outweigh the risk for proximity-induced damage? Presumably this is the case, given that membraneless organelles have been selected during evolution to play essential regulatory roles in gene expression. It is also likely that these proximity-induced damaged RNAs will provide substrates for the RNA damage surveillance and processing pathways mentioned previously.

Our studies of glycoRNAs identified their expression in both the prokaryote and eukaryote domains of life and four of the seven kingdoms (bacteria, animals, plants and fungi). As such, it appears likely that glycoRNAs are expressed in all life forms. In fact, throughout our studies we found only one condition where glycoRNAs were essentially absent – within the senescent leaves of *Arabidopsis thaliana*. Our work identified common O-glycosylation moieties present on RNA, as well as confirming the previously accepted N-glycan domains and their conjugation via OST^43^. The three canonical forms of glycosylation (O, N and lipid) are conserved across the domains of life^61^. The conservation of glycoRNAs indicates that RNA does not hijack the enzymes of other biomolecules to become glycosylated but in all likelihood that these mechanisms of conjugation coevolved for RNA and protein substrates.

Our observation of the conservation of glycoRNAs was only possible due to the AquIRE methodology with its key advantages of requiring low inputs and not relying on metabolic labelling. Unlike the use of bio-orthogonal agents, our method directly interrogates what the glycans are comprised of, independent of how they are made. Furthermore, the ability to switch lectins to analyse additional moieties will allow a rich picture of glycoRNA expression to be drawn. We have shown a reciprocal functional relationship between RNA-damaging drugs and glycoRNA. We believe that AquIRE is the first method to detect glycoRNAs that is amenable to high throughput analyses. This will be important as further roles for glycoRNAs in disease emerge, placing glycoRNAs in line with the misregulation of glycoproteins and glycolipids in human pathology^62^.

## Resource availability

This study did not create any new reagents

## Acknowledgements

We thank the facilities at the CRUK Manchester Institute; the BRU Experimental Team for mouse tissues and the Molecular Biology Core for Agilent Bioanalyzer experiments. We thank the BioMS facility at the University of Manchester for proteomic analysis (RRID: SCR_020987). We thank the University of Manchester Biological Service Facility for technical support in our *Xenopus* studies. Research samples were obtained from the Manchester Cancer Research Centre (MCRC) Biobank, UK. The role of the MCRC Biobank is to distribute samples and therefore, cannot endorse studies performed or the interpretation of results. Some figures were created in BioRender. Knight, J (2025) https://BioRender.com/f11o153, https://BioRender.com/t94t009, https://BioRender.com/g55d965, https://BioRender.com/m06l035, https://BioRender.com/d61e008, https://BioRender.com/s88×264.

We are thankful for funding from the Lister Institute of Preventative Medicine (JRPK), the China Scholarship Council (ZZ, DT, JRPK), The Christie Charity (ZVK, MB, MS, JRPK). PS is funded by The Christie Charity, in partnership with NIHR Manchester Biomedical Research Centre. Research in the lab of DLJL was supported by the Belgian Fonds de la Recherche Scientifique (F.R.S./FNRS), EOS [CD-INFLADIS, 40007512], Région Wallonne (SPW EER) Win4SpinOff [RIBOGENESIS], the COST action TRANSLACORE (CA21154), the European Joint Programme on Rare Diseases (EJP-RD) RiboEurope and DBAGeneCure. Research in the lab of MA was supported by the BBSRC (BB/Y005783/1). Research in the lab of HA was supported by a Wellcome Trust Discovery Award (227415/Z/23/Z)

## Author contributions

Conceptualization: ZZ, ZVK, EH, MA, HA, DLJL, PG, PS, DT, JRPK. Formal Analysis: ZZ, ZVK, KD, XK, NS, TWY, AS, DLJL, JRPK. Investigation: ZZ, ZVK, KD, XK, NS, TWY, AS, LFB, EH, MC, CDH, JRPK. Resources: LFB, EH, TQ, KTK, MA, HA, PG, MB, MS, PS, DT. Writing – original draft: ZZ, ZVK, KTK, LFB, EH, PG, JRPK. Writing – review & editing: all authors. Supervision: NS, MA, HA, DL, MB, MS, DT, JRPK. Funding Acquisition: MA, HA, PG, DLJL, MB, MS, DT, JRPK.

## Supplemental Information

Tables: Supplemental Tables 1-4: Figures:

Supplemental Figures 1-7

## Supplemental legends

Supplemental Table 1

Survival analysis statistics produced by kmplot.com. Analyses are for colon cancer patients with the specific transcript, Affy ID and expression cut offs state. Gene Expression Omnibus accession numbers, along with the number of patients in each data set, are also listed.

Supplemental Table 2

Raw data from the oxaliplatin-RNA-crosslinked proteome. Protein accessions, names and normalised abundances are given, as well as the fold changes and adjusted P values for each protein. Full data can be found on proteomeXchange with the dataset identifier PXD06117

Supplemental Table 3

Tables outline the data represented for the digital holographic microscopy experiments. The number of nuclei analysed per condition are listed and descriptive statistics for each parameter – mean, standard deviation and P values for all comparisons are listed.

Supplemental Table 4

Details of the patients and selected tumour characteristics for the samples used in this study to identify cellular and cell-free glycoRNA expression.

Supplemental Figure 1

A) RNA from *in vitro* transcriptions (IVT) with the stated percentage of 5FUTP included in place of UTP or from HCT116 cells treated with 2.5µM 5FU for 24hours was analysed by RNA dot blot. 5FU:RNA was detected using anti-BrdU antibody with an HRP-conjugated secondary antibody. Total RNA was visualised using methylene blue. B) Pixel intensity was calculated for the IVT RNA and plotted against the known incorporation of 5FUTP. The orange line plots a simple linear regression and the dashed line is the 95% confidence interval. Details of the linear regression fit are inset into the graph. Data are from one replicate. C) The same samples as in B were analysed by AquIRE and again expressed as a linear regression. Data are from at least two technical replicates (i.e. different AquIRE assays) of the same IVT RNA sample. D) Graph plots the mean relative fluorescence intensity for 5FU incorporation into RNA at the shown timepoints. These data are from 3 biological replicates, one of which is represented in Figure 1D. Significance was tested using an ANOVA with Šídák multiple comparison testing. E) RNA recovery was determined as the amount of RNA recovered as a percentage of RNA input, following the AquIRE assay shown in Figure 1E. Data are technical triplicates from one AquIRE assay. F) Data as in D but with one of the biological replicates depicted in Figure 1G. Significance compared for all timepoints relative to 0 hours was tested using an ANOVA with Šídák multiple comparison testing. * *P*<0.05, ** *P*<0.01, *** *P*<0.001, **** *P*<0.001

Supplemental Figure 2

A) Representative TapeStation traces of RNA extracted from HCT116 cells following 24-hour treatment with the indicated does of oxaliplatin. The mean RIN score from the indicated number of biological replicates is inset into the traces ±SEM. * *P*<0.05

Supplemental Figure 3

A) Schematic of the detection of m6A using a specific antibody integrated into the AquIRE protocol. B) Different amounts of IVT RNA made with 50% m6ATP were analysed by AquIRE. Mean fluorescence reads ±SEM from 3 technical replicates were plotted against the known quantity of m6A. The purple line plots a simple linear regression and dashed lines are the 95% confidence interval. Details of the linear regression fit are inset into the graph. C) RNA was isolated from 5 different colorectal cancer cell lines and analysed for m6A content by AquIRE. Values are expressed as the mean of 3 biological replicates ±SEM relative fluorescence compared to the cell line with the lowest m6A levels (HCT116). Significance between samples was tested by ANOVA. D) AquIRE was used to determine m6A levels in RNA samples from 4 tissues from 3 mice. Each colour coded bar represents a tissue with the numbers in the annotation indicating the same animal. Values are presented as the raw fluorescence read from 1 technical replicate per tissue per animal. E) RNA was extracted from *Drosophila* (*y*^1^ *w*^67c^^23^) embryos at the indicated timepoints or adult ovary tissue then m6A levels quantified by AquIRE. The graph plots the fold fluorescence change compared to the 0-1 hour timepoints from at least 3 biological replicates. Within the embryo samples, significance compared to 2-4 hour timepoint was tested using an ANOVA with Šídák multiple comparison testing. F) RNA was extracted from HCT116 cells following treatment with STM2457 for 24h at the indicated concentrations. The graphs show AquIRE data plotted as relative fluorescence compared to vehicle treatment for total RNA, left, or polyA RNA, right. Data are from 3 biological replicates and show the mean ±SEM. For both, significance compared to vehicle was tested using an ANOVA with Šídák multiple comparison testing. G) Schematic of the detection of pseudouridine (Ψ) using a specific antibody within the AquIRE protocol. H) IVT RNA made with the indicated percentage of pseudoUTP were analysed by AquIRE. Mean fluorescence reads ±SEM from 3 technical replicates were plotted against the known quantity of Ψ. The purple line plots a simple linear regression and the dashed lines are the 95% confidence interval. Details of the linear regression fit are inset into the graph. I) RNA was isolated from 5 colorectal cancer cell lines and analysed for Ψ content by AquIRE. Values are expressed as the mean of 3 biological replicates ±SEM relative fluorescence compared to HCT116, the cell line with the lowest Ψ levels. Significance between samples was tested by ANOVA. J) AquIRE was used to determine Ψ levels in RNA samples from 4 tissues from 3 mice. Each colour coded bar represents a tissue with the numbers in the annotation indicating the same animal. Values are presented as the raw fluorescence read from 1 technical replicate per tissue per animal. K) Schematic indicating that 5FUridine, unlike uridine, cannot be converted to pseudouridine. The modified part of the Ψ base is shown in green and the hindering fluorine atom in 5FUridine in red. L) RNA was extracted from HCT116 or DLD1 cells following treatment with 5FU at 10µM for 72h. The graphs show AquIRE data plotted as relative fluorescence compared to vehicle treatment. Data are from 3 biological replicates and show the mean ±SEM. For both cell lines, significance was tested using an unpaired t test. * *P*<0.05, ** *P*<0.01, *** *P*<0.001, **** *P*<0.001

Supplemental Figure 4

A) HCT116 cells were crosslinked with UV at 254nm at the shown intensities directly prior to lysis and RNA extraction. NPM1 crosslinking to total RNA was detected by AquIRE with the additional protein digest step or a parallel incubation without enzyme. Data is plotted as raw fluorescence values. Bars represent the mean of 3 biological replicates ±SEM. Statistical significance was determined by ANOVA using Šídák multiple comparison testing. The significance between samples where the experimental variable is UV exposure are shown. B) HCT116 cells were treated with 200mM salt for the indicated times with or without UV 254nm crosslinking directly prior to lysis. G3BP1 crosslinking to RNA was subsequently detected by AquIRE. The mean fluorescence is shown for 3 biological replicates ±SEM. C) Schematic of the experimental protocol to test retention of non-covalent G3BP1 binding through the AquIRE protocol. Differing amounts of recombinant G3BP1 was preincubated with RNA samples then analysed by AquIRE for G3BP1 crosslinking. D) Relative fluorescence signals for G3BP1 crosslinking are plotted from 3 independent AquIRE assays using RNA isolated from 200mM NaCl, right, or vehicle, left, treated HCT116 cells. Mean values from each replicate are plotted ±SEM. E) In parallel to the experiment in Figure 3K, HCT116 cells were treated with 2mM NAC compared to vehicle treatment. Representative images of G3BP1 localisation as determined by immunofluorescence in parallel to DAPI staining. Scale bar 50µm. * *P*<0.05

Supplemental Figure 5

A) RNA from DLD1 colorectal cancer cells was isolated after treatment with 10µM 5FU for 72hours and analysed from protein crosslinking of the indicated enzymes. Data are the mean normalised fluorescence compared to vehicle treated for 3 biological replicates ±SEM. Significance was tested using unpaired t tests for each protein. B) HCT116 cell were UV 254nm exposed directly prior to lysis for RNA extraction and the crosslinking of 3 enzymes analysed by AquIRE. The graphs show the mean raw fluorescence reads from 3 biological replicates ±SEM. Significance was tested for each protein by unpaired student t test. C) Top, HCT116 cells were treated with 2.5µM for 48 hours and protein extracted in the presence of nuclease or DNase. After digest, lysates were separated by SDS-PAGE and analysed by western blotting for expression of specific enzymes. P53 expression serves as a control for 5FU efficacy and β-actin as a loading control. † indicates non-specific bands. Blots are representative of n=3 biological replicates. Quantification of these replicates are represented in the graphs below, showing the relative amount of DUS3L and TRMT2A present as protein or as protein crosslinked to RNA. Significance was tested using a 2-way ANOVA with Šídák multiple comparison testing. D) Kaplan-Meier plots produced using kmplot.com show the effects of tumour expression of *TRMT2A* on colon cancer patient survival. Data are plotted for 278 adjuvant therapy treated colon cancer patients split into low in black (n=152) and high in red (n=126). Numbers below the graph indicate the number of surviving patients at each time. Marks on the graph indicate censored patients. The hazard ration (HR) and exact log rank *P* value are shown for each transcript. * *P*<0.05, ** *P*<0.01, *** *P*<0.001

Supplemental Figure 6

A) Top, experimental overview of treatment regime with HCT116 cells treated with oxaliplatin at 2.5µM for 24 hours, and diagram of bifunctional oxaliplatin crosslinking of RNA and protein. Bottom, RNA from this analysis was analysed for crosslinking of 3 proteins. Graphs show the raw fluorescence values for RNA from cells with and without oxaliplatin, compared to the elution buffer, water, in column 1. Data are the average of 3 biological replicates ±SEM. Statistical comparison was performed using an ANOVA with Dunnett multiple comparison testing. B) Top, schematic of the on-bead protease digest using proteinase K. RNA was extracted from HCT116 cells after treatment with oxaliplatin at 50µM or vehicle for 6 hours. Two equal amounts of RNA were immobilised on beads, with one digested with proteinase K and the other incubated without enzyme in parallel. Following digestion, the crosslinking of fibrillarin was tested by AquIRE. The graph plots the mean raw fluorescence of 3 biological replicates and the negative control (water) ±SEM. Significance was tested using an ANOVA with Šídák multiple comparison testing. C) The nucleotide abundance for the annotated RNA sequences for the human pre-rRNAs (External Transcribed Spacer 1 (ETS1), Internal Transcribed Spacer 1 (ITS1), ITS2 and ETS2) and the polycistronic rRNAs (18S, 5.8S and 28S) were calculated and the mean value plotted ±SEM. Significance was tested using a 2-way ANOVA with Šídák multiple comparison testing. D) RNA from HCT116 cells treated with the indicated concentrations of oxaliplatin for 6 hours was reverse transcribed in a cDNA stalling assay. Graphs show the change in cDNA levels for two pre-rRNA sites in the ETS1 plotted as the mean fold change compared to vehicle treatment for 3 biological replicates ±SEM. Statistical difference compared to vehicle treatment was calculated using an ANOVA with Šídák multiple comparison testing. E) Total RNA was isolated from HCT116 cells that were treated with 50µM cisplatin for 6 hours then analysed for crosslinking of the indicted proteins. Data are the mean of two biological replicates ±SEM. The first column shows the fluorescence of the negative control, water. F) DHM images of HeLa cells as shown in Figure 5F were analysed for nucleolar circularity and nucleolar roundness. Data are plotted from 8 independent experiments and significance determined by ANOVA analysis with Holm-Šídák multiple comparison test. G) RNA from HCT116 cells was extracted after pre-lysis crosslinking with UV_254nm_ at either 0 or 100J/cm^2^. These RNAs were analysed for CST3 and CSTB covalent linking by AquIRE. Data are the mean fluorescence of 3 biological replicates ±SEM. Significance for each protein was determined by an unpaired t test. H) RNA extracted from cells treated with 2.5µM oxaliplatin for 24 hours or vehicle was analysed for crosslinked CST3 or CSTB using AquIRE. Data are from 3 biological replicates plotted as the fold change in signal from the vehicle ±SEM. Significance was tested by unpaired t test. I) Kaplan-Meier plots produced using kmplot.com stratifying the tumour expression of *CST3* and *EPM1* (the gene encoding CSTB) on colon cancer patient survival treated with any type of adjuvant chemotherapy. For *CST3* data are plotted for 278 colon cancer patients split into low in black (n=96) and high in red (n=182). For *EPM1* the numbers were 185 for low expression and 93 for high expression. Numbers below the graph indicate the surviving patients at each time. Marks on the graph indicate censored patients. The hazard ration (HR) and exact log rank P value are shown for each transcript. * *P*<0.05, ** *P*<0.01, *** *P*<0.001, **** *P*<0.001

Supplemental Figure 7

A) Representation of the binding specificities of the lectins used in this study. Lectin names are showed in coloured text, matching the colours in Figure 6C. B) RNA precipitated from the cell-free growth media of four colorectal cancer cell lines was analysed for glycoRNA levels using the Peanut agglutinin (PNA) lectin. Values are plotted as the fluorescence per µg of RNA from 3 biological replicates ±SEM after normalisation to *in vitro* transcribed (IVT) RNA. HCT116 cells (white bar) are non-mucinous in origin, while the other cells lines (red bars) are mucinous. C) The amount of RNA isolated from the growth media of the same four cell lines as in B are plotted as the mean ng/mL of media from at least 3 biological replicates ±SEM. D) LS174T cells were treated with the N-glycosylation inhibitor NGI-1 for 24 hours at the indicated concentrations and RNA precipitated from cell-free media. PNA glycoRNA levels are expressed as the fold change in fluorescence normalised against IVT RNA compared to vehicle treatment. E) The percentage of RNA recovered from AquIRE analyses from different mouse samples and an IVT RNA sample in Figure 6G are plotted. Each shape represents an individual animal of 3 biological replicates. F) The scatter graph plots the fluorescence data from Figure 6G against the percent recovery of RNA in E. The red line shows a simple linear regression and the dashed lines are the 95% confidence interval. Data of the fit of the linear regression are inset into the plot. G) RNA was extracted from *Arabidopsis* leaves (as analysed in Figure 6H) and compared to RNA extracted from whole 7-day old seedlings. Data are plotted as the mean fold change in fluorescence from the leaf samples for 3 independent plants. H) Dark-induced senescence was induced in *Arabidopsis thaliana* Col-0 by wrapping selected leaves in aluminium foil while on the plant. Representative images of i) a 7-day old seedling ii) a leaf exposed to light, 8h per day, for 6 days. iii) a leaf after 4 days of the dark and iv) a leaf after 6 days of the dark. The leaves come from the same plant. I) W303-1A *Saccharomyces* were grown to OD0.8 then pelleted at low speed for cell RNA extraction. The supernatant was spun again to isolate cell debris, leaving a cell-free supernatant. RNA was extracted from all 3 fractions (cell pellet, cell debris and cell-free media) and analysed by PNA AquIRE. Data are the fraction, as a percent, of the RNA and glycoRNA signal found in each fraction from two biological replicates ±SEM. Significance was tested using a 2-way ANOVA with Šídák multiple comparison testing comparing the cell-free signal to the sum of the two other fractions. J) A single graph showing the relative levels of glycoRNA across the biological samples analysed in this work. Data are plotted as the raw (not normalised) values for each sample, including an IVT control in the final row. Data are from a minimum of 2 biological replicates, with individual replicate numbers visible from the small grey circles on the figure. Data are the mean values plotted ±SEM. Cell/tissue-based samples are plotted in rank order in grey. Cell-free samples are plotted in rank order below this in red. K) RNA was extracted from U251 glioblastoma cells following treatment with 2mM temozolomide for the indicated times. Left is a graph of the mean m7G signal detected by AquIRE from at least 3 biological replicates, plotted with the SEM. Significance was determined by mixed affects analysis. Right shows the methylation of guanosine at position 7 (orange) that is being detected.

## STAR methods

### Lead contact

Request for information or resources should be directed to John Knight

### Materials availability

This study did not create any new reagents

### Data availability

Proteomic data have been deposited on ProteomeXchange and are publicly available on the date of publication. Accession numbers are in the key resources table. Original western blot images are publicly available on Mendeley Data (doi: 10.17632/4ckcxhcyhg.1). All microscopy images are available from the lead contact on request. No original code was used in this study. The mass spectrometry proteomics data have been deposited to the ProteomeXchange Consortium via the PRIDE^63^ partner repository with the dataset identifier PXD06117. Any additional requests for information can be sent to the lead contact.

### Experimental models and subject details

#### *In vivo* animal studies

*Mus musculus*: adult wild-type mixed background animals of both genders were used in this study. Animals were kept under Establishment Licence number X44772EDA, granted by the UK Home Office, in individually ventilated cages with *ad libidum* access to water and diet in a 12:12 light cycle. *Xenopus tropicalis*: Adult male and female frogs primed with 15 units of pregnant mare serum gonadotrophin (MSD Animal Health) 18–24 hours prior to ovulation. Mating was subsequently induced with 50 units of human chorionic gonadotrophin (MSD Animal Health) in males and 75 units in females. Hormone injection in adults was performed under United Kingdom Home Office animal project licence number PFDA14F2D. All data presented in this study were obtained from pre-feeding stage embryos (approximately 3–4 days of development from fertilisation) which are not considered protected animals for regulated procedures under the Animals (Scientific Procedures) Act 1986. All experiments using *Xenopus tropicalis* animals are reported according to applicable ARRIVE guidelines for this species. *Drosophila melanogaster*: *y*^1^ *w*^67c^^23^ flies were housed in standard conditions. For ovary tissue samples, 8-10 ovary pairs from 2–5-day old non-virgin females were used. Embryos were harvested at the specified timepoints after being laid on apple juice agar plates supplemented with yeast paste in small cages.

#### Human participants

Access to colorectal primary or metastatic tumours was granted by the Manchester Cancer Research Centre Biobank, application number 23_JOKN_01. Approval is under the MCRC Biobank Research Tissue Bank Ethics, reference 22/NW/0237. Anonymised details of the patients are available in Supplemental Table 4. Survival analyses were based on publicly available dataset, analysed using the KMplot online tool^64^. Details of the datasets used in these analyses, expression ranges and cut offs are in Supplementary Table 1.

#### Cell lines

All cell lines were maintained under standardized conditions at 37°C in a humidified atmosphere containing 5% CO_2_. Medium were refreshed every 2-3 days, and cells were passaged using trypsin-EDTA when they reached approximately 80% confluence. Specific medium compositions for each cell line are detailed: LS174T, RKO and DLD1 cells were grown in MEM supplemented with 10% FBS, 2 mM L-glutamine, 1x NEAA, and 1% PenStrep. A172 and U251 were grown in DMEM supplemented with 10% FBS, 2 mM L-glutamine and 1% PenStrep. HCT116 were grown in either the MEM or DMEM base media listed above. JVE-127 were cultured in RPMI supplemented with 10% FBS, 2 mM L-glutamine and 1% PenStrep. JVE-253 were grown in RPMI supplemented with 10% FBS, 2 mM GlutaMAX and 1% PenStrep. HeLa-FBL-GFP cells were grown in DMEM medium supplemented with 10% FBS and 1% PenStrep mix.

#### Microorganisms

*Saccharomyces cerevisiae* W303-1A were grown in SCD media (1x Yeast Nitrogen Base w/o amino acid, 1x Kaiser Complete SC media and 2% D-glucose) and harvested at an OD_600_ of 0.8. One Shot TOP10 Chemically Competent *E. coli* were cultured in Luria broth and harvested in exponential growth phase.

#### Plant models

Seedling growth conditions: *Arabidopsis thaliana Col-0* seeds were sterilised for 10 minutes under rotation with a sterilising solution (50% ethanol and 0.5% Triton X100) followed by five subsequent washes using sterile distilled water. Sterile seeds were sown onto sterile plates containing 1% glucose, 0.8% agar and ½ Murashige and Skoog basal medium (Duchefa) and stratified for 2 days at 4°C in darkness. Seeds were grown in continuous light (69 μmol/m^2^s) at 24°C in a growth cabinet (Perceval, Perry, IA, USA) for 7 days. Plant growth conditions: Seeds were planted in Levington Advance F2, grown in 8 cm pots and watered regularly. They were grown in controlled growth cabinets (Perceval, Perry, IA, USA), under a short-day photoperiod of 8h light (112 μmol/m^2^s intensity) at 22°C during the day and 17°C during the night.o

### Method details

#### Aqueous Identification of RNA Elements (AquIRE)

The input material for all AquIRE assays was purified RNA. Equal amounts of RNA per sample were used within each experiment. The RNA was polyadenylated using *E.Coli* polyA polymerase (NEB), unless otherwise stated, as per the manufacturer’s protocol recommendations. Incubation of the polyadenylated RNA in denaturing buffer (20mM TRIS pH7.5, 200mM NaCl and 2% w/v SDS) for five minutes at 65°C ensured removal of RNA secondary structures, following which the samples were rapidly cooled to 4°C. Magnetic oligodT beads (Cytiva) were washed twice with denaturing buffer before binding the RNA to the beads at a ratio of one µg of RNA per 10 µl of washed beads. A five-minute-long incubation at room temperature allowed oligodT:poly-A-tail base-pairing, which was followed by two washes, one with denaturing buffer and a subsequent one with wash buffer (20mM TRIS pH7.5, 200mM NaCl and 1mM EDTA).

*Immunodetection:* Primary antibodies were applied with 1:200 and 1:500 dilution, then recognised by species-specific biotinylated secondary antibodies. Secondary antibodies were subsequently bound by fluorescently-labelled streptavidin (Invitrogen). Wash buffer was used as the diluent for all antibodies and streptavidin. Degradation of RNA was prevented by the addition of RiboLock (Thermo Scientific), at 0.1U/μL. All incubation steps were performed on a shaker for an hour at room temperature except the primary antibody incubation, which lasted for an hour and a half. Each incubation step was followed by two wash steps using wash buffer removing any unbound material. *Glycan detection:* All glycan detection steps were the same as the AquIRE steps described above apart from the immunodetection step, replaced by the use of glycan-binding biotinylated lectins. A dilution factor of 1:200 was used for the preparation of the biotinylated lectin incubation solution. The lectin incubation was performed on a shaker for 1.5 hours at room temperature. *Aqueous elution and signal detection:* The complexes of RNAs and antibodies / lectins of interest were eluted using nuclease-free water, which was incubated for 10 minutes at room temperature. Fluorescence reading in black opaque 96 well plates was conducted using a microplate reader and SkanIt™ Software (Thermo Scientific). The specified excitation and emission wavelengths were 488 nm and 515 nm, respectively. *NanoOrange:* A dilution factor of 1:5000 was used for the preparation of the NanoOrange (Invitrogen) incubation solution. The NanoOrange reagent was diluted with wash buffer and was allowed to bind to crosslinked proteins for 10 minutes at room temperature. A single wash step using wash buffer preceded the nuclease-free water-based elution, which was performed for 10 minutes at room temperature. The fluorescence signal was detected by specifying the excitation and emission wavelengths as 485 nm and 590 nm, respectively. The NanoOrange incubation solution served as a positive control for fluorescence. *Data analysis:* The raw fluorescence values presented are an average of triplicate reads from the final eluate of each sample after removal of outliers. Where the AquIRE signal has been normalised, the signal of the negative control is subtracted then values presented relative to a given sample, either set to 1 or 0. For glycoRNA samples the fluorescence is additionally normalised against IVT RNA (i.e. with no glycoRNAs). Where the fluorescence is presented per µg of RNA, the mean fluorescence reads are divided by the amount of input RNA.

#### RNA isolation

RNA was isolated by Zymo’s Quick-RNA^TM^ Miniprep Kit with DNase digestion, unless otherwise stated, as per the manufacturer’s protocol recommendations. The lysis step differed across sample types. Human cell pellets were lysed by immediate incubation with Zymo RNA lysis buffer. Bacterial cell pellets, obtained by a 15-minute-long centrifugation at 3000 x g at 4°C, were resuspended in PBS and then lysed in 3 volumes of Zymo RNA lysis buffer. Mouse tissues were sampled into RNAlater (Invitrogen) and stored at-80°C prior to extraction. These were lysed in Zymo RNA lysis buffer using gentleMACS M tubes (Militenyi). Whole mouse blood from cardiac puncture was placed into heparin containing tubes (Teklab) and separated into blood cells and plasma by centrifugation at 10,000g for 5mins. Blood cells were directly lysed in Zymo RNA lysis buffer while plasma was mixed with the same buffer at a ratio of 1 part plasma to 3 parts lysis buffer. Five whole *Xenopus* embryos were pooled and lysed directly in Zymo RNA lysis buffer with repeated pipetting. For large volume liquid samples (cell line media, bacterial broth, cell-free human tumour material, polysome profile fractions, *Xenopus* MMR media, commercially available sera) these were mixed with an equal volume of 7.7M guanidine hydrochloride, then an equal volume again of 100% ethanol, giving a final ratio of 1:1:2. This was stored at-20°C for at least 24 hours then RNA pelleted at 4000g for 45 minutes. RNA pellets were dissolved in Zymo RNA lysis buffer. For *Drosophila* RNA extractions, embryos were dechorionated for 2 minutes in 50% bleach (2.5% final concentration of sodium hypochlorite diluted in distilled water) and rinsed thoroughly in distilled water. For ovaries, these were dissected in 1x PBS on ice and transferred to 1.5mL microcentrifuge tubes and the PBS removed. 50μL TRIzol Reagent (Invitrogen) was added to the embryos or ovaries and samples were crushed and homogenised using a disposable pestle (Fisher Scientific). An additional 450μL Trizol was added to rinse the pestle and then RNA was extracted and purified according to the manufacturer’s protocol. For *Saccharomyces cerevisiae* RNA extractions, cells were pelleted at 1500g for 15 minutes at 4°C and immediately lysed in Trizol. Supernatant was centrifuged again at 15000g for 30 minutes at 4°C to yield cell debris, which was placed in Trizol. The supernatant was then concentrated ∼20-fold using a Vivaspin 20 column with 5kDa cutoff (Sartorius) and RNA extracted using Trizol. For *Arabidopsis* RNA extraction, the Aurum Total RNA Mini Kit from BioRad was used, following the manufacturer’s guidelines. RNA was extracted from 10 pooled seedlings or one leaf (up to 100mg).

#### Plant senescence model

To induce senescence, individual 5-week-old leaves were wrapped in aluminium foil without detaching the leaves from the plants. After 4 and 6 days, the dark-induced leaves were unwrapped, photographed and total RNA extracted. The control leaves, unwrapped, were collected at day 6.

#### Ionomycin secretagogue protocol

Experiments were performed as previously published ^65^. Batches of 50-60 *Xenopus tropicalis* embryos at NF stage 40 were transferred into 3 ml 0.01 X MMR media (0.1M NaCl, 2mM KCl, 1mM MgSO_4_, 2mM CaCl_2_, 5mM HEPES (pH 7.8), 0.1mM EDTA) in single wells of a 12-well plate, and incubated at 25°C. Embryos were exposed to 4 µM ionomycin (Sigma Aldrich), and incubated at room temperature for 10 minutes with gentle swirling each minute. Media were centrifuged at 10000 g for 5 minutes to pellet cellular debris, and RNA extracted as outlined above for large volume liquid samples.

#### RNA analysis, crosslinking and digestion

*RNA integrity number (RIN) analysis*: A 2100 Bioanalyzer was used with RNA 6000 Pico Kit set to Eukaryote Total RNA. RIN values were calculated using the Bioanalyzer software. RNA concentrations were routinely determined using a NanoDrop. *UV crosslinking*: At experiment end, culture medium was aspirated from cells and pre-warmed PBS was added to rinse the cells, followed by thorough aspiration of the PBS. The cells were then placed in a UV_254nm_ crosslinker, with the UV intensity set to 100 J/m². Upon completion of the countdown, the RNA, with crosslinked proteins, was extracted. Control cells were incubated in the absence of PBS or UV light for the same duration. *PNGase F digests*: 1.5µg RNA was incubated with recombinant PNGase F (NEB) using 0.75µL enzyme in a 10µL reaction for 2 hours at 37°C. A no enzyme control was incubated in parallel. Subsequently, RNA was purified using RNA Clean and Concentrator columns (Zymo) before quantification and analysis of equal amounts of RNA by AquIRE. *RNase I digests*: Total RNA was incubated with 0.4U/µL RNase I (Thermo) at 37°C for 1 hour. Protein was precipitated by the addition of trichloroacetic acid to 12.5%, washed twice with a buffer of 50mM TRIS (pH7.5), 70% acetone and 20% ethanol. Finally, samples were dissolved in 6M urea. *Proteinase K digests*: On-bead digest used 50 µg/mL Proteinase K (Roche) directly prior to primary antibody incubation. AquIRE wash buffer was used as a diluent of proteinase K, which was incubated with bead-immobilised RNA for an hour at 37°C, followed by two wash steps using AquIRE wash buffer. Once the washing was complete, the RNA was analysed from the primary antibody incubation step as normal. *RNase A digests*: Cells were treated with non-cell permeable RNase A (NEB) concurrently with drug treatments by diluting the enzyme directly in cell media.

#### *In vitro* transcription

All reactions used the HighYield T7 RNA Synthesis Kit from Jena Bioscience following the manufacturer’s instructions. For incorporation of non-canonical nucleotides, these were substituted for the canonical nucleotide at the stated ratios. Specifically, 5FUTP or pseudoUTP were substituted for UTP or m6ATP for ATP. To template transcription we used either the DNA provided with the HighYield kit or a pUC57-Curlcake 3 DNA, expressed and digested as previously published ^66^. Reactions were incubated for 1.5 hours at 37°C in a thermal cycler. The reaction volume was then adjusted to 50 µL and RNA purified using the Monarch Spin RNA cleanup kit, and the RNA product stored at-80°C.

#### RNA dot blot

Nitrocellulose membranes (Fisher Scientific) were soaked in sterile distilled water, washed in 10X SSC (1.5M NaCl, 150mM tri-sodium citrate), and air-dried. RNA samples were thawed on ice, mixed with 3 volume of RNA incubation solution (66% formamide, 8.5% formaldehyde, 150mM MOPS, 70mM sodium acetate, 7.7mM EDTA at pH 7) heated at 65°C for 5 minutes, and cooled on ice. An equal volume of ice-cold 10X SSC was added. The RNA solution was dotted onto the membrane and air dried. Membranes were exposed to 130kJ of UV_254nM_ in a crosslinker. Total RNA was visualised with methylene blue for 5 minutes, washed with water until the membrane turned white again and air-dried. For immunodetection, the membranes were blocked with 5% non-fat dry milk in TBST for 1 hour, then incubated overnight at 4°C with primary antibodies. After washing, the membranes were incubated with HRP-conjugated secondary antibody for 1 hour. Protein bands were detected using Clarity™ Western ECL substrate (Fisher Scientific) and visualized. Dot intensity was determined using ImageJ.

#### Human tumour processing

1000-4000mg of tumour was place in ADF media and processed within 16 hours of surgery. Tumours were cut into small pieces using scissors and/or scalpels then placed in gentleMACS C tubes (Militenyi) in at least 5mLs of ADF supplemented with 10mM EDTA and 1.5mg/mL collagenase II and run on the 37C_h_TDK_1 gentleMACS program. After digestion, 150µM collagenase inhibitor was added and the sample passed through a 100µm cell strainer (Starlab), which was washed with 0.5 volumes of ADF. Strained cells were then pelleted at 600g and washed twice in 5mLs of ADF. All three supernatants from these centrifugations were pooled and centrifuged at 1000g for 5mins. RNA from this cell-free media was then processed as detailed above for large volume RNA samples. A fraction of the cell pellet was directly lysed in Zymo RNA lysis buffer.

#### Drug treatments

5FU, STM2457, temozolomide, ionomycin and NGI-1 were dissolved in DMSO, stored at-20°C and used with 6 months or after 5 freeze-thaw cycles, whichever was sooner. Oxaliplatin and carboplatin were dissolved in water and kept at 4°C for a maximum of 1 week. Cisplatin was dissolved directly in cell culture media and kept at 4°C for a maximum of 1 week. All vehicle treatments used the same volume of drug-free DMSO/water/media.

#### Proteomics

7.72µg of RNA-bound protein extracted from all samples, previously released by RNase I digestion were reduced and alkylated using dithiothreitol and iodoacetamide. Samples were then processed using S-Trap (Protifi) columns following the manufacturer’s guidelines, with final elution in 30% aqueous acetonitrile containing 0.1% formic acid. Samples were desalted using FiltrEx desalt filter plates (Corning). Separation was performed on a Thermo RSLC system consisting of a NCP3200RS nano pump, WPS3000TPS autosampler and TCC3000RS column oven configured with buffer A as 0.1% formic acid in water and buffer B as 0.1% formic acid in acetonitrile. An injection volume of 2μL was loaded into the end of a 5μL loop and reversed flushed on to the analytical column (Waters nanoEase M/Z Peptide CSH C18 Column, 130Å, 1.7µm, 75µm X 250mm) kept at 35°C at a flow rate of 300nL/minute for 8 minutes with an initial pulse of 500nL/minute for 0.3 minutes to rapidly re-pressurise the column. The injection valve was set to load before a separation consisting of a multistage gradient of 1% B to 6% B over 2 minutes, 6% B to 18% B over 44 minutes, 18% B to 29% B over 7 minutes and 29% B to 65% B over 1 minute before washing for 4 minutes at 65% B and dropping down to 2% B in 1 minute. The complete method time was 75 minutes. The analytical column was connected to a Thermo Exploris 480 mass spectrometry system via a Thermo nanospray Flex Ion source via a 20μm ID fused silica capillary. The capillary was connected to a stainless-steel emitter with an outer diameter of 150μm and an inner diameter of 30μm (Thermo Scientific, ES542) via a butt-to-butt connection in a steel union. The nanospray voltage was set at 1900 V and the ion transfer tube temperature set to 275°C. Data was acquired in a data-dependent manner using a fixed cycle time of 1.5 sec, an expected peak width of 15 seconds and a default charge state of 2. Full mass spectrum data was acquired in positive mode over a scan range of 300 to 1750 Th, with a resolution of 120,000, a normalised AGC target of 300% and a max fill time of 25 mS for a single microscan. Fragmentation data was obtained from signals with a charge state of +2 or +3 and an intensity over 5,000 and they were dynamically excluded from further analysis for a period of 15 seconds after a single acquisition within a 10ppm window. Fragmentation spectra were acquired with a resolution of 15,000 with a normalised collision energy of 30%, a normalised AGC target of 300%, first mass of 110 Th and a max fill time of 25 mS for a single microscan. All data was collected in profile mode. The resulting data was analysed using Proteome Discoverer (v3.1). The data was processed using the consensus workflow provided with PD in the file ‘CWF_Comprehensive_Enhanced_Annotation_LFQ_and_Precursor_Quan’, and the processing workflow provided with PD in the file ‘PWF_OT_Precursor_Quan_and_FQ_CID_SequestHT_Percolator’. The processing workflow set the search engine SequestHT to search the SwissProt database against human proteins (TaxID = 9606, v27/3/2024). The protein identification algorithm was provided with trypsin as the cleavage enzyme using the strict trypsin specificity; cleavage at Lysines and Arginines except where the presence of a C-terminal Proline obstructs cleavage. A maximum of 2 missed cleavages were permitted, charge states +1 to +6 included, fixed modifications of carbamidomethyl (+57.021Da) to cysteines, and dynamic modifications of oxidation (+15.995Da) to methionine. A precursor tolerance of 10ppm, and a fragment tolerance of 0.02Da were used to search, with an FDR cut off of 0.01. An FDR is calculated for both the protein level and the peptide level by PD. Proteins are labelled with High Confidence FDR 0.05. The raw output of this analysis is available in Supplemental Table 2. These high confidence proteins were analysed for Gene Ontology. Selected GO terms were reported to focus on terms with the fewest members. Terms higher in the hierarchy were omitted if no additional factors were added.

#### Digital holographic microscopy and data analysis

HeLa-FBL-GFP cells ^67^ were seeded onto ibidi µ channel slides. At the end of the experiment, cells were fixed with ice-cold methanol for 5 minutes and washed 3x with PBS. Slides were imaged with a Zeiss Axio Observer Z1 driven by MetaXpress, equipped with a QMOD off-axis differential interferometer and a Retiga R3 camera ^38^. Holograms were captured using transmitted light (HAL lamp fitted with a single-band bandpass optical filter). Holograms were imaged with a 20x (0.5 NA) EC Plan Neofluar objective and converted to DHM phase using the OsOne software. Images were exported as.tiff and.bin files. ImageJ was used for the analysis of the images. To generate the nucleolar masks, the background was subtracted using a rolling ball of a 50 px radius. Images were transformed into 8-bit format. Dead cells were excluded from the analysis. The image was thresholded using the MaxEntropy algorithm before being converted to a mask. Particles in the size spectrum of 4-150 px and the circularity 0.55-1.00 were included in the analysis. The nucleolar area, roundness (calculated as, 4*area/(π*major_axis^2^)), and circularity (calculated as, 4π*area/perimeter^2^) was extracted from the data. For the analysis of nucleolar optical thickness, ROIs of 12 px centred in each nucleolus were used to determine the nucleolar phase shift. As reference, the nuclear phase shift was measured in 12 px ROIs adjacent to the nucleolus. Phase shift values were transformed into OPL values following OPL= (phase shift*λ)/2π. The OPL of each nucleolus was normalized to the OPL of each corresponding nucleus by subtraction ^38^.

#### Immunofluorescence and analysis

Staining and single-cell analysis of 5FU incorporation into RNA was performed as previously described ^21^. For stress granule staining with G3BP1, cells were washed in PBS and fixed in pre-chilled 70% ethanol for 30 minutes at-20°C. Cells were blocked with PBS supplemented with 2.5% bovine serum albumin (BSA), 10% normal goat serum (NGS) and 0.5% Triton X-100 for 1 hour. Anti-G3BP1 primary antibody (PoteinTech) at 1:1000 dilution in 1% BSA / 5% NGS / PBS was added overnight at 4°C. Anti-rabbit CoraLite 594 (ProteinTech) secondary antibody at 1:300 dilution in 1% BSA / PBS was added for one hour. Cells were then counterstained with DAPI (Merck), and coverslips mounted onto microscopy slides. Images were taken using confocal microscopy (Olympus Spinning Disk) at 40x magnification. At least 5 fields of view were captured per condition and the percentage of stress granule positive (at least 3 cytoplasmic G3BP1 granules) cells scored manually.

#### Western blotting

Protein samples were standardised to the same mass and separated by SDS-PAGE (NuPAGE). The separated proteins were transferred onto nitrocellulose membranes, which were blocked with 5% non-fat dry milk in TBST for 1 hour at room temperature. Next, membranes were incubated with primary antibodies specific to the target proteins, diluted in blocking buffer, overnight at 4°C. After primary antibody incubation, the membranes were washed three times with TBST and then incubated with HRP-conjugated secondary antibody for 1 hour at room temperature. After a final series of washes, protein bands were detected using a Clarity^TM^ Western ECL substrate (Fisher Scientific) and visualized using a ChemiDoc. Band intensities were determined using Image J.

#### Cellular ROS analysis

Intracellular ROS levels were assessed using the DCFDA Cellular ROS Detection Assay Kit (Abcam) according to the manufacturer’s protocol. Cells were cultured in a black 96-well plates. The concentration of the positive control [tert-butyl hydroperoxide (TBHP)] was 100μM. Fluorescence at 485nm / 535nm was recorded using the microplate reader at the end of the experiment. Background fluorescence was subtracted from each value before fold increase in fluorescence intensity relative to the negative control was determined.

#### cDNA stalling assays

SuperScript™ II Reverse Transcriptase kit (Invitrogen), was used with random primers (Promega), dNTP Mix (Invitrogen) and RiboLock (Thermo) to generate cDNAs from total RNA in a total reaction volume of 20μl as per the kit recommendations. The amount of random primers used for an RNA input amount of 1000 ng was 200 ng and it was decreased or increased proportionally if the RNA input was smaller or greater, respectively. The generated cDNAs were then used in a 10μL qPCR including PowerTrack™ SYBR™ Green Master Mix (Applied Biosystems) and target-specific primers (Integrated DNA Technologies) as per the kit recommendations. qPCRs were run on the QuantStudio 5-C Real-Time PCR System within 384-well format plates covered with adhesive seal using Thermo Scientific™ Design and Analysis Software 2. The run method involved a 2 minute-long hold step at 21° C and a subsequent 2 minute long Taq DNA polymerase-activating step at 95°C, which was followed by 40 thermal cycles of 15-second-long denaturation at 95°C and a minute long primer annealing and extension at 60°C. The data was analysed via the relative quantification ΔΔCt method. Normalisation was done against the transcripts from the housekeeping genes actin beta (*ACTB*) and glyceraldehyde-3-phosphate dehydrogenase (*GAPDH*), which was followed by normalisation against the control groups. Normalisation was done against *ACTB* and *GAPDH* as their raw Ct values were not changed, which was speculated to be due to their low abundance compared to the RNAs of interest. Interpretation of results was based on the direct proportionality between RT-qPCR signal reduction and drug-induced RNA damage.

#### Polysome profiling

This was performed as previously ^68^. Briefly, exponentially growing cells were treated with 200µg/mL cycloheximide for three minutes then transferred to ice. Media was removed and replaced with ice cold PBS also containing cycloheximide for detachment by scraping. Pelleted and washed cells were lysed (300mM NaCl, 15mM MgCl_2_, 15mM TRIS pH7.5, 100µg/mL cycloheximide, 0.1% Triton X-100, 2mM DTT and 0.2U/mL RiboLock). Post-nuclear supernatants were then layered on 10-50% sucrose gradients containing the same concentration of NaCl, MgCl_2_, TRIS and cycloheximide as in the lysis buffer. These were then centrifuged in a JXN-30 centrifuge with a JS24.15 rotor at 79,000g at 4°C for 3.5hours. Gradients were then separated through a live UV_254nm_ detector and distributed into nine 1.4mL fractions. For RNA isolation, sub-polysome and polysome fractions were pooled and RNA extracted from the resulting large volume liquid samples as described above.

#### Statistical analysis

All statistical analyses on data produced in this project were performed using GraphPad Prism 10, with details of the individual test used found in the figure legends. Data is presented as the mean and error bars are the standard error of the mean (SEM), unless otherwise stated. The number and type of replicates are also depicted in each figure legend also. *P* values of 0.05 or less were considered significant.

## Key resources table

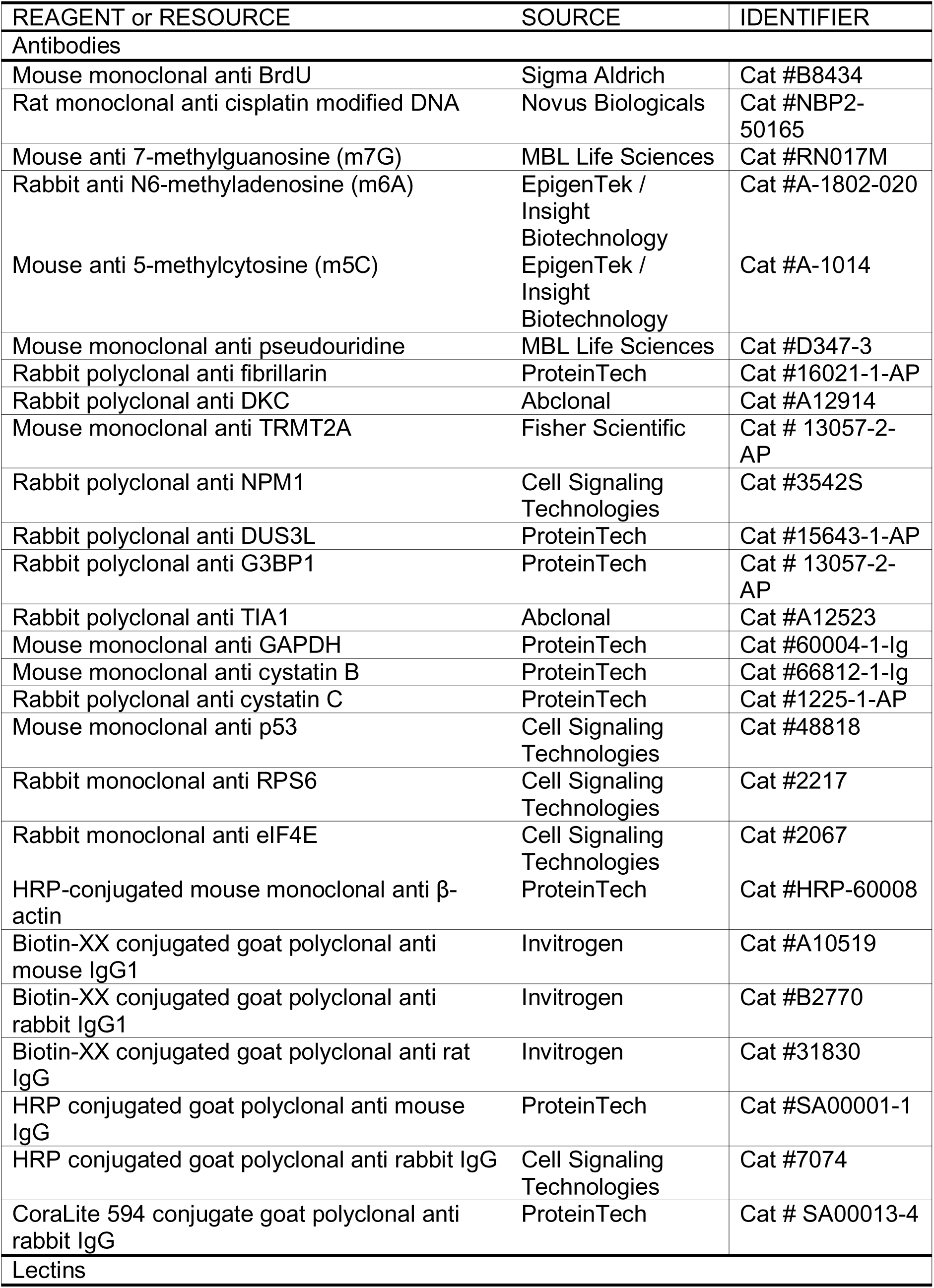
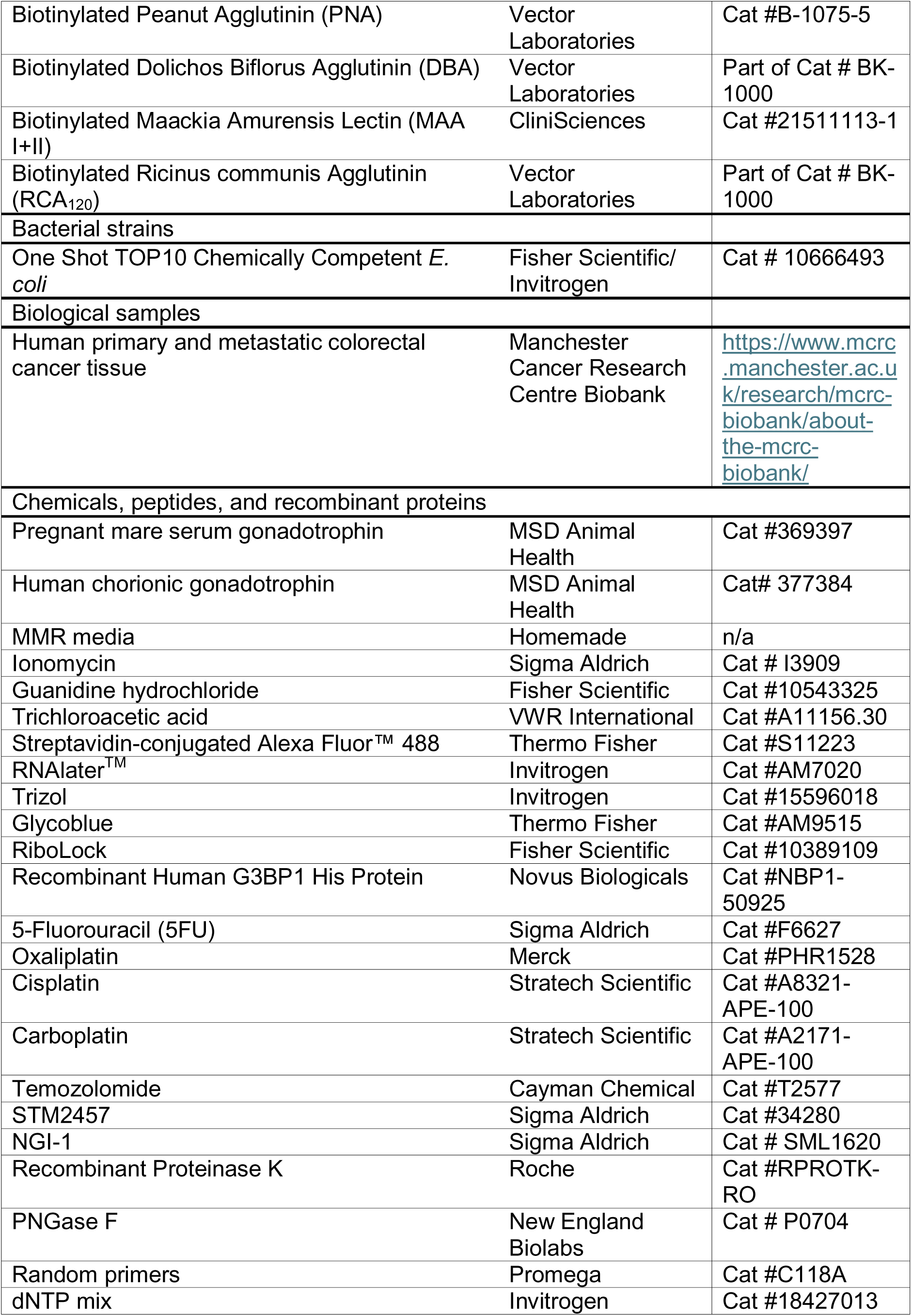
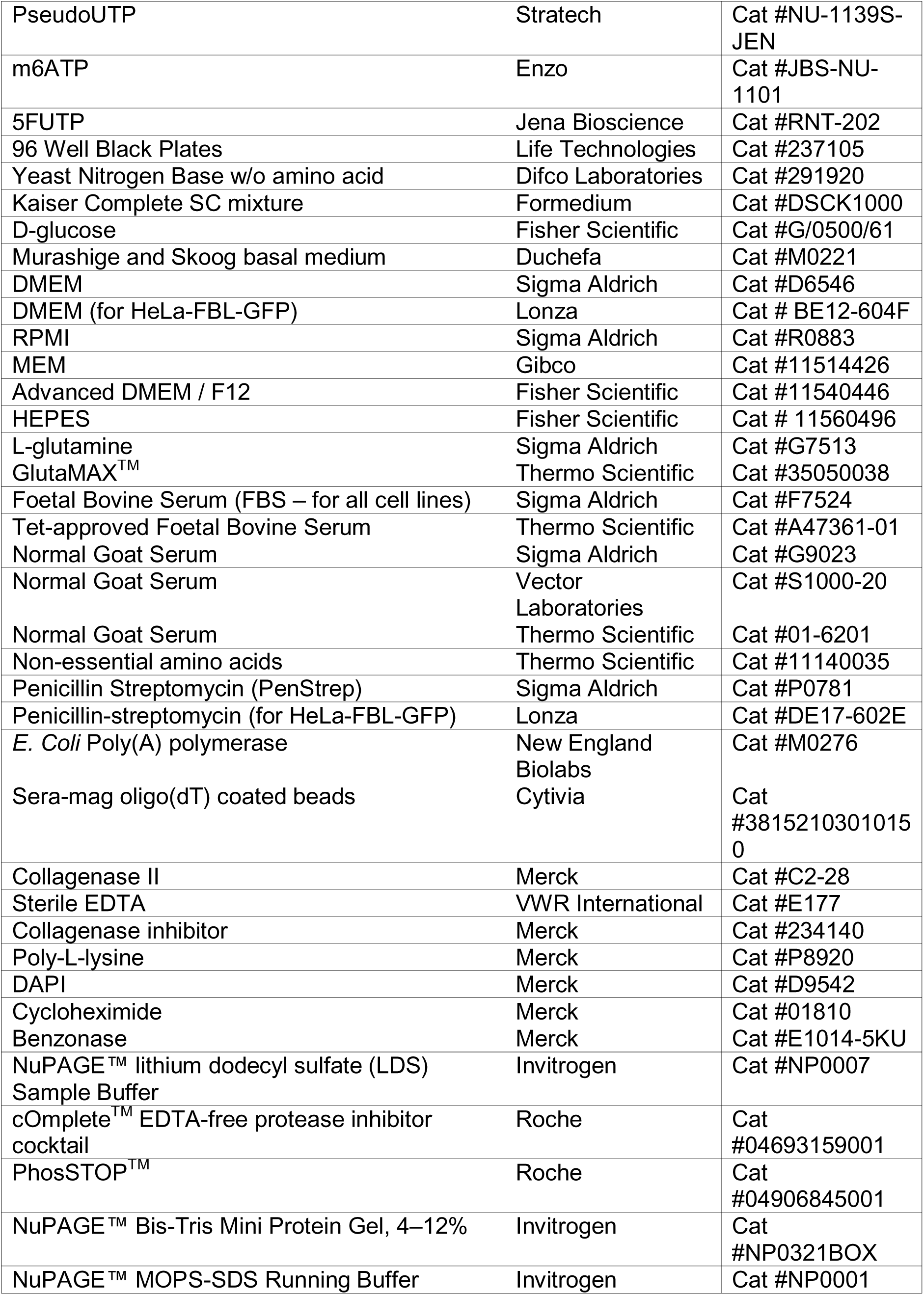
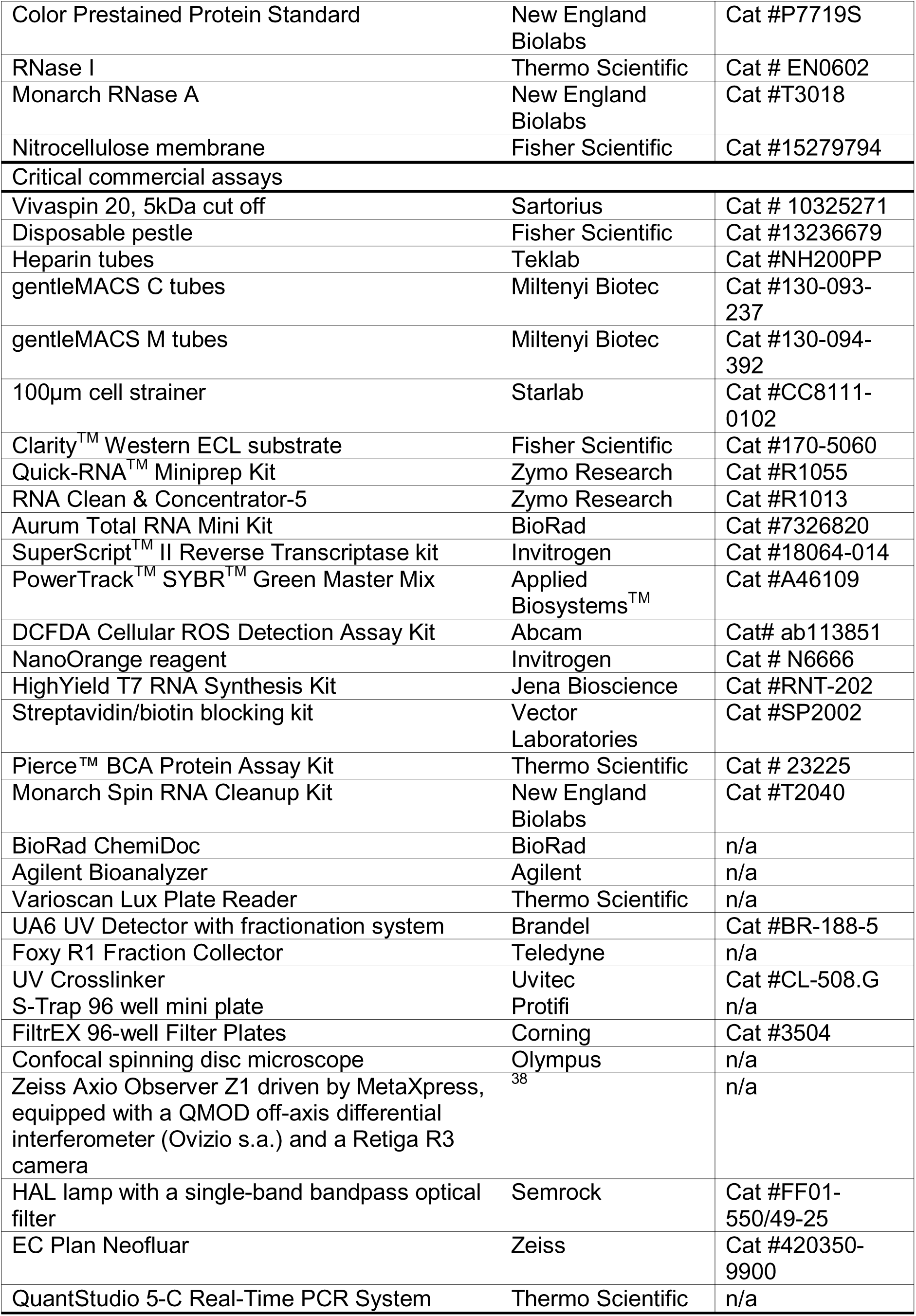
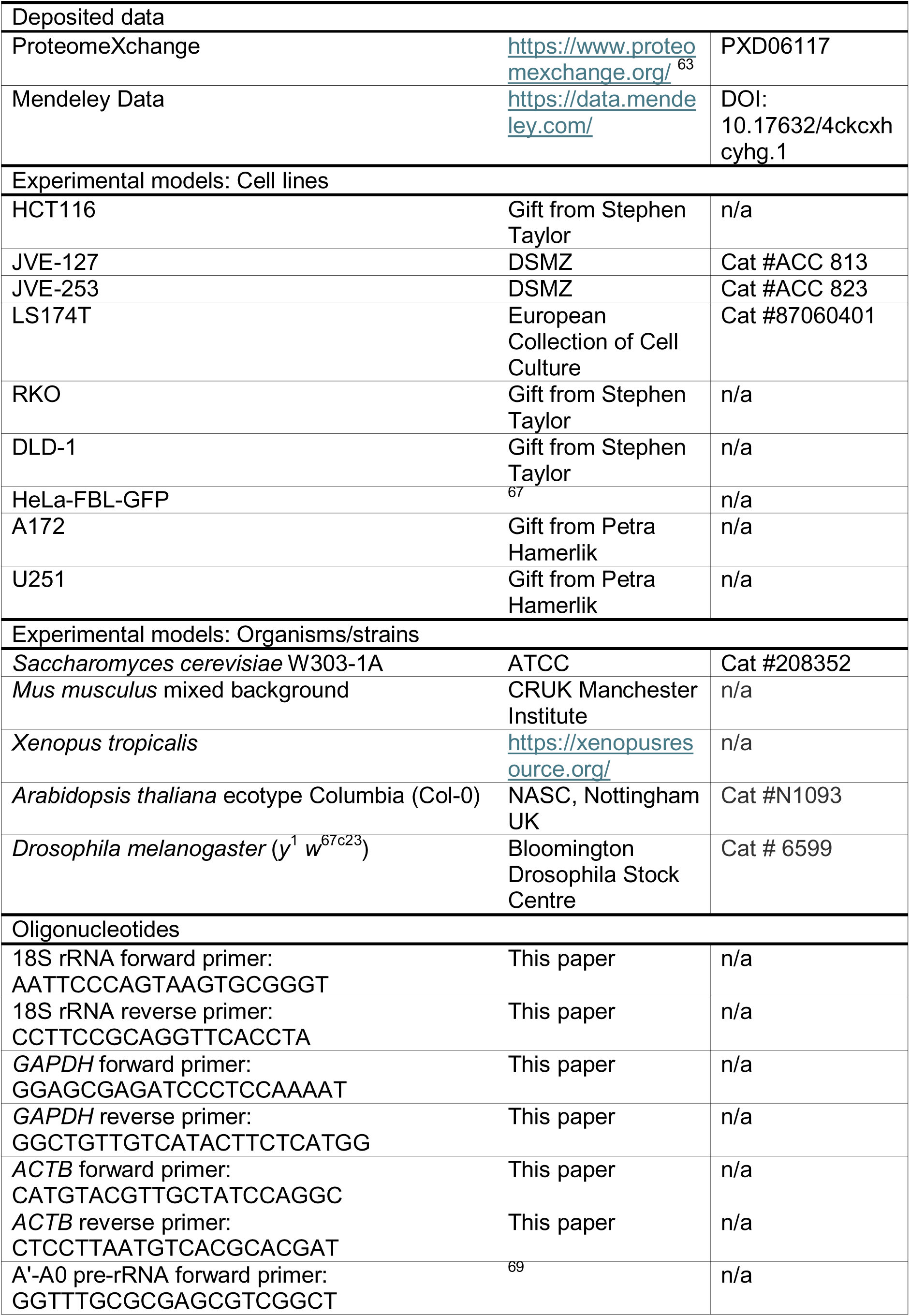
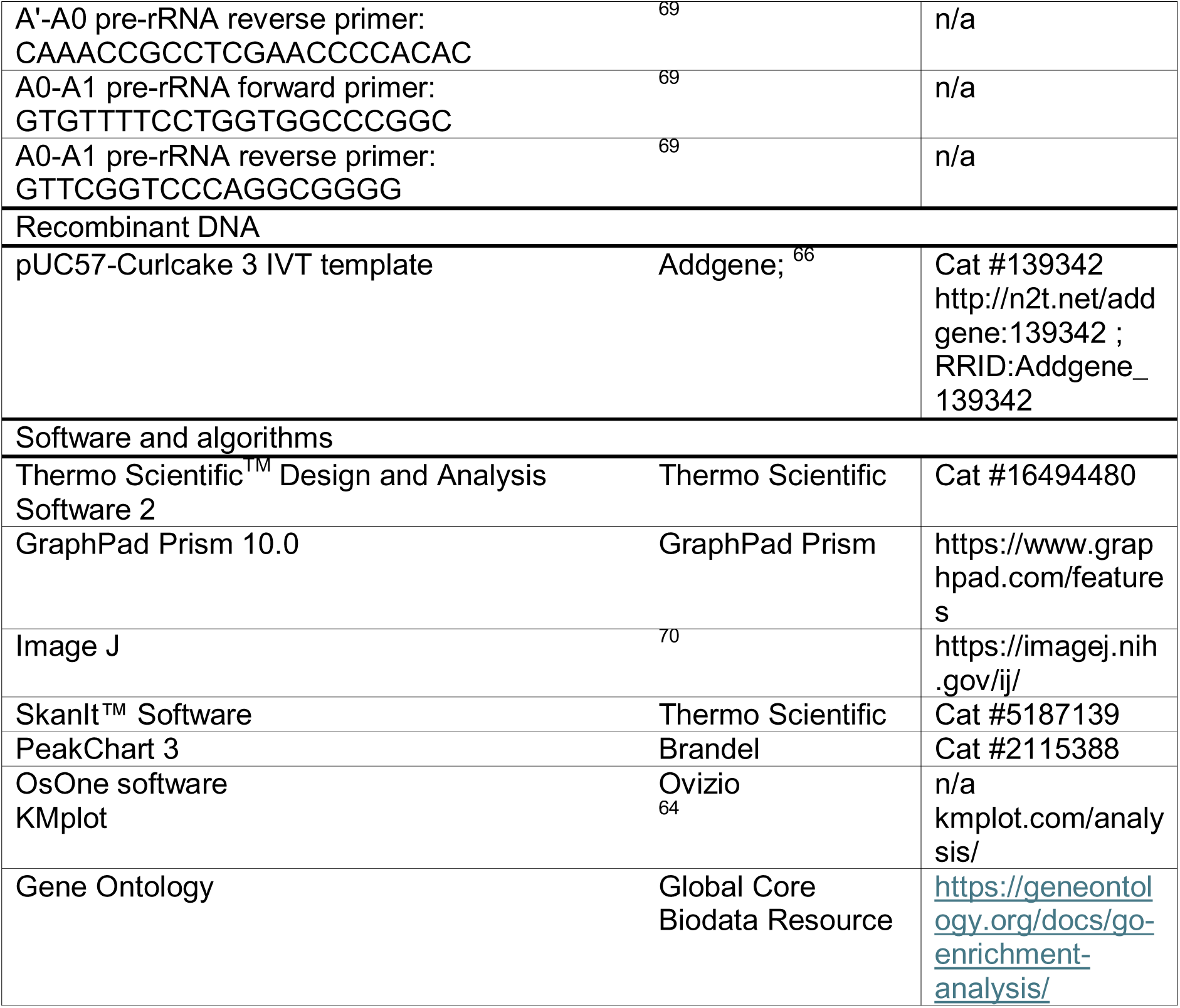

