## Supplementary figures and images for "AquIRE reveals multiple mechanisms of clinically induced RNA damage and the conservation and dynamics of glycoRNAs"

### Supplemental figure 1

Supplemental Figure 1

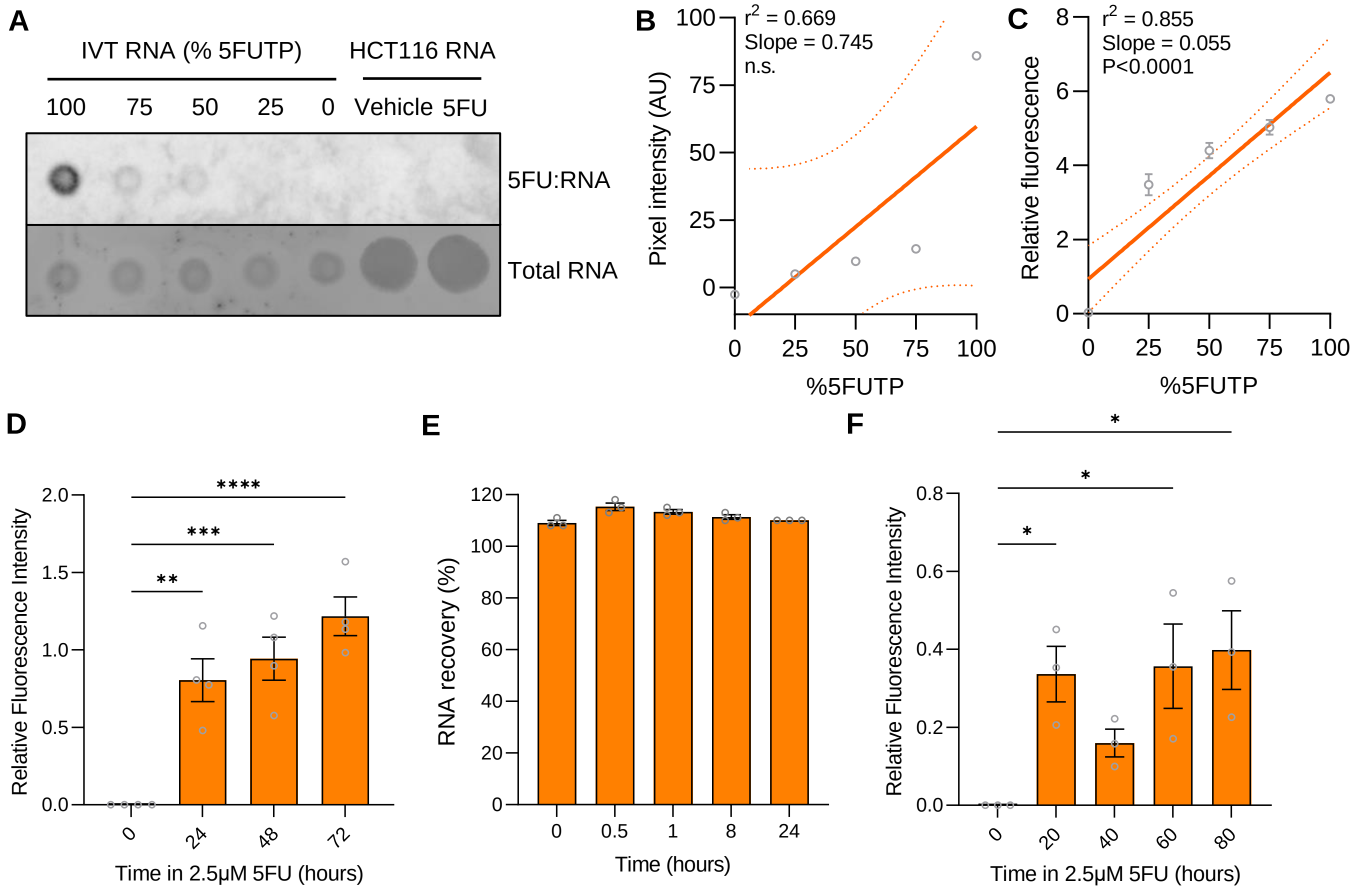

### Supplemental figure 2

**Supplemental Figure 2**

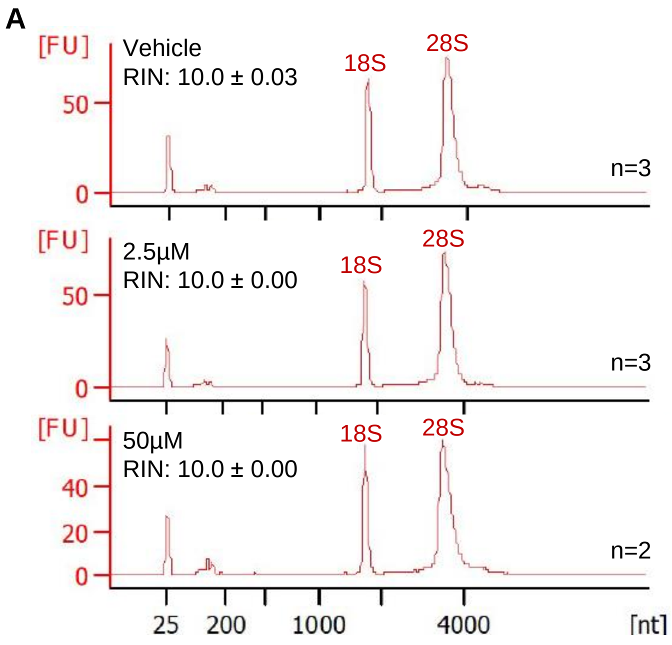

### Supplemental figure 3

#### Supplemental Figure 3

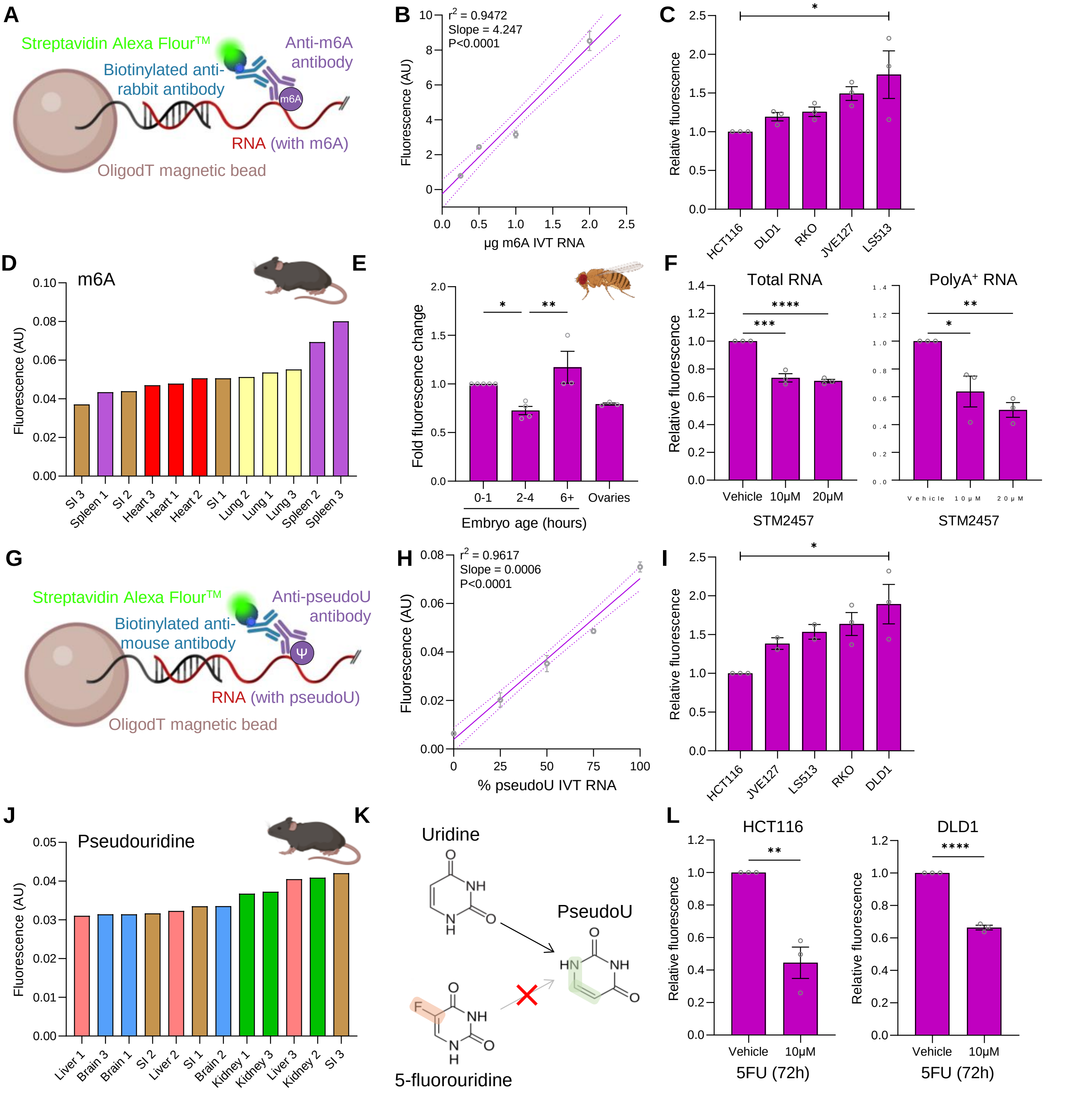

### Supplemental figure 4

##### Supplemental Figure 4

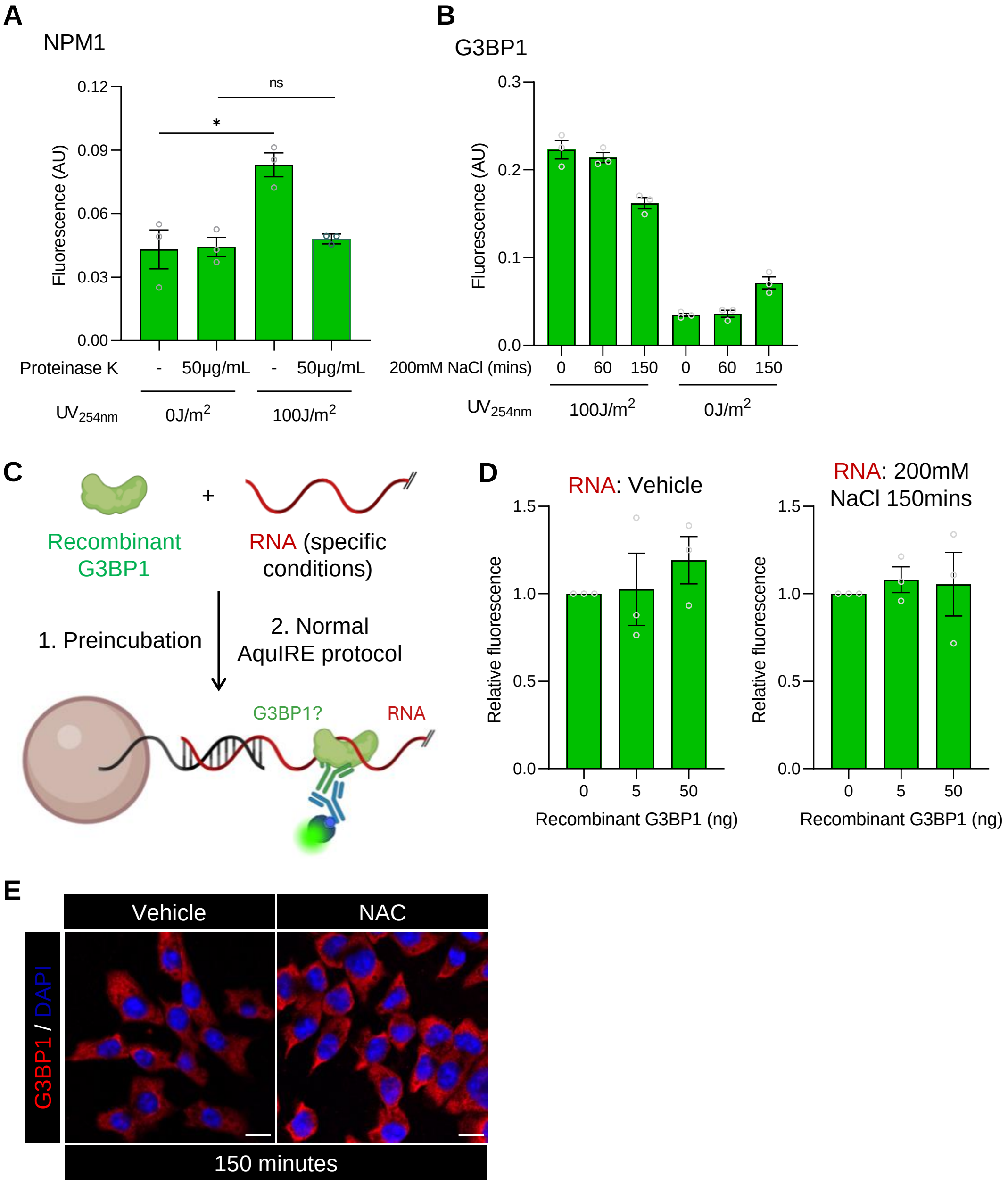

### Supplemental figure 5

#### Supplemental Figure 5

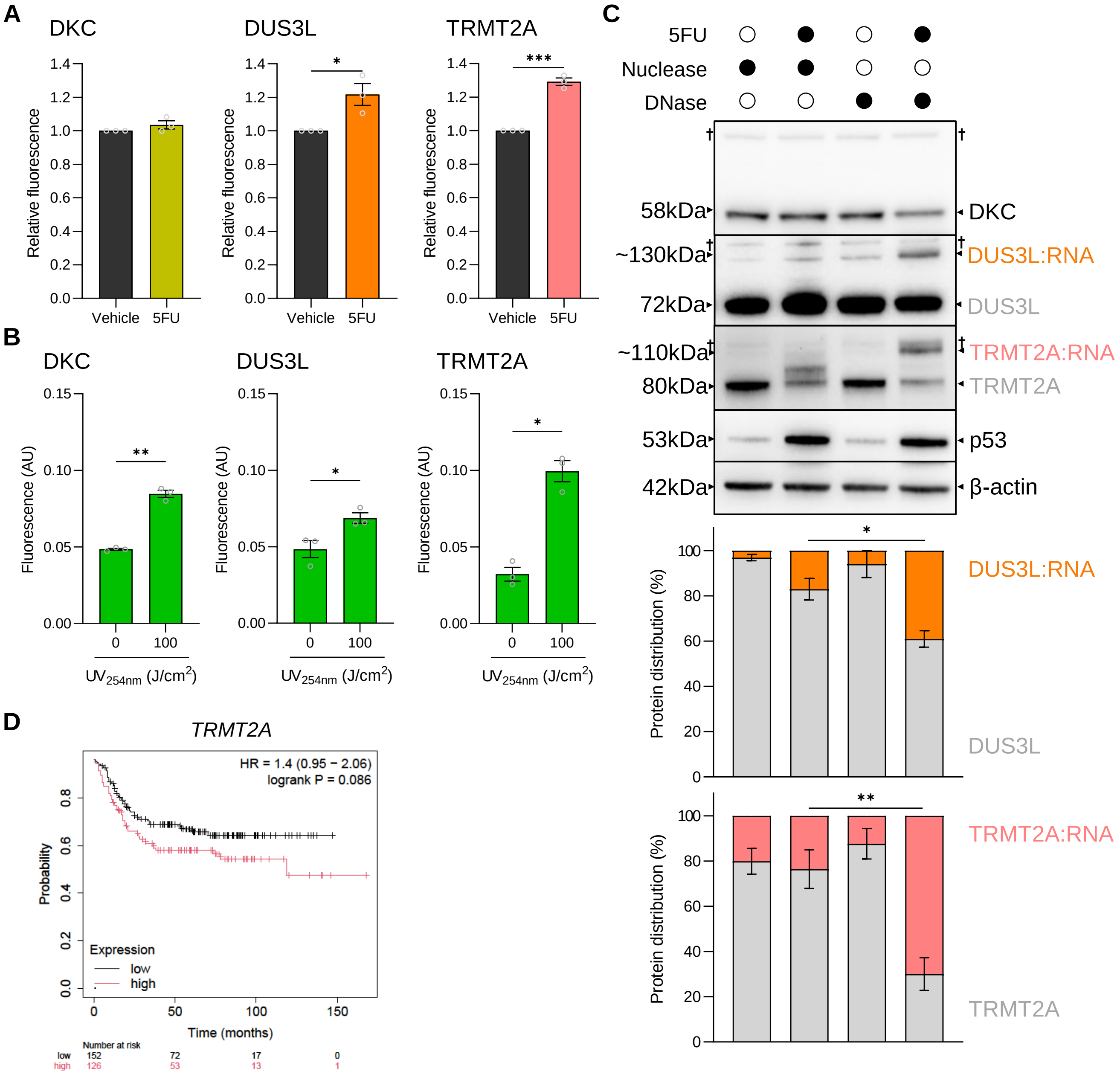

### Supplemental figure 6

#### Supplemental Figure 6

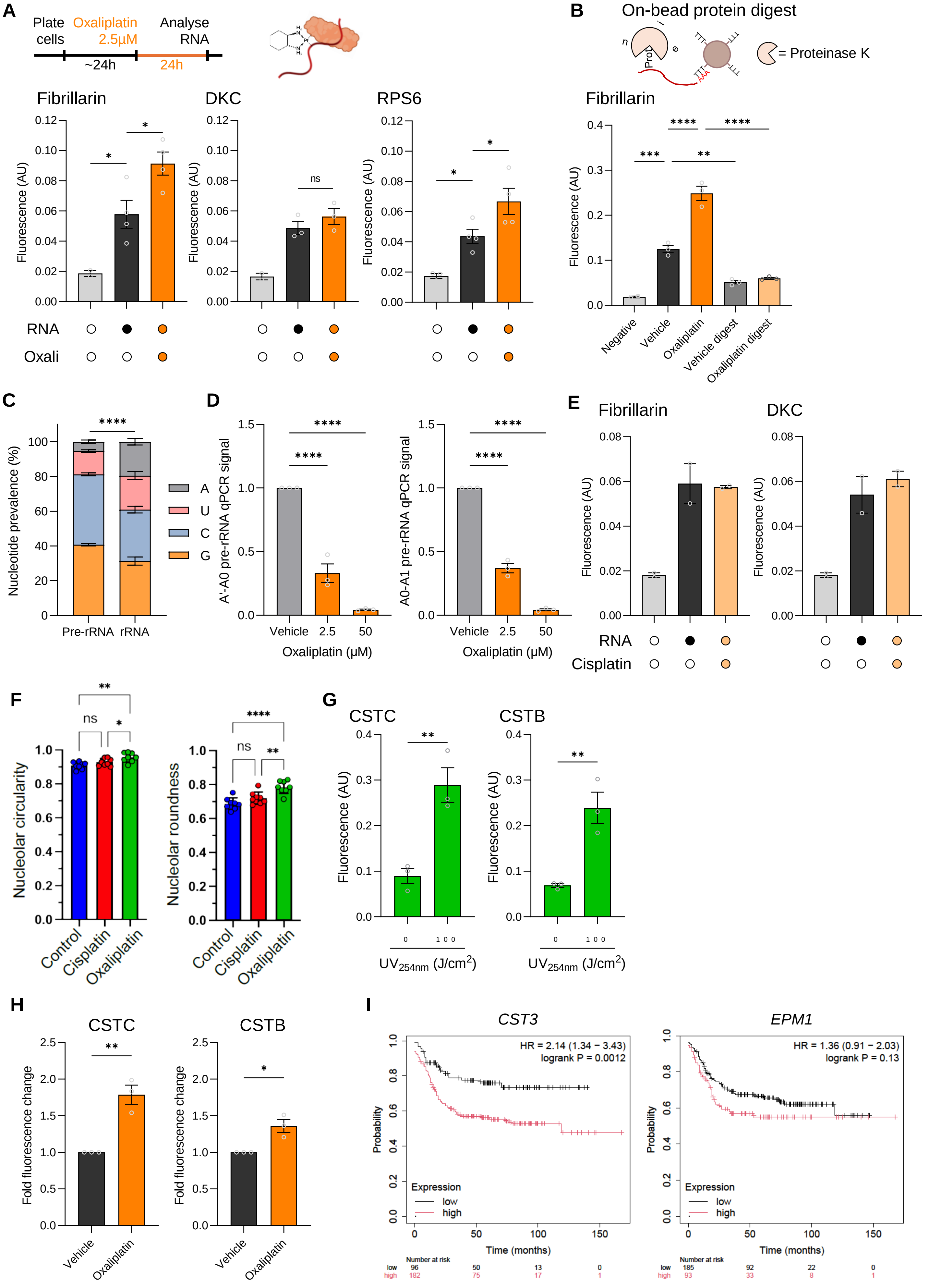

### Supplemental figure 7

#### Supplemental Figure 7

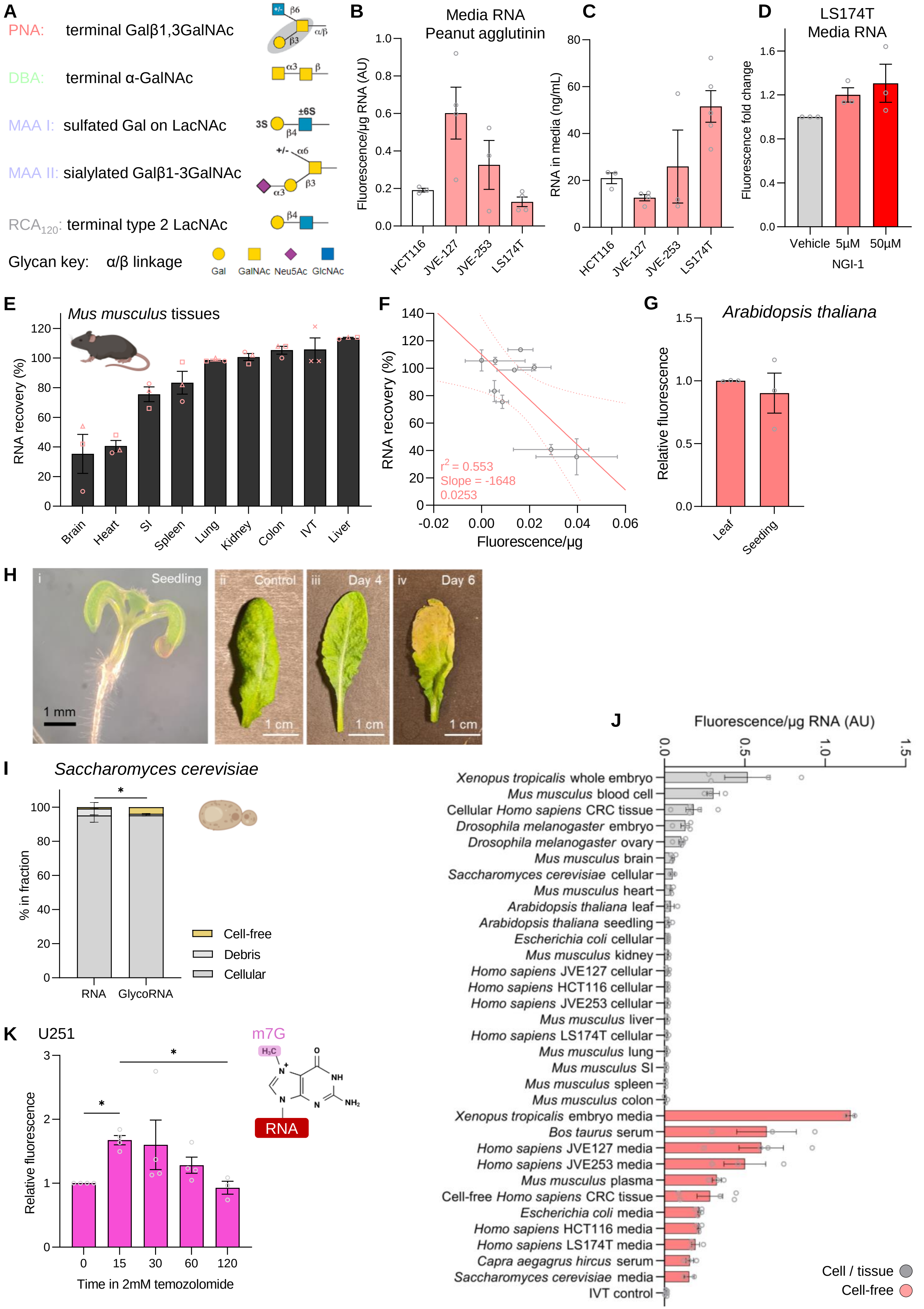
