## Supplemental table 1 for "AquIRE reveals multiple mechanisms of clinically induced RNA damage and the conservation and dynamics of glycoRNAs"

| Figure | Transcript | Affy ID | Data set | Cutoff used | Expression range | Upper quartile survival | | Hazard ratio | *P* value | FDR |
|---|---|---|---|---|---|---|---|---|---|---|
| | | | | | | Low | High | | | |
| Figure 4D | *DKC1* | 201479_at | Adjuvant | 2551 | 783 - 7984 | 13.0 | 27.6 | 0.55 | 2.8E-03 | 50% |
| Figure 4D | *DUS3L* | 224966_s_at | Adjuvant | 243 | 80 - 865 | 11.0 | 32.0 | 0.48 | 1.9E-04 | 3% |
| Supplementary Figure 5D | *TRMT2A* | 218475_at | Adjuvant | 108 | 4 - 355 | 22.0 | 15.0 | 1.40 | 8.6E-02 | 100% |
| Figure 5I | *CST3* | 201360_at | All cases | 3927 | 233 - 20826 | n/a | n/a | 2.06 | 3.2E-08 | 1% |
| Figure 5I | *EPM1* | 201201_at | All cases | 6034 | 1124 - 25430 | n/a | n/a | 1.76 | 5.3E-05 | 1% |
| Supplementary Figure 6I | *CST3* | 201360_at | Adjuvant | 5063 | 851 - 17795 | 70.5 | 14.0 | 2.14 | 1.2E-03 | 20% |
| Supplementary Figure 6I | *EPM1* | 201201_at | Adjuvant | 8922 | 2582 - 20693 | 22.4 | 16.0 | 1.36 | 1.3E-01 | 100% |

| Datasets: | n= |
|---|---|
| GSE12645 | 62 |
| GSE13294 | 155 |
| GSE14333 | 123 |
| GSE143985 | 91 |
| GSE17583 | 232 |
| GSE18088 | 53 |
| GSE26682 | 331 |
| GSE30540 | 35 |
| GSE31595 | 37 |
| GSE33114 | 90 |
| GSE34489 | 33 |
| GSE37892 | 62 |
| GSE38832 | 70 |
| GSE39582 | 514 |
| GSE41258 | 185 |
| GSE92921 | 59 |

Győrffy B: Integrated Analysis of Public Datasets for the Discovery and Validation of Survival-Associated Genes in Solid Tumors, **The Innovation**, 2024, DOI: 10.1016/j.xinn.2024.100625
