## Supplemental table 2 for "AquIRE reveals multiple mechanisms of clinically induced RNA damage and the conservation and dynamics of glycoRNAs"

| Accession | Description | Abundance Ratio (log2): (oxali) / (control) | Abundance Ratio Adj. P-Value: (oxali) / (control) | Abundance, Vehicle 1 | Abundance, Vehicle 2 | Abundance, Vehicle 3 | Abundance, Oxali 1 | Abundance, Oxali 2 | Abundance, Oxali 3 | Normalised abundances, Vehicle 1 | Normalised abundances, Vehicle 2 | Normalised abundances, Vehicle 3 | Normalised abundances, oxali 1 | Normalised abundances, oxali 2 | Normalised abundances, oxali 3 | Modifications |
|---|---|---|---|---|---|---|---|---|---|---|---|---|---|---|---|---|
| P01833 | Polymeric immunoglobulin receptor [OS=Homo sapiens] | 6.64 | 8.30303E-17 | | | | | 98831043.59 | 44581.17969 | | | | | 102668332 | 377254.0651 | |
| P24158 | Myeloblastin [OS=Homo sapiens] | 6.64 | 8.30303E-17 | | | | | 4054840.625 | | | | | | 4212276.915 | | |
| P00156 | Cytochrome b [OS=Homo sapiens] | 6.64 | 8.30303E-17 | | | 415063.125 | 47910.49219 | 450510.875 | | | | 415063.125 | 350218.6014 | 468002.7489 | | |
| P12429 | Annexin A3 [OS=Homo sapiens] | 6.64 | 8.30303E-17 | | | 236459.5781 | 60232.39844 | 4749389.438 | | | | 236459.5781 | 440289.911 | 4933792.802 | | Met-loss+Acetyl [N-Term] |
| A6NKN8 | Purkinje cell protein 4-like protein 1 [OS=Homo sapiens] | 6.64 | 8.30303E-17 | | | 2214424 | 182370.1719 | 1444177.75 | 455003.9375 | | | 2214424 | 1333098.944 | 1500250.481 | 3850326.219 | |
| P23280 | Carbonic anhydrase 6 [OS=Homo sapiens] | 6.64 | 8.30303E-17 | | | | | 34342059.78 | | | | | | 35675450.41 | | |
| P80303 | Nucleobindin-2 [OS=Homo sapiens] | 6.64 | 8.30303E-17 | | | | | 3631580.219 | | | | | | 3772582.682 | | |
| P00395 | Cytochrome c oxidase subunit 1 [OS=Homo sapiens] | 6.64 | 8.30303E-17 | | | 1339835.625 | 407712.5313 | 1199662.125 | | | | 1339835.625 | 2980318.213 | 1246241.108 | | |
| Q16378 | Proline-rich protein 4 [OS=Homo sapiens] | 6.64 | 8.30303E-17 | | | | | 1146703.875 | | | | | | 1191226.662 | | |
| Q99808 | Equilibrative nucleoside transporter 1 [OS=Homo sapiens] | 6.64 | 8.30303E-17 | 218717.3047 | | | 471732.1719 | 560489.0781 | 85734.60742 | 1478772.88 | | | 3448292.303 | 582251.0484 | 725501.8686 | |
| Q969E2 | Secretory carrier-associated membrane protein 4 [OS=Homo sapiens] | 6.64 | 8.30303E-17 | | | | 210910.3594 | | | | | | 1541723.487 | | | |
| Q969H8 | Myeloid-derived growth factor [OS=Homo sapiens] | 6.64 | 8.30303E-17 | | | 599240.3125 | 81856.71875 | 442498.3438 | 31346.42383 | | | 599240.3125 | 598360.4894 | 459679.1171 | 265259.1497 | |
| P01036 | Cystatin-S [OS=Homo sapiens] | 6.64 | 8.30303E-17 | | | | | 11185763.19 | | | | | | 11620070.04 | | |
| P15514 | Amphiregulin [OS=Homo sapiens] | 6.64 | 8.30303E-17 | 145117.375 | | | 434220.8438 | 430549.0313 | 175945.1094 | 981155.2814 | | | 3174090.051 | 447265.8516 | 1488879.572 | |
| P36405 | ADP-ribosylation factor-like protein 3 [OS=Homo sapiens] | 6.64 | 8.30303E-17 | | | | | 956800.0625 | | | | | | 993949.5009 | | |
| P11215 | Integrin alpha-M [OS=Homo sapiens] | 6.64 | 8.30303E-17 | | | | | 275832.0938 | | | | | | 286541.758 | | |
| P55058 | Phospholipid transfer protein [OS=Homo sapiens] | 6.64 | 8.30303E-17 | | | | 14612.00684 | 727170.75 | | | | | 106811.6056 | 755404.4281 | | |
| Q8TDL5 | BPI fold-containing family B member 1 [OS=Homo sapiens] | 6.64 | 8.30303E-17 | | | | | 2802299.5 | | | | | | 2911103.687 | | |
| P20061 | Transcobalamin-1 [OS=Homo sapiens] | 6.64 | 8.30303E-17 | | | | | 1550089.625 | | | | | | 1610274.57 | | |
| P26583 | High mobility group protein B2 [OS=Homo sapiens] | 6.64 | 8.30303E-17 | 45265.58594 | | | 41987.55078 | 1690007.281 | 44697.1582 | 306045.8385 | | | 306922.777 | 1755624.774 | 378235.4964 | |
| P06870 | Kallikrein-1 [OS=Homo sapiens] | 6.64 | 8.30303E-17 | | | | | 5362447.875 | | | | | | 5570654.307 | | |
| P0DOX7 | Immunoglobulin kappa light chain [OS=Homo sapiens] | 6.64 | 8.30303E-17 | | | | | 300521 | | | | | | 312189.2542 | | |
| Q6P5S2 | Protein LEG1 homolog [OS=Homo sapiens] | 6.64 | 8.30303E-17 | | | | | 35395624.81 | | | | | | 36769921.95 | | |
| P16422 | Epithelial cell adhesion molecule [OS=Homo sapiens] | 6.64 | 8.30303E-17 | | | | 91048.01563 | | | | | | 665547.5082 | | | |
| Q8N3C0 | Activating signal cointegrator 1 complex subunit 3 [OS=Homo sapiens] | 6.64 | 8.30303E-17 | | | | 133839.8281 | 480461.4063 | 252827.125 | | | | 978349.3192 | 499116.162 | 2139469.196 | |
| P08493 | Matrix Gla protein [OS=Homo sapiens] | 6.64 | 8.30303E-17 | | | | | 2349992.5 | | | | | | 2441235.075 | | |
| P04080 | Cystatin-B [OS=Homo sapiens] | 6.64 | 8.30303E-17 | | | | 164721.6484 | 23059267.06 | 46091.60156 | | | | 1204090.852 | 23954583.5 | 390035.5302 | |
| P17213 | Bactericidal permeability-increasing protein [OS=Homo sapiens] | 6.64 | 8.30303E-17 | | | | | 539071.0625 | | | | | | 560001.4408 | | |
| P60033 | CD81 antigen [OS=Homo sapiens] | 6.64 | 8.30303E-17 | | | | 2042332.656 | 2303770 | 194726.1094 | | | | 14929149.21 | 2393217.906 | 1647807.816 | |
| P11908 | Ribose-phosphate pyrophosphokinase 2 [OS=Homo sapiens] | 6.64 | 8.30303E-17 | | | 843660.0938 | 32665.43359 | 209798.5 | | | | 843660.0938 | 238779.4812 | 217944.2943 | | Met-loss [N-Term] |
| Q9H9B4 | Sideroflexin-1 [OS=Homo sapiens] | 6.64 | 8.30303E-17 | | | | | 506975.25 | | | | | | 526659.4521 | | |
| P31947 | 14-3-3 protein sigma [OS=Homo sapiens] | 6.64 | 8.30303E-17 | | | 122395.3125 | 125926.8984 | 5253925.125 | 113600.1484 | | | 122395.3125 | 920506.9753 | 5457917.971 | 961305.1537 | |
| P0DOX2 | Immunoglobulin alpha-2 heavy chain [OS=Homo sapiens] | 6.64 | 8.30303E-17 | | | | | 3916343.875 | | | | | | 4068402.786 | | |
| Q9UBC9 | Small proline-rich protein 3 [OS=Homo sapiens] | 6.64 | 8.30303E-17 | | | | | 520447.7188 | | | | | | 540655.0131 | | |
| Q9HC84 | Mucin-5B [OS=Homo sapiens] | 6.64 | 8.30303E-17 | | | | | 13338537 | | | | | | 13856429.06 | | |
| Q15269 | Periodic tryptophan protein 2 homolog [OS=Homo sapiens] | 6.64 | 8.30303E-17 | | | | 188207.125 | | | | | | 1375766.207 | | | |
| P09228 | Cystatin-SA [OS=Homo sapiens] | 6.64 | 8.30303E-17 | | | | | 1032279.625 | | | | | | 1072359.69 | | |
| P07711 | Procathepsin L [OS=Homo sapiens] | 6.64 | 8.30303E-17 | | | | 215603.5625 | 786837.9375 | 311444 | | | | 1576030.107 | 817388.2987 | 2635495.872 | |
| P17655 | Calpain-2 catalytic subunit [OS=Homo sapiens] | 6.64 | 8.30303E-17 | | | 1960435.469 | 109163.7344 | 1187732.781 | | | | 1960435.469 | 797970.729 | 1233848.587 | | |
| Q9Y253 | DNA polymerase eta [OS=Homo sapiens] | 6.64 | 8.30303E-17 | | | | | 349468.3438 | | | | | | 363037.0644 | | |
| Q6ZVX7 | F-box only protein 50 [OS=Homo sapiens] | 6.64 | 8.30303E-17 | 175577.7188 | | | | 307131.75 | 81381.47656 | 1187101.173 | | | | 319056.6782 | 688664.8821 | |
| Q7L266 | Isoaspartyl peptidase/L-asparaginase [OS=Homo sapiens] | 6.64 | 8.30303E-17 | | | | | 772505.8125 | | | | | | 802499.7039 | | |
| Q6P4E1 | Protein GOLM2 [OS=Homo sapiens] | 6.64 | 8.30303E-17 | | | | 20313.41211 | 815874.9375 | 40437.61719 | | | | 148488.0337 | 847552.7111 | 342190.4842 | |
| P41218 | Myeloid cell nuclear differentiation antigen [OS=Homo sapiens] | 6.64 | 8.30303E-17 | | | | | 3586268.688 | | | | | | 3725511.851 | | |
| Q96FQ6 | Protein S100-A16 [OS=Homo sapiens] | 6.64 | 8.30303E-17 | | | | | 370989.7188 | | | | | | 385394.0444 | | |
| P61769 | Beta-2-microglobulin [OS=Homo sapiens] | 6.64 | 8.30303E-17 | | | | | 3101472.5 | 30922.44922 | | | | | 3221892.603 | 261671.3993 | |
| P18510 | Interleukin-1 receptor antagonist protein [OS=Homo sapiens] | 6.64 | 8.30303E-17 | | | | | 354973.625 | | | | | | 368756.0978 | | |
| P13796 | Plastin-2 [OS=Homo sapiens] | 6.64 | 8.30303E-17 | | | | | 511368.9063 | | | | | | 531223.6998 | | |
| P34096 | Ribonuclease 4 [OS=Homo sapiens] | 6.64 | 8.30303E-17 | | | | | 828316.4375 | | | | | | 860477.2741 | | |
| P53990 | IST1 homolog [OS=Homo sapiens] | 6.64 | 8.30303E-17 | | | 1235810.688 | 26592.29492 | 795809.125 | | | | 1235810.688 | 194385.737 | 826707.8083 | | |
| Q12800 | Alpha-globin transcription factor CP2 [OS=Homo sapiens] | 6.64 | 8.30303E-17 | | | 372241.2813 | 69711.72656 | 241197.4063 | 66816.42969 | | | 372241.2813 | 509582.3955 | 250562.3181 | 565412.8018 | |
| O00566 | U3 small nucleolar ribonucleoprotein protein MPP10 [OS=Homo sapiens] | 6.64 | 8.30303E-17 | | | | | 433559.25 | 28537.20117 | | | | | 450392.9473 | 241486.9957 | |
| Q8N4F0 | BPI fold-containing family B member 2 [OS=Homo sapiens] | 6.64 | 8.30303E-17 | | | | | 6024847.688 | | | | | | 6258772.953 | | |
| P54108 | Cysteine-rich secretory protein 3 [OS=Homo sapiens] | 6.64 | 8.30303E-17 | | | | | 2016493.25 | | | | | | 2094787.133 | | |
| Q8TAX7 | Mucin-7 [OS=Homo sapiens] | 6.64 | 8.30303E-17 | | 22501.92578 | | | 205702088.5 | 84503.125 | | 161496.9344 | | | 213688832.4 | 715080.8399 | |
| O15127 | Secretory carrier-associated membrane protein 2 [OS=Homo sapiens] | 6.64 | 8.30303E-17 | 325262.9375 | | | 880560.5313 | 896730.6875 | 77701.79688 | 2199140.172 | | | 6436767.055 | 931547.8272 | 657526.7622 | |
| P52566 | Rho GDP-dissociation inhibitor 2 [OS=Homo sapiens] | 6.64 | 8.30303E-17 | | | | | 798191.8125 | | | | | | 829183.0078 | | |
| Q9Y2W1 | Thyroid hormone receptor-associated protein 3 [OS=Homo sapiens] | 6.64 | 8.30303E-17 | 48832.375 | | 512320.3438 | 94012.83594 | 234296.125 | 77974.35938 | 330161.31 | | 512320.3438 | 687219.9055 | 243393.0825 | 659833.2357 | |
| P08246 | Neutrophil elastase [OS=Homo sapiens] | 6.64 | 8.30303E-17 | | | | | 6064458 | 64645.40625 | | | | | 6299921.205 | 547041.2059 | |
| P02810 | Salivary acidic proline-rich phosphoprotein 1/2 [OS=Homo sapiens] | 6.64 | 8.30303E-17 | | | | | 32500085.59 | | | | | | 33761958.35 | | |
| P02790 | Hemopexin [OS=Homo sapiens] | 6.64 | 8.30303E-17 | | | | | 544650.625 | | | | | | 565797.6396 | | |
| Q9NRP0 | Oligosaccharyltransferase complex subunit OSTC [OS=Homo sapiens] | 6.64 | 8.30303E-17 | | | 432791.6563 | 147077.8438 | 423679.5938 | | | | 432791.6563 | 1075117.253 | 440129.6961 | | |
| P06312 | Immunoglobulin kappa variable 4-1 [OS=Homo sapiens] | 6.64 | 8.30303E-17 | | | | | 1132818.25 | | | | | | 1176801.903 | | |
| P14780 | Matrix metalloproteinase-9 [OS=Homo sapiens] | 6.64 | 8.30303E-17 | | | | | 1309799.938 | | | | | | 1360655.214 | | |
| P01591 | Immunoglobulin J chain [OS=Homo sapiens] | 6.64 | 8.30303E-17 | | | | | 19344701.56 | | | | | | 20095793.49 | | |
| Q9UKV3 | Apoptotic chromatin condensation inducer in the nucleus [OS=Homo sapier | 6.64 | 8.30303E-17 | | | 417359.125 | 51402.91406 | 214766.7656 | | | | 417359.125 | 375747.688 | 223105.4615 | | |
| Q9UM07 | Protein-arginine deiminase type-4 [OS=Homo sapiens] | 6.64 | 8.30303E-17 | | | | | 572381.4375 | | | | | | 594605.1494 | | |
| P35613 | Basigin [OS=Homo sapiens] | 6.64 | 8.30303E-17 | 836248.0938 | | | 472002.0625 | 834900.3906 | 638081.1563 | 5653969.649 | | | 3450265.164 | 867316.8608 | 5399559.001 | |
| P28325 | Cystatin-D [OS=Homo sapiens] | 6.64 | 8.30303E-17 | | | | | 6206352.25 | | | | | | 6447324.748 | | |
| P01824 | Immunoglobulin heavy variable 4-39 [OS=Homo sapiens] | 6.64 | 8.30303E-17 | | | | | 435561.0625 | | | | | | 452472.4837 | | |
| Q6FI81 | Anamorsin [OS=Homo sapiens] | 6.64 | 8.30303E-17 | | | 477999.25 | 55318.32031 | 316245.1563 | 22366.26758 | | | 477999.25 | 404368.7277 | 328523.9284 | 189267.4313 | |
| P20292 | Arachidonate 5-lipoxygenase-activating protein [OS=Homo sapiens] | 6.64 | 8.30303E-17 | | | | | 475680.4688 | | | | | | 494149.5962 | | |
| P08311 | Cathepsin G [OS=Homo sapiens] | 6.64 | 8.30303E-17 | | | | | 4215940.313 | | | | | | 4379631.58 | | |
| Q08380 | Galectin-3-binding protein [OS=Homo sapiens] | 6.64 | 8.30303E-17 | | | | | 2263980.344 | | | | | | 2351883.346 | | |
| Q8N983 | Large ribosomal subunit protein mL43 [OS=Homo sapiens] | 6.64 | 8.30303E-17 | 476990.2188 | | | 397273.7344 | 303819.9688 | 237328.1016 | 3224985.79 | | | 2904012.154 | 315616.3112 | 2008313.636 | |
| Q8N108 | Mesoderm induction early response protein 1 [OS=Homo sapiens] | 6.64 | 8.30303E-17 | 82854.44531 | | | 62554.41797 | 189954.4531 | 64701.48047 | 560188.4447 | | | 457263.5298 | 197329.7675 | 547515.7161 | |
| P05164 | Myeloperoxidase [OS=Homo sapiens] | 6.64 | 8.30303E-17 | | | | | 23377824.09 | | | | | | 24285509.06 | | |
| O75400 | Pre-mRNA-processing factor 40 homolog A [OS=Homo sapiens] | 6.64 | 8.30303E-17 | | | | 121811.9688 | 199775.9844 | 75701.41406 | | | | 890427.4488 | 207532.6369 | 640599.1584 | |
| P15515 | Histatin-1 [OS=Homo sapiens] | 6.64 | 8.30303E-17 | | | | | 2748947 | | | | | | 2855679.683 | | |

| | | | | | | | | | | | | | | | | |
|---|---|---|---|---|---|---|---|---|---|---|---|---|---|---|---|---|
| P0DOX8 | Immunoglobulin lambda-1 light chain [OS=Homo sapiens] | 6.64 | 8.30303E-17 | | | | | 3767947.438 | 13535.44043 | | | | | 3914244.597 | 114539.363 | |
| Q99816 | Tumor susceptibility gene 101 protein [OS=Homo sapiens] | 6.64 | 8.30303E-17 | | | 296951.875 | 61933.42578 | 586640.9063 | 60183.05859 | | | 296951.875 | 452724.1689 | 609418.267 | 509280.007 | |
| P10909 | Clusterin [OS=Homo sapiens] | 6.64 | 8.30303E-17 | | | | 32986.32422 | 979063.8125 | | | | | 241125.1441 | 1017077.68 | | |
| P80188 | Neutrophil gelatinase-associated lipocalin [OS=Homo sapiens] | 6.64 | 8.30303E-17 | | | | | 4040681.938 | | | | | | 4197568.492 | | |
| P01037 | Cystatin-SN [OS=Homo sapiens] | 6.57 | 1.30156E-06 | | 127556.0938 | 333227.2188 | 173835.9688 | 115480238 | 484703.9004 | | 915473.56 | 333227.2188 | 1270715.183 | 119963960.5 | 4101652.717 | |
| Q96DA0 | Pancreatic adenocarcinoma up-regulated factor [OS=Homo sapiens] | 6.56 | 5.88968E-07 | 23185.41016 | 121701.2266 | | 149179.6133 | 257370786.3 | 252386.5234 | 156759.2277 | 873453.0187 | | 1090480.877 | 267363658 | 2135740.745 | |
| P01024 | Complement C3 [OS=Homo sapiens] | 4.33 | 0.007408981 | | 102888.2891 | | 86817.01172 | 5523995.438 | | | 738432.0537 | | 634619.4963 | 5738474.237 | | |
| P17931 | Galectin-3 [OS=Homo sapiens] | 4.33 | 0.011374485 | | | 186237.3906 | | 620790.3125 | 89027.25 | | | 186237.3906 | | 644893.5838 | 753364.81 | |
| P01034 | Cystatin-C [OS=Homo sapiens] | 4.14 | 0.000211121 | 243109.7656 | 30343.33008 | | 158678.4219 | 9675618.125 | 130025.5469 | 1643693.117 | 217774.9067 | | 1159915.76 | 10051290.95 | 1100299.868 | |
| O15118 | NPC intracellular cholesterol transporter 1 [OS=Homo sapiens] | 4.01 | 0.0305647 | 187555.1914 | | | 433929.1875 | 270065.4688 | | 1268082.244 | | | 3171958.087 | 280551.2336 | | |
| Q9H1C7 | Cysteine-rich and transmembrane domain-containing protein 1 [OS=Homo | 3.94 | 0.007672891 | 104985.0625 | | | 229413.9844 | 507685.5625 | | 709816.0957 | | | 1676982.245 | 527397.3438 | | |
| Q96G23 | Ceramide synthase 2 [OS=Homo sapiens] | 3.91 | 0.009916497 | 169716.6563 | | | 355567.7813 | | 79022.52344 | 1147473.854 | | | 2599147.814 | | 668702.991 | |
| O75882 | Attractin [OS=Homo sapiens] | 3.61 | 0.002091393 | 37571.91406 | | | 51973.02734 | | 21002.81055 | 254028.0371 | | | 379915.1316 | | 177729.6095 | |
| P13010 | X-ray repair cross-complementing protein 5 [OS=Homo sapiens] | 3.58 | 0.001547471 | 120377.8828 | | | 159797.125 | | 37107.58594 | 813888.7262 | | | 1168093.315 | | 314011.1531 | |
| P12110 | Collagen alpha-2(VI) chain [OS=Homo sapiens] | 3.52 | 0.002028446 | 113401 | | | 138310.8594 | | 99698.60156 | 766717.2181 | | | 1011031.895 | | 843667.7312 | |
| O15511 | Actin-related protein 2/3 complex subunit 5 [OS=Homo sapiens] | 3.34 | 0.004234111 | 49895.65234 | | 365096.2188 | 47084.68359 | 2370826.875 | 59911.11328 | 337350.2506 | | 365096.2188 | 344182.0629 | 2462878.381 | 506978.7562 | |
| Q96PU5 | E3 ubiquitin-protein ligase NEDD4-like [OS=Homo sapiens] | 3.31 | 0.034964583 | | | 287909.5 | | 252728.9531 | 33280.92578 | | | 287909.5 | | 262541.5974 | 281629.2576 | |
| Q8N9T8 | Protein KRI1 homolog [OS=Homo sapiens] | 3.19 | 0.002767664 | 80650.71094 | | | 62445.82422 | 312030.2188 | | 545288.7429 | | | 456469.7256 | 324145.3386 | | |
| Q13595 | Transformer-2 protein homolog alpha [OS=Homo sapiens] | 3.17 | 0.004295584 | 128777.0938 | | | 95889.75781 | | 86825.85156 | 870676.7584 | | | 700939.9264 | | 734736.1753 | |
| Q8N573 | Oxidation resistance protein 1 [OS=Homo sapiens] | -3.93 | 0.048634869 | | 64569.30078 | 371403.0938 | | 191288.5781 | | | 463415.6309 | 371403.0938 | | 198715.6922 | | |
| Q96FZ7 | Charged multivesicular body protein 6 [OS=Homo sapiens] | -3.97 | 0.040959529 | | 73762.82813 | 347036.8438 | | 208938.3906 | | | 529397.8271 | 347036.8438 | | 217050.7896 | | |
| P23763 | Vesicle-associated membrane protein 1 [OS=Homo sapiens] | -6.64 | 8.30303E-17 | | 235997.9531 | 986835.75 | | | | | 1693763.739 | 986835.75 | | | | |
| Q9UBE0 | SUMO-activating enzyme subunit 1 [OS=Homo sapiens] | -6.64 | 8.30303E-17 | | 57717.83594 | 428130.4063 | | | | | 414242.481 | 428130.4063 | | | | |
| Q8IWB7 | WD repeat and FYVE domain-containing protein 1 [OS=Homo sapiens] | -6.64 | 8.30303E-17 | 20272.83398 | 262661.0781 | 4186995 | | | | 137066.9648 | 1885125.714 | 4186995 | | | | Met-loss+Acetyl [N-Term] |
| Q96PU8 | KH domain-containing RNA-binding protein QKI [OS=Homo sapiens] | -6.64 | 8.30303E-17 | | 72764.32031 | 443744.1563 | | | | | 522231.5093 | 443744.1563 | | | | |
| O94925 | Glutaminase kidney isoform, mitochondrial [OS=Homo sapiens] | -6.64 | 8.30303E-17 | 175094.9688 | 2065057.523 | 27482266.47 | | 14445177.38 | | 1183837.244 | 14820974.11 | 27482266.47 | | 15006036.69 | | |
| Q9Y6R0 | Numb-like protein [OS=Homo sapiens] | -6.64 | 8.30303E-17 | | | 460706.625 | | | | | | 460706.625 | | | | |
| Q99615 | DnaJ homolog subfamily C member 7 [OS=Homo sapiens] | -6.64 | 8.30303E-17 | | | 251884.3594 | | | | | | 251884.3594 | | | | |
| Q8IW45 | ATP-dependent (S)-NAD(P)H-hydrate dehydratase [OS=Homo sapiens] | -6.64 | 8.30303E-17 | | | 440928.75 | | | | | | 440928.75 | | | | |
| Q9Y2Z0 | Protein SGT1 homolog [OS=Homo sapiens] | -6.64 | 8.30303E-17 | | | 938639.1875 | | | | | | 938639.1875 | | | | |
| Q9NYI0 | PH and SEC7 domain-containing protein 3 [OS=Homo sapiens] | -6.64 | 8.30303E-17 | 22350.28516 | 308314.1484 | 2880242.516 | | 2643710.594 | | 151112.8514 | 2212779.043 | 2880242.516 | | 2746357.289 | | |
| Q15843 | NEDD8 [OS=Homo sapiens] | -6.64 | 8.30303E-17 | 85766.50781 | 338324.4063 | 5309243.063 | | 3748451.906 | | 579877.2346 | 2428163.481 | 5309243.063 | | 3893992.118 | | |
| O15083 | ERC protein 2 [OS=Homo sapiens] | -6.64 | 8.30303E-17 | 182386.1563 | 37415.3125 | 1167069.719 | | 786159.8438 | | 1233133.803 | 268530.7172 | 1167069.719 | | 816683.8768 | | |
| P46109 | Crk-like protein [OS=Homo sapiens] | -6.64 | 8.30303E-17 | | 110946.4453 | 1336769.375 | | | | | 796265.6607 | 1336769.375 | | | | |
| Q9H0C8 | Integrin-linked kinase-associated serine/threonine phosphatase 2C [OS=Ho | -6.64 | 8.30303E-17 | | | 261635.4844 | | | | | | 261635.4844 | | | | |
| Q13620 | Cullin-4B [OS=Homo sapiens] | -6.64 | 8.30303E-17 | 17690.4375 | 34699.92969 | 611011.75 | | 460200.5313 | | 119607.0848 | 249042.3407 | 611011.75 | | 478068.623 | | |
| P40939 | Trifunctional enzyme subunit alpha, mitochondrial [OS=Homo sapiens] | -6.64 | 8.30303E-17 | 198105.4453 | 1620036.75 | 11201085.31 | | 9455975.25 | | 1339413.726 | 11627047.89 | 11201085.31 | | 9823120.053 | | |
| P61018 | Ras-related protein Rab-4B [OS=Homo sapiens] | -6.64 | 8.30303E-17 | | 30408.80273 | 500190.375 | | | | | 218244.8057 | 500190.375 | | | | |
| P61221 | ATP-binding cassette sub-family E member 1 [OS=Homo sapiens] | -6.64 | 8.30303E-17 | | 65758.39063 | 448678.7813 | | | | | 471949.7611 | 448678.7813 | | | | |
| O00194 | Ras-related protein Rab-27B [OS=Homo sapiens] | -6.64 | 8.30303E-17 | 87469.1875 | 100198.8359 | 721477.625 | | 772903.1875 | | 591389.248 | 719129.7754 | 721477.625 | | 802912.5076 | | |
| P42167 | Lamina-associated polypeptide 2, isoforms beta/gamma [OS=Homo sapier | -6.64 | 8.30303E-17 | 114849.3828 | 503616.6328 | 3287283.328 | | 1899220.094 | | 776509.9011 | 3614470.294 | 3287283.328 | | 1972960.641 | | Met-loss [N-Term] |
| Q9H254 | Spectrin beta chain, non-erythrocytic 4 [OS=Homo sapiens] | -6.64 | 8.30303E-17 | | 18522.01758 | 340949.7813 | | | | | 132933.0248 | 340949.7813 | | | | |
| Q9Y3E7 | Charged multivesicular body protein 3 [OS=Homo sapiens] | -6.64 | 8.30303E-17 | | | 386384.6875 | | | | | | 386384.6875 | | | | |
| Q8N6C5 | Immunoglobulin superfamily member 1 [OS=Homo sapiens] | -6.64 | 8.30303E-17 | | 27615.34961 | 350820.4375 | | | | | 198196.1165 | 350820.4375 | | | | |
| Q6YN16 | Hydroxysteroid dehydrogenase-like protein 2 [OS=Homo sapiens] | -6.64 | 8.30303E-17 | | | 311431.9688 | | | | | | 311431.9688 | | | | |
| Q96T17 | MAP7 domain-containing protein 2 [OS=Homo sapiens] | -6.64 | 8.30303E-17 | | | 284593.125 | | | | | | 284593.125 | | | | |
| Q9Y2W6 | Tudor and KH domain-containing protein [OS=Homo sapiens] | -6.64 | 8.30303E-17 | | 35109.37891 | 320567.5313 | | | | | 251980.9689 | 320567.5313 | | | | |
| P15374 | Ubiquitin carboxyl-terminal hydrolase isozyme L3 [OS=Homo sapiens] | -6.64 | 8.30303E-17 | | 124445.0469 | 832619.0625 | | 1764780.375 | | | 893145.4918 | 832619.0625 | | 1833301.064 | | |
| Q9P1U1 | Actin-related protein 3B [OS=Homo sapiens] | -6.64 | 8.30303E-17 | | | 383220 | | | | | | 383220 | | | | |
| Q15819 | Ubiquitin-conjugating enzyme E2 variant 2 [OS=Homo sapiens] | -6.64 | 8.30303E-17 | 369649.3125 | 785608.125 | 8600964.563 | | 6002823.125 | | 2499241.564 | 5638330.917 | 8600964.563 | | 6235893.248 | | |
| Q13492 | Phosphatidylinositol-binding clathrin assembly protein [OS=Homo sapiens] | -6.64 | 8.30303E-17 | | 58712.76563 | 662153.125 | | 301629.125 | | | 421383.1185 | 662153.125 | | 313340.4041 | | |
| O43924 | Retinal rod rhodopsin-sensitive cGMP 3',5'-cyclic phosphodiesterase subun | -6.64 | 8.30303E-17 | | | 327546.375 | | | | | | 327546.375 | | | | |
| Q5TH69 | Brefeldin A-inhibited guanine nucleotide-exchange protein 3 [OS=Homo sap | -6.64 | 8.30303E-17 | | 65844.09375 | 375514 | | | | | 472564.8547 | 375514 | | | | |
| Q05086 | Ubiquitin-protein ligase E3A [OS=Homo sapiens] | -6.64 | 8.30303E-17 | | | 379797.2813 | | | | | | 379797.2813 | | | | |
| Q96TC7 | Regulator of microtubule dynamics protein 3 [OS=Homo sapiens] | -6.64 | 8.30303E-17 | | 70265.01563 | 717247.3125 | | | | | 504293.9315 | 717247.3125 | | | | |
| O60861 | Growth arrest-specific protein 7 [OS=Homo sapiens] | -6.64 | 8.30303E-17 | | | 529169.1875 | | | | | | 529169.1875 | | | | |
| P31146 | Coronin-1A [OS=Homo sapiens] | -6.64 | 8.30303E-17 | 21082.81641 | 807106.0938 | 7463758.781 | | 7460975.188 | | 142543.3492 | 5792622.424 | 7463758.781 | | 7750660.618 | | |
| Q7L0J3 | Synaptic vesicle glycoprotein 2A [OS=Homo sapiens] | -6.64 | 8.30303E-17 | 58004.70313 | 3831744.279 | 38983684.13 | | 27916629.69 | | 392176.4766 | 27500533.09 | 38983684.13 | | 29000541.74 | | |
| Q00688 | Peptidyl-prolyl cis-trans isomerase FKBP3 [OS=Homo sapiens] | -6.64 | 8.30303E-17 | | | 235024.5 | | | | | | 235024.5 | | | | |
| Q9UM54 | Unconventional myosin-VI [OS=Homo sapiens] | -6.64 | 8.30303E-17 | | | 413701.3125 | | | | | | 413701.3125 | | | | |
| O75077 | Disintegrin and metalloproteinase domain-containing protein 23 [OS=Homo | -6.64 | 8.30303E-17 | | 68146.13281 | 2996195.938 | | 621052.7031 | | | 489086.6518 | 2996195.938 | | 645166.1622 | | |
| O75569 | Interferon-inducible double-stranded RNA-dependent protein kinase activat | -6.64 | 8.30303E-17 | | 55313.20703 | 619310.0469 | | | | | 396984.3938 | 619310.0469 | | | | |
| Q6KCM7 | Mitochondrial adenyl nucleotide antiporter SLC25A25 [OS=Homo sapiens] | -6.64 | 8.30303E-17 | 4105720.75 | 344181.4688 | 2199803.125 | | 5564256.781 | | 27759250.73 | 2470199.778 | 2199803.125 | | 5780298.798 | | |
| Q96FE5 | Leucine-rich repeat and immunoglobulin-like domain-containing nogo recep | -6.64 | 8.30303E-17 | | | 384031.25 | | | | | | 384031.25 | | | | |
| Q9UJU6 | Drebrin-like protein [OS=Homo sapiens] | -6.64 | 8.30303E-17 | 13060.91992 | 80331.03125 | 1818783.281 | | 1172175.578 | | 88306.38344 | 576537.9998 | 1818783.281 | | 1217687.348 | | |
| P52565 | Rho GDP-dissociation inhibitor 1 [OS=Homo sapiens] | -6.64 | 8.30303E-17 | 31777.44336 | 3063744.945 | 30650163.38 | | 21524304.09 | | 214850.9535 | 21988580.94 | 30650163.38 | | 22360022.9 | | Met-loss+Acetyl [N-Term] |
| P16870 | Carboxypeptidase E [OS=Homo sapiens] | -6.64 | 8.30303E-17 | 102321.6328 | 1364034.711 | 12865777.91 | | 12515557.06 | | 691808.3409 | 9789714.283 | 12865777.91 | | 13001495.49 | | |
| Q14498 | RNA-binding protein 39 [OS=Homo sapiens] | -6.64 | 8.30303E-17 | | | 330648.0313 | | | | | | 330648.0313 | | | | |
| P62306 | Small nuclear ribonucleoprotein F [OS=Homo sapiens] | -6.64 | 8.30303E-17 | | 42987.30469 | 350921.4063 | | | | | 308521.0569 | 350921.4063 | | | | |
| Q9UJW0 | Dynactin subunit 4 [OS=Homo sapiens] | -6.64 | 8.30303E-17 | | | 291665.2188 | | | | | | 291665.2188 | | | | |
| P15954 | Cytochrome c oxidase subunit 7C, mitochondrial [OS=Homo sapiens] | -6.64 | 8.30303E-17 | 114078.2266 | 935552.1875 | 5336669 | | 4314861 | | 771296.0249 | 6714483.538 | 5336669 | | 4482393.037 | | |
| Q9NZJ7 | Mitochondrial carrier homolog 1 [OS=Homo sapiens] | -6.64 | 8.30303E-17 | 34174.64063 | 587764.8691 | 6027627.438 | | 5048440.344 | | 231058.6802 | 4218404.479 | 6027627.438 | | 5244454.883 | | |
| P22061 | Protein-L-isoaspartate(D-aspartate) O-methyltransferase [OS=Homo sapier | -6.64 | 8.30303E-17 | 31291.16602 | 547060.5781 | 9876562.469 | | 8439912.031 | | 211563.1763 | 3926268.674 | 9876562.469 | | 8767606.4 | | |
| P23258 | Tubulin gamma-1 chain [OS=Homo sapiens] | -6.64 | 8.30303E-17 | | | 305127.7813 | | | | | | 305127.7813 | | | | |
| Q86YM7 | Homer protein homolog 1 [OS=Homo sapiens] | -6.64 | 8.30303E-17 | | | 342242.5 | | | | | | 342242.5 | | | | |
| Q9Y3B8 | Oligoribonuclease, mitochondrial [OS=Homo sapiens] | -6.64 | 8.30303E-17 | | | 529719.125 | | | | | | 529719.125 | | | | |
| P23434 | Glycine cleavage system H protein, mitochondrial [OS=Homo sapiens] | -6.64 | 8.30303E-17 | | | 503849.4063 | | | | | | 503849.4063 | | | | |
| Q8WVM8 | Sec1 family domain-containing protein 1 [OS=Homo sapiens] | -6.64 | 8.30303E-17 | | 10290.80566 | 734019.25 | | 347482.5 | | | 73857.39262 | 734019.25 | | 360974.1167 | | |
| P48147 | Prolyl endopeptidase [OS=Homo sapiens] | -6.64 | 8.30303E-17 | | | 935086.1563 | | | | | | 935086.1563 | | | | |
| O00233 | 26S proteasome non-ATPase regulatory subunit 9 [OS=Homo sapiens] | -6.64 | 8.30303E-17 | 17491.79297 | 63108.57031 | 344722.625 | | 361991.375 | | 118264.0263 | 452931.9285 | 344722.625 | | 376046.3242 | | |
| P22079 | Lactoperoxidase [OS=Homo sapiens] | -6.64 | 8.30303E-17 | 18959.7168 | 124549.7734 | 254621.375 | | 25882815.81 | | 128188.8283 | 893897.1172 | 254621.375 | | 26887761.48 | | |
| P62166 | Neuronal calcium sensor 1 [OS=Homo sapiens] | -6.64 | 8.30303E-17 | | | 298099.6563 | | | | | | 298099.6563 | | | | |

| | | | | | | | | | | | | | | | | |
|---|---|---|---|---|---|---|---|---|---|---|---|---|---|---|---|---|
| P22059 | Oxysterol-binding protein 1 [OS=Homo sapiens] | -6.64 | 8.30303E-17 | | 64961.88281 | 960516.75 | | | | | 466233.2028 | 960516.75 | | | | |
| O60218 | Aldo-keto reductase family 1 member B10 [OS=Homo sapiens] | -6.64 | 8.30303E-17 | 60956.08984 | 1805879.32 | 761630.5625 | | 637407.25 | | 412131.142 | 12960845.08 | 761630.5625 | | 662155.703 | | |
| P46060 | Ran GTPase-activating protein 1 [OS=Homo sapiens] | -6.64 | 8.30303E-17 | | | 792402.1875 | | | | | | 792402.1875 | | | | |
| Q00537 | Cyclin-dependent kinase 17 [OS=Homo sapiens] | -6.64 | 8.30303E-17 | | | 381691.9063 | | | | | | 381691.9063 | | | | |
| P34897 | Serine hydroxymethyltransferase, mitochondrial [OS=Homo sapiens] | -6.64 | 8.30303E-17 | 240415.3281 | 145048.6406 | 312959.6563 | | | 135325.875 | 1625475.715 | 1041018.046 | 312959.6563 | | | 1145152.21 | |
| Q14576 | ELAV-like protein 3 [OS=Homo sapiens] | -6.64 | 8.30303E-17 | | | 251561.375 | | | | | | 251561.375 | | | | |
| Q9UBQ5 | Eukaryotic translation initiation factor 3 subunit K [OS=Homo sapiens] | -6.64 | 8.30303E-17 | | 44341.77734 | 389193.8438 | | | | | 318242.1441 | 389193.8438 | | | | |
| Q8WXG6 | MAP kinase-activating death domain protein [OS=Homo sapiens] | -6.64 | 8.30303E-17 | | | 434759.4063 | | | | | | 434759.4063 | | | | |
| P34949 | Mannose-6-phosphate isomerase [OS=Homo sapiens] | -6.64 | 8.30303E-17 | | | 979404.5 | | | | | | 979404.5 | | | | |
| Q96EP5 | DAZ-associated protein 1 [OS=Homo sapiens] | -6.64 | 8.30303E-17 | | | 680554.125 | | | | | | 680554.125 | | | | |
| Q15121 | Astrocytic phosphoprotein PEA-15 [OS=Homo sapiens] | -6.64 | 8.30303E-17 | 38384.76953 | 456134.1133 | 16734866.31 | | 5974732.625 | | 259523.8465 | 3273686.958 | 16734866.31 | | 6206712.085 | | |
| O43237 | Cytoplasmic dynein 1 light intermediate chain 2 [OS=Homo sapiens] | -6.64 | 8.30303E-17 | | | 373481.7813 | | | | | | 373481.7813 | | | | |
| P61163 | Alpha-centractin [OS=Homo sapiens] | -6.64 | 8.30303E-17 | 41837.91406 | 1155888.945 | 14561058.69 | | 10311278.34 | | 282870.9543 | 8295846.452 | 14561058.69 | | 10711631.79 | | |
| Q14697 | Neutral alpha-glucosidase AB [OS=Homo sapiens] | -6.64 | 8.30303E-17 | 43440.72656 | 247585.1289 | 3131755.281 | | 2371289.25 | | 293707.7541 | 1776925.215 | 3131755.281 | | 2463358.709 | | |
| Q8TCD5 | 5'(3')-deoxyribonucleotidase, cytosolic type [OS=Homo sapiens] | -6.64 | 8.30303E-17 | 19433.82813 | 80210.65234 | 963021.6875 | | 327365.4375 | | 131394.3496 | 575674.0372 | 963021.6875 | | 340075.9741 | | |
| Q9Y584 | Mitochondrial import inner membrane translocase subunit Tim22 [OS=Hom | -6.64 | 8.30303E-17 | | | 335287.125 | | | | | | 335287.125 | | | | |
| Q9BY11 | Protein kinase C and casein kinase substrate in neurons protein 1 [OS=Hom | -6.64 | 8.30303E-17 | 44195.96094 | 3284163.961 | 26009705.69 | | 11876175.19 | | 298813.9807 | 23570534.22 | 26009705.69 | | 12337288.49 | | |
| Q13740 | CD166 antigen [OS=Homo sapiens] | -6.64 | 8.30303E-17 | 37049.78516 | 620033.0781 | 7117514.031 | | 4739549.281 | | 250497.8634 | 4449994.294 | 7117514.031 | | 4923570.584 | | |
| O14917 | Protocadherin-17 [OS=Homo sapiens] | -6.64 | 8.30303E-17 | | | 177153.3594 | | | | | | 177153.3594 | | | | |
| P17096 | High mobility group protein HMG-I/HMG-Y [OS=Homo sapiens] | -6.64 | 8.30303E-17 | 40671.14063 | 111414.8672 | 2317512.5 | | 1497709.875 | | 274982.2647 | 799627.5372 | 2317512.5 | | 1555861.084 | | |
| P46779 | Large ribosomal subunit protein eL28 [OS=Homo sapiens] | -6.64 | 8.30303E-17 | | | 394756.9063 | | | | | | 394756.9063 | | | | Met-loss+Acetyl [N-Term] |
| P61086 | Ubiquitin-conjugating enzyme E2 K [OS=Homo sapiens] | -6.64 | 8.30303E-17 | | 68473.00781 | 1475952.063 | | 395836.4375 | | | 491432.6426 | 1475952.063 | | 411205.4806 | | |
| P53618 | Coatomer subunit beta [OS=Homo sapiens] | -6.64 | 8.30303E-17 | 23030.63086 | 265392.2266 | 2108645.438 | | 1603125.344 | | 155712.747 | 1904727.24 | 2108645.438 | | 1665369.493 | | |
| Q96DD7 | Protein shisa-4 [OS=Homo sapiens] | -6.64 | 8.30303E-17 | | | 390650.0313 | | | | | | 390650.0313 | | | | |
| Q96BM9 | ADP-ribosylation factor-like protein 8A [OS=Homo sapiens] | -6.64 | 8.30303E-17 | | | 222834.6875 | | | | | | 222834.6875 | | | | |
| Q15042 | Rab3 GTPase-activating protein catalytic subunit [OS=Homo sapiens] | -6.64 | 8.30303E-17 | | 94246.90625 | 982440.5938 | | 208167.625 | | | 676412.6139 | 982440.5938 | | 216250.0977 | | |
| Q9UBT2 | SUMO-activating enzyme subunit 2 [OS=Homo sapiens] | -6.64 | 8.30303E-17 | 66996.35938 | 128287.4688 | 1803333.375 | | 1548051.938 | | 452970.0998 | 920722.6583 | 1803333.375 | | 1608157.766 | | |
| Q9NQG5 | Regulation of nuclear pre-mRNA domain-containing protein 1B [OS=Homo s | -6.64 | 8.30303E-17 | | | 167514.1719 | | | | | | 167514.1719 | | | | |
| P48163 | NADP-dependent malic enzyme [OS=Homo sapiens] | -6.64 | 8.30303E-17 | | 128039.8516 | 1010341.375 | | | | | 918945.503 | 1010341.375 | | | | |
| Q92499 | ATP-dependent RNA helicase DDX1 [OS=Homo sapiens] | -6.64 | 8.30303E-17 | 81176.28906 | 306276.9844 | 4294803.188 | | 1354126.609 | | 548842.2371 | 2198158.261 | 4294803.188 | | 1406702.947 | | |
| Q7RTV2 | Glutathione S-transferase A5 [OS=Homo sapiens] | -6.64 | 8.30303E-17 | 17966.21484 | 587509.8438 | 3152228.875 | | 2476053.25 | | 121471.6472 | 4216574.155 | 3152228.875 | | 2572190.355 | | |
| P09471 | Guanine nucleotide-binding protein G(o) subunit alpha [OS=Homo sapiens] | -6.64 | 8.30303E-17 | 43571.72656 | 16841372.14 | 140795684.1 | | 104068586.3 | 68458.18848 | 294593.4602 | 120870986.7 | 140795684.1 | | 108109231.4 | 579305.6638 | |
| O95831 | Apoptosis-inducing factor 1, mitochondrial [OS=Homo sapiens] | -6.64 | 8.30303E-17 | | | 633301.0625 | | | | | | 633301.0625 | | | | |
| Q9UF56 | F-box/LRR-repeat protein 17 [OS=Homo sapiens] | -6.64 | 8.30303E-17 | | | 509426.6563 | | | | | | 509426.6563 | | | | |
| Q96F85 | CB1 cannabinoid receptor-interacting protein 1 [OS=Homo sapiens] | -6.64 | 8.30303E-17 | 54888.14063 | 347064.375 | 2434054.625 | | 1924159.125 | | 371105.0387 | 2490890.475 | 2434054.625 | | 1998867.974 | | |
| Q14257 | Reticulocalbin-2 [OS=Homo sapiens] | -6.64 | 8.30303E-17 | | 80601.39063 | 1292330.875 | | | | | 578478.3764 | 1292330.875 | | | | |
| P49721 | Proteasome subunit beta type-2 [OS=Homo sapiens] | -6.64 | 8.30303E-17 | 23434.45313 | 413394.4141 | 3197211.688 | | 2874802.063 | | 158443.0359 | 2966942.972 | 3197211.688 | | 2986421.288 | | |
| O75145 | Liprin-alpha-3 [OS=Homo sapiens] | -6.64 | 8.30303E-17 | | | 1010439.25 | | | | | | 1010439.25 | | | | |
| P11532 | Dystrophin [OS=Homo sapiens] | -6.64 | 8.30303E-17 | | | 237119.9219 | | | | | | 237119.9219 | | | | |
| P42704 | Leucine-rich PPR motif-containing protein, mitochondrial [OS=Homo sapien | -6.64 | 8.30303E-17 | | 94297.67188 | 729888.8125 | | | | | 676776.9601 | 729888.8125 | | | | |
| Q8NFW8 | N-acylneuraminate cytidylyltransferase [OS=Homo sapiens] | -6.64 | 8.30303E-17 | 40350.87109 | 704315.0977 | 3943490.031 | | 3166289.063 | | 272816.8855 | 5054888.644 | 3943490.031 | | 3289225.782 | | |
| Q5T5C0 | Syntaxin-binding protein 5 [OS=Homo sapiens] | -6.64 | 8.30303E-17 | 112800.4531 | 180834.9844 | 2574465.406 | | 1379986.281 | | 762656.8516 | 1297857.61 | 2574465.406 | | 1433566.666 | | |
| O14980 | Exportin-1 [OS=Homo sapiens] | -6.64 | 8.30303E-17 | 33545.24219 | 607524.4023 | 5519217.406 | | 4378803.641 | | 226803.2449 | 4360219.187 | 5519217.406 | | 4548818.362 | | |
| Q9NZQ3 | NCK-interacting protein with SH3 domain [OS=Homo sapiens] | -6.64 | 8.30303E-17 | | | 465068.0625 | | | | | | 465068.0625 | | | | |
| Q09666 | Neuroblast differentiation-associated protein AHNAK [OS=Homo sapiens] | -6.64 | 8.30303E-17 | | | 500194.0938 | | | | | | 500194.0938 | | | | |
| Q9HCK5 | Protein argonaute-4 [OS=Homo sapiens] | -6.64 | 8.30303E-17 | | | 323570.5 | | | | | | 323570.5 | | | | |
| A3KMH1 | von Willebrand factor A domain-containing protein 8 [OS=Homo sapiens] | -6.64 | 8.30303E-17 | 33281.81641 | 48538.08984 | 428310.1563 | | 294603.5313 | | 225022.1928 | 348359.1932 | 428310.1563 | | 306042.0294 | | |
| Q15119 | [Pyruvate dehydrogenase (acetyl-transferring)] kinase isozyme 2, mitochond | -6.64 | 8.30303E-17 | 48251.69141 | 115982.7266 | 840857.125 | | 575517.7656 | | 326235.2414 | 832411.1884 | 840857.125 | | 597863.251 | | |
| P55327 | Tumor protein D52 [OS=Homo sapiens] | -6.64 | 8.30303E-17 | | | 443209.7813 | | | | | | 443209.7813 | | | | |
| O00534 | von Willebrand factor A domain-containing protein 5A [OS=Homo sapiens] | -6.64 | 8.30303E-17 | | | 703872.1875 | | | | | | 703872.1875 | | | | |
| Q9P121 | Neurotrimin [OS=Homo sapiens] | -6.64 | 8.30303E-17 | 991839.25 | 1914592.645 | 19058132.44 | | 12963913.03 | | 6705939.371 | 13741083.57 | 19058132.44 | | 13467259.66 | | |
| Q96JD6 | 1,5-anhydro-D-fructose reductase [OS=Homo sapiens] | -6.64 | 8.30303E-17 | | | 526899.8125 | | | | | | 526899.8125 | | | | |
| Q86SX6 | Glutaredoxin-related protein 5, mitochondrial [OS=Homo sapiens] | -6.64 | 8.30303E-17 | | 63217.20703 | 1318752.719 | | 208475.6875 | | | 453711.6172 | 1318752.719 | | 216570.1213 | | |
| Q14832 | Metabotropic glutamate receptor 3 [OS=Homo sapiens] | -6.64 | 8.30303E-17 | | | 291327.6875 | | | | | | 291327.6875 | | | | |
| Q9BQG1 | Synaptotagmin-3 [OS=Homo sapiens] | -6.64 | 8.30303E-17 | | | 278278.5313 | | | | | | 278278.5313 | | | | |
| P23368 | NAD-dependent malic enzyme, mitochondrial [OS=Homo sapiens] | -6.64 | 8.30303E-17 | | | 824714.125 | | | | | | 824714.125 | | | | |
| Q07817 | Bcl-2-like protein 1 [OS=Homo sapiens] | -6.64 | 8.30303E-17 | | | 394671 | | | | | | 394671 | | | | |
| O75382 | Tripartite motif-containing protein 3 [OS=Homo sapiens] | -6.64 | 8.30303E-17 | | 37392.875 | 678569.3125 | | | | | 268369.6826 | 678569.3125 | | | | |
| P30566 | Adenylosuccinate lyase [OS=Homo sapiens] | -6.64 | 8.30303E-17 | | | 508774.7188 | | | | | | 508774.7188 | | | | |
| Q08174 | Protocadherin-1 [OS=Homo sapiens] | -6.64 | 8.30303E-17 | | | 740807.3438 | | | | | | 740807.3438 | | | | |
| Q9Y4J8 | Dystrobrevin alpha [OS=Homo sapiens] | -6.64 | 8.30303E-17 | | | 302297.5625 | | | | | | 302297.5625 | | | | |
| Q93045 | Stathmin-2 [OS=Homo sapiens] | -6.64 | 8.30303E-17 | | | 277889.4063 | | | | | | 277889.4063 | | | | |
| Q92614 | Unconventional myosin-XVIIIa [OS=Homo sapiens] | -6.64 | 8.30303E-17 | | | 310332.8125 | | | | | | 310332.8125 | | | | |
| P11137 | Microtubule-associated protein 2 [OS=Homo sapiens] | -6.64 | 8.30303E-17 | 653556.3125 | 6422906.238 | 53606601.5 | | 35571555.33 | | 4418769.48 | 46097373.06 | 53606601.5 | | 36952683.27 | | |
| Q15075 | Early endosome antigen 1 [OS=Homo sapiens] | -6.64 | 8.30303E-17 | | | 312038.125 | | | | | | 312038.125 | | | | |
| Q16543 | Hsp90 co-chaperone Cdc37 [OS=Homo sapiens] | -6.64 | 8.30303E-17 | 42265.32422 | 220007.6172 | 1435164 | | 962292.6875 | | 285760.7235 | 1579000.663 | 1435164 | | 999655.3867 | | |
| P22694 | cAMP-dependent protein kinase catalytic subunit beta [OS=Homo sapiens] | -6.64 | 8.30303E-17 | | | 1289889.375 | | | | | | 1289889.375 | | | | |
| P78330 | Phosphoserine phosphatase [OS=Homo sapiens] | -6.64 | 8.30303E-17 | | | 336536.4688 | | | | | | 336536.4688 | | | | |
| Q9UBB6 | Neurochondrin [OS=Homo sapiens] | -6.64 | 8.30303E-17 | 134802.6719 | 2186456.646 | 31083827.81 | | 17827530.34 | | 911416.3859 | 15692258.92 | 31083827.81 | | 18519715.44 | | |
| O14787 | Transportin-2 [OS=Homo sapiens] | -6.64 | 8.30303E-17 | | 55655.91797 | 281486.5625 | | | | | 399444.0396 | 281486.5625 | | | | |
| P40123 | Adenylyl cyclase-associated protein 2 [OS=Homo sapiens] | -6.64 | 8.30303E-17 | 514143.9688 | 338425.1875 | 2931344.031 | | 2219856.063 | | 3476186.572 | 2428886.791 | 2931344.031 | | 2306045.863 | | |
| P48556 | 26S proteasome non-ATPase regulatory subunit 8 [OS=Homo sapiens] | -6.64 | 8.30303E-17 | 78676.78906 | 312407.3906 | 2788016.125 | | 824078.375 | | 531942.8298 | 2242156.354 | 2788016.125 | | 856074.6614 | | |
| P09661 | U2 small nuclear ribonucleoprotein A' [OS=Homo sapiens] | -6.64 | 8.30303E-17 | | | 336775.9063 | | | | | | 336775.9063 | | | | Met-loss [N-Term] |
| O76054 | SEC14-like protein 2 [OS=Homo sapiens] | -6.64 | 8.30303E-17 | | | 400553.5313 | | | | | | 400553.5313 | | | | |
| Q9Y2G0 | Protein EFR3 homolog B [OS=Homo sapiens] | -6.64 | 8.30303E-17 | | | 436751.7813 | | | | | | 436751.7813 | | | | |
| Q16401 | 26S proteasome non-ATPase regulatory subunit 5 [OS=Homo sapiens] | -6.64 | 8.30303E-17 | | | 927324.0313 | | | | | | 927324.0313 | | | | |
| Q14C86 | GTPase-activating protein and VPS9 domain-containing protein 1 [OS=Hom | -6.64 | 8.30303E-17 | | | 684806.375 | | | | | | 684806.375 | | | | |
| P78417 | Glutathione S-transferase omega-1 [OS=Homo sapiens] | -6.64 | 8.30303E-17 | 25956.07031 | 111689.7656 | 858609.3125 | | 811819.25 | | 175491.98 | 801600.4908 | 858609.3125 | | 843339.5544 | | |
| Q6U841 | Sodium-driven chloride bicarbonate exchanger [OS=Homo sapiens] | -6.64 | 8.30303E-17 | | 42413.77734 | 335067.8438 | | | | | 304404.8355 | 335067.8438 | | | | |
| Q15555 | Microtubule-associated protein RP/EB family member 2 [OS=Homo sapiens | -6.64 | 8.30303E-17 | 14484.76074 | 603942.5469 | 7481762.531 | | 4391047.625 | | 97933.13517 | 4334512.113 | 7481762.531 | | 4561537.741 | | |
| Q8N9N7 | Leucine-rich repeat-containing protein 57 [OS=Homo sapiens] | -6.64 | 8.30303E-17 | | | 347279.3125 | | | | | | 347279.3125 | | | | |
| Q96NL8 | Cilia- and flagella-associated protein 418 [OS=Homo sapiens] | -6.64 | 8.30303E-17 | | 81743.02344 | 514907.5 | | | | | 586671.9062 | 514907.5 | | | | |

| | | | | | | | | | | | | | | | | |
|---|---|---|---|---|---|---|---|---|---|---|---|---|---|---|---|---|
| Q9UKE5 | TRAF2 and NCK-interacting protein kinase [OS=Homo sapiens] | -6.64 | 8.30303E-17 | | | 357278.5 | | | | | | 357278.5 | | | | |
| Q8N568 | Serine/threonine-protein kinase DCLK2 [OS=Homo sapiens] | -6.64 | 8.30303E-17 | | 63966.41016 | 1102845.219 | | 357663.6875 | | | 459088.6684 | 1102845.219 | | 371550.6067 | | |
| Q9UI09 | NADH dehydrogenase [ubiquinone] 1 alpha subcomplex subunit 12 [OS=Hc | -6.64 | 8.30303E-17 | | | 316271.25 | | | | | | 316271.25 | | | | |
| Q13907 | Isopentenyl-diphosphate Delta-isomerase 1 [OS=Homo sapiens] | -6.64 | 8.30303E-17 | | | 421003 | | | | | | 421003 | | | | |
| O43815 | Striatin [OS=Homo sapiens] | -6.64 | 8.30303E-17 | | | 336487.375 | | | | | | 336487.375 | | | | |
| Q8NB37 | Glutamine amidotransferase-like class 1 domain-containing protein 1 [OS=I | -6.64 | 8.30303E-17 | | | 514991.6875 | | | | | | 514991.6875 | | | | |
| P33121 | Long-chain-fatty-acid--CoA ligase 1 [OS=Homo sapiens] | -6.64 | 8.30303E-17 | | 64920.49609 | 402167.3125 | | | | | 465936.1692 | 402167.3125 | | | | |
| P48730 | Casein kinase I isoform delta [OS=Homo sapiens] | -3.84 | 0.055226077 | | 51480.03906 | 345599.1875 | | 173542.7344 | | | 369473.6429 | 345599.1875 | | 180280.835 | | |
| Q8WZA2 | Rap guanine nucleotide exchange factor 4 [OS=Homo sapiens] | -3.58 | 0.05754144 | | 70349.41992 | 483145.1563 | | 109079.3438 | | | 504899.7034 | 483145.1563 | | 113314.5403 | | |
| Q9H0F7 | ADP-ribosylation factor-like protein 6 [OS=Homo sapiens] | 3.22 | 0.060793131 | | | 618107.3438 | | 471985.2031 | 36176.47656 | | | 618107.3438 | | 490310.8554 | 306131.9359 | |
| P19013 | Keratin, type II cytoskeletal 4 [OS=Homo sapiens] | 5.41 | 0.066105987 | | 180917.9609 | 1154483.063 | 3180931.844 | 249896649.1 | 127533.2148 | | 1298453.135 | 1154483.063 | 23252140.62 | 259599324.2 | 1079209.3 | |
| P14867 | Gamma-aminobutyric acid receptor subunit alpha-1 [OS=Homo sapiens] | -3.75 | 0.070898743 | | 67019.5625 | 523046.9375 | | 257135.9844 | | | 481001.226 | 523046.9375 | | 267119.7394 | | |
| Q01081 | Splicing factor U2AF 35 kDa subunit [OS=Homo sapiens] | 2.93 | 0.078486529 | | 130418.5625 | | 58504.28125 | 522691.7188 | | | 936017.5762 | | 427657.6303 | 542986.1403 | | |
| Q7L576 | Cytoplasmic FMR1-interacting protein 1 [OS=Homo sapiens] | -3.69 | 0.082944343 | | 105327.9688 | 528159.5625 | | 436528.25 | | | 755941.7013 | 528159.5625 | | 453477.224 | | |
| P36873 | Serine/threonine-protein phosphatase PP1-gamma catalytic subunit [OS=H | -3.69 | 0.082944343 | | 101250.8672 | 947451.5 | | 417854.2188 | | | 726680.2323 | 947451.5 | | 434078.1407 | | Met-loss+Acetyl [N-Term] |
| Q15758 | Neutral amino acid transporter B(0) [OS=Homo sapiens] | 3.52 | 0.083253539 | 286287.3281 | | 121354.3672 | 344446.5 | 601721.1094 | 38240.28125 | 1935621.589 | | 121354.3672 | 2517852.896 | 625083.9855 | 323596.2272 | |
| Q6PIU2 | Neutral cholesterol ester hydrolase 1 [OS=Homo sapiens] | -3.6 | 0.08389382 | | 50729.44531 | 919203.625 | | 238226.9531 | | | 364086.6111 | 919203.625 | | 247476.5319 | | |
| O15173 | Membrane-associated progesterone receptor component 2 [OS=Homo sapi | -3.67 | 0.085985729 | | 109759.2344 | 514233.4063 | | 467627.9688 | | | 787745.0154 | 514233.4063 | | 485784.4438 | | |
| Q13368 | MAGUK p55 subfamily member 3 [OS=Homo sapiens] | -3.56 | 0.08948212 | | 47558.23438 | 220098.2969 | | 235642 | | | 341326.7438 | 220098.2969 | | 244791.2134 | | |
| P60201 | Myelin proteolipid protein [OS=Homo sapiens] | -6.55 | 0.091454472 | 20291.11523 | 29992060.94 | 418632197.4 | | 296613506.2 | 573674.1641 | 137190.5664 | 215253838.5 | 418632197.4 | | 308130045.1 | 4854535.298 | Met-loss [N-Term] |
| Q99653 | Calcineurin B homologous protein 1 [OS=Homo sapiens] | -3.62 | 0.091816041 | | 157053.3555 | 918225.5 | | 264003.9375 | | | 1127176.211 | 918225.5 | | 274254.3528 | | |
| P61225 | Ras-related protein Rap-2b [OS=Homo sapiens] | -3.67 | 0.092961628 | | 79699.70313 | 1117969.375 | | 340903.9063 | | | 572006.9407 | 1117969.375 | | 354140.0976 | | |
| P36915 | Guanine nucleotide-binding protein-like 1 [OS=Homo sapiens] | -3.79 | 0.092961628 | | 91979.54688 | 734666.1875 | | 331400.3438 | | | 660139.7138 | 734666.1875 | | 344267.5426 | | |
| P48444 | Coatomer subunit delta [OS=Homo sapiens] | -3.59 | 0.093908301 | | 83148.17969 | 815909.4688 | | 398090.375 | | | 596756.7509 | 815909.4688 | | 413546.9311 | | |
| Q96DA2 | Ras-related protein Rab-39B [OS=Homo sapiens] | -3.55 | 0.093908301 | | 90694.17969 | 564363.6875 | | 456468.75 | | | 650914.5985 | 564363.6875 | | 474191.9488 | | |
| O43399 | Tumor protein D54 [OS=Homo sapiens] | -3.87 | 0.096361976 | | 84659.83594 | 589356.75 | | 272354.25 | | | 607605.949 | 589356.75 | | 282928.8808 | | |
| P23193 | Transcription elongation factor A protein 1 [OS=Homo sapiens] | -3.84 | 0.10066095 | | 74479.51563 | 363120.7188 | | 252028.5938 | | | 534541.5128 | 363120.7188 | | 261814.0454 | | |
| Q9P2R3 | Rabankyrin-5 [OS=Homo sapiens] | -3.52 | 0.101995668 | | 54784.37109 | 478045.6875 | | 288949 | | | 393188.9239 | 478045.6875 | | 300167.9511 | | |
| Q13424 | Alpha-1-syntrophin [OS=Homo sapiens] | -3.33 | 0.105756061 | | 25708.49805 | 498593.625 | | 174841.125 | | | 184510.5909 | 498593.625 | | 181629.6379 | | |
| O43670 | BUB3-interacting and GLEBS motif-containing protein ZNF207 [OS=Homo sa | -3.8 | 0.106271406 | | 63844.74219 | 567565.5 | | 226324.2188 | | | 458215.4541 | 567565.5 | | 235111.653 | | |
| O00217 | NADH dehydrogenase [ubiquinone] iron-sulfur protein 8, mitochondrial [OS | -3.54 | 0.11495452 | 25898.39453 | 518214.7188 | 3458788.375 | | 2835560.031 | 16158.23828 | 175102.0274 | 3719241.155 | 3458788.375 | | 2945655.616 | 136733.9563 | |
| O00764 | Pyridoxal kinase [OS=Homo sapiens] | -3.68 | 0.118388124 | | 89842.19531 | 727003.5625 | | 376428.1875 | | | 644799.8835 | 727003.5625 | | 391043.6713 | | |
| Q5JVS0 | Intracellular hyaluronan-binding protein 4 [OS=Homo sapiens] | -3.42 | 0.118774448 | | 47029.33203 | 608710.9375 | | 282964.6875 | | | 337530.7973 | 608710.9375 | | 293951.2872 | | |
| Q9Y276 | Mitochondrial chaperone BCS1 [OS=Homo sapiens] | -3.57 | 0.119911172 | | 80046.36719 | 517179 | | 393796.75 | | | 574494.9581 | 517179 | | 409086.5986 | | |
| Q9UIA9 | Exportin-7 [OS=Homo sapiens] | -3.62 | 0.120346979 | | 72874.25 | 1171716.375 | | 334545.6563 | | | 523020.4777 | 1171716.375 | | 347534.9774 | | |
| P62310 | U6 snRNA-associated Sm-like protein LSm3 [OS=Homo sapiens] | -3.42 | 0.120346979 | | 52196.48047 | 359985.75 | | 314103.6875 | | | 374615.5624 | 359985.75 | | 326299.3134 | | |
| Q8ND76 | Cyclin-Y [OS=Homo sapiens] | -3.69 | 0.120346979 | | 60114.03516 | 423178.75 | | 471089.3594 | | | 431440.0681 | 423178.75 | | 489380.229 | | |
| P07741 | Adenine phosphoribosyltransferase [OS=Homo sapiens] | -3.45 | 0.122830512 | | 52739.3125 | 358941.8125 | | 306852.5938 | | | 378511.4827 | 358941.8125 | | 318766.6832 | | |
| O75323 | Protein NipSnap homolog 2 [OS=Homo sapiens] | -3.44 | 0.125378234 | | 67841.40625 | 967870.7188 | | 396581.5 | | | 486899.621 | 967870.7188 | | 411979.4714 | | |
| O75317 | Ubiquitin carboxyl-terminal hydrolase 12 [OS=Homo sapiens] | -3.7 | 0.125378234 | | 76022.28125 | 411110.0313 | | 311860.9375 | | | 545613.9837 | 411110.0313 | | 323969.4847 | | |
| P40616 | ADP-ribosylation factor-like protein 1 [OS=Homo sapiens] | -3.47 | 0.125378234 | | 85701.39844 | 737092.75 | | 480565.125 | | | 615081.2715 | 737092.75 | | 499223.9078 | | |
| Q9H1V8 | Sodium-dependent neutral amino acid transporter SLC6A17 [OS=Homo sap | -3.44 | 0.125378234 | | 74921.15625 | 561250.3125 | | 436732.4688 | | | 537711.1796 | 561250.3125 | | 453689.3719 | | |
| Q9Y285 | Phenylalanine--tRNA ligase alpha subunit [OS=Homo sapiens] | -3.5 | 0.128029939 | | 88857.05469 | 552321 | | 481982.8125 | | | 637729.5023 | 552321 | | 500696.6395 | | |
| Q86VS8 | Protein Hook homolog 3 [OS=Homo sapiens] | -3.58 | 0.128619567 | | 65500.73047 | 904914.75 | | 502480.7656 | | | 470100.5271 | 904914.75 | | 521990.4615 | | |
| Q8NFP9 | Neurobeachin [OS=Homo sapiens] | -3.48 | 0.128619567 | | 51869.81641 | 448276.375 | | 288718 | | | 372271.0855 | 448276.375 | | 299927.9821 | | |
| O00629 | Importin subunit alpha-3 [OS=Homo sapiens] | -3.48 | 0.131044317 | | 90700.71094 | 575754.6875 | | 502247.0938 | | | 650961.4735 | 575754.6875 | | 521747.7169 | | |
| Q9H977 | WD repeat-containing protein 54 [OS=Homo sapiens] | -3.56 | 0.131101138 | | 56732.63672 | 700295.5 | | 439269.125 | | | 407171.6794 | 700295.5 | | 456324.5182 | | |
| Q9BS26 | Endoplasmic reticulum resident protein 44 [OS=Homo sapiens] | -3.36 | 0.133404167 | | 72291.14844 | 415309.875 | | 471717.2188 | | | 518835.5419 | 415309.875 | | 490032.4661 | | |
| O00303 | Eukaryotic translation initiation factor 3 subunit F [OS=Homo sapiens] | -3.41 | 0.137582493 | | 55322.03906 | 618291.9531 | | 501339.4063 | | | 397047.7816 | 618291.9531 | | 520804.7869 | | |
| Q9UKY7 | Protein CDV3 homolog [OS=Homo sapiens] | -3.52 | 0.139859966 | | 93024.42188 | 592861.9375 | | 485096 | | | 667638.8102 | 592861.9375 | | 503930.7019 | | |
| Q9H479 | Fructosamine-3-kinase [OS=Homo sapiens] | -3.55 | 0.142103826 | | 53042.37891 | 341150.7188 | | 267796.3125 | | | 380686.5985 | 341150.7188 | | 278193.9734 | | |
| P30519 | Heme oxygenase 2 [OS=Homo sapiens] | -3.5 | 0.142192201 | | 91783.71094 | 689433.0625 | | 493506.6563 | | | 658734.194 | 689433.0625 | | 512667.9166 | | |
| Q96B23 | Protein ARK2N [OS=Homo sapiens] | 2.79 | 0.14344642 | 542797.5625 | | 143303.6563 | 345422.0938 | 612545.0625 | 117267.1172 | 3669916.819 | | 143303.6563 | 2524984.341 | 636328.1975 | 992335.7113 | |
| O14495 | Phospholipid phosphatase 3 [OS=Homo sapiens] | -3.54 | 0.144900951 | | 85445.83594 | 1372208.25 | | 433736.5 | | | 613247.0924 | 1372208.25 | | 450577.0794 | | |
| P36969 | Phospholipid hydroperoxide glutathione peroxidase GPX4 [OS=Homo sapier | -3.5 | 0.144900951 | | 79471.23438 | 988772.9688 | | 428574.2188 | | | 570367.2143 | 988772.9688 | | 445214.3635 | | |
| P36507 | Dual specificity mitogen-activated protein kinase kinase 2 [OS=Homo sapier | -3.54 | 0.14562653 | | 61496.06641 | 366121.6563 | | 314939.25 | | | 441358.944 | 366121.6563 | | 327167.3181 | | |
| O00445 | Synaptotagmin-5 [OS=Homo sapiens] | -3.54 | 0.14562653 | | 65608.20313 | 281440.8125 | | 335614.5625 | | | 470871.8613 | 281440.8125 | | 348645.3858 | | |
| Q99963 | Endophilin-A3 [OS=Homo sapiens] | -3.42 | 0.146322266 | | 83480.74219 | 499116.7188 | | 502829.4063 | | | 599143.5611 | 499116.7188 | | 522352.6387 | | Met-loss+Acetyl [N-Term] |
| Q8NAT1 | Protein O-linked-mannose beta-1,4-N-acetylglucosaminyltransferase 2 [OS | -3.57 | 0.150282194 | | 71357.36719 | 302218.6875 | | 348687.6563 | | | 512133.7684 | 302218.6875 | | 362226.0653 | | |
| Q9NVT9 | Armadillo repeat-containing protein 1 [OS=Homo sapiens] | -3.51 | 0.153784045 | | 48613.65234 | 455695.5313 | | 257698.8594 | | | 348901.5073 | 455695.5313 | | 267704.469 | | |
| Q9NX40 | OCIA domain-containing protein 1 [OS=Homo sapiens] | -3.38 | 0.157390711 | | 77996.88281 | 564257.0625 | | 498025.7813 | | | 559785.7529 | 564257.0625 | | 517362.5046 | | |
| P50579 | Methionine aminopeptidase 2 [OS=Homo sapiens] | -3.5 | 0.157667853 | | 75225.99219 | 707711.4219 | | 406579.9375 | | | 539898.9954 | 707711.4219 | | 422366.1158 | | |
| O95777 | U6 snRNA-associated Sm-like protein LSm8 [OS=Homo sapiens] | -3.24 | 0.160563503 | | 28710.91406 | 236102.2656 | | 222924.4219 | | | 206059.0124 | 236102.2656 | | 231579.853 | | Met-loss+Acetyl [N-Term] |
| P02766 | Transthyretin [OS=Homo sapiens] | 2.01 | 0.1614045 | | 56868.69141 | 128953.7031 | 78668.42969 | 2822023.844 | 61523.03906 | | 408148.1476 | 128953.7031 | 575054.5686 | 2931593.863 | 520619.1658 | |
| P34931 | Heat shock 70 kDa protein 1-like [OS=Homo sapiens] | -3.67 | 0.162159501 | | 63164.95313 | 356497.2813 | | 270475.4063 | | | 453336.5895 | 356497.2813 | | 280977.0877 | | |
| Q14444 | Caprin-1 [OS=Homo sapiens] | -3.46 | 0.163819317 | | 91332.26563 | 599382.75 | | 520518.9375 | | | 655494.1587 | 599382.75 | | 540728.997 | | |
| Q92783 | Signal transducing adapter molecule 1 [OS=Homo sapiens] | -3.49 | 0.163819317 | | 152659.0078 | 930011 | | 413442.3125 | | | 1095637.858 | 930011 | | 429494.9345 | | |
| P21359 | Neurofibromin [OS=Homo sapiens] | -3.32 | 0.165053498 | | 46484.56641 | 696694.5938 | | 524436.125 | | | 333620.9995 | 696694.5938 | | 544798.2762 | | |
| Q13404 | Ubiquitin-conjugating enzyme E2 variant 1 [OS=Homo sapiens] | -3.62 | 0.170510695 | | 84167.46094 | 634756.875 | | 383024.7813 | | | 604072.1602 | 634756.875 | | 397896.3893 | | Met-loss+Acetyl [N-Term] |
| P10155 | RNA-binding protein RO60 [OS=Homo sapiens] | -3.37 | 0.170600755 | | 65878.125 | 611191.125 | | 424997.125 | | | 472809.0979 | 611191.125 | | 441498.3828 | | |
| O15079 | Syntaphilin [OS=Homo sapiens] | -3.4 | 0.17233973 | | 36174.91406 | 742089.6563 | | 225601.9375 | | | 259628.3438 | 742089.6563 | | 234361.3279 | | |
| Q8N1G4 | Leucine-rich repeat-containing protein 47 [OS=Homo sapiens] | -3.31 | 0.175151883 | | 47133.80469 | 484635.25 | | 332640.9375 | | | 338280.6004 | 484635.25 | | 345556.3046 | | |
| O15371 | Eukaryotic translation initiation factor 3 subunit D [OS=Homo sapiens] | 2.66 | 0.176223077 | | 83061.64844 | 1063354.031 | 116398.3281 | 604860.8125 | 28834.44531 | | 596135.7138 | 1063354.031 | 850854.5377 | 628345.5931 | 244002.3298 | |
| Q9H4A4 | Aminopeptidase B [OS=Homo sapiens] | -3.4 | 0.176255725 | | 51955.99219 | 444491.375 | | 324246.6875 | | | 372889.5714 | 444491.375 | | 336836.1331 | | |
| O14576 | Cytoplasmic dynein 1 intermediate chain 1 [OS=Homo sapiens] | -3.52 | 0.176322562 | | 61245.46484 | 634025.3125 | | 322898.5938 | | | 439560.3698 | 634025.3125 | | 335435.6973 | | |
| Q9UBS8 | E3 ubiquitin-protein ligase RNF14 [OS=Homo sapiens] | -3.19 | 0.177887046 | | 58208.5625 | 439230.75 | | 485062.5 | | | 417764.4389 | 439230.75 | | 503895.9012 | | |
| Q5JU85 | IQ motif and SEC7 domain-containing protein 2 [OS=Homo sapiens] | -3.45 | 0.179023194 | | 71966.97656 | 498981.875 | | 417415.375 | | | 516508.9515 | 498981.875 | | 433622.2581 | | |
| Q99733 | Nucleosome assembly protein 1-like 4 [OS=Homo sapiens] | -3.38 | 0.179543892 | | 44230.92969 | 333779.375 | | 282328.4375 | | | 317446.5874 | 333779.375 | | 293290.3336 | | |
| Q9NZ72 | Stathmin-3 [OS=Homo sapiens] | -3.47 | 0.182515568 | | 48192.94922 | 340327.3125 | | 272741.875 | | | 345882.1095 | 340327.3125 | | 283331.5561 | | |
| Q8N2K0 | Lysophosphatidylserine lipase ABHD12 [OS=Homo sapiens] | -3.46 | 0.18275723 | | 54726.88672 | 936723.0313 | | 562051.5625 | | | 392776.357 | 936723.0313 | | 583874.199 | | |
| O75110 | Probable phospholipid-transporting ATPase IIA [OS=Homo sapiens] | -3.26 | 0.18275723 | | 36741.26953 | 726948.4531 | | 475229.0156 | | | 263693.0924 | 726948.4531 | | 493680.6146 | | |
| P01834 | Immunoglobulin kappa constant [OS=Homo sapiens] | 3 | 0.184231751 | 307099.3164 | 316162.918 | 446059.375 | 501349.2891 | 44412635.16 | 430771.0078 | 2076333.838 | 2269109.876 | 446059.375 | 3664789.043 | 46137033.51 | 3645262.753 | |

| | | | | | | | | | | | | | | | | |
|---|---|---|---|---|---|---|---|---|---|---|---|---|---|---|---|---|
| P25311 | Zinc-alpha-2-glycoprotein [OS=Homo sapiens] | 2.66 | 0.186132212 | 143009.2715 | 72736.42773 | 149080.0625 | 191345.6211 | 43615008.14 | 288337.9121 | 966902.1508 | 522031.3235 | 149080.0625 | 1398708.148 | 45308437.23 | 2439967.947 | |
| Q9UMS0 | NFU1 iron-sulfur cluster scaffold homolog, mitochondrial [OS=Homo sapie | -3.48 | 0.187289924 | | 73506.39844 | 924210.2188 | | 535773.7344 | | | 527557.4242 | 924210.2188 | | 556576.0881 | | |
| Q9UL46 | Proteasome activator complex subunit 2 [OS=Homo sapiens] | -3.75 | 0.188176121 | | 99418.95313 | 286984.1875 | | 380298.0313 | | | 713532.5352 | 286984.1875 | | 395063.7685 | | |
| Q96KR1 | Zinc finger RNA-binding protein [OS=Homo sapiens] | -3.53 | 0.190443733 | | 79881.96094 | 479277.2813 | | 413972.4375 | | | 573315.01 | 479277.2813 | | 430045.6425 | | |
| Q15118 | [Pyruvate dehydrogenase (acetyl-transferring)] kinase isozyme 1, mitochond | -3.15 | 0.190443733 | | 25888.08984 | 302059.125 | | 225752.6719 | | | 185799.5261 | 302059.125 | | 234517.9148 | | |
| Q04323 | UBX domain-containing protein 1 [OS=Homo sapiens] | -3.33 | 0.193006704 | | 48733.37109 | 416264.1875 | | 330865.5 | | | 349760.732 | 416264.1875 | | 343711.9326 | | |
| P54750 | Dual specificity calcium/calmodulin-dependent 3',5'-cyclic nucleotide phos | -3.51 | 0.193980891 | | 81974 | 406938.0313 | | 433525.6875 | | | 588329.6313 | 406938.0313 | | 450358.0817 | | |
| Q9H1P3 | Oxysterol-binding protein-related protein 2 [OS=Homo sapiens] | -3.21 | 0.194662049 | | 38302.83594 | 451968.9063 | | 308576.4063 | | | 274900.4971 | 451968.9063 | | 320557.4258 | | |
| P12955 | Xaa-Pro dipeptidase [OS=Homo sapiens] | -3.37 | 0.195569364 | | 48961.34375 | 726012.6875 | | 314727.7813 | | | 351396.898 | 726012.6875 | | 326947.6387 | | |
| O00168 | Phospholemman [OS=Homo sapiens] | -3.33 | 0.197309954 | | 83705.64063 | 527714.5625 | | 572648.375 | | | 600757.6633 | 527714.5625 | | 594882.4513 | | |
| P04003 | C4b-binding protein alpha chain [OS=Homo sapiens] | -3.04 | 0.199093534 | 144158.9375 | 185921.5313 | | 196416.7656 | | | 974675.1751 | 1334363.895 | | 1435777.463 | | | |
| P17252 | Protein kinase C alpha type [OS=Homo sapiens] | -3.26 | 0.20033997 | | 65513.14063 | 727226.25 | | 491534.6875 | | | 470189.5951 | 727226.25 | | 510619.3828 | | |
| Q9NSD9 | Phenylalanine--tRNA ligase beta subunit [OS=Homo sapiens] | -3.48 | 0.201025551 | | 76355.61719 | 521084.7188 | | 421642.1875 | | | 548006.3448 | 521084.7188 | | 438013.184 | | |
| O14994 | Synapsin-3 [OS=Homo sapiens] | -3.33 | 0.205608843 | | 69466.375 | 821367.1875 | | 474720.875 | | | 498562.0659 | 821367.1875 | | 493152.7445 | | |
| P05067 | Amyloid-beta precursor protein [OS=Homo sapiens] | -3.6 | 0.205608843 | | 74539.11719 | 349623.1563 | | 351253.625 | | | 534969.2748 | 349623.1563 | | 364891.6623 | | |
| Q12904 | Aminoacyl tRNA synthase complex-interacting multifunctional protein 1 [OS | -3.46 | 0.207393733 | | 107565.7813 | 941857.1875 | | 285829.2188 | | | 772002.5426 | 941857.1875 | | 296927.0389 | | |
| P53365 | Arfaptin-2 [OS=Homo sapiens] | -3.5 | 0.207416296 | | 65485.84766 | 399792.75 | | 354085.625 | | | 469993.7127 | 399792.75 | | 367833.6196 | | |
| Q9UPN3 | Microtubule-actin cross-linking factor 1, isoforms 1/2/3/4/5 [OS=Homo sap | -3.65 | 0.20860557 | | 73619.01563 | 838805.4688 | | 322432.875 | | | 528365.6808 | 838805.4688 | | 334951.8962 | | |
| P48426 | Phosphatidylinositol 5-phosphate 4-kinase type-2 alpha [OS=Homo sapiens | -3.35 | 0.210717791 | | 71396.38281 | 1215400.469 | | 476531.4688 | | | 512413.7846 | 1215400.469 | | 495033.6378 | | Met-loss+Acetyl [N-Term] |
| Q9P2U7 | Vesicular glutamate transporter 1 [OS=Homo sapiens] | -3.7 | 0.210793319 | | 82909.95313 | 523308.7813 | | 338836.0313 | | | 595046.9924 | 523308.7813 | | 351991.9337 | | |
| P41227 | N-alpha-acetyltransferase 10 [OS=Homo sapiens] | -3.36 | 0.211324318 | | 56667.67969 | 376352 | | 369926.5313 | | | 406705.481 | 376352 | | 384289.5768 | | |
| P47869 | Gamma-aminobutyric acid receptor subunit alpha-2 [OS=Homo sapiens] | -3.43 | 0.219153082 | | 52661.375 | 239960.1719 | | 314643.9063 | | | 377952.1231 | 239960.1719 | | 326860.5071 | | |
| O76094 | Signal recognition particle subunit SRP72 [OS=Homo sapiens] | -3.5 | 0.219631733 | | 75788.85938 | 398108.5313 | | 407682.8438 | | | 543938.7086 | 398108.5313 | | 423511.8443 | | |
| Q53FP2 | Novel acetylcholine receptor chaperone [OS=Homo sapiens] | -3.33 | 0.219759015 | | 53529.70703 | 487174.4063 | | 363193.6563 | | | 384184.1657 | 487174.4063 | | 377295.2862 | | |
| Q9H1K1 | Iron-sulfur cluster assembly enzyme ISCU [OS=Homo sapiens] | -3.48 | 0.220065472 | | 63483.61719 | 1099502.75 | | 351409.2813 | | | 455623.6501 | 1099502.75 | | 365053.3622 | | |
| Q8NCW5 | NAD(P)H-hydrate epimerase [OS=Homo sapiens] | -3.22 | 0.227073921 | | 63083.70313 | 1246469.75 | | 502078.5313 | | | 452753.456 | 1246469.75 | | 521572.6097 | | |
| P35498 | Sodium channel protein type 1 subunit alpha [OS=Homo sapiens] | -2.99 | 0.227073921 | | 43287.5 | 695787.25 | | 471621.2188 | | | 310675.5668 | 695787.25 | | 489932.7387 | | |
| Q9NQL2 | Ras-related GTP-binding protein D [OS=Homo sapiens] | -3.23 | 0.227073921 | | 58604.94531 | 462406.7813 | | 459725.3125 | | | 420609.2892 | 462406.7813 | | 477574.9531 | | |
| Q9BQG0 | Myb-binding protein 1A [OS=Homo sapiens] | -3.63 | 0.230260703 | | 79897.64063 | 482083.375 | | 362163.5313 | | | 573427.5435 | 482083.375 | | 376225.1647 | | |
| Q96QG7 | Myotubularin-related protein 9 [OS=Homo sapiens] | -3.62 | 0.230662616 | | 67282.25781 | 1003410.344 | | 306015.75 | | | 482886.5974 | 1003410.344 | | 317897.3475 | | |
| Q14108 | Lysosome membrane protein 2 [OS=Homo sapiens] | -3.35 | 0.236424734 | | 64844.06641 | 570445.5 | | 429706.125 | | | 465387.6312 | 570445.5 | | 446390.218 | | |
| Q6UWR7 | Glycerophosphocholine cholinephosphodiesterase ENPP6 [OS=Homo sapie | -3.6 | 0.238863088 | | 74741.17969 | 629571.1875 | | 352289.5 | | | 536419.4829 | 629571.1875 | | 365967.7569 | | |
| Q9H2M9 | Rab3 GTPase-activating protein non-catalytic subunit [OS=Homo sapiens] | -3.37 | 0.240767684 | | 62276.10938 | 515984.6875 | | 402687.4688 | | | 446957.3337 | 515984.6875 | | 418322.5151 | | |
| Q99627 | COP9 signalosome complex subunit 8 [OS=Homo sapiens] | -2.97 | 0.24141692 | | 46899.47266 | 431509.4688 | | 526118.25 | | | 336598.793 | 431509.4688 | | 546545.7127 | | |
| Q3YEC7 | Rab-like protein 6 [OS=Homo sapiens] | -3.53 | 0.242268269 | | 73489.23438 | 478774.1563 | | 382977.2813 | | | 527434.2373 | 478774.1563 | | 397847.0451 | | |
| O43765 | Small glutamine-rich tetratricopeptide repeat-containing protein alpha [OS= | -3.68 | 0.243833518 | | 126553.3594 | 629944.5 | | 530000.3125 | | | 908276.9081 | 629944.5 | | 550578.503 | | |
| Q9ULC3 | Ras-related protein Rab-23 [OS=Homo sapiens] | -3.61 | 0.246485046 | | 79929.63281 | 528383 | | 368628.75 | | | 573657.1523 | 528383 | | 382941.4069 | | |
| Q9Y2R0 | Cytochrome c oxidase assembly factor 3 homolog, mitochondrial [OS=Hom | -3.19 | 0.257364036 | | 67817.91406 | 662446.8125 | | 564192.8125 | | | 486731.0169 | 662446.8125 | | 586098.5867 | | |
| P06396 | Gelsolin [OS=Homo sapiens] | 1.86 | 0.261619701 | | 60152.10547 | 651562.6875 | 46858.96875 | 2814161.656 | 59136.41406 | | 431713.2998 | 651562.6875 | 342532.1208 | 2923426.412 | 500423.1103 | |
| P47224 | Guanine nucleotide exchange factor MSS4 [OS=Homo sapiens] | -3.17 | 0.262904378 | | 46913.94531 | 748561.125 | | 400666.25 | | | 336702.6636 | 748561.125 | | 416222.819 | | |
| P10606 | Cytochrome c oxidase subunit 5B, mitochondrial [OS=Homo sapiens] | -2.69 | 0.268001571 | 81760.15625 | 177364.3906 | 1354657.25 | 11680.62305 | 998612.9375 | | 552789.8304 | 1272949.064 | 1354657.25 | 85383.62434 | 1037385.834 | | |
| Q9ULJ8 | Neurabin-1 [OS=Homo sapiens] | -3.1 | 0.270928954 | | 37479.10547 | 416099.7188 | | 351278.2188 | | | 268988.561 | 416099.7188 | | 364917.2109 | | |
| Q99447 | Ethanolamine-phosphate cytidylyltransferase [OS=Homo sapiens] | -3.47 | 0.271861555 | | 55845.47656 | 849036.5625 | | 313205.4375 | | | 400804.5068 | 849036.5625 | | 325366.1872 | | |
| O60739 | Eukaryotic translation initiation factor 1b [OS=Homo sapiens] | -3.4 | 0.271861555 | | 61083.89063 | 677563.5625 | | 377628.8438 | | | 438400.7472 | 677563.5625 | | 392290.9451 | | |
| Q5VW32 | BRO1 domain-containing protein BROX [OS=Homo sapiens] | -3.34 | 0.277397482 | | 54002.09375 | 396573.9375 | | 364043.0625 | | | 387574.4981 | 396573.9375 | | 378177.6721 | | |
| Q96CN7 | Isochorismatase domain-containing protein 1 [OS=Homo sapiens] | -3.13 | 0.277397482 | | 63386.47266 | 572162.25 | | 574311.875 | | | 454926.4412 | 572162.25 | | 596610.5396 | | |
| Q7Z6G3 | N-terminal EF-hand calcium-binding protein 2 [OS=Homo sapiens] | -3.44 | 0.287352568 | | 69671.45313 | 404424.9688 | | 408460.6563 | | | 500033.9172 | 404424.9688 | | 424319.8567 | | |
| Q9Y2B0 | Protein canopy homolog 2 [OS=Homo sapiens] | -3.54 | 0.288090637 | | 122851.8691 | 964325.9063 | | 466243.1875 | | | 881711.2118 | 964325.9063 | | 484345.896 | | |
| P01009 | Alpha-1-antitrypsin [OS=Homo sapiens] | 2.43 | 0.290014187 | 128230.9805 | 168529.6699 | | 374814.4414 | 7007663.813 | 322322.7461 | 866984.4236 | 1209542.033 | | 2739838.049 | 7279748.635 | 2727553.804 | |
| P13716 | Delta-aminolevulinic acid dehydratase [OS=Homo sapiens] | -3.27 | 0.290575036 | | 75822.8125 | 732499.3125 | | 565884.25 | | | 544182.3911 | 732499.3125 | | 587855.6973 | | |
| Q9UEE9 | Craniofacial development protein 1 [OS=Homo sapiens] | 2.26 | 0.294439995 | | 81749.53906 | 388972.3125 | 313890.8125 | 866040.125 | 220603.4375 | | 586718.6691 | 388972.3125 | 2294495.346 | 899665.6499 | 1866786.481 | |
| O95502 | Neuronal pentraxin receptor [OS=Homo sapiens] | -3.41 | 0.297488211 | | 66233.15625 | 448594.375 | | 405029.4688 | | | 475357.1669 | 448594.375 | | 420755.4474 | | |
| P09211 | Glutathione S-transferase P [OS=Homo sapiens] | 2.09 | 0.299058118 | | 65817.1875 | 1430971.969 | 76519.07813 | 2404771.719 | | | 472371.7478 | 1430971.969 | 559343.1271 | 2498141.193 | | Met-loss [N-Term] |
| P49840 | Glycogen synthase kinase-3 alpha [OS=Homo sapiens] | -2.92 | 0.310852668 | | 44046.01172 | 1035619.25 | | 528699.875 | | | 316119.4261 | 1035619.25 | | 549227.5738 | | |
| Q8TAF3 | WD repeat-containing protein 48 [OS=Homo sapiens] | -3.36 | 0.316032478 | | 58222.99609 | 508898.625 | | 379561.9688 | | | 417868.0292 | 508898.625 | | 394299.1271 | | |
| Q8N9R8 | Protein SCAI [OS=Homo sapiens] | -3.51 | 0.320307083 | | 73201.99219 | 1244826.578 | | 521526.4844 | | | 525372.6923 | 1244826.578 | | 541775.6637 | | |
| O75251 | NADH dehydrogenase [ubiquinone] iron-sulfur protein 7, mitochondrial [OS | -3.92 | 0.320621624 | | 161246.9063 | 744410.875 | | 486010.5 | | | 1157273.439 | 744410.875 | | 504880.709 | | |
| P61626 | Lysozyme C [OS=Homo sapiens] | 3.39 | 0.324482604 | 960788.1445 | 967369.25 | 1361537.469 | 1517394.648 | 403324101.6 | 2078375.125 | 6495999.272 | 6942835.463 | 1361537.469 | 11091930.12 | 418983866.3 | 17587588.98 | |
| O95219 | Sorting nexin-4 [OS=Homo sapiens] | -3.05 | 0.325118748 | | 36350.48438 | 424412.75 | | 366709 | | | 260888.4167 | 424412.75 | | 380947.1193 | | |
| Q6NYC1 | Bifunctional arginine demethylase and lysyl-hydroxylase JMJD6 [OS=Homo s | 2.06 | 0.32973777 | 729611.3125 | 34810.84375 | | 436796.75 | 1005768 | 202593.25 | 4932986.092 | 249838.374 | | 3192919.544 | 1044818.704 | 1714380.993 | |
| Q9Y3U8 | Large ribosomal subunit protein eL36 [OS=Homo sapiens] | -2.53 | 0.33171323 | 24740.1543 | 126333.4453 | 1636394.563 | | 1460743 | 15914.56055 | 167271.0318 | 906698.5789 | 1636394.563 | | 1517458.906 | 134671.9109 | |
| Q9H019 | Mitochondrial fission regulator 1-like [OS=Homo sapiens] | -3.25 | 0.336245171 | | 49595.15234 | 607482.75 | | 378257.3438 | | | 355945.7596 | 607482.75 | | 392943.8477 | | |
| Q15046 | Lysine--tRNA ligase [OS=Homo sapiens] | -3.39 | 0.337266267 | | 65340.23828 | 429917.5625 | | 412086.5625 | | | 468948.6703 | 429917.5625 | | 428086.5451 | | |
| Q92572 | AP-3 complex subunit sigma-1 [OS=Homo sapiens] | -3.28 | 0.34602717 | | 92623.05078 | 968626.8906 | | 254753.7188 | | | 664758.1589 | 968626.8906 | | 264644.9781 | | |
| P05109 | Protein S100-A8 [OS=Homo sapiens] | 3.61 | 0.348961505 | 904362.5234 | 461113.4258 | 763480.0313 | 776336.7656 | 44017114.63 | 2555583.703 | 6114499.151 | 3309423.62 | 763480.0313 | 5674906.766 | 45726156.2 | 21625814.91 | |
| Q96FN4 | Copine-2 [OS=Homo sapiens] | -2.95 | 0.37109601 | | 29126.11523 | 801976 | | 337782 | | | 209038.9226 | 801976 | | 350896.9778 | | |
| Q9NX47 | E3 ubiquitin-protein ligase MARCHF5 [OS=Homo sapiens] | -3.32 | 0.37123524 | | 59271.63281 | 364253 | | 410966.0625 | | | 425394.1235 | 364253 | | 426922.5398 | | |
| P02787 | Serotransferrin [OS=Homo sapiens] | 1.98 | 0.378733739 | 85811.69531 | 45268.91406 | 552143.5625 | 241053.7148 | 11682076.34 | 137712.5977 | 580182.7525 | 324896.2295 | 552143.5625 | 1762066.951 | 12135653.42 | 1165349.092 | |
| P29475 | Nitric oxide synthase 1 [OS=Homo sapiens] | -2.8 | 0.378868552 | | 21287.68555 | 356154.7188 | | 305266.2813 | | | 152782.2991 | 356154.7188 | | 317118.7794 | | |
| P01876 | Immunoglobulin heavy constant alpha 1 [OS=Homo sapiens] | 3.57 | 0.382740315 | 278713.125 | 268276.0195 | 228675.9375 | 796446.6758 | 203268497.6 | 788121.625 | 1884411.53 | 1925424.301 | 228675.9375 | 5821907.23 | 211160753.1 | 6669228.784 | |
| Q02952 | A-kinase anchor protein 12 [OS=Homo sapiens] | 2.73 | 0.402572685 | 1217887.945 | | 156148.7031 | 1898030.914 | 1606447.172 | 566332.8652 | 8234280.628 | | 156148.7031 | 13874324.85 | 1668820.297 | 4792411.89 | |
| P02511 | Alpha-crystallin B chain [OS=Homo sapiens] | -2.77 | 0.403242567 | 29652.96484 | 240830.4609 | 2433696.875 | | 1583907.375 | 23733.53906 | 200487.1096 | 1728446.779 | 2433696.875 | | 1645405.353 | 200837.5317 | |
| Q15043 | Metal cation symporter ZIP14 [OS=Homo sapiens] | 4.08 | 0.403525068 | 240879.9531 | | | 877242.375 | 1349451.063 | | 1628617.098 | | | 6412511.824 | 1401845.864 | | |
| P50150 | Guanine nucleotide-binding protein G(I)/G(S)/G(O) subunit gamma-4 [OS=H | -3.09 | 0.405538974 | | 41797.17578 | 541786.9375 | | 398540.375 | | | 299979.4693 | 541786.9375 | | 414014.4032 | | |
| P0DOY2 | Immunoglobulin lambda constant 2 [OS=Homo sapiens] | 1.82 | 0.440824708 | 180722.6875 | 145595.0938 | 763360.8906 | 269638.6875 | 26948074.88 | 235062.9531 | 1221886.899 | 1044939.954 | 763360.8906 | 1971018.867 | 27994381.09 | 1989145.537 | |
| P19623 | Spermidine synthase [OS=Homo sapiens] | -2.57 | 0.465725685 | | 28872.82031 | 981928.9688 | | 569196.4375 | | | 207221.018 | 981928.9688 | | 591296.4862 | | |
| Q9NY46 | Sodium channel protein type 3 subunit alpha [OS=Homo sapiens] | -2.96 | 0.476312372 | | 38430.91797 | 508657.5 | | 440527.0938 | | | 275819.7453 | 508657.5 | | 457631.3298 | | |
| P55087 | Aquaporin-4 [OS=Homo sapiens] | -3.61 | 0.477711717 | | 90306.44531 | 1176726.875 | | 624326.7031 | | | 648131.8183 | 1176726.875 | | 648567.281 | | |
| Q9Y6M5 | Proton-coupled zinc antiporter SLC30A1 [OS=Homo sapiens] | -2.99 | 0.485909306 | | 37774.59766 | 468455.9063 | | 415178.8438 | | | 271109.3165 | 468455.9063 | | 431298.8896 | | |
| Q9NPH2 | Inositol-3-phosphate synthase 1 [OS=Homo sapiens] | -3.65 | 0.491371848 | | 100540.4219 | 1850148.688 | | 440494.2813 | | | 721581.347 | 1850148.688 | | 457597.2433 | | |
| Q96B54 | Zinc finger protein 428 [OS=Homo sapiens] | 2.73 | 0.492089983 | 987437.875 | | 233076.875 | 1175797.5 | 1714896.625 | 756554.9688 | 6676181.168 | | 233076.875 | 8594905.566 | 1781480.49 | 6402105.987 | |
| P11182 | Lipoamide acyltransferase component of branched-chain alpha-keto acid d | -3.7 | 0.492230137 | | 148346.9063 | 464449 | | 605379.75 | | | 1064689.788 | 464449 | | 628884.6792 | | |

| | | | | | | | | | | | | | | | | |
|---|---|---|---|---|---|---|---|---|---|---|---|---|---|---|---|---|
| Q96HU8 | GTP-binding protein Di-Ras2 [OS=Homo sapiens] | -3.48 | 0.492848032 | | 100459.3906 | 787203.6875 | | 560815.25 | | | 720999.7835 | 787203.6875 | | 582589.8845 | | |
| Q9ULV4 | Coronin-1C [OS=Homo sapiens] | 1.18 | 0.525874839 | | 290749.3477 | 10203758.63 | 81275.95117 | 6872932.875 | | | 2086715.989 | 10203758.63 | 594115.1645 | 7139786.533 | | |
| Q15785 | Mitochondrial import receptor subunit TOM34 [OS=Homo sapiens] | -4.89 | 0.526497886 | | 439643.6484 | 2186385.25 | | 345145.7813 | | | 3155334.443 | 2186385.25 | | 358546.6708 | | |
| P13646 | Keratin, type I cytoskeletal 13 [OS=Homo sapiens] | 2.93 | 0.526497886 | 405756.9102 | 316689.1602 | 148272.8125 | 2681303.867 | 245505200.8 | 194748.8906 | 2743369.189 | 2272886.73 | 148272.8125 | 19599934.11 | 255037370.3 | 1648000.595 | |
| P06702 | Protein S100-A9 [OS=Homo sapiens] | 3.17 | 0.537105258 | 1699705.738 | 699367.3906 | 2321825.602 | 2047285.992 | 124431366.5 | 4319069.516 | 11491906.2 | 5019378.816 | 2321825.602 | 14965357.36 | 129262632.3 | 36548753.16 | |
| Q13616 | Cullin-1 [OS=Homo sapiens] | -3.56 | 0.537105258 | | 60572.39844 | 631770.5 | | 680520.0313 | | | 434729.7538 | 631770.5 | | 706942.4135 | | |
| Q14194 | Dihydropyrimidinase-related protein 1 [OS=Homo sapiens] | -4.29 | 0.544401015 | 36008.25 | 5738168.355 | 72556319.13 | | 48205035.25 | 73653.73047 | 243455.9243 | 41182990.62 | 72556319.13 | | 50076680.18 | 623271.2867 | |
| P33176 | Kinesin-1 heavy chain [OS=Homo sapiens] | -3.53 | 0.550353009 | | 78870.35938 | 1252569.656 | | 660175.4844 | | | 566054.718 | 1252569.656 | | 685807.954 | | |
| P04083 | Annexin A1 [OS=Homo sapiens] | 3.54 | 0.550616394 | 267326.168 | 198362.2168 | 101600.7813 | 1013560.051 | 121975353 | 108078.6035 | 1807423.002 | 1423651.034 | 101600.7813 | 7408973.843 | 126711259.7 | 914580.8345 | Met-loss+Acetyl [N-Term] |
| Q96HS1 | Serine/threonine-protein phosphatase PGAM5, mitochondrial [OS=Homo sa | -3.51 | 0.551148957 | | 107315.709 | 889221.2813 | | 565344.3906 | | | 770207.7671 | 889221.2813 | | 587294.8769 | | |
| Q99829 | Copine-1 [OS=Homo sapiens] | -3.35 | 0.556096594 | | 61104.12109 | 799597.6563 | | 717453.4063 | | | 438545.9418 | 799597.6563 | | 745309.7915 | | |
| Q96EY1 | DnaJ homolog subfamily A member 3, mitochondrial [OS=Homo sapiens] | -3.27 | 0.557497302 | | 81149.25 | 773979.25 | | 605533.4375 | | | 582410.3781 | 773979.25 | | 629044.3339 | | |
| P54886 | Delta-1-pyrroline-5-carboxylate synthase [OS=Homo sapiens] | -3.68 | 0.568322271 | | 96603.65625 | 844258.625 | | 703168.8438 | | | 693327.0728 | 844258.625 | | 730470.6058 | | |
| P05455 | Lupus La protein [OS=Homo sapiens] | 2.58 | 0.586378848 | | | 834764.125 | | 202353.0469 | 35447.34766 | | | 834764.125 | | 210209.7584 | 299961.9143 | |
| Q9ULP9 | TBC1 domain family member 24 [OS=Homo sapiens] | -3.36 | 0.586378848 | | 69954.82031 | 1059345.844 | | 672575.6406 | | | 502067.6513 | 1059345.844 | | 698689.568 | | |
| Q08170 | Serine/arginine-rich splicing factor 4 [OS=Homo sapiens] | 1.8 | 0.586626322 | 158476.9844 | | 225414.7656 | 238306.25 | 721910.125 | 92669.03125 | 1071481.139 | | 225414.7656 | 1741983.39 | 749939.5502 | 784182.2264 | |
| P60880 | Synaptosomal-associated protein 25 [OS=Homo sapiens] | -4.11 | 0.607729563 | 34400.46875 | 6228337.863 | 61531791 | | 42203551.02 | 28897.01172 | 232585.5301 | 44700950.53 | 61531791 | | 43842177.81 | 244531.7781 | Met-loss+Acetyl [N-Term] |
| Q13564 | NEDD8-activating enzyme E1 regulatory subunit [OS=Homo sapiens] | -3.47 | 0.617887313 | | 77353.15625 | 1150720.719 | | 622386.9375 | | | 555165.7098 | 1150720.719 | | 646552.2006 | | |
| O60925 | Prefoldin subunit 1 [OS=Homo sapiens] | -3.28 | 0.626740922 | | 163449.2539 | 1411929.438 | | 475204.7188 | | | 1173079.748 | 1411929.438 | | 493655.3744 | | |
| P0DOX5 | Immunoglobulin gamma-1 heavy chain [OS=Homo sapiens] | 2.21 | 0.63757335 | 714460.543 | 572064.0967 | 410491.2109 | 1610391.596 | 23082787 | 1419659.641 | 4830549.995 | 4105719.607 | 410491.2109 | 11771724.03 | 23979016.64 | 12013418.54 | |
| P23435 | Cerebellin-1 [OS=Homo sapiens] | -3.46 | 0.643548732 | | 42706.28906 | 965652.2188 | | 678565.7656 | | | 306504.2001 | 965652.2188 | | 704912.2701 | | |
| P62834 | Ras-related protein Rap-1A [OS=Homo sapiens] | -3.54 | 0.645441072 | | 114170.1484 | 1402190.438 | | 586558.625 | | | 819402.2658 | 1402190.438 | | 609332.7911 | | |
| O00220 | Tumor necrosis factor receptor superfamily member 10A [OS=Homo sapien | 3.17 | 0.657188082 | 811762.625 | | | 606414.4688 | 534295.7656 | | 5488420.575 | | | 4432799.944 | 555040.7346 | | |
| P47929 | Galectin-7 [OS=Homo sapiens] | 3.12 | 0.672417768 | 49576.64063 | 40675.20703 | | | 4344794.75 | 319408.7891 | 335193.3756 | 291927.069 | | | 4513489.017 | 2702895.368 | |
| Q9Y5L4 | Mitochondrial import inner membrane translocase subunit Tim13 [OS=Hom | -3.43 | 0.681709864 | | 94685.67188 | 1323245 | | 562174.6875 | | | 679561.6467 | 1323245 | | 584002.1045 | | |
| P17568 | NADH dehydrogenase [ubiquinone] 1 beta subcomplex subunit 7 [OS=Hom | -3.23 | 0.699196901 | | 82764.75 | 1248549.375 | | 651032.5 | | | 594004.8656 | 1248549.375 | | 676309.9772 | | |
| P39687 | Acidic leucine-rich nuclear phosphoprotein 32 family member A [OS=Homo | 2.74 | 0.705748477 | 27895329.41 | 8288011.258 | 29668300.41 | 74433638.73 | 104067829.8 | 31017122.15 | 188603534 | 59483282.6 | 29668300.41 | 544098874 | 108108445.6 | 262472538.8 | |
| P26447 | Protein S100-A4 [OS=Homo sapiens] | 1.14 | 0.714001396 | 77988.3125 | 78596.70313 | | 158742.1406 | 549865.625 | 120465.7344 | 527287.9605 | 564090.6796 | | 1160381.535 | 571215.1211 | 1019402.993 | |
| P63279 | SUMO-conjugating enzyme UBC9 [OS=Homo sapiens] | -3.23 | 0.714001396 | | 82673.25 | 587499.6875 | | 644680.6875 | | | 593348.1676 | 587499.6875 | | 669711.5445 | | |
| Q9Y3A3 | MOB-like protein phocein [OS=Homo sapiens] | -3.17 | 0.714001396 | | 78352.82031 | 823280.75 | | 670473.6875 | | | 562340.3261 | 823280.75 | | 696506.0029 | | |
| Q6IQ22 | Ras-related protein Rab-12 [OS=Homo sapiens] | -3.41 | 0.723809653 | | 68965.14844 | 804489.5625 | | 670814.1719 | | | 494964.7492 | 804489.5625 | | 696859.7072 | | |
| P01860 | Immunoglobulin heavy constant gamma 3 [OS=Homo sapiens] | 1.16 | 0.728196163 | 36884.87109 | 27667.7793 | | 63514.42188 | 425825.6875 | 78538.96094 | 249382.8604 | 198572.4058 | | 464281.0162 | 442359.1157 | 664610.997 | |
| Q9BY67 | Cell adhesion molecule 1 [OS=Homo sapiens] | -3.82 | 0.730218367 | | 413515.4102 | 6356464.688 | | 2501770.813 | | | 2967811.365 | 6356464.688 | | 2598906.447 | | |
| Q92522 | Histone H1.10 [OS=Homo sapiens] | -3.08 | 0.73212506 | | 70878.04688 | 965060.9375 | | 682149.375 | | | 508693.6734 | 965060.9375 | | 708635.0194 | | |
| Q9C004 | Protein sprouty homolog 4 [OS=Homo sapiens] | 1.77 | 0.734171684 | 666685.3125 | 47240.11328 | | 2163879.031 | 2212572.219 | 42311.08203 | 4507536.161 | 339043.58 | | 15817635.2 | 2298479.211 | 358044.0851 | |
| O15540 | Fatty acid-binding protein, brain [OS=Homo sapiens] | -3.8 | 0.738929891 | | 724632.1875 | 2839868.5 | | 2594370.75 | | | 5200704.952 | 2839868.5 | | 2695101.739 | | |
| O43633 | Charged multivesicular body protein 2a [OS=Homo sapiens] | -3.37 | 0.741459634 | | 114295.7266 | 651425.6875 | | 739134.8125 | | | 820303.5435 | 651425.6875 | | 767833.0163 | | |
| O43768 | Alpha-endosulfine [OS=Homo sapiens] | -3.34 | 0.742010726 | | 95053.78125 | 807661.6875 | | 637456.1875 | | | 682203.5777 | 807661.6875 | | 662206.5406 | | |
| P04844 | Dolichyl-diphosphooligosaccharide--protein glycosyltransferase subunit 2 [ | -3.4 | 0.742010726 | | 73987.99219 | 748538.7813 | | 795039.5313 | | | 531013.8357 | 748538.7813 | | 825908.3337 | | |
| Q8N6T3 | ADP-ribosylation factor GTPase-activating protein 1 [OS=Homo sapiens] | -3.07 | 0.747486855 | | 61589.06641 | 1087701.25 | | 601423.875 | | | 442026.4076 | 1087701.25 | | 624775.2105 | | |
| Q9BZC1 | CUGBP Elav-like family member 4 [OS=Homo sapiens] | -3.41 | 0.747486855 | | 48196.90234 | 738823.375 | | 854246.3438 | | | 345910.4812 | 738823.375 | | 887413.9544 | | |
| P56381 | ATP synthase subunit epsilon, mitochondrial [OS=Homo sapiens] | -3.76 | 0.747486855 | | 976526.5781 | 5663232.25 | | 3661150.125 | | | 7008557.856 | 5663232.25 | | 3803300.692 | | |
| P08962 | CD63 antigen [OS=Homo sapiens] | 1.51 | 0.749574851 | 793112.1875 | 276838.8438 | | 2088720.625 | 1957548.75 | 234518.875 | 5362322.82 | 1986879.924 | | 15268238.39 | 2033554.009 | 1984541.449 | |
| P12273 | Prolactin-inducible protein [OS=Homo sapiens] | 2.7 | 0.756948044 | 442925.375 | 212350.4844 | 1077306 | 413593.0313 | 98663952.63 | 829429.0625 | 2994669.459 | 1524045.212 | 1077306 | 3023303.797 | 102494753.4 | 7018779.846 | |
| P0C7P4 | Putative cytochrome b-c1 complex subunit Rieske-like protein 1 [OS=Homo | -3.74 | 0.756948044 | | 1793424.25 | 9197665.875 | | 7651816.25 | | | 12871454.7 | 9197665.875 | | 7948911.42 | | |
| P09496 | Clathrin light chain A [OS=Homo sapiens] | -3.42 | 0.759185637 | | 1685082.258 | 18609779.91 | | 10962420.97 | | | 12093881.27 | 18609779.91 | | 11388056.17 | | |
| Q9UDT6 | CAP-Gly domain-containing linker protein 2 [OS=Homo sapiens] | -3.15 | 0.759185637 | | 50955.375 | 2156408.875 | | 677592.1719 | | | 365708.1146 | 2156408.875 | | 703900.8749 | | |
| Q9NZN3 | EH domain-containing protein 3 [OS=Homo sapiens] | -3.46 | 0.759185637 | | 603594.9063 | 5159389.5 | | 3932427.625 | | | 4332017.087 | 5159389.5 | | 4085111.016 | | |
| Q15056 | Eukaryotic translation initiation factor 4H [OS=Homo sapiens] | -3.44 | 0.759185637 | | 592904.4375 | 6986956.063 | | 4389454.156 | | | 4255291.301 | 6986956.063 | | 4559882.403 | | Met-loss+Acetyl [N-Term] |
| Q96I99 | Succinate--CoA ligase [GDP-forming] subunit beta, mitochondrial [OS=Hom | -3.51 | 0.759185637 | | 736047.875 | 6942970 | | 3944789.75 | | | 5282635.652 | 6942970 | | 4097953.122 | | |
| Q8TDJ6 | DmX-like protein 2 [OS=Homo sapiens] | -3.39 | 0.759185637 | | 972173.9629 | 9335880.594 | | 6257003.523 | | | 6977319.018 | 9335880.594 | | 6499942.646 | | |
| O14773 | Tripeptidyl-peptidase 1 [OS=Homo sapiens] | 1.34 | 0.759185637 | | 25235.12695 | | | 440708.2188 | | | 181113.1937 | | | 457819.4873 | | |
| P35573 | Glycogen debranching enzyme [OS=Homo sapiens] | -3.23 | 0.759185637 | | 60215.60938 | 801852.25 | | 679759.6719 | | | 432169.069 | 801852.25 | | 706152.5319 | | |
| Q92905 | COP9 signalosome complex subunit 5 [OS=Homo sapiens] | -3.46 | 0.759185637 | | 520152.8867 | 4574266.5 | | 3378856.094 | | | 3733151.439 | 4574266.5 | | 3510046.1 | | |
| Q9BTT0 | Acidic leucine-rich nuclear phosphoprotein 32 family member E [OS=Homo | 2.39 | 0.759185637 | 4054648.238 | 1773891.469 | 4435842.344 | 18497415.45 | 18741778.13 | 5826856.875 | 27413943.6 | 12731267.4 | 4435842.344 | 135213367.1 | 19469460.49 | 49307924.51 | |
| Q92930 | Ras-related protein Rab-8B [OS=Homo sapiens] | -3.43 | 0.759185637 | | 497265.5117 | 3585920.438 | | 3046020.781 | | | 3568888.125 | 3585920.438 | | 3164287.874 | | |
| P28070 | Proteasome subunit beta type-4 [OS=Homo sapiens] | -3.41 | 0.759185637 | | 467308.2656 | 4285236.313 | | 3791920.75 | | | 3353884.154 | 4285236.313 | | 3939148.715 | | |
| P51674 | Neuronal membrane glycoprotein M6-a [OS=Homo sapiens] | -3.44 | 0.759185637 | | 9958558.047 | 64622185.94 | | 50398436.69 | | | 71472842.42 | 64622185.94 | | 52355244.27 | | |
| P09417 | Dihydropteridine reductase [OS=Homo sapiens] | -3.37 | 0.759185637 | | 1046588.359 | 13911817.13 | | 7114812.375 | | | 7511393.169 | 13911817.13 | | 7391057.428 | | |
| Q05586 | Glutamate receptor ionotropic, NMDA 1 [OS=Homo sapiens] | -3.23 | 0.759185637 | | 67501.59766 | 1181055.156 | | 723042.7813 | | | 484460.8054 | 1181055.156 | | 751116.1838 | | |
| Q92777 | Synapsin-2 [OS=Homo sapiens] | -3.53 | 0.759185637 | | 6462867.303 | 57497120.94 | | 38956949.81 | | | 46384174.71 | 57497120.94 | | 40469521.63 | | |
| Q9NWB1 | RNA binding protein fox-1 homolog 1 [OS=Homo sapiens] | -3.18 | 0.759185637 | | 94812.0625 | 512094.0625 | | 792107.8125 | | | 680468.7557 | 512094.0625 | | 822862.7858 | | |
| Q9UBQ0 | Vacuolar protein sorting-associated protein 29 [OS=Homo sapiens] | -3.52 | 0.759185637 | | 668062.3125 | 8328825.313 | | 5531629.547 | | | 4794701.418 | 8328825.313 | | 5746404.754 | | |
| Q9NR46 | Endophilin-B2 [OS=Homo sapiens] | -3.38 | 0.759185637 | | 946492.8359 | 12069613.03 | | 6782229.469 | | | 6793004.87 | 12069613.03 | | 7045561.408 | | |
| P63098 | Calcineurin subunit B type 1 [OS=Homo sapiens] | -3.56 | 0.759185637 | | 691925.7031 | 4955848.875 | | 5446985.25 | | | 4965969.622 | 4955848.875 | | 5658473.994 | | |
| Q9H492 | Microtubule-associated proteins 1A/1B light chain 3A [OS=Homo sapiens] | -3.44 | 0.759185637 | | 1389994.211 | 9895627.125 | | 7496802.406 | | | 9976026.321 | 9895627.125 | | 7787878.892 | | |
| Q4KMZ1 | IQ domain-containing protein C [OS=Homo sapiens] | -3.09 | 0.759185637 | | 67706.28906 | 711971.5 | | 647982.5625 | | | 485929.8812 | 711971.5 | | 673141.6206 | | |
| P05771 | Protein kinase C beta type [OS=Homo sapiens] | -3.39 | 0.759185637 | | 497051.6641 | 4479494.719 | | 3341294.875 | | | 3567353.335 | 4479494.719 | | 3471026.501 | | |
| Q9NZR1 | Tropomodulin-2 [OS=Homo sapiens] | -3.44 | 0.759185637 | | 437564.8203 | 4340085.5 | | 3045000.516 | | | 3140414.637 | 4340085.5 | | 3163227.995 | | |
| P54652 | Heat shock-related 70 kDa protein 2 [OS=Homo sapiens] | -3.61 | 0.759185637 | | 652515.0625 | 5450118.75 | | 4185635.25 | | | 4683118.381 | 5450118.75 | | 4348149.871 | | |
| Q08752 | Peptidyl-prolyl cis-trans isomerase D [OS=Homo sapiens] | -3.56 | 0.759185637 | | 1366880.023 | 12413024.31 | | 9096661.281 | | | 9810135.168 | 12413024.31 | | 9449855.09 | | |
| Q14118 | Dystroglycan 1 [OS=Homo sapiens] | 1.67 | 0.759185637 | 135445.6406 | 71223.20313 | 385124.1875 | 399971.75 | 722460.6875 | 185547.3438 | 915763.5717 | 511170.8693 | 385124.1875 | 2923734.249 | 750511.4892 | 1570135.429 | |
| Q9Y5K8 | V-type proton ATPase subunit D [OS=Homo sapiens] | -3.41 | 0.759185637 | | 1794332.68 | 17686135.97 | | 12234370.19 | | | 12877974.53 | 17686135.97 | | 12709391.04 | | |
| Q9NR56 | Muscleblind-like protein 1 [OS=Homo sapiens] | -2.98 | 0.759185637 | | 70019.875 | 717321.5 | | 780549.125 | | | 502534.5505 | 717321.5 | | 810855.3121 | | |
| Q9HC38 | Glyoxalase domain-containing protein 4 [OS=Homo sapiens] | -3.4 | 0.759185637 | | 588577.6289 | 3448164.75 | | 2793817 | | | 4224237.678 | 3448164.75 | | 2902291.839 | | |
| P49588 | Alanine--tRNA ligase, cytoplasmic [OS=Homo sapiens] | -3.37 | 0.759185637 | | 586607.3828 | 6678294.438 | | 3875150.313 | | | 4210097.168 | 6678294.438 | | 4025609.811 | | |
| P48509 | CD151 antigen [OS=Homo sapiens] | 1.77 | 0.759185637 | 662517.125 | 232509.6172 | 806920.7188 | 2451624.063 | 3005568.375 | 330636.1953 | 4479354.565 | 1668727.857 | 806920.7188 | 17921008.76 | 3122264.832 | 2797903.726 | |
| Q16181 | Septin-7 [OS=Homo sapiens] | -3.38 | 0.759185637 | | 4860881.629 | 44596783.38 | | 32415736.42 | | | 34886679.88 | 44596783.38 | | 33674334.17 | | |
| P16401 | Histone H1.5 [OS=Homo sapiens] | -3.36 | 0.759185637 | | 535917.6992 | 2540342.156 | | 7432593.563 | | | 3846295.928 | 2540342.156 | | 7721177.027 | | |
| Q9Y4W6 | AFG3-like protein 2 [OS=Homo sapiens] | -3.49 | 0.759185637 | | 384890.8672 | 4150619.438 | | 3341669.469 | | | 2762372.24 | 4150619.438 | | 3471415.639 | | |
| P10636 | Microtubule-associated protein tau [OS=Homo sapiens] | -3.47 | 0.759185637 | | 11120281.02 | 84457238.75 | | 63880157.19 | | | 79810559.84 | 84457238.75 | | 66360416.19 | | |
| Q9UH03 | Neuronal-specific septin-3 [OS=Homo sapiens] | -3.54 | 0.759185637 | | 5239072.027 | 43058536.47 | | 29987801.73 | | | 37600962.68 | 43058536.47 | | 31152130.66 | | |

| | | | | | | | | | | | | | | | | |
|---|---|---|---|---|---|---|---|---|---|---|---|---|---|---|---|---|
| Q9NZ32 | Actin-related protein 10 [OS=Homo sapiens] | -3.11 | 0.759185637 | | 83374.46094 | 1019421.875 | | 702463.5313 | | | 598380.7776 | 1019421.875 | | 729737.9083 | | |
| P49418 | Amphiphysin [OS=Homo sapiens] | -3.4 | 0.759185637 | | 1840279.16 | 17526364.88 | | 11758349.66 | | | 13207733.67 | 17526364.88 | | 12214888.18 | | |
| P09497 | Clathrin light chain B [OS=Homo sapiens] | -3.42 | 0.759185637 | | 757496.4688 | 13944751.66 | | 9055080.313 | | | 5436572.794 | 13944751.66 | | 9406659.667 | | |
| O95490 | Adhesion G protein-coupled receptor L2 [OS=Homo sapiens] | -3.49 | 0.759185637 | | 1861761.25 | 14463521 | | 10249285 | | | 13361911.21 | 14463521 | | 10647231.44 | | |
| O43759 | Synaptogyrin-1 [OS=Homo sapiens] | -3.52 | 0.759185637 | | 1153716.125 | 7629443.75 | | 6308608.25 | | | 8280252.061 | 7629443.75 | | 6553551.016 | | |
| P13861 | cAMP-dependent protein kinase type II-alpha regulatory subunit [OS=Homo | -3.37 | 0.759185637 | | 2188384.188 | 21163813.84 | | 16110057.63 | | | 15706092.93 | 21163813.84 | | 16735558.83 | | |
| Q10567 | AP-1 complex subunit beta-1 [OS=Homo sapiens] | -3.47 | 0.759185637 | | 934541.7285 | 9934420.844 | | 5781909.781 | | | 6707231.447 | 9934420.844 | | 6006402.557 | | |
| Q96FC7 | Phytanoyl-CoA hydroxylase-interacting protein-like [OS=Homo sapiens] | -3.47 | 0.759185637 | 41600.03906 | 2866397.949 | 22354076.19 | 69846.28906 | 18040120.88 | | 281262.654 | 20572216.17 | 22354076.19 | 510566.0274 | 18740560.17 | | |
| P35611 | Alpha-adducin [OS=Homo sapiens] | -3.42 | 0.759185637 | | 1393943.621 | 11972464.38 | | 8881902.75 | | | 10004371.35 | 11972464.38 | | 9226758.183 | | |
| Q07960 | Rho GTPase-activating protein 1 [OS=Homo sapiens] | -3.44 | 0.759185637 | | 735160.5 | 4778633.375 | | 3881588.125 | | | 5276266.937 | 4778633.375 | | 4032297.583 | | |
| P13693 | Translationally-controlled tumor protein [OS=Homo sapiens] | -3.47 | 0.759185637 | | 651330 | 5204492.594 | | 4401042.688 | | | 4674613.155 | 5204492.594 | | 4571920.879 | | |
| P20339 | Ras-related protein Rab-5A [OS=Homo sapiens] | 1.47 | 0.759185637 | | | 2228988.125 | | 1052233.156 | 20087.79297 | | | 2228988.125 | | 1093087.952 | 169986.564 | |
| Q04917 | 14-3-3 protein eta [OS=Homo sapiens] | -3.44 | 0.759185637 | | 5630607.008 | 37482765.45 | | 32431246.64 | | | 40411019.9 | 37482765.45 | | 33690446.6 | | Met-loss+Acetyl [N-Term] |
| Q16352 | Alpha-internexin [OS=Homo sapiens] | -3.48 | 0.759185637 | | 5193071.598 | 45455816.97 | | 31401154.84 | | | 37270816.34 | 45455816.97 | | 32620359.69 | | |
| Q8TEA8 | D-aminoacyl-tRNA deacylase 1 [OS=Homo sapiens] | -3.37 | 0.759185637 | | 582899.9648 | 4788727.656 | | 3443727.094 | | | 4183488.928 | 4788727.656 | | 3577435.831 | | |
| P36871 | Phosphoglucomutase-1 [OS=Homo sapiens] | -3.53 | 0.759185637 | | 4881627.328 | 35052088.88 | | 27282210.94 | | | 35035572.33 | 35052088.88 | | 28341490.57 | | |
| P05204 | Non-histone chromosomal protein HMG-17 [OS=Homo sapiens] | -3.37 | 0.759185637 | | 712263.375 | 5502986.5 | | 4609687.5 | | | 5111933.648 | 5502986.5 | | 4788666.692 | | |
| O00499 | Myc box-dependent-interacting protein 1 [OS=Homo sapiens] | -3.48 | 0.759185637 | | 1057017.5 | 12750382 | | 7626797.313 | | | 7586243.395 | 12750382 | | 7922921.078 | | |
| Q9H598 | Vesicular inhibitory amino acid transporter [OS=Homo sapiens] | -3.58 | 0.759185637 | | 699090.957 | 10075646.38 | | 6325378.5 | | | 5017394.845 | 10075646.38 | | 6570972.4 | | |
| Q6S8J3 | POTE ankyrin domain family member E [OS=Homo sapiens] | -3.65 | 0.759185637 | | 2769522.75 | 16750988 | | 12087397 | | | 19876940.23 | 16750988 | | 12556711.36 | | |
| Q96FW1 | Ubiquitin thioesterase OTUB1 [OS=Homo sapiens] | -3.51 | 0.759185637 | 50173.68359 | 2978272.207 | 43538887.13 | | 24767795.3 | 112820.5391 | 339230.0518 | 21375140.77 | 43538887.13 | | 25729448.33 | 954707.9571 | Met-loss+Acetyl [N-Term] |
| Q9BUL8 | Programmed cell death protein 10 [OS=Homo sapiens] | -3.7 | 0.759185637 | | 113261.4375 | 908716.6563 | | 936203.0938 | | | 812880.4227 | 908716.6563 | | 972552.8188 | | |
| Q9BT78 | COP9 signalosome complex subunit 4 [OS=Homo sapiens] | -3.59 | 0.759185637 | | 939158.0859 | 7282617.5 | | 4900527.813 | | | 6740363.17 | 7282617.5 | | 5090799.389 | | |
| P06454 | Prothymosin alpha [OS=Homo sapiens] | 2.28 | 0.759185637 | 8755096.594 | 4188464.891 | 9293347.875 | 48017158.38 | 24367459.81 | 15912676.21 | 59194216.14 | 30060726.63 | 9293347.875 | 350998315.4 | 25313569.12 | 134655965.3 | Met-loss+Acetyl [N-Term] |
| Q9BW30 | Tubulin polymerization-promoting protein family member 3 [OS=Homo sapi | -3.36 | 0.759185637 | | 1949677.227 | 14220996.88 | | 12012126.88 | | | 13992886.57 | 14220996.88 | | 12478518.74 | | |
| Q92599 | Septin-8 [OS=Homo sapiens] | -3.55 | 0.759185637 | | 666809.0781 | 6720463.188 | | 4787572.5 | | | 4785706.92 | 6720463.188 | | 4973458.389 | | |
| Q8TB36 | Ganglioside-induced differentiation-associated protein 1 [OS=Homo sapien | -3.59 | 0.759185637 | | 461074.8281 | 6726198.063 | | 3468295.656 | | | 3309146.603 | 6726198.063 | | 3602958.31 | | |
| Q92752 | Tenascin-R [OS=Homo sapiens] | -3.4 | 0.759185637 | | 1878341.266 | 16132457.44 | | 11473706.28 | | | 13480906.44 | 16132457.44 | | 11919193.03 | | |
| O60256 | Phosphoribosyl pyrophosphate synthase-associated protein 2 [OS=Homo s | -3.61 | 0.759185637 | | 484053.4375 | 4311880.625 | | 2795855.656 | | | 3474064.709 | 4311880.625 | | 2904409.65 | | |
| Q13825 | Methylglutaconyl-CoA hydratase, mitochondrial [OS=Homo sapiens] | -3.42 | 0.759185637 | | 659403.6055 | 7241043.563 | | 3777129.156 | | | 4732557.641 | 7241043.563 | | 3923782.812 | | |
| Q96G61 | Diphosphoinositol polyphosphate phosphohydrolase 3-beta [OS=Homo sap | -3.5 | 0.759185637 | | 263869.9336 | 5315899.781 | | 4378762.094 | | | 1893801.702 | 5315899.781 | | 4548775.202 | | |
| P21397 | Amine oxidase [flavin-containing] A [OS=Homo sapiens] | -3.5 | 0.759185637 | | 902724.0781 | 7049692.656 | | 5538955.031 | | | 6478875.303 | 7049692.656 | | 5754014.663 | | |
| P49189 | 4-trimethylaminobutyraldehyde dehydrogenase [OS=Homo sapiens] | -3.58 | 0.759185637 | | 883875.1094 | 4260196.438 | | 3882663.688 | | | 6343595.741 | 4260196.438 | | 4033414.906 | | Met-loss+Acetyl [N-Term] |
| Q9Y6G9 | Cytoplasmic dynein 1 light intermediate chain 1 [OS=Homo sapiens] | -3.47 | 0.759185637 | | 637641.2813 | 6530152.516 | | 4961504.906 | | | 4576368.848 | 6530152.516 | | 5154144.025 | | |
| Q96NT1 | Nucleosome assembly protein 1-like 5 [OS=Homo sapiens] | -3.37 | 0.759185637 | | 2284832.805 | 7894216.531 | | 14521768 | | | 16398307.29 | 7894216.531 | | 15085601.08 | | |
| Q9H936 | Mitochondrial glutamate carrier 1 [OS=Homo sapiens] | -3.53 | 0.759185637 | | 4602216.5 | 25700421.13 | | 23659357.88 | | | 33030233.2 | 25700421.13 | | 24577973.89 | | |
| O94760 | N(G),N(G)-dimethylarginine dimethylaminohydrolase 1 [OS=Homo sapiens] | -3.44 | 0.759185637 | | 739185.332 | 7958464.656 | | 4736855.063 | | | 5305153.267 | 7958464.656 | | 4920771.758 | | |
| O95782 | AP-2 complex subunit alpha-1 [OS=Homo sapiens] | -3.36 | 0.759185637 | 17190.58789 | 7759061.93 | 83622779.41 | 92152.25 | 55769997.72 | | 116227.5441 | 55686998.86 | 83622779.41 | 673619.2979 | 57935365.57 | | |
| P54577 | Tyrosine--tRNA ligase, cytoplasmic [OS=Homo sapiens] | -3.5 | 0.759185637 | | 421406.6914 | 5508949.938 | | 3459807.25 | | | 3024447.305 | 5508949.938 | | 3594140.327 | | |
| P78356 | Phosphatidylinositol 5-phosphate 4-kinase type-2 beta [OS=Homo sapiens] | -3.49 | 0.759185637 | | 395846.1992 | 4324059.063 | | 3411161.297 | | | 2840998.956 | 4324059.063 | | 3543605.61 | | |
| P53007 | Tricarboxylate transport protein, mitochondrial [OS=Homo sapiens] | -3.38 | 0.759185637 | | 785242.2031 | 5881388.469 | | 4406881.938 | | | 5635704.685 | 5881388.469 | | 4577986.848 | | |
| P09960 | Leukotriene A-4 hydrolase [OS=Homo sapiens] | -3.58 | 0.759185637 | | 358088.6875 | 3058021.297 | | 3394715.063 | | | 2570012.266 | 3058021.297 | | 3526520.821 | | |
| Q00765 | Receptor expression-enhancing protein 5 [OS=Homo sapiens] | -2.75 | 0.759185637 | | 41656.06641 | 1068297.75 | | 636882.4375 | | | 298966.7235 | 1068297.75 | | 661610.5138 | | |
| Q16740 | ATP-dependent Clp protease proteolytic subunit, mitochondrial [OS=Homo | -3.46 | 0.759185637 | | 452639.5234 | 3675438.406 | | 3158865.781 | | | 3248606.192 | 3675438.406 | | 3281514.279 | | |
| P52306 | Rap1 GTPase-GDP dissociation stimulator 1 [OS=Homo sapiens] | -3.43 | 0.759185637 | | 1516183.641 | 12683194.84 | | 9086043.219 | | | 10881691.3 | 12683194.84 | | 9438824.763 | | |
| Q02818 | Nucleobindin-1 [OS=Homo sapiens] | -3.58 | 0.759185637 | | 99797.79688 | 980422.5625 | | 969654.2813 | | | 716251.5071 | 980422.5625 | | 1007302.807 | | |
| Q96MZ0 | Ganglioside-induced differentiation-associated protein 1-like 1 [OS=Homo s | -3.43 | 0.759185637 | | 416028.5781 | 4759791.813 | | 3555328.438 | | | 2985848.44 | 4759791.813 | | 3693370.292 | | |
| P60763 | Ras-related C3 botulinum toxin substrate 3 [OS=Homo sapiens] | -3.03 | 0.759185637 | | 72750.78125 | 904263.75 | | 749231.1875 | | | 522134.3391 | 904263.75 | | 778321.4008 | | |
| O95970 | Leucine-rich glioma-inactivated protein 1 [OS=Homo sapiens] | -3.36 | 0.759185637 | | 854550.6406 | 6703185.563 | | 5233831.625 | | | 6133133.229 | 6703185.563 | | 5437044.306 | | |
| Q96GD0 | Chronophin [OS=Homo sapiens] | -3.36 | 0.759185637 | | 1172392.488 | 12138488.19 | | 8198494.719 | | | 8414292.829 | 12138488.19 | | 8516815.637 | | |
| Q9BQI5 | SH3-containing GRB2-like protein 3-interacting protein 1 [OS=Homo sapiens | -3.51 | 0.759185637 | | 530671.8711 | 3863266.625 | | 2665152.344 | | | 3808646.477 | 3863266.625 | | 2768631.552 | | |
| A6NHQ2 | rRNA/tRNA 2'-O-methyltransferase fibrillarin-like protein 1 [OS=Homo sapie | -2.73 | 0.759185637 | | 46095.46875 | 953930.5 | | 723322 | | | 330828.4351 | 953930.5 | | 751406.2437 | | |
| P29966 | Myristoylated alanine-rich C-kinase substrate [OS=Homo sapiens] | -3.49 | 0.759185637 | | 2305313.609 | 16796092.25 | | 12480021.25 | | | 16545298.58 | 16796092.25 | | 12964579.94 | | |
| P55081 | Microfibrillar-associated protein 1 [OS=Homo sapiens] | 1.6 | 0.759185637 | 1511156.578 | 357797.2188 | 696738.9531 | 2457768.953 | 7458654.313 | 1523660.602 | 10217103.62 | 2567920.387 | 696738.9531 | 17965927 | 7748249.631 | 12893493.62 | |
| Q9C040 | Tripartite motif-containing protein 2 [OS=Homo sapiens] | -3.52 | 0.759185637 | | 799410.2383 | 7875303.75 | | 5418506.594 | | | 5737389.06 | 7875303.75 | | 5628889.604 | | |
| P35612 | Beta-adducin [OS=Homo sapiens] | -3.56 | 0.759185637 | | 688484.2031 | 4102371.563 | | 3234456.188 | | | 4941269.883 | 4102371.563 | | 3360039.615 | | |
| O15394 | Neural cell adhesion molecule 2 [OS=Homo sapiens] | -3.53 | 0.759185637 | | 489866.5703 | 5037213.656 | | 3064619.094 | | | 3515785.721 | 5037213.656 | | 3183608.298 | | |
| Q86WG3 | Caytaxin [OS=Homo sapiens] | -3.54 | 0.759185637 | | 5573346.969 | 40308126.06 | | 28696581.69 | | | 40000063.04 | 40308126.06 | | 29810776.74 | | |
| P31323 | cAMP-dependent protein kinase type II-beta regulatory subunit [OS=Homo s | -3.49 | 0.759185637 | | 561962.3125 | 4913360.594 | | 3026986.078 | | | 4033218.827 | 4913360.594 | | 3144514.115 | | |
| Q01813 | ATP-dependent 6-phosphofructokinase, platelet type [OS=Homo sapiens] | 0.37 | 0.759185637 | 487223.8535 | 1534776.856 | 14756682.34 | 955435.8633 | 13235173.22 | 270406.8281 | 3294176.573 | 11015135.31 | 14756682.34 | 6984094.641 | 13749052 | 2288231.847 | |
| Q96C19 | EF-hand domain-containing protein D2 [OS=Homo sapiens] | -3.37 | 0.759185637 | | 1222763.477 | 9355212.813 | | 7646148.063 | | | 8775806.785 | 9355212.813 | | 7943023.155 | | |
| Q92561 | Phytanoyl-CoA hydroxylase-interacting protein [OS=Homo sapiens] | -3.53 | 0.759185637 | | 600526.7344 | 4625541.813 | | 4122223.938 | | | 4309996.734 | 4625541.813 | | 4282276.503 | | |
| P78508 | ATP-sensitive inward rectifier potassium channel 10 [OS=Homo sapiens] | -3.55 | 0.759185637 | | 947442.8672 | 11088697 | | 7408355.25 | | | 6799823.27 | 11088697 | | 7695997.62 | | |
| Q15185 | Prostaglandin E synthase 3 [OS=Homo sapiens] | 1.64 | 0.759185637 | 451670.8203 | 197573.293 | 7371318.156 | 1458962.023 | 5395717.063 | 517344.9063 | 3053798.422 | 1417988.906 | 7371318.156 | 10664796.29 | 5605215.229 | 4377866.855 | |
| Q9UHG2 | ProSAAS [OS=Homo sapiens] | -3.5 | 0.759185637 | | 10845356.41 | 84107368.5 | | 66846029.25 | 24017.56445 | | 77837418.4 | 84107368.5 | | 69441443.43 | 203241.0063 | |
| Q15102 | Platelet-activating factor acetylhydrolase IB subunit alpha1 [OS=Homo sapi | -3.46 | 0.759185637 | | 978042.1016 | 7072382.906 | | 6272333.563 | | | 7019434.809 | 7072382.906 | | 6515867.9 | | |
| Q9UNS2 | COP9 signalosome complex subunit 3 [OS=Homo sapiens] | -3.62 | 0.759185637 | | 520845.418 | 5428499.031 | | 3279564.563 | | | 3738121.755 | 5428499.031 | | 3406899.401 | | |
| Q9UBI6 | Guanine nucleotide-binding protein G(I)/G(S)/G(O) subunit gamma-12 [OS= | -3.46 | 0.759185637 | | 483017.9375 | 4363380.75 | | 3275187.125 | | | 3466632.897 | 4363380.75 | | 3402352.002 | | |
| E9PAV3 | Nascent polypeptide-associated complex subunit alpha, muscle-specific fo | 2.37 | 0.759185637 | 1543367.656 | 1724955.496 | 8825161.516 | 9234335.742 | 15658959.66 | 5378941.891 | 10434886.43 | 12380052.59 | 8825161.516 | 67501626.49 | 16266946.19 | 45517586.3 | |
| P00491 | Purine nucleoside phosphorylase [OS=Homo sapiens] | -3.48 | 0.759185637 | | 121311.7813 | 1230818.75 | | 668908.75 | | | 870657.9591 | 1230818.75 | | 694880.3039 | | |
| Q9Y5J7 | Mitochondrial import inner membrane translocase subunit Tim9 [OS=Homo | -3.54 | 0.759185637 | | 690337.2266 | 4834211.563 | | 3738418.219 | | | 4954569.083 | 4834211.563 | | 3883568.855 | | Met-loss+Acetyl [N-Term] |
| O94812 | BAI1-associated protein 3 [OS=Homo sapiens] | -3.45 | 0.759185637 | | 371657.0234 | 3278654.875 | | 2845762.375 | | | 2667392.583 | 3278654.875 | | 2956254.084 | | |
| Q9ULU8 | Calcium-dependent secretion activator 1 [OS=Homo sapiens] | -3.41 | 0.759185637 | | 2136354.693 | 27518670.09 | | 17754462.13 | | | 15332675.84 | 27518670.09 | | 18443810.22 | | |
| P62993 | Growth factor receptor-bound protein 2 [OS=Homo sapiens] | -3.4 | 0.759185637 | | 715917.2578 | 5098331.813 | | 5060959.75 | | | 5138157.664 | 5098331.813 | | 5257460.378 | | |
| Q9NVA2 | Septin-11 [OS=Homo sapiens] | -3.46 | 0.759185637 | | 7642284.465 | 52593257.56 | | 38841290.78 | | | 54848883.81 | 52593257.56 | | 40349371.93 | | Met-loss+Acetyl [N-Term] |
| P78559 | Microtubule-associated protein 1A [OS=Homo sapiens] | -3.4 | 0.759185637 | | 1601146.738 | 19901559.69 | | 11548713.63 | | | 11491473.76 | 19901559.69 | | 11997112.66 | | |
| Q96HN2 | Adenosylhomocysteinase 3 [OS=Homo sapiens] | -3.43 | 0.759185637 | | 565512.875 | 6927919.172 | | 3665378.875 | | | 4058701.311 | 6927919.172 | | 3807693.63 | | Met-loss+Acetyl [N-Term] |
| P20916 | Myelin-associated glycoprotein [OS=Homo sapiens] | -3.55 | 0.759185637 | | 698459.3906 | 11117581.53 | | 4102910.156 | | | 5012862.076 | 11117581.53 | | 4262212.83 | | |
| Q92871 | Phosphomannomutase 1 [OS=Homo sapiens] | -3.39 | 0.759185637 | | 117103.5859 | 1048296.906 | | 719633.6875 | | | 840455.6267 | 1048296.906 | | 747574.726 | | |
| Q16623 | Syntaxin-1A [OS=Homo sapiens] | -3.53 | 0.759185637 | | 1120166.816 | 10807831.31 | | 7114265.156 | | | 8039467.76 | 10807831.31 | | 7390488.963 | | |
| Q9P2X3 | Protein IMPACT [OS=Homo sapiens] | -3.41 | 0.759185637 | | 2563453.977 | 15863704.47 | | 12809491.89 | | | 18397979.03 | 15863704.47 | | 13306842.85 | | |
| P10114 | Ras-related protein Rap-2a [OS=Homo sapiens] | -3.49 | 0.759185637 | | 928064.9453 | 7230969.063 | | 5420543.727 | | | 6660747.397 | 7230969.063 | | 5631005.832 | | |

| | | | | | | | | | | | | | | | | |
|---|---|---|---|---|---|---|---|---|---|---|---|---|---|---|---|---|
| Q12926 | ELAV-like protein 2 [OS=Homo sapiens] | -3.5 | 0.759185637 | | 778028.5391 | 6934263.563 | | 5118941.594 | | | 5583932.022 | 6934263.563 | | 5317693.468 | | |
| Q13557 | Calcium/calmodulin-dependent protein kinase type II subunit delta [OS=Ho | -3.46 | 0.759185637 | | 1482689.891 | 11441753.94 | | 8881786.594 | | | 10641305.74 | 11441753.94 | | 9226637.516 | | |
| Q8NCB2 | CaM kinase-like vesicle-associated protein [OS=Homo sapiens] | -3.49 | 0.759185637 | | 1154598.219 | 9493280.375 | | 7720216.188 | | | 8286582.872 | 9493280.375 | | 8019967.104 | | Met-loss [N-Term] |
| Q7L5N1 | COP9 signalosome complex subunit 6 [OS=Homo sapiens] | -3.4 | 0.759185637 | | 521991.6719 | 6231458.969 | | 2772654.719 | | | 3746348.45 | 6231458.969 | | 2880307.895 | | |
| P59768 | Guanine nucleotide-binding protein G(I)/G(S)/G(O) subunit gamma-2 [OS=H | -3.55 | 0.759185637 | | 1047453.305 | 6393531.688 | | 7340858.625 | | | 7517600.905 | 6393531.688 | | 7625880.32 | | Met-loss+Acetyl [N-Term] |
| P26196 | Probable ATP-dependent RNA helicase DDX6 [OS=Homo sapiens] | -3.37 | 0.759185637 | | 510738.2852 | 4740823.594 | | 3557112 | | | 3665582.587 | 4740823.594 | | 3695223.104 | | |
| Q9BPX5 | Actin-related protein 2/3 complex subunit 5-like protein [OS=Homo sapiens | -3.68 | 0.759185637 | | 554510.9688 | 4989873.5 | | 3195552.688 | | | 3979740.33 | 4989873.5 | | 3319625.618 | | |
| P23381 | Tryptophan--tRNA ligase, cytoplasmic [OS=Homo sapiens] | -3.59 | 0.759185637 | | 428915.25 | 4499506.938 | | 3086768.781 | | | 3078336.435 | 4499506.938 | | 3206617.986 | | |
| Q9NZW5 | Protein PALS2 [OS=Homo sapiens] | -3.41 | 0.759185637 | | 481159.9219 | 8547137.5 | | 4254740.422 | | | 3453297.868 | 8547137.5 | | 4419938.172 | | |
| Q01105 | Protein SET [OS=Homo sapiens] | 2.24 | 0.759185637 | 3925967.836 | 2969731.957 | 10013909.87 | 27081047.3 | 36512309.38 | 9789768.262 | 26543920.58 | 21313847.16 | 10013909.87 | 197958444.5 | 37929963.74 | 82842802.69 | |
| O95670 | V-type proton ATPase subunit G 2 [OS=Homo sapiens] | -3.52 | 0.759185637 | | 609624.5859 | 4420424.688 | | 2726290.063 | | | 4375292.262 | 4420424.688 | | 2832143.05 | | Met-loss+Acetyl [N-Term] |
| Q8WU79 | Stromal membrane-associated protein 2 [OS=Homo sapiens] | -3.37 | 0.759185637 | | 111263.9766 | 1055695.719 | | 718337.9375 | | | 798544.5911 | 1055695.719 | | 746228.6662 | | |
| O95294 | RasGAP-activating-like protein 1 [OS=Homo sapiens] | -3.43 | 0.759185637 | | 709162.4648 | 4869407.344 | | 3770062.766 | | | 5089678.331 | 4869407.344 | | 3916442.056 | | |
| Q14012 | Calcium/calmodulin-dependent protein kinase type 1 [OS=Homo sapiens] | -3.36 | 0.759185637 | | 582616.5547 | 4097213.938 | | 3482212.375 | | | 4181454.886 | 4097213.938 | | 3617415.37 | | |
| Q14203 | Dynactin subunit 1 [OS=Homo sapiens] | -3.48 | 0.759185637 | | 780310.3672 | 8499825.406 | | 5167904.094 | | | 5600308.764 | 8499825.406 | | 5368557.023 | | |
| P62714 | Serine/threonine-protein phosphatase 2A catalytic subunit beta isoform [OS | -3.45 | 0.759185637 | | 1375892.578 | 17905652.63 | | 11570658.97 | | | 9874818.519 | 17905652.63 | | 12019910.07 | | |
| Q9UGM3 | Deleted in malignant brain tumors 1 protein [OS=Homo sapiens] | 1.75 | 0.759185637 | 948598.9375 | 203563.8086 | 171192.2344 | 1352947.313 | 23947861.75 | 316615.3555 | 6413586.639 | 1460983 | 171192.2344 | 9889844.456 | 24877679.43 | 2679256.824 | |
| P04350 | Tubulin beta-4A chain [OS=Homo sapiens] | -3.53 | 0.759185637 | | 7917211.047 | 79090393.63 | | 49208670.38 | 26771.18945 | | 56822039.38 | 79090393.63 | | 51119283.2 | 226542.6828 | |
| P10644 | cAMP-dependent protein kinase type I-alpha regulatory subunit [OS=Homo | -3.58 | 0.759185637 | | 383047.7305 | 3494551.969 | | 2782646.188 | | | 2749143.997 | 3494551.969 | | 2890687.3 | | |
| Q13449 | Limbic system-associated membrane protein [OS=Homo sapiens] | -3.41 | 0.759185637 | | 2167182.488 | 19519949.5 | | 14812871.47 | | | 15553927.76 | 19519949.5 | | 15388007.15 | | |
| P09972 | Fructose-bisphosphate aldolase C [OS=Homo sapiens] | -3.43 | 0.759185637 | | 6775033.352 | 74230767.97 | | 46696368.84 | | | 48624598.95 | 74230767.97 | | 48509437.16 | | Met-loss [N-Term] |
| O43761 | Synaptogyrin-3 [OS=Homo sapiens] | -3.54 | 0.759185637 | | 715006.625 | 3568438.594 | | 3630385.5 | | | 5131622.028 | 3568438.594 | | 3771341.576 | | Acetyl [N-Term] |
| P57105 | Synaptojanin-2-binding protein [OS=Homo sapiens] | -3.58 | 0.759185637 | | 119611.6094 | 1910751.5 | | 577127.4375 | | | 858455.7792 | 1910751.5 | | 599535.4212 | | |
| P21333 | Filamin-A [OS=Homo sapiens] | 1.18 | 0.759185637 | 138472.4336 | 37642.80078 | | 277597.7656 | 620263.9375 | 71977.60156 | 936228.0674 | 270163.4068 | | 2029198.549 | 644346.7714 | 609087.5785 | |
| Q00577 | Transcriptional activator protein Pur-alpha [OS=Homo sapiens] | -3.36 | 0.759185637 | | 5287854.297 | 39160081.91 | | 37116430.06 | | | 37951074.35 | 39160081.91 | | 38557540.48 | | |
| O15075 | Serine/threonine-protein kinase DCLK1 [OS=Homo sapiens] | -3.42 | 0.759185637 | | 336482.7148 | 4392266.313 | | 3177218.313 | | | 2414945.612 | 4392266.313 | | 3300579.379 | | |
| O15217 | Glutathione S-transferase A4 [OS=Homo sapiens] | -3.47 | 0.759185637 | | 507397.875 | 3868312.625 | | 2851054.25 | | | 3641608.373 | 3868312.625 | | 2961751.425 | | |
| Q12979 | Active breakpoint cluster region-related protein [OS=Homo sapiens] | -3.51 | 0.759185637 | | 577335.7891 | 5444727.906 | | 4044411.906 | | | 4143554.687 | 5444727.906 | | 4201443.283 | | |
| Q7Z6L0 | Proline-rich transmembrane protein 2 [OS=Homo sapiens] | -3.74 | 0.759185637 | | 174239.2656 | 973221.3125 | | 676433.5625 | | | 1250519.956 | 973221.3125 | | 702697.2805 | | |
| Q13098 | COP9 signalosome complex subunit 1 [OS=Homo sapiens] | -3.43 | 0.759185637 | | 719494.9648 | 4075924.469 | | 3486315.125 | | | 5163834.965 | 4075924.469 | | 3621677.417 | | |
| P08247 | Synaptophysin [OS=Homo sapiens] | -3.44 | 0.759185637 | | 5569260.813 | 26738807.75 | | 26203593.41 | | | 39970736.58 | 26738807.75 | | 27220993.82 | | |
| Q16653 | Myelin-oligodendrocyte glycoprotein [OS=Homo sapiens] | -3.36 | 0.759185637 | | 1031544.613 | 10181116.59 | | 5966532.719 | | | 7403423.793 | 10181116.59 | | 6198193.803 | | |
| O43301 | Heat shock 70 kDa protein 12A [OS=Homo sapiens] | -3.38 | 0.759185637 | | 1882535.926 | 20007442.78 | | 12790228.47 | | | 13511011.63 | 20007442.78 | | 13286831.49 | | |
| P35998 | 26S proteasome regulatory subunit 7 [OS=Homo sapiens] | -3.56 | 0.759185637 | | 649365.2344 | 5386121.125 | | 2189912.563 | | | 4660511.979 | 5386121.125 | | 2274939.754 | | Met-loss [N-Term] |
| Q9P0L0 | Vesicle-associated membrane protein-associated protein A [OS=Homo sap | -3.62 | 0.759185637 | | 929184.0625 | 5102904.5 | | 4033039.438 | | | 6668779.332 | 5102904.5 | | 4189629.258 | | |
| P55036 | 26S proteasome non-ATPase regulatory subunit 4 [OS=Homo sapiens] | 1.22 | 0.759185637 | | 46950.03516 | 505179.9375 | | 757207.9375 | 66570.5 | | 336961.6814 | 505179.9375 | | 786607.8621 | 563331.6999 | |
| O95757 | Heat shock 70 kDa protein 4L [OS=Homo sapiens] | -3.48 | 0.759185637 | | 2025985.906 | 18137467.06 | | 12740548.53 | | | 14540556.05 | 18137467.06 | | 13235222.65 | | |
| Q15223 | Nectin-1 [OS=Homo sapiens] | -2.99 | 0.759185637 | | 64689.15625 | 499660.2188 | | 710825.375 | | | 464275.8368 | 499660.2188 | | 738424.4153 | | |
| Q5H9L2 | Transcription elongation factor A protein-like 5 [OS=Homo sapiens] | -3.41 | 0.759185637 | | 466681.0469 | 3432541.125 | | 2842851.25 | | | 3349382.588 | 3432541.125 | | 2953229.93 | | |
| Q4G0F5 | Vacuolar protein sorting-associated protein 26B [OS=Homo sapiens] | -3.41 | 0.759185637 | | 527707.9355 | 4927681.813 | | 2828394.594 | | | 3787374.23 | 4927681.813 | | 2938211.968 | | |
| P61081 | NEDD8-conjugating enzyme Ubc12 [OS=Homo sapiens] | -3.55 | 0.759185637 | | 437004.9453 | 3867002.281 | | 3067881.656 | | | 3136396.398 | 3867002.281 | | 3186997.536 | | |
| P49591 | Serine--tRNA ligase, cytoplasmic [OS=Homo sapiens] | -3.43 | 0.759185637 | | 696606.8281 | 8278798.313 | | 5403489.516 | | | 4999566.184 | 8278798.313 | | 5613289.461 | | Met-loss [N-Term] |
| Q9NTK5 | Obg-like ATPase 1 [OS=Homo sapiens] | -3.47 | 0.759185637 | | 1117326.973 | 9367977.719 | | 7064093.719 | | | 8019086.124 | 9367977.719 | | 7338369.531 | | |
| P40394 | All-trans-retinol dehydrogenase [NAD(+)] ADH7 [OS=Homo sapiens] | -3.36 | 0.759185637 | | 449366.3438 | 4430905.5 | | 2928962 | | | 3225114.492 | 4430905.5 | | 3042684.08 | | |
| Q9ULD0 | 2-oxoglutarate dehydrogenase-like, mitochondrial [OS=Homo sapiens] | -3.47 | 0.759185637 | | 759308.375 | 7248119.75 | | 5466037.031 | | | 5449576.894 | 7248119.75 | | 5678265.494 | | |
| P05413 | Fatty acid-binding protein, heart [OS=Homo sapiens] | -3.61 | 0.759185637 | | 1094585.563 | 6104105.25 | | 5047125.75 | | | 7855870.403 | 6104105.25 | | 5243089.248 | | |
| Q00169 | Phosphatidylinositol transfer protein alpha isoform [OS=Homo sapiens] | -3.36 | 0.759185637 | | 1226374.828 | 10728605.53 | | 8695000.969 | | | 8801725.553 | 10728605.53 | | 9032599.612 | | |
| Q06481 | Amyloid beta precursor like protein 2 [OS=Homo sapiens] | -2.11 | 0.759185637 | | 105078.9219 | 239349.8281 | 359503.3828 | 643468.9844 | | | 754154.2851 | 239349.8281 | 2627916.479 | 668452.7949 | | |
| Q99962 | Endophilin-A1 [OS=Homo sapiens] | -3.37 | 0.759185637 | | 3304278.65 | 21659520.89 | | 13659630.03 | | | 23714897.9 | 21659520.89 | | 14189989.09 | | Met-loss+Acetyl [N-Term] |
| P54578 | Ubiquitin carboxyl-terminal hydrolase 14 [OS=Homo sapiens] | -3.35 | 0.759389029 | | 428560.1563 | 4697637.563 | | 2667227.188 | | | 3075787.917 | 4697637.563 | | 2770786.955 | | Met-loss [N-Term] |
| Q7L1I2 | Synaptic vesicle glycoprotein 2B [OS=Homo sapiens] | -3.72 | 0.760220815 | | 133796.9531 | 1111477.656 | | 948375.5938 | | | 960264.3778 | 1111477.656 | | 985197.9374 | | |
| P28066 | Proteasome subunit alpha type-5 [OS=Homo sapiens] | -3.32 | 0.760220815 | | 606454 | 6417519.438 | | 5133951.219 | | | 4352536.88 | 6417519.438 | | 5333285.868 | | |
| Q8WXD9 | Caskin-1 [OS=Homo sapiens] | -3.32 | 0.760220815 | | 241536.7773 | 5113223.5 | | 3257908.063 | | | 1733516.032 | 5113223.5 | | 3384402.05 | | |
| P63151 | Serine/threonine-protein phosphatase 2A 55 kDa regulatory subunit B alpha | -3.32 | 0.760220815 | | 660288.4121 | 8683901.813 | | 6727129.219 | | | 4738907.923 | 8683901.813 | | 6988321.795 | | |
| O00429 | Dynamin-1-like protein [OS=Homo sapiens] | -3.32 | 0.760220815 | | 3138482.742 | 37269094.02 | | 24722232.09 | | | 22524976.15 | 37269094.02 | | 25682116.06 | | |
| Q15257 | Serine/threonine-protein phosphatase 2A activator [OS=Homo sapiens] | -3.33 | 0.760220815 | | 548080.1875 | 4254482.25 | | 3597086.625 | | | 3933586.438 | 4254482.25 | | 3736749.814 | | |
| Q9BY32 | Inosine triphosphate pyrophosphatase [OS=Homo sapiens] | -3.33 | 0.760220815 | | 512739.2148 | 4781971.75 | | 3449080.031 | | | 3679943.314 | 4781971.75 | | 3582996.605 | | Met-loss+Acetyl [N-Term] |
| Q6ZVM7 | TOM1-like protein 2 [OS=Homo sapiens] | -3.33 | 0.760220815 | | 451159.8906 | 4742887.25 | | 2921936.719 | | | 3237986.827 | 4742887.25 | | 3035386.03 | | |
| Q9H4G0 | Band 4.1-like protein 1 [OS=Homo sapiens] | -3.32 | 0.760220815 | | 1187233.172 | 15690609.31 | | 11006826.41 | | | 8520804.82 | 15690609.31 | | 11434185.72 | | |
| P41236 | Protein phosphatase inhibitor 2 [OS=Homo sapiens] | -3 | 0.760220815 | | 75037.80469 | 733947.8125 | | 805580.3125 | | | 538548.3686 | 733947.8125 | | 836858.3793 | | |
| Q9Y376 | Calcium-binding protein 39 [OS=Homo sapiens] | -3.35 | 0.760220815 | | 979260.7891 | 7315917.969 | | 6569997.375 | | | 7028181.362 | 7315917.969 | | 6825089.032 | | |
| Q13554 | Calcium/calmodulin-dependent protein kinase type II subunit beta [OS=Hor | -3.34 | 0.760220815 | | 3365066.543 | 47647468.75 | | 31243319.63 | | | 24151174.26 | 47647468.75 | | 32456396.24 | | Met-loss+Acetyl [N-Term] |
| P00568 | Adenylate kinase isoenzyme 1 [OS=Homo sapiens] | -3.33 | 0.760220815 | | 1217155.656 | 16064782.13 | | 7172625 | | | 8735559.306 | 16064782.13 | | 7451114.73 | | |
| P62195 | 26S proteasome regulatory subunit 8 [OS=Homo sapiens] | -3.34 | 0.760220815 | | 488142.4727 | 6797551.156 | | 4206005.781 | | | 3503411.825 | 6797551.156 | | 4369311.323 | | |
| P30085 | UMP-CMP kinase [OS=Homo sapiens] | -3.33 | 0.760220815 | | 643583.2656 | 7865078.469 | | 4853219.656 | | | 4619014.631 | 7865078.469 | | 5041654.411 | | |
| Q01433 | AMP deaminase 2 [OS=Homo sapiens] | -3.32 | 0.760220815 | | 310103.5391 | 3616085.375 | | 3320727.813 | | | 2225621.548 | 3616085.375 | | 3449660.887 | | |
| P51649 | Succinate-semialdehyde dehydrogenase, mitochondrial [OS=Homo sapiens | -3.31 | 0.760222974 | | 816556.0156 | 6744212.375 | | 5804446.688 | | | 5860444.771 | 6744212.375 | | 6029814.498 | | |
| O00562 | Membrane-associated phosphatidylinositol transfer protein 1 [OS=Homo sa | -3.35 | 0.760222974 | | 148488.9453 | 1498020.438 | | 812072.9688 | | | 1065709.206 | 1498020.438 | | 843603.1243 | | |
| Q9NP90 | Ras-related protein Rab-9B [OS=Homo sapiens] | -2.78 | 0.760222974 | | 38125.98828 | 2715792.313 | | 555895.25 | | | 273631.2566 | 2715792.313 | | 577478.8568 | | |
| Q9H0Q3 | FXYD domain-containing ion transport regulator 6 [OS=Homo sapiens] | -3.31 | 0.76049737 | | 897104.625 | 7776102.938 | | 6327898.5 | | | 6438544.335 | 7776102.938 | | 6573590.244 | | |
| P42658 | Dipeptidyl aminopeptidase-like protein 6 [OS=Homo sapiens] | -3.3 | 0.763778241 | | 1730011.539 | 17327132.16 | | 12242331.5 | | | 12416339.95 | 17327132.16 | | 12717661.47 | | |
| Q6IBS0 | Twinfilin-2 [OS=Homo sapiens] | -3.64 | 0.765683993 | | 137816.875 | 900763.0938 | | 901411.4063 | | | 989115.4666 | 900763.0938 | | 936410.2831 | | Met-loss+Acetyl [N-Term] |
| Q9H444 | Charged multivesicular body protein 4b [OS=Homo sapiens] | -3.09 | 0.765683993 | | 83326.14844 | 718063.875 | | 794962.6875 | | | 598034.0375 | 718063.875 | | 825828.5064 | | |
| P62333 | 26S proteasome regulatory subunit 10B [OS=Homo sapiens] | -3.37 | 0.765683993 | | 294855.7617 | 3249912.063 | | 2967474.594 | | | 2116187.835 | 3249912.063 | | 3082691.993 | | |
| Q9BPU6 | Dihydropyrimidinase-related protein 5 [OS=Homo sapiens] | -3.28 | 0.765683993 | | 5618448.867 | 54872581.28 | | 41419785.13 | | | 40323760.59 | 54872581.28 | | 43027980.84 | | |
| O60282 | Kinesin heavy chain isoform 5C [OS=Homo sapiens] | -3.28 | 0.765683993 | | 740431.9766 | 8730387.813 | | 5633786.719 | | | 5314100.468 | 8730387.813 | | 5852528.357 | | |
| Q7Z3B1 | Neuronal growth regulator 1 [OS=Homo sapiens] | -3.3 | 0.765683993 | | 634631.7031 | 8249547.688 | | 4457845.938 | | | 4554769.023 | 8249547.688 | | 4630929.615 | | |
| Q7L099 | Protein RUFY3 [OS=Homo sapiens] | -3.29 | 0.765683993 | | 1062719.297 | 10350513.19 | | 8775730 | | | 7627165.347 | 10350513.19 | | 9116463.09 | | |
| P48539 | Calmodulin regulator protein PCP4 [OS=Homo sapiens] | -3.29 | 0.765683993 | | 1194519.813 | 10899712.22 | | 8519509 | | | 8573101.238 | 10899712.22 | | 8850293.86 | | |
| P20336 | Ras-related protein Rab-3A [OS=Homo sapiens] | -3.29 | 0.765683993 | | 7295697.156 | 78464236.19 | | 59172507.66 | | | 52361417.26 | 78464236.19 | | 61469983.92 | | |
| Q9UI15 | Transgelin-3 [OS=Homo sapiens] | -3.29 | 0.765683993 | | 1916271.059 | 20695024.22 | | 14146208.78 | | | 13753129.6 | 20695024.22 | | 14695460.12 | | |
| O15020 | Spectrin beta chain, non-erythrocytic 2 [OS=Homo sapiens] | -3.27 | 0.768488877 | | 732218.3242 | 13790191.31 | | 6651048.172 | | | 5255150.862 | 13790191.31 | | 6909286.768 | | |

| | | | | | | | | | | | | | | | | |
|---|---|---|---|---|---|---|---|---|---|---|---|---|---|---|---|---|
| Q9NUP9 | Protein lin-7 homolog C [OS=Homo sapiens] | -3.24 | 0.768816825 | | 682592.7109 | 7852586.688 | | 5737637 | | | 4898986.483 | 7852586.688 | | 5960410.807 | | |
| P35080 | Profilin-2 [OS=Homo sapiens] | -3.27 | 0.768816825 | | 2530198.781 | 30304366.28 | | 17752262.44 | | | 18159305.59 | 30304366.28 | | 18441525.12 | | |
| P54920 | Alpha-soluble NSF attachment protein [OS=Homo sapiens] | -3.23 | 0.768816825 | | 543372.8125 | 9302253.156 | | 5669845.172 | | | 3899801.479 | 9302253.156 | | 5889986.842 | | |
| Q9P0L2 | Serine/threonine-protein kinase MARK1 [OS=Homo sapiens] | -3.19 | 0.768816825 | | 97618.05469 | 755301.0625 | | 810434.75 | | | 700607.438 | 755301.0625 | | 841901.2988 | | |
| O14531 | Dihydropyrimidinase-related protein 4 [OS=Homo sapiens] | -3.24 | 0.768816825 | | 826316.1172 | 12207135.97 | | 6930197 | | | 5930493.286 | 12207135.97 | | 7199274.038 | | |
| P62140 | Serine/threonine-protein phosphatase PP1-beta catalytic subunit [OS=Hom | -3.24 | 0.768816825 | | 442423.5098 | 6149578.75 | | 3774431.938 | | | 3175285.583 | 6149578.75 | | 3920980.869 | | Met-loss+Acetyl [N-Term] |
| Q16849 | Receptor-type tyrosine-protein phosphatase-like N [OS=Homo sapiens] | -3.25 | 0.768816825 | | 508148.2188 | 5985340.719 | | 5058061.75 | | | 3646993.61 | 5985340.719 | | 5254449.858 | | |
| Q15084 | Protein disulfide-isomerase A6 [OS=Homo sapiens] | -3.44 | 0.768816825 | | 111539.2891 | 1868564.688 | | 654148.5 | | | 800520.5164 | 1868564.688 | | 679546.9614 | | |
| P53985 | Monocarboxylate transporter 1 [OS=Homo sapiens] | 1.06 | 0.768816825 | 341964.5391 | 131516.1523 | 205721.2969 | 476636.625 | 1681215.656 | 230688.3047 | 2312061.623 | 943895.0084 | 205721.2969 | 3484143.129 | 1746491.799 | 1952126.465 | |
| Q16851 | UTP--glucose-1-phosphate uridylyltransferase [OS=Homo sapiens] | -3.26 | 0.768816825 | | 379524.4453 | 2792067.656 | | 3031499.406 | | | 2723857.284 | 2792067.656 | | 3149202.681 | | |
| P07602 | Prosaposin [OS=Homo sapiens] | 1.08 | 0.768816825 | | 99125.74219 | 1292851.5 | 91150.17188 | 1444106 | | | 711428.1523 | 1292851.5 | 666294.2552 | 1500175.945 | | |
| P39060 | Collagen alpha-1(XVIII) chain [OS=Homo sapiens] | 1.03 | 0.768816825 | 70752.42188 | | | 221242.4688 | | | 478365.2708 | | | 1617249.676 | | | |
| P27361 | Mitogen-activated protein kinase 3 [OS=Homo sapiens] | -3.22 | 0.768816825 | | 687401.3828 | 7386717.344 | | 6042224.125 | | | 4933498.452 | 7386717.344 | | 6276824.061 | | |
| P13521 | Secretogranin-2 [OS=Homo sapiens] | -3.26 | 0.768816825 | | 2624380.598 | 64856867.81 | | 89058503.34 | | | 18835251.05 | 64856867.81 | | 92516355.74 | | |
| Q08945 | FACT complex subunit SSRP1 [OS=Homo sapiens] | 1.17 | 0.768816825 | 1388417.422 | 162186.0664 | 1190792.75 | 852447.0078 | 2326619.75 | 350734.3984 | 9387250.054 | 1164013.817 | 1190792.75 | 6231261.363 | 2416954.837 | 2967978.382 | Met-loss+Acetyl [N-Term] |
| Q9NUQ9 | CYFIP-related Rac1 interactor B [OS=Homo sapiens] | -3.23 | 0.768816825 | | 691686.7266 | 10962008.91 | | 6343679.219 | | | 4964254.481 | 10962008.91 | | 6589983.677 | | |
| Q07866 | Kinesin light chain 1 [OS=Homo sapiens] | -3.25 | 0.768816825 | | 462930.3594 | 5218114.625 | | 3006943.344 | | | 3322463.801 | 5218114.625 | | 3123693.186 | | |
| Q9BVA1 | Tubulin beta-2B chain [OS=Homo sapiens] | -3.25 | 0.768816825 | | 2324357.5 | 18535658.75 | | 14735261.25 | | | 16681977.1 | 18535658.75 | | 15307383.58 | | |
| Q9UNZ2 | NSFL1 cofactor p47 [OS=Homo sapiens] | -3.26 | 0.768816825 | | 195712.4688 | 4704673.438 | | 3452491.75 | | | 1404633.72 | 4704673.438 | | 3586540.79 | | |
| O75955 | Flotillin-1 [OS=Homo sapiens] | -3.22 | 0.768816825 | | 1011813.563 | 10906728.75 | | 8035622.313 | | | 7261813.504 | 10906728.75 | | 8347619.425 | | |
| Q12974 | Protein tyrosine phosphatase type IVA 2 [OS=Homo sapiens] | -3.38 | 0.768816825 | | 63174.5 | 1214618.219 | | 836050.9688 | | | 453405.1076 | 1214618.219 | | 868512.1121 | | |
| Q16537 | Serine/threonine-protein phosphatase 2A 56 kDa regulatory subunit epsilon | -3.25 | 0.768816825 | | 646681.8906 | 5941200.844 | | 4924081.156 | | | 4641253.547 | 5941200.844 | | 5115267.233 | | Met-loss+Acetyl [N-Term] |
| P23470 | Receptor-type tyrosine-protein phosphatase gamma [OS=Homo sapiens] | 1.24 | 0.768816825 | 1372192.609 | 29455.96289 | 261819.1563 | 2496578.438 | 3050577.203 | 641192.3828 | 9277552.228 | 211406.2482 | 261819.1563 | 18249618.58 | 3169021.207 | 5425886.767 | |
| Q96E17 | Ras-related protein Rab-3C [OS=Homo sapiens] | -3.22 | 0.769364266 | | 704257.4688 | 7194550.938 | | 6156980.25 | | | 5054475.04 | 7194550.938 | | 6396035.793 | | |
| P19086 | Guanine nucleotide-binding protein G(z) subunit alpha [OS=Homo sapiens] | -3.2 | 0.778780376 | | 1156928.605 | 10233538.31 | | 9439656 | | | 8303308.121 | 10233538.31 | | 9806167.18 | | |
| O15260 | Surfeit locus protein 4 [OS=Homo sapiens] | 1.4 | 0.778780376 | 742886.6563 | 243227.1094 | 546317.9375 | 1856514.188 | 1657449.625 | 365807.625 | 5022742.22 | 1745647.591 | 546317.9375 | 13570843.72 | 1721803.01 | 3095530.772 | |
| Q9UBC2 | Epidermal growth factor receptor substrate 15-like 1 [OS=Homo sapiens] | -3.57 | 0.780222102 | | 156721.668 | 1311795.406 | | 714144.9375 | | | 1124795.681 | 1311795.406 | | 741872.8657 | | |
| Q9Y281 | Cofilin-2 [OS=Homo sapiens] | -3.19 | 0.780222102 | | 756921.375 | 17095884.69 | | 9199939.719 | | | 5432445.33 | 17095884.69 | | 9557143.494 | | Met-loss+Acetyl [N-Term] |
| P32320 | Cytidine deaminase [OS=Homo sapiens] | 1.49 | 0.780222102 | | 41284.61719 | | | 799729.125 | | | 296300.8223 | | | 830780.0092 | | |
| O14964 | Hepatocyte growth factor-regulated tyrosine kinase substrate [OS=Homo sa | -3.42 | 0.784506598 | | 153734.4727 | 830864.625 | | 913831.8438 | | | 1103356.499 | 830864.625 | | 949312.9658 | | |
| P55060 | Exportin-2 [OS=Homo sapiens] | -3.18 | 0.784506598 | | 351502.375 | 3655410.531 | | 3258170.984 | | | 2522742.122 | 3655410.531 | | 3384675.18 | | |
| Q8NE86 | Calcium uniporter protein, mitochondrial [OS=Homo sapiens] | -3.38 | 0.784897336 | | 137876.4766 | 1065263.656 | | 893411.3438 | | | 989543.2286 | 1065263.656 | | 928099.6041 | | |
| O00505 | Importin subunit alpha-4 [OS=Homo sapiens] | -3.55 | 0.785678178 | | 83273.07031 | 1992992.813 | | 830141 | | | 597653.0944 | 1992992.813 | | 862372.6785 | | |
| Q9Y4C0 | Neurexin-3 [OS=Homo sapiens] | -3.17 | 0.786222977 | | 261753.9453 | 5949405.813 | | 3978941.664 | | | 1878615.197 | 5949405.813 | | 4133431.044 | | |
| O75915 | PRA1 family protein 3 [OS=Homo sapiens] | 1.26 | 0.786222977 | 95530.30469 | | 575895.8281 | 316867.0625 | 1044545.656 | | 645891.3895 | | 575895.8281 | 2316251.293 | 1085101.971 | | |
| P21926 | CD9 antigen [OS=Homo sapiens] | 2.02 | 0.786857631 | 4307045.793 | 1351017.805 | 1171089.875 | 15747060.17 | 18030795.41 | 1528270.5 | 29120432.53 | 9696291.592 | 1171089.875 | 115108677.4 | 18730872.62 | 12932503.42 | |
| P43004 | Excitatory amino acid transporter 2 [OS=Homo sapiens] | -3.16 | 0.786857631 | | 7418772.676 | 85602268.88 | | 56441345.09 | | | 53244733.61 | 85602268.88 | | 58632779.18 | | Met-loss+Acetyl [N-Term] |
| O14827 | Ras-specific guanine nucleotide-releasing factor 2 [OS=Homo sapiens] | -2.66 | 0.786857631 | | 33448.875 | 2775855 | | 577585.9219 | | | 240063.4872 | 2775855 | | 600011.707 | | |
| O95292 | Vesicle-associated membrane protein-associated protein B/C [OS=Homo sa | -3.16 | 0.786857631 | | 386477.2734 | 4058823.375 | | 2950079.563 | | | 2773757.921 | 4058823.375 | | 3064621.569 | | |
| Q8NFZ8 | Cell adhesion molecule 4 [OS=Homo sapiens] | -3.16 | 0.787094295 | | 344632.9922 | 4282522.375 | | 3089980.75 | | | 2473440.374 | 4282522.375 | | 3209954.665 | | |
| P07196 | Neurofilament light polypeptide [OS=Homo sapiens] | -3.16 | 0.787094295 | | 3595785.73 | 54774569.63 | | 36069681.58 | | | 25807052.15 | 54774569.63 | | 37470150.15 | | |
| P43307 | Translocon-associated protein subunit alpha [OS=Homo sapiens] | -3.5 | 0.787094295 | | 149850.3125 | 744964.3125 | | 810952.4375 | | | 1075479.775 | 744964.3125 | | 842439.0864 | | |
| Q9Y617 | Phosphoserine aminotransferase [OS=Homo sapiens] | -3.74 | 0.78741743 | | 171696.1563 | 1535758.5 | | 664001.625 | | | 1232267.991 | 1535758.5 | | 689782.6512 | | |
| Q9UHD9 | Ubiquilin-2 [OS=Homo sapiens] | -3.15 | 0.78741743 | | 462249.9102 | 6905752.906 | | 5609292.453 | | | 3317580.199 | 6905752.906 | | 5827083.058 | | |
| Q5TGZ0 | MICOS complex subunit MIC10 [OS=Homo sapiens] | -3.87 | 0.788805599 | | 221985.7188 | 1113812 | | 721111 | | | 1593197.551 | 1113812 | | 749109.3977 | | Met-loss+Acetyl [N-Term] |
| P32004 | Neural cell adhesion molecule L1 [OS=Homo sapiens] | -3.15 | 0.788805599 | | 348443.1309 | 5471573.938 | | 3755041.234 | | | 2500785.843 | 5471573.938 | | 3900837.288 | | |
| Q9H1E3 | Nuclear ubiquitous casein and cyclin-dependent kinase substrate 1 [OS=Ho | 1.16 | 0.788805599 | 4348961.75 | 360377.6875 | 1855796.313 | 819252.5469 | 4911649.5 | 1224458.969 | 29403831.14 | 2586440.481 | 1855796.313 | 5988614.77 | 5102352.896 | 10361594.89 | |
| Q99471 | Prefoldin subunit 5 [OS=Homo sapiens] | -3.55 | 0.790446789 | | 382362.9766 | 3111245.875 | | 2831276.563 | | | 2744229.5 | 3111245.875 | | 2941205.835 | | |
| P27816 | Microtubule-associated protein 4 [OS=Homo sapiens] | -3.45 | 0.794810284 | | 142072.4531 | 1317862.25 | | 827212.4375 | | | 1019657.847 | 1317862.25 | | 859330.4094 | | |
| Q6PUV4 | Complexin-2 [OS=Homo sapiens] | -3.13 | 0.796024495 | | 398712.2813 | 9090250.875 | | 5582615.75 | | | 2861568.905 | 9090250.875 | | 5799370.586 | | |
| P49720 | Proteasome subunit beta type-3 [OS=Homo sapiens] | -3.12 | 0.798838212 | | 413308.5234 | 3656901.969 | | 3075195.25 | | | 2966326.533 | 3656901.969 | | 3194595.092 | | |
| Q5SW79 | Centrosomal protein of 170 kDa [OS=Homo sapiens] | 1.08 | 0.807194254 | | 153162.2188 | 741938.0938 | 81676.82813 | | 109204.2031 | | 1099249.417 | 741938.0938 | 597045.5156 | | 924105.8635 | |
| Q13177 | Serine/threonine-protein kinase PAK 2 [OS=Homo sapiens] | -3.5 | 0.809126105 | | 50960.70703 | 1347810.313 | | 940739.4375 | | | 365746.3828 | 1347810.313 | | 977265.294 | | |
| Q14195 | Dihydropyrimidinase-related protein 3 [OS=Homo sapiens] | -3.09 | 0.809126105 | 233264.7031 | 3661040.074 | 62152236.91 | | 36259247.75 | 410436.3789 | 1577805.628 | 26275384.36 | 62152236.91 | | 37667076.56 | 3473187.418 | |
| P04792 | Heat shock protein beta-1 [OS=Homo sapiens] | 1.9 | 0.809314703 | 700675.0938 | 557247.9531 | 248239.9219 | 986859.2266 | 13366933.59 | 667196.3047 | 4737344.986 | 3999383.741 | 248239.9219 | 7213794.773 | 13885928.21 | 5645936.691 | |
| Q2M2I8 | AP2-associated protein kinase 1 [OS=Homo sapiens] | -3.09 | 0.809314703 | | 536078.4219 | 6529143.438 | | 4168040.281 | | | 3847449.439 | 6529143.438 | | 4329871.746 | | |
| Q9ULA0 | Aspartyl aminopeptidase [OS=Homo sapiens] | -2.79 | 0.809314703 | | 49616.33984 | 702720.375 | | 712295.1875 | | | 356097.8228 | 702720.375 | | 739951.2959 | | |
| Q9H4M9 | EH domain-containing protein 1 [OS=Homo sapiens] | -3.44 | 0.814440091 | | 45056.42969 | 1403274.25 | | 906271.1875 | | | 323371.2234 | 1403274.25 | | 941458.7539 | | |
| O75122 | CLIP-associating protein 2 [OS=Homo sapiens] | -3.44 | 0.814840899 | | 118060.8828 | 1808072.969 | | 697465.125 | | | 847326.1724 | 1808072.969 | | 724545.4302 | | |
| Q7Z5P4 | 17-beta-hydroxysteroid dehydrogenase 13 [OS=Homo sapiens] | -3.51 | 0.816179292 | | 93555.02344 | 1660596.938 | | 938615.4688 | | | 671446.9521 | 1660596.938 | | 975058.8585 | | |
| Q92823 | Neuronal cell adhesion molecule [OS=Homo sapiens] | -3.35 | 0.816179292 | | 57054.125 | 2424057.688 | | 750337.125 | | | 409479.0095 | 2424057.688 | | 779470.2782 | | |
| O15427 | Monocarboxylate transporter 4 [OS=Homo sapiens] | 1.11 | 0.816179292 | 804550.9531 | 226428.1758 | | 2052788.438 | 1825090.844 | 540186.082 | 5439661.631 | 1625081.186 | | 15005579.42 | 1895953.193 | 4571153.046 | Met-loss [N-Term] |
| Q9P032 | NADH dehydrogenase [ubiquinone] 1 alpha subcomplex assembly factor 4 [ | -1.89 | 0.816179292 | | 6934.620605 | 664401.5625 | | 350084.125 | | | 49769.96101 | 664401.5625 | | 363676.7544 | | |
| P68032 | Actin, alpha cardiac muscle 1 [OS=Homo sapiens] | 0.41 | 0.818139647 | 178969.2188 | 2095927.555 | 92644132.09 | 284694.4219 | 53855111.59 | 399227.3906 | 1210031.495 | 15042529.16 | 92644132.09 | 2081074.055 | 55946130.64 | 3378334.917 | |
| Q08495 | Dematin [OS=Homo sapiens] | -3.36 | 0.818139647 | | 134398.5195 | 1135623.438 | | 878345.5938 | | | 964581.8363 | 1135623.438 | | 912448.9 | | |
| Q14141 | Septin-6 [OS=Homo sapiens] | -3.04 | 0.819809013 | | 282272.5313 | 4292687 | | 2875099.5 | | | 2025877.647 | 4292687 | | 2986730.274 | | |
| Q5VZM2 | Ras-related GTP-binding protein B [OS=Homo sapiens] | -3.19 | 0.820101177 | | 69476.76563 | 1554537.188 | | 913251.5938 | | | 498636.6397 | 1554537.188 | | 948710.1866 | | |
| O94819 | Kelch repeat and BTB domain-containing protein 11 [OS=Homo sapiens] | -3.23 | 0.820101177 | | 62477.32422 | 1026413.75 | | 858452.0313 | | | 448401.4581 | 1026413.75 | | 891782.9351 | | |
| P78357 | Contactin-associated protein 1 [OS=Homo sapiens] | -4.05 | 0.821092507 | | 137611.625 | 2429064.125 | | 982480 | | | 987642.3818 | 2429064.125 | | 1020626.507 | | |
| Q9H299 | SH3 domain-binding glutamic acid-rich-like protein 3 [OS=Homo sapiens] | -3.03 | 0.821092507 | | 378553.2031 | 3582044.625 | | 3773530.5 | | | 2716886.652 | 3582044.625 | | 3920044.432 | | |
| Q93034 | Cullin-5 [OS=Homo sapiens] | -3.35 | 0.821092507 | | 115948.6875 | 1076895.813 | | 999187.3594 | | | 832166.8891 | 1076895.813 | | 1037982.559 | | |
| Q8WWM9 | Cytoglobin [OS=Homo sapiens] | -3.43 | 0.821092507 | | 134809.0313 | 862882.625 | | 798342.5625 | | | 967528.0901 | 862882.625 | | 829339.6109 | | |
| Q14978 | Nucleolar and coiled-body phosphoprotein 1 [OS=Homo sapiens] | 0.93 | 0.821172429 | 225652.9531 | | 348145.8125 | 59119.57031 | 477575.0625 | 78643.73438 | 1525665.598 | | 348145.8125 | 432155.3022 | 496117.7509 | 665497.6089 | |
| P67936 | Tropomyosin alpha-4 chain [OS=Homo sapiens] | -3.34 | 0.821566185 | | 54885.25 | 1255272.688 | | 1098780.031 | | | 393912.9346 | 1255272.688 | | 1141442.091 | | Met-loss+Acetyl [N-Term] |
| P00492 | Hypoxanthine-guanine phosphoribosyltransferase [OS=Homo sapiens] | -3 | 0.830241181 | | 2932566.879 | 34409370.91 | | 23457256.13 | | | 21047112.39 | 34409370.91 | | 24368025.18 | | |
| P21266 | Glutathione S-transferase Mu 3 [OS=Homo sapiens] | -2.38 | 0.830241181 | | 26412.93164 | 946564.8125 | | 677289.9375 | | | 189566.33 | 946564.8125 | | 703586.9057 | | |
| O14744 | Protein arginine N-methyltransferase 5 [OS=Homo sapiens] | -3.31 | 0.830241181 | | 73373.78125 | 983126.5625 | | 881592.3438 | | | 526605.6271 | 983126.5625 | | 915821.7108 | | |
| P31946 | 14-3-3 protein beta/alpha [OS=Homo sapiens] | -3 | 0.830241181 | | 3888536.719 | 30192633.25 | | 24626055.25 | 15667.3584 | | 27908133.97 | 30192633.25 | | 25582204.98 | 132580.0413 | Met-loss+Acetyl [N-Term] |
| P53992 | Protein transport protein Sec24C [OS=Homo sapiens] | -3 | 0.830241181 | | 580480.5 | 12442265 | | 6259138 | | | 4166124.363 | 12442265 | | 6502159.997 | | |
| Q9GZN7 | Protein rogdi homolog [OS=Homo sapiens] | -3.39 | 0.831571916 | | 161223.0664 | 994946.125 | | 953570.2813 | | | 1157102.34 | 994946.125 | | 990594.3178 | | |
| P63220 | Small ribosomal subunit protein eS21 [OS=Homo sapiens] | -3.69 | 0.831853169 | | 240006.5781 | 871196.8125 | | 1001852.5 | | | 1722533.75 | 871196.8125 | | 1040751.178 | | |
| P46934 | E3 ubiquitin-protein ligase NEDD4 [OS=Homo sapiens] | -2.64 | 0.831853169 | | 30610.75 | 1116681.188 | | 806941 | | | 219694.1867 | 1116681.188 | | 838271.8978 | | |

| | | | | | | | | | | | | | | | | |
|---|---|---|---|---|---|---|---|---|---|---|---|---|---|---|---|---|
| Q86UW7 | Calcium-dependent secretion activator 2 [OS=Homo sapiens] | -3.23 | 0.835866857 | | 103192.6406 | 1022861.188 | | 806686.125 | | | 740616.3932 | 1022861.188 | | 838007.1269 | | |
| P35813 | Protein phosphatase 1A [OS=Homo sapiens] | -3.47 | 0.835866857 | | 200995.3203 | 3026742.531 | | 2135508.25 | | | 1442548.89 | 3026742.531 | | 2218423.099 | | |
| Q9UIW2 | Plexin-A1 [OS=Homo sapiens] | -3.19 | 0.835866857 | | 79213.19922 | 1609348.875 | | 900437.6875 | | | 568515.2889 | 1609348.875 | | 935398.758 | | |
| Q9P0M2 | A-kinase anchor protein 7 isoform gamma [OS=Homo sapiens] | -3.63 | 0.835866857 | | 171537.7109 | 1113692.156 | | 756851.75 | | | 1231130.825 | 1113692.156 | | 786237.845 | | |
| Q8ND24 | RING finger protein 214 [OS=Homo sapiens] | -3.37 | 0.835866857 | | 128569.2578 | 885818.125 | | 801191.875 | | | 922745.0661 | 885818.125 | | 832299.553 | | |
| Q9NVH1 | DnaJ homolog subfamily C member 11 [OS=Homo sapiens] | -3.48 | 0.835866857 | | 131740.1719 | 1191218.5 | | 804151.0469 | | | 945502.8027 | 1191218.5 | | 835373.6199 | | |
| P37840 | Alpha-synuclein [OS=Homo sapiens] | -2.98 | 0.835866857 | | 2043911.609 | 15893654.31 | | 14972117.13 | | | 14669209.31 | 15893654.31 | | 15553435.8 | | |
| Q9UJ70 | N-acetyl-D-glucosamine kinase [OS=Homo sapiens] | -3.29 | 0.835866857 | | 138297.9219 | 966615.0938 | | 1000231.375 | | | 992567.9531 | 966615.0938 | | 1039067.11 | | |
| O94772 | Lymphocyte antigen 6H [OS=Homo sapiens] | -2.93 | 0.835866857 | | 1757924.516 | 21705093.72 | | 18550480.56 | | | 12616672.14 | 21705093.72 | | 19270735.47 | | |
| P09486 | SPARC [OS=Homo sapiens] | -3.34 | 0.835866857 | | 131750.0156 | 1707864.156 | | 885674.3125 | | | 945573.4516 | 1707864.156 | | 920062.1691 | | |
| Q9Y5S9 | RNA-binding protein 8A [OS=Homo sapiens] | -3.3 | 0.835866857 | | 132985.8086 | 1038798.125 | | 984820.9375 | | | 954442.7714 | 1038798.125 | | 1023058.335 | | |
| Q9H2H9 | Sodium-coupled neutral amino acid symporter 1 [OS=Homo sapiens] | 1.02 | 0.835866857 | 123195.6328 | 146853.3281 | 576007.8594 | 652131.9375 | 1000842.531 | 170754.1406 | 832939.8584 | 1053970.337 | 576007.8594 | 4766987.869 | 1039701.996 | 1444952.649 | |
| Q9HD45 | Transmembrane 9 superfamily member 3 [OS=Homo sapiens] | 0.88 | 0.836186999 | 103966.6602 | 93891.60938 | 449500.5781 | 246010.8828 | 817161.125 | 106383.4551 | 702930.5602 | 673862.6385 | 449500.5781 | 1798303.114 | 848888.8371 | 900236.1796 | |
| Q8WXF7 | Atlastin-1 [OS=Homo sapiens] | -2.6 | 0.836186999 | 21749.62305 | 628497.4297 | 3956350.688 | | 2966377.844 | 24958.74609 | 147051.7057 | 4510743.176 | 3956350.688 | | 3081552.66 | 211205.4568 | |
| Q14008 | Cytoskeleton-associated protein 5 [OS=Homo sapiens] | -3.26 | 0.836186999 | | 110341.1406 | 2742921.781 | | 791655.4688 | | | 791921.3724 | 2742921.781 | | 822392.8791 | | |
| Q3ZCW2 | Galectin-related protein [OS=Homo sapiens] | -3 | 0.837150446 | | 218246.2734 | 4789356.094 | | 2903519.313 | | | 1566359.45 | 4789356.094 | | 3016253.536 | | |
| P62837 | Ubiquitin-conjugating enzyme E2 D2 [OS=Homo sapiens] | 1.09 | 0.838988879 | 143293.2031 | 271554.2656 | 5327800.469 | 1438110.875 | 4565769.75 | 326133.4375 | 968821.8453 | 1948952.363 | 5327800.469 | 10512377.48 | 4743043.758 | 2759800.569 | |
| Q9Y320 | Thioredoxin-related transmembrane protein 2 [OS=Homo sapiens] | -3.59 | 0.839738201 | | 117818.5547 | 1319530.469 | | 1030754.188 | | | 845586.9769 | 1319530.469 | | 1070775.024 | | |
| Q13232 | Nucleoside diphosphate kinase 3 [OS=Homo sapiens] | -3.57 | 0.840172154 | | 196647.2188 | 1243550.313 | | 969030.2813 | | | 1411342.446 | 1243550.313 | | 1006654.579 | | |
| P53004 | Biliverdin reductase A [OS=Homo sapiens] | 0.72 | 0.840987431 | | 35803.55078 | 615252.1875 | 33249.57031 | 407091.6563 | 38498.70313 | | 256963.0594 | 615252.1875 | 243049.4341 | 422897.7029 | 325783.0402 | |
| P28289 | Tropomodulin-1 [OS=Homo sapiens] | -3.43 | 0.840987431 | | 135363.543 | 1163087.906 | | 802439.0313 | | | 971507.8358 | 1163087.906 | | 833595.1323 | | |
| Q13618 | Cullin-3 [OS=Homo sapiens] | -3.76 | 0.840987431 | | 559548.4844 | 3618840.625 | | 2314771.438 | | | 4015894.717 | 3618840.625 | | 2404646.493 | | |
| Q8N163 | Cell cycle and apoptosis regulator protein 2 [OS=Homo sapiens] | -2.85 | 0.840987431 | | 65649.01563 | 527739.1875 | | 880761.5938 | | | 471164.7737 | 527739.1875 | | 914958.7054 | | |
| Q12765 | Secernin-1 [OS=Homo sapiens] | 1.33 | 0.840987431 | | 560866.5391 | 14835818.13 | 34680.50391 | 8277014.984 | | | 4025354.431 | 14835818.13 | 253509.3467 | 8598384.59 | | |
| P38919 | Eukaryotic initiation factor 4A-III [OS=Homo sapiens] | -3.2 | 0.840987431 | | 89981.88672 | 1819073.5 | | 738517.875 | | | 645802.4525 | 1819073.5 | | 767192.1252 | | |
| P55145 | Mesencephalic astrocyte-derived neurotrophic factor [OS=Homo sapiens] | -3.25 | 0.842305472 | | 123338.3984 | 1207301.5 | | 937985.125 | | | 885203.0459 | 1207301.5 | | 974404.0405 | | |
| Q9Y3P9 | Rab GTPase-activating protein 1 [OS=Homo sapiens] | 0.95 | 0.842527634 | | 102269.2891 | 687886.4375 | 246760.0938 | | 156671.2031 | | 733989.4739 | 687886.4375 | 1803779.735 | | 1325780.266 | |
| Q5T1C6 | Acyl-coenzyme A thioesterase THEM4 [OS=Homo sapiens] | -3.41 | 0.842527634 | | 123338.3594 | 1167385.625 | | 754894.875 | | | 885202.7655 | 1167385.625 | | 784204.9909 | | |
| P30531 | Sodium- and chloride-dependent GABA transporter 1 [OS=Homo sapiens] | -2.87 | 0.842527634 | | 419861.6758 | 3507020.719 | | 3974202.25 | | | 3013358.686 | 3507020.719 | | 4128507.614 | | |
| P42356 | Phosphatidylinositol 4-kinase alpha [OS=Homo sapiens] | -3.53 | 0.848300563 | | 140049.9609 | 1572522.219 | | 1003675.938 | | | 1005142.385 | 1572522.219 | | 1042645.414 | | |
| P09936 | Ubiquitin carboxyl-terminal hydrolase isozyme L1 [OS=Homo sapiens] | -2.85 | 0.851181641 | | 3075631.73 | 57549759.84 | | 38209922.28 | 33434.26953 | | 22073892.72 | 57549759.84 | | 39693489.45 | 282926.8804 | |
| Q15365 | Poly(rC)-binding protein 1 [OS=Homo sapiens] | 0.82 | 0.851181641 | | 638100.25 | 10857078.5 | 77169.47852 | 7391247 | 146365.3574 | | 4579662.879 | 10857078.5 | 564097.4576 | 7678225.112 | 1238570.322 | |
| O75487 | Glypican-4 [OS=Homo sapiens] | 1.7 | 0.856313287 | 1436147.547 | 211591.5352 | 594369.625 | 5107817.406 | 4616963.344 | 338019.8359 | 9709958.924 | 1518598.212 | 594369.625 | 37337388.67 | 4796225.033 | 2860385.438 | |
| O00483 | Cytochrome c oxidase subunit NDUFA4 [OS=Homo sapiens] | -2.84 | 0.856313287 | | 1641597.707 | 16104188.38 | | 13151791.38 | | | 11781791.47 | 16104188.38 | | 13662432.71 | | |
| P19823 | Inter-alpha-trypsin inhibitor heavy chain H2 [OS=Homo sapiens] | 0.96 | 0.860895242 | 189948.25 | 42508.47656 | 185266.4844 | 341217.4375 | 537189.3125 | 122707.3594 | 1284261.989 | 305084.4944 | 185266.4844 | 2494248.927 | 558046.6285 | 1038372.032 | |
| P48066 | Sodium- and chloride-dependent GABA transporter 3 [OS=Homo sapiens] | -2.83 | 0.861198995 | | 4433895.297 | 43475450.88 | | 37582464.5 | 46642.35938 | | 31822187.34 | 43475450.88 | | 39041669.52 | 394696.1431 | |
| Q05329 | Glutamate decarboxylase 2 [OS=Homo sapiens] | -2.83 | 0.862149386 | | 1668382.484 | 30455203.88 | | 21331091.22 | | | 11974026.54 | 30455203.88 | | 22159308.2 | | |
| P19022 | Cadherin-2 [OS=Homo sapiens] | -3.12 | 0.868413789 | | 103965.8828 | 1670668 | | 951029.875 | | | 746165.9734 | 1670668 | | 987955.2758 | | |
| Q8N126 | Cell adhesion molecule 3 [OS=Homo sapiens] | -2.81 | 0.870142917 | | 316415.8359 | 8112521.5 | | 4664757 | | | 2270925.075 | 8112521.5 | | 4845874.362 | | |
| Q99250 | Sodium channel protein type 2 subunit alpha [OS=Homo sapiens] | -3.52 | 0.872425619 | | 312616.7656 | 2198124.906 | | 2049086.969 | | | 2243659.044 | 2198124.906 | | 2128646.36 | | |
| Q9GZT8 | NIF3-like protein 1 [OS=Homo sapiens] | -2.93 | 0.872425619 | | 79477.02344 | 1423617.375 | | 945340.3125 | | | 570408.7625 | 1423617.375 | | 982044.8061 | | |
| Q9UQM7 | Calcium/calmodulin-dependent protein kinase type II subunit alpha [OS=Ho | -2.8 | 0.872425619 | | 12447398.62 | 103631631.5 | | 86583283.53 | 67670.86719 | | 89335319.04 | 103631631.5 | | 89945031.18 | 572643.2076 | Met-loss+Acetyl [N-Term] |
| P62888 | Large ribosomal subunit protein eL30 [OS=Homo sapiens] | 0.88 | 0.872425619 | 129336.4238 | 364322.9219 | 5542874.75 | 413663.6328 | 4394606 | 79856.74219 | 874458.4535 | 2614755.536 | 5542874.75 | 3023819.884 | 4565234.276 | 675762.302 | |
| P60520 | Gamma-aminobutyric acid receptor-associated protein-like 2 [OS=Homo sa | -3.61 | 0.872425619 | | 387756.8828 | 4810460.844 | | 2489196.563 | | | 2782941.712 | 4810460.844 | | 2585843.979 | | |
| P26038 | Moesin [OS=Homo sapiens] | 0.17 | 0.872425619 | 1131101.262 | 1272330.422 | 8916550.906 | 1192767.402 | 12614295.53 | 560690.9883 | 7647505.867 | 9131550.102 | 8916550.906 | 8718953.038 | 13104067.65 | 4744669.299 | |
| P12035 | Keratin, type II cytoskeletal 3 [OS=Homo sapiens] | 1.33 | 0.87496601 | | 41090.54297 | | | 1511344.188 | | | 294907.9464 | | | 1570024.773 | | |
| Q9NQX3 | Gephyrin [OS=Homo sapiens] | -3.52 | 0.87496601 | | 293954.7188 | 3068784.875 | | 1957794.25 | | | 2109721.025 | 3068784.875 | | 2033809.041 | | Met-loss+Acetyl [N-Term] |
| P12081 | Histidine--tRNA ligase, cytoplasmic [OS=Homo sapiens] | -3.63 | 0.87496601 | | 362404.0469 | 5411708.063 | | 2158883.813 | | | 2600983.718 | 5411708.063 | | 2242706.258 | | |
| P48651 | Phosphatidylserine synthase 1 [OS=Homo sapiens] | 0.88 | 0.876911518 | 89053.74219 | 35631.02344 | | 215045.1094 | 317401.4063 | | 602102.6047 | 255724.826 | | 1571947.897 | 329725.0718 | | |
| P39023 | Large ribosomal subunit protein uL3 [OS=Homo sapiens] | 1.62 | 0.88098477 | 1257738.879 | 1751449.059 | 10865233.78 | 7876242.656 | 17651607.03 | 4654761.727 | 8503717.378 | 12570197.61 | 10865233.78 | 57574167.19 | 18336961.59 | 39389441.81 | |
| O60331 | Phosphatidylinositol 4-phosphate 5-kinase type-1 gamma [OS=Homo sapie | -3.06 | 0.883136109 | | 88195.43359 | 795345.8438 | | 871789.625 | | | 632981.0297 | 795345.8438 | | 905638.3843 | | |
| O94973 | AP-2 complex subunit alpha-2 [OS=Homo sapiens] | 0.74 | 0.888665578 | 75758 | 1381474.934 | 20128183.66 | 116721.1953 | 16034406.81 | | 512208.5609 | 9914883.236 | 20128183.66 | 853214.649 | 16656970.74 | | |
| Q16143 | Beta-synuclein [OS=Homo sapiens] | -2.74 | 0.888665578 | | 976719.7891 | 26430442.25 | | 15211563 | | | 7009944.536 | 26430442.25 | | 15802178.58 | | |
| P41208 | Centrin-2 [OS=Homo sapiens] | -3.55 | 0.888665578 | | 201646.7813 | 1151229 | | 1018514.375 | | | 1447224.443 | 1151229 | | 1058059.98 | | |
| Q8NC51 | SERPINE1 mRNA-binding protein 1 [OS=Homo sapiens] | 0.86 | 0.890679059 | | 90833.45313 | 1261585.625 | 132701.5781 | 1076752.844 | 286506.4453 | | 651914.1678 | 1261585.625 | 970028.8803 | 1118559.659 | 2424469.741 | Met-loss [N-Term] |
| P51970 | NADH dehydrogenase [ubiquinone] 1 alpha subcomplex subunit 8 [OS=Hon | -2.7 | 0.892233662 | | 50951.94141 | 1070829.25 | | 836220.3125 | | | 365683.4716 | 1070829.25 | | 868688.0309 | | Met-loss [N-Term] |
| Q13332 | Receptor-type tyrosine-protein phosphatase S [OS=Homo sapiens] | -3.39 | 0.893101279 | | 85178.57813 | 1254657.25 | | 867973.5938 | | | 611328.9758 | 1254657.25 | | 901674.1889 | | |
| P42262 | Glutamate receptor 2 [OS=Homo sapiens] | -3.54 | 0.895494494 | | 345507.4883 | 3193677.156 | | 2352802.766 | | | 2479716.657 | 3193677.156 | | 2444154.454 | | |
| Q9Y696 | Chloride intracellular channel protein 4 [OS=Homo sapiens] | 0.63 | 0.897301083 | 20059.20703 | 139361.7578 | 2804167.625 | 32344.66602 | 963651.1719 | 66047.71094 | 135622.6084 | 1000203.133 | 2804167.625 | 236434.7177 | 1001066.617 | 558907.7636 | |
| Q8NC96 | Adaptin ear-binding coat-associated protein 1 [OS=Homo sapiens] | -3.13 | 0.900927279 | | 88206.29688 | 3759590.594 | | 798325 | | | 633058.9958 | 3759590.594 | | 829321.3665 | | |
| O75822 | Eukaryotic translation initiation factor 3 subunit J [OS=Homo sapiens] | -2.99 | 0.900927279 | | 50752.19531 | 1191008.188 | | 897413.2188 | | | 364249.8885 | 1191008.188 | | 932256.859 | | |
| P48067 | Sodium- and chloride-dependent glycine transporter 1 [OS=Homo sapiens] | -3.77 | 0.900927279 | | 228373.3047 | 1801649.938 | | 1123258.922 | | | 1639041.43 | 1801649.938 | | 1166871.417 | | |
| Q9NP79 | Vacuolar protein sorting-associated protein VTA1 homolog [OS=Homo sapie | -3.43 | 0.900927279 | | 155798.2031 | 1555624.938 | | 928615.25 | | | 1118167.948 | 1555624.938 | | 964670.3637 | | |
| Q92915 | Fibroblast growth factor 14 [OS=Homo sapiens] | -3.37 | 0.900927279 | | 161644.1094 | 1037178.063 | | 1042748.938 | | | 1160124.177 | 1037178.063 | | 1083235.492 | | |
| P43686 | 26S proteasome regulatory subunit 6B [OS=Homo sapiens] | -3.44 | 0.900927279 | | 313606.6211 | 3444036.344 | | 2825938.375 | | | 2250763.264 | 3444036.344 | | 2935660.383 | | |
| Q15120 | [Pyruvate dehydrogenase (acetyl-transferring)] kinase isozyme 3, mitochond | -3.36 | 0.900927279 | | 84446.46094 | 1480697.75 | | 836820.4375 | | | 606074.5508 | 1480697.75 | | 869311.4568 | | |
| P26885 | Peptidyl-prolyl cis-trans isomerase FKBP2 [OS=Homo sapiens] | -3.41 | 0.900927279 | | 326924.2578 | 2620307.25 | | 1972587.75 | | | 2346344.305 | 2620307.25 | | 2049176.925 | | |
| O43236 | Septin-4 [OS=Homo sapiens] | -3.34 | 0.900927279 | | 122869.625 | 1904736.625 | | 942025.1719 | | | 881838.6461 | 1904736.625 | | 978600.9493 | | |
| P37235 | Hippocalcin-like protein 1 [OS=Homo sapiens] | -2.67 | 0.902101064 | | 3489886.395 | 18547604.09 | | 20725823.06 | 16604.16016 | | 25047009.73 | 18547604.09 | | 21530539.45 | 140507.4285 | |
| Q93009 | Ubiquitin carboxyl-terminal hydrolase 7 [OS=Homo sapiens] | 0.82 | 0.902101064 | | | 256007.2188 | | | 53519.04688 | | | 256007.2188 | | | 452887.9257 | |
| Q9UMY4 | Sorting nexin-12 [OS=Homo sapiens] | -3.47 | 0.902101064 | | 146825.9375 | 1408339.625 | | 1043046.609 | | | 1053773.754 | 1408339.625 | | 1083544.721 | | |
| Q53H12 | Acylglycerol kinase, mitochondrial [OS=Homo sapiens] | -3.36 | 0.902101064 | | 150866.0469 | 1669654 | | 977638.3438 | | | 1082769.729 | 1669654 | | 1015596.865 | | |
| P58549 | FXYD domain-containing ion transport regulator 7 [OS=Homo sapiens] | -3.76 | 0.90357887 | | 361023 | 1871934.25 | | 1356182.625 | | | 2591071.907 | 1871934.25 | | 1408838.791 | | |
| O14810 | Complexin-1 [OS=Homo sapiens] | -3.52 | 0.904052095 | | 394834.5 | 3885082.5 | | 2059921.5 | | | 2833737.964 | 3885082.5 | | 2139901.561 | | |
| Q05639 | Elongation factor 1-alpha 2 [OS=Homo sapiens] | 0.64 | 0.904052095 | 134382.4063 | 2739791.297 | 43497091.03 | 349446.4531 | 28277118.63 | 196923.2344 | 908574.9216 | 19663556.78 | 43497091.03 | 2554401.812 | 29375027.29 | 1666400.288 | |
| O00330 | Pyruvate dehydrogenase protein X component, mitochondrial [OS=Homo sa | -3.69 | 0.904541076 | | 243614.4844 | 1936853 | | 1016089.125 | | | 1748427.791 | 1936853 | | 1055540.565 | | |
| P55011 | Solute carrier family 12 member 2 [OS=Homo sapiens] | 0.76 | 0.905774455 | 53580.01563 | 125214 | 1339164.375 | 522160.9609 | 975769.9688 | 50586.20703 | 362260.6549 | 898664.2894 | 1339164.375 | 3816919.282 | 1013655.947 | 428069.7005 | |
| P01116 | GTPase KRas [OS=Homo sapiens] | -3.4 | 0.905779831 | | 377823.3984 | 2888504.25 | | 2368392.125 | | | 2711648.824 | 2888504.25 | | 2460349.098 | | |
| Q96N66 | Lysophospholipid acyltransferase 7 [OS=Homo sapiens] | -2.9 | 0.906070333 | | 77139.77344 | 1040087.875 | | 957191.5625 | | | 553634.2556 | 1040087.875 | | 994356.2016 | | |
| Q86X29 | Lipolysis-stimulated lipoprotein receptor [OS=Homo sapiens] | 1.11 | 0.908699731 | 1432386.578 | 53066.78906 | 232745.1719 | 3163258.188 | 3743156.891 | 474953.125 | 9684530.582 | 380861.7909 | 232745.1719 | 23122948.81 | 3888491.514 | 4019139.879 | |
| O75689 | Arf-GAP with dual PH domain-containing protein 1 [OS=Homo sapiens] | -3 | 0.908699731 | | 104907.9238 | 1091362.438 | | 1012235.125 | | | 752927.0275 | 1091362.438 | | 1051536.927 | | |
| Q3SXM5 | Inactive hydroxysteroid dehydrogenase-like protein 1 [OS=Homo sapiens] | -3.54 | 0.908699731 | | 391808.7344 | 1997181.813 | | 1991571.75 | | | 2812021.961 | 1997181.813 | | 2068898.012 | | |

| | | | | | | | | | | | | | | | | |
|---|---|---|---|---|---|---|---|---|---|---|---|---|---|---|---|---|
| Q12955 | Ankyrin-3 [OS=Homo sapiens] | -3.79 | 0.913715464 | | 114971.8047 | 2627866.281 | | 1620488.719 | | | 825155.7746 | 2627866.281 | | 1683407.032 | | |
| P12277 | Creatine kinase B-type [OS=Homo sapiens] | -2.63 | 0.913715464 | 43966.24609 | 19001684.67 | 292309445.8 | 355128.6172 | 165381306.5 | 806800.5918 | 297260.8522 | 136375608.6 | 292309445.8 | 2595937.59 | 171802525.5 | 6827293.604 | |
| Q8WVV9 | Heterogeneous nuclear ribonucleoprotein L-like [OS=Homo sapiens] | -3.4 | 0.91412212 | | 164639.2539 | 1359334.844 | | 1031212.813 | | | 1181620.411 | 1359334.844 | | 1071251.456 | | |
| P55735 | Protein SEC13 homolog [OS=Homo sapiens] | 0.6 | 0.91412212 | | 29616.86914 | 371668.375 | 70705.76563 | 310454.625 | | | 212561.0767 | 371668.375 | 516848.6738 | 322508.5696 | | |
| O95674 | Phosphatidate cytidylyltransferase 2 [OS=Homo sapiens] | -3.68 | 0.915529026 | | 473195.9063 | 2034458.25 | | 1995987 | | | 3396139.911 | 2034458.25 | | 2073484.692 | | |
| O00410 | Importin-5 [OS=Homo sapiens] | -3.66 | 0.916074998 | | 216999.1797 | 2913129.125 | | 2049355.125 | | | 1557409.025 | 2913129.125 | | 2128924.928 | | |
| Q99805 | Transmembrane 9 superfamily member 2 [OS=Homo sapiens] | 0.57 | 0.916606119 | 470241.8125 | 225289.0859 | 984585.4688 | 1536663.844 | 2614903.875 | 127186.7031 | 3179359.039 | 1616905.908 | 984585.4688 | 11232785.09 | 2716432.098 | 1076277.055 | |
| P49407 | Beta-arrestin-1 [OS=Homo sapiens] | -3.32 | 0.916606119 | | 138284.8164 | 3288518.438 | | 2212014.938 | | | 992473.8948 | 3288518.438 | | 2297900.292 | | |
| P30419 | Glycylpeptide N-tetradecanoyltransferase 1 [OS=Homo sapiens] | -3.51 | 0.917827872 | | 132341.0938 | 1151695.563 | | 1166585.594 | | | 949815.6354 | 1151695.563 | | 1211880.323 | | |
| O94967 | WD repeat-containing protein 47 [OS=Homo sapiens] | -3.04 | 0.917827872 | | 85277.03125 | 1796237.438 | | 967992.6875 | | | 612035.5766 | 1796237.438 | | 1005576.699 | | |
| P02768 | Albumin [OS=Homo sapiens] | 1.41 | 0.917827872 | 615159817.1 | 33901602.76 | 309792682.2 | 385988050.3 | 918014842.3 | 219226962.9 | 4159166354 | 243312726.6 | 309792682.2 | 2821515475 | 953658376.6 | 1855138503 | |
| Q9Y3T9 | Nucleolar complex protein 2 homolog [OS=Homo sapiens] | 0.89 | 0.917827872 | 4871839.203 | 446231.7227 | 658633.9063 | 2878604.145 | 6907525.922 | 1487576.484 | 32939065.8 | 3202617.231 | 658633.9063 | 21042169.92 | 7175722.716 | 12588143.24 | |
| Q15019 | Septin-2 [OS=Homo sapiens] | -3.53 | 0.917827872 | | 138599.9219 | 2069398.344 | | 1064290.344 | | | 994735.4154 | 2069398.344 | | 1105613.281 | | |
| P29218 | Inositol monophosphatase 1 [OS=Homo sapiens] | -3.55 | 0.917827872 | | 344158.6875 | 2471142.063 | | 1900015.375 | | | 2470036.277 | 2471142.063 | | 1973786.8 | | |
| Q8WUH6 | Transmembrane protein 263 [OS=Homo sapiens] | -3.53 | 0.917827872 | | 286208.9063 | 1774370.906 | | 1480152.75 | | | 2054129.118 | 1774370.906 | | 1537622.273 | | |
| Q9NX63 | MICOS complex subunit MIC19 [OS=Homo sapiens] | -3.67 | 0.917827872 | | 365043.5547 | 2213108.125 | | 1628699.594 | | | 2619927.537 | 2213108.125 | | 1691936.708 | | |
| Q12959 | Disks large homolog 1 [OS=Homo sapiens] | -3.37 | 0.917827872 | | 245621.875 | 3243578.469 | | 2603775.313 | | | 1762834.889 | 3243578.469 | | 2704871.45 | | |
| Q5T0D9 | Tumor protein p63-regulated gene 1-like protein [OS=Homo sapiens] | -3.65 | 0.917827872 | | 275060 | 1780336.625 | | 1213167.125 | | | 1974113.114 | 1780336.625 | | 1260270.464 | | |
| P23471 | Receptor-type tyrosine-protein phosphatase zeta [OS=Homo sapiens] | -3.72 | 0.917827872 | | 397471.5313 | 2355628 | | 1570637.375 | | | 2852664.009 | 2355628 | | 1631620.122 | | |
| Q96FJ2 | Dynein light chain 2, cytoplasmic [OS=Homo sapiens] | -2.53 | 0.917827872 | | 4715922.836 | 46124958.03 | | 35543405.19 | 33902.64453 | | 33846306.68 | 46124958.03 | | 36923440.15 | 286890.355 | |
| Q06323 | Proteasome activator complex subunit 1 [OS=Homo sapiens] | -3.67 | 0.917827872 | | 352885.125 | 2480057.375 | | 2194528.563 | | | 2532666.156 | 2480057.375 | | 2279734.978 | | |
| Q15369 | Elongin-C [OS=Homo sapiens] | -3.62 | 0.917827872 | | 180170.5938 | 1560612.609 | | 1055968.297 | | | 1293089.26 | 1560612.609 | | 1096968.116 | | |
| P58546 | Myotrophin [OS=Homo sapiens] | -3.68 | 0.917827872 | | 404050.1953 | 2492727.094 | | 1552519.375 | | | 2899879.26 | 2492727.094 | | 1612798.659 | | Met-loss+Acetyl [N-Term] |
| P02538 | Keratin, type II cytoskeletal 6A [OS=Homo sapiens] | 1.3 | 0.917827872 | 24137182.09 | 12652822.09 | 32423491.5 | 23311540.93 | 287509739 | 36310307.98 | 163194267.3 | 90809648.88 | 32423491.5 | 170403911.3 | 298672808.2 | 307264441.8 | Met-loss+Acetyl [N-Term] |
| O43719 | 17S U2 SnRNP complex component HTATSF1 [OS=Homo sapiens] | 0.33 | 0.917827872 | 3334881.578 | 121100.0313 | 564914.2813 | 2165239.453 | 4008956.063 | 1469546.922 | 22547518.33 | 869138.2236 | 564914.2813 | 15827579.69 | 4164610.804 | 12435573.8 | |
| P13929 | Beta-enolase [OS=Homo sapiens] | -2.44 | 0.917827872 | | 1200942.273 | 27469568.59 | | 7272071.969 | 63476.05078 | | 8619195.415 | 27469568.59 | | 7554422.901 | 537145.9068 | |
| Q9GZY8 | Mitochondrial fission factor [OS=Homo sapiens] | -3.59 | 0.917827872 | | 211175.6797 | 1744411.844 | | 1188118.984 | | | 1515613.606 | 1744411.844 | | 1234249.785 | | |
| P04259 | Keratin, type II cytoskeletal 6B [OS=Homo sapiens] | 1.04 | 0.917827872 | 897780.3281 | 1245701.555 | 3247787.188 | 1609727.406 | 18553535.81 | 2276116.172 | 6069996.17 | 8940433.996 | 3247787.188 | 11766868.91 | 19273909.34 | 19260909.75 | Met-loss+Acetyl [N-Term] |
| Q96GR2 | Long-chain-fatty-acid--CoA ligase ACSBG1 [OS=Homo sapiens] | -3.15 | 0.917827872 | | 167605.9492 | 3894902.25 | | 2559802.625 | | | 1202912.464 | 3894902.25 | | 2659191.446 | | |
| Q16134 | Electron transfer flavoprotein-ubiquinone oxidoreductase, mitochondrial [O | -3.6 | 0.917827872 | | 271690.7734 | 1454720.938 | | 1262399.438 | | | 1949932.083 | 1454720.938 | | 1311414.307 | | |
| Q99436 | Proteasome subunit beta type-7 [OS=Homo sapiens] | -3.44 | 0.917827872 | | 211963.25 | 1512205.75 | | 1249588.75 | | | 1521266.02 | 1512205.75 | | 1298106.222 | | |
| Q9H0R4 | Haloacid dehalogenase-like hydrolase domain-containing protein 2 [OS=Ho | -3.45 | 0.917827872 | | 432101.4688 | 4271035.25 | | 2492735.5 | | | 3101204.014 | 4271035.25 | | 2589520.322 | | |
| O00743 | Serine/threonine-protein phosphatase 6 catalytic subunit [OS=Homo sapier | -3.1 | 0.917827872 | | 252341.1797 | 1542496.313 | | 707724.75 | | | 1811059.521 | 1542496.313 | | 735203.4031 | | Met-loss [N-Term] |
| Q9UHD8 | Septin-9 [OS=Homo sapiens] | -3.43 | 0.917827872 | | 349122.7402 | 2566495.906 | | 1913958.766 | | | 2505663.418 | 2566495.906 | | 1988217.567 | | |
| Q92896 | Golgi apparatus protein 1 [OS=Homo sapiens] | -3.39 | 0.917827872 | | 94273.1875 | 1750158.125 | | 1146490.219 | | | 676601.2351 | 1750158.125 | | 1191004.71 | | |
| Q13427 | Peptidyl-prolyl cis-trans isomerase G [OS=Homo sapiens] | 0.85 | 0.917827872 | 396525.0781 | 37449.75781 | 214522.7969 | 331477.7969 | 853122.2813 | 207069.2031 | 2680951.71 | 268777.9321 | 214522.7969 | 2423053.597 | 886246.2482 | 1752257.324 | |
| Q12792 | Twinfilin-1 [OS=Homo sapiens] | -3.51 | 0.917827872 | | 203751.3672 | 1096465.656 | | 1087527.406 | | | 1462329.113 | 1096465.656 | | 1129752.563 | | Met-loss+Acetyl [N-Term] |
| P31942 | Heterogeneous nuclear ribonucleoprotein H3 [OS=Homo sapiens] | -3.49 | 0.917827872 | | 211473.7734 | 1217586.813 | | 1151155.188 | | | 1517753.033 | 1217586.813 | | 1195850.804 | | |
| P27338 | Amine oxidase [flavin-containing] B [OS=Homo sapiens] | -3.4 | 0.917827872 | | 351643.4609 | 2440693.188 | | 1936399.063 | | | 2523754.699 | 2440693.188 | | 2011583.148 | | |
| Q9BXS5 | AP-1 complex subunit mu-1 [OS=Homo sapiens] | -3.54 | 0.917827872 | | 318894.168 | 2574489.531 | | 1953715.5 | | | 2288712.132 | 2574489.531 | | 2029571.927 | | Met-loss+Acetyl [N-Term] |
| Q96SB3 | Neurabin-2 [OS=Homo sapiens] | -2.94 | 0.917827872 | | 138253.3867 | 1524485.188 | | 896215.25 | | | 992248.3231 | 1524485.188 | | 931012.377 | | |
| P29992 | Guanine nucleotide-binding protein subunit alpha-11 [OS=Homo sapiens] | -3.23 | 0.917827872 | | 380554.0859 | 3194474.188 | | 2201554.375 | | | 2731247.043 | 3194474.188 | | 2287033.58 | | |
| O75844 | CAAX prenyl protease 1 homolog [OS=Homo sapiens] | 0.99 | 0.917827872 | 280587.6094 | 127861.4102 | 882023.2656 | 562933.666 | 1201740.531 | 101950.6563 | 1897085.134 | 917664.8242 | 882023.2656 | 4114961.717 | 1248400.212 | 862725.0283 | |
| O75396 | Vesicle-trafficking protein SEC22b [OS=Homo sapiens] | -3.39 | 0.917827872 | | 277448.1895 | 5229091.938 | | 2523264.281 | | | 1991253.215 | 5229091.938 | | 2621234.437 | | Met-loss [N-Term] |
| P06703 | Protein S100-A6 [OS=Homo sapiens] | 0.65 | 0.917827872 | 124760.6016 | 181209.75 | 937042.5 | 187358.4531 | 1959033.125 | 86133.64844 | 843520.7923 | 1300547.313 | 937042.5 | 1369562.541 | 2035096.018 | 728878.6264 | |
| Q00535 | Cyclin-dependent kinase 5 [OS=Homo sapiens] | -3.43 | 0.917827872 | | 379546.6328 | 3975418.063 | | 1980396 | | | 2724016.524 | 3975418.063 | | 2057288.344 | | |
| P84085 | ADP-ribosylation factor 5 [OS=Homo sapiens] | -3.79 | 0.917827872 | | 384329.125 | 3721698.125 | | 1382308.125 | | | 2758340.601 | 3721698.125 | | 1435978.659 | | |
| P17174 | Aspartate aminotransferase, cytoplasmic [OS=Homo sapiens] | -2.6 | 0.917827872 | | 7875466.648 | 72470660.63 | | 46740913.38 | 46518.45557 | | 56522438.7 | 72470660.63 | | 48555711.21 | 393647.6465 | |
| P54687 | Branched-chain-amino-acid aminotransferase, cytosolic [OS=Homo sapien | -3.54 | 0.917827872 | | 194633.6094 | 1873459.406 | | 1224076.844 | | | 1396890.717 | 1873459.406 | | 1271603.771 | | |
| Q8WWI5 | Choline transporter-like protein 1 [OS=Homo sapiens] | 0.66 | 0.917827872 | 182934.6602 | 113601.4063 | 1172721.625 | 476208.1797 | 1389564.875 | 44352.33691 | 1236842.301 | 815320.3877 | 1172721.625 | 3481011.258 | 1443517.165 | 375317.5558 | |
| O95989 | Diphosphoinositol polyphosphate phosphohydrolase 1 [OS=Homo sapiens] | -3.56 | 0.917827872 | | 398032.5156 | 2149678.75 | | 792775.3125 | | | 2856690.209 | 2149678.75 | | 823556.2027 | | |
| Q8TBF2 | Prostamide/prostaglandin F synthase [OS=Homo sapiens] | -3.43 | 0.917827872 | | 212312.3594 | 1637926 | | 1263103.844 | | | 1523771.588 | 1637926 | | 1312146.063 | | |
| P53677 | AP-3 complex subunit mu-2 [OS=Homo sapiens] | -3.73 | 0.917827872 | | 274274.9844 | 2388685.219 | | 1137306.563 | | | 1968479.035 | 2388685.219 | | 1181464.482 | | |
| Q9Y2X7 | ARF GTPase-activating protein GIT1 [OS=Homo sapiens] | -3.35 | 0.917827872 | | 117960.7891 | 1893068.406 | | 1057928.594 | | | 846607.7969 | 1893068.406 | | 1099004.525 | | |
| P15531 | Nucleoside diphosphate kinase A [OS=Homo sapiens] | -3.63 | 0.917827872 | 31366.20117 | 1285909.805 | 10791367.56 | | 9216347.5 | 75706.96875 | 212070.4977 | 9229009.702 | 10791367.56 | | 9574188.336 | 640646.1631 | |
| Q9UN86 | Ras GTPase-activating protein-binding protein 2 [OS=Homo sapiens] | 0.55 | 0.917827872 | | 416053.0664 | 2423538.438 | 136734.0977 | 2632948.688 | | | 2986024.193 | 2423538.438 | 999506.0008 | 2735177.532 | | Met-loss [N-Term] |
| Q13835 | Plakophilin-1 [OS=Homo sapiens] | 0.72 | 0.917827872 | 352039.2188 | 160902.8945 | 217179.9063 | 187316.9355 | 2299182.25 | 406550.6328 | 2380177.692 | 1154804.458 | 217179.9063 | 1369259.053 | 2388452.028 | 3440305.527 | |
| Q96QR8 | Transcriptional activator protein Pur-beta [OS=Homo sapiens] | -3.61 | 0.917827872 | | 265709.0078 | 2622768.25 | | 1993059.281 | | | 1907000.788 | 2622768.25 | | 2070443.299 | | |
| P63215 | Guanine nucleotide-binding protein G(I)/G(S)/G(O) subunit gamma-3 [OS=H | -3.46 | 0.917827872 | | 208603.1484 | 1457966.938 | | 1236440.844 | | | 1497150.48 | 1457966.938 | | 1284447.825 | | |
| P42025 | Beta-centractin [OS=Homo sapiens] | -3.57 | 0.917827872 | | 311363.2227 | 2288372.938 | | 1329946.375 | | | 2234662.332 | 2288372.938 | | 1381583.873 | | |
| P51452 | Dual specificity protein phosphatase 3 [OS=Homo sapiens] | -3.32 | 0.917827872 | | 129027.9375 | 1592036.313 | | 1081113.813 | | | 926037.0227 | 1592036.313 | | 1123089.95 | | |
| P11441 | Ubiquitin-like protein 4A [OS=Homo sapiens] | -3.13 | 0.917827872 | | 62431.52344 | 518588.1875 | | 1005082.906 | | | 448072.7446 | 518588.1875 | | 1044107.011 | | |
| Q92769 | Histone deacetylase 2 [OS=Homo sapiens] | 0.71 | 0.917827872 | | 72408.00781 | 386721.3125 | 72206.34375 | 815790.375 | | | 519674.2448 | 386721.3125 | 527817.6776 | 847464.8653 | | |
| O60831 | PRA1 family protein 2 [OS=Homo sapiens] | -3.71 | 0.917827872 | | 268954.6797 | 1695745.563 | | 1201180.406 | | | 1930295.064 | 1695745.563 | | 1247818.34 | | |
| Q06124 | Tyrosine-protein phosphatase non-receptor type 11 [OS=Homo sapiens] | -3.5 | 0.917827872 | | 404079.2773 | 3256570.469 | | 2447580.484 | | | 2904609.51 | 3256570.469 | | 2542612.084 | | |
| C9JRZ8 | Aldo-keto reductase family 1 member B15 [OS=Homo sapiens] | -3.05 | 0.917827872 | | 89783.6875 | 1133924.375 | | 906286.6875 | | | 644379.9713 | 1133924.375 | | 941474.8557 | | |
| Q9UPR5 | Sodium/calcium exchanger 2 [OS=Homo sapiens] | -3.5 | 0.917827872 | | 265189.582 | 4009654.688 | | 2695866.953 | | | 1903272.855 | 4009654.688 | | 2800538.71 | | |
| Q96QK1 | Vacuolar protein sorting-associated protein 35 [OS=Homo sapiens] | -3.35 | 0.917827872 | 17391.92773 | 2181994.326 | 19159037.47 | 87242.6875 | 13792191.97 | | 117588.8259 | 15660232.73 | 19159037.47 | 637731.1232 | 14327697.98 | | |
| P25789 | Proteasome subunit alpha type-4 [OS=Homo sapiens] | 0.53 | 0.917827872 | | 610576.7734 | 6007030.938 | 63758.23633 | 4999634.438 | 27985.40234 | | 4382126.138 | 6007030.938 | 466063.2637 | 5193754.002 | 236817.5735 | |
| P49006 | MARCKS-related protein [OS=Homo sapiens] | -3.58 | 0.917827872 | | 251250.2344 | 1824236.594 | | 1475637.531 | | | 1803229.777 | 1824236.594 | | 1532931.743 | | |
| O95861 | 3'(2'),5'-bisphosphate nucleotidase 1 [OS=Homo sapiens] | -2.13 | 0.917827872 | 110503.4883 | 437568.4063 | 4006312.281 | | 3106900.125 | 96439.27734 | 747126.8078 | 3140440.373 | 4006312.281 | | 3227530.965 | 816086.7358 | |
| P31151 | Protein S100-A7 [OS=Homo sapiens] | 1.3 | 0.917827872 | 1183630.844 | 1104243.348 | 2876152.438 | 2890930.563 | 4018103.656 | 3857322 | 8002664.419 | 7925184.591 | 2876152.438 | 21132274.21 | 4174113.569 | 32641361.56 | |
| Q9ULP0 | Protein NDRG4 [OS=Homo sapiens] | -3.44 | 0.917827872 | | 352488.1094 | 4032220.125 | | 2577650.938 | | | 2529816.764 | 4032220.125 | | 2677732.75 | | |
| P02788 | Lactotransferrin [OS=Homo sapiens] | 1.28 | 0.917827872 | 646488.9199 | 272037.4219 | 148453.4844 | 691920.8359 | 64453529.44 | 824190.3555 | 4370986.025 | 1952419.988 | 148453.4844 | 5057838.823 | 66956050.62 | 6974448.953 | |
| Q9P2U8 | Vesicular glutamate transporter 2 [OS=Homo sapiens] | -3.57 | 0.917827872 | | 429464.1875 | 2671595 | | 2098447.75 | | | 3082276.174 | 2671595 | | 2179923.66 | | |
| P35241 | Radixin [OS=Homo sapiens] | -3.6 | 0.917827872 | | 101010.9297 | 2615820.531 | | 1656977.781 | | | 724958.1943 | 2615820.531 | | 1721312.846 | | |
| P17152 | Transmembrane protein 11, mitochondrial [OS=Homo sapiens] | -2.96 | 0.917827872 | | 76890.63281 | 3423933.875 | | 876960.3125 | | | 551846.1665 | 3423933.875 | | 911009.8328 | | |
| P25098 | Beta-adrenergic receptor kinase 1 [OS=Homo sapiens] | -3.12 | 0.917827872 | | 90660.03125 | 1417453.344 | | 1075763.75 | | | 650669.5143 | 1417453.344 | | 1117532.162 | | |
| Q16881 | Thioredoxin reductase 1, cytoplasmic [OS=Homo sapiens] | 0.7 | 0.917827872 | 83772.29688 | 172267.4063 | 2545906.625 | 126064.8828 | 2118068.563 | | 566394.1448 | 1236367.868 | 2545906.625 | 921515.6206 | 2200306.285 | | |
| O94919 | Endonuclease domain-containing 1 protein [OS=Homo sapiens] | 0.73 | 0.917827872 | 168110.7813 | 93043.03906 | | 408221.8984 | 951756.3438 | | 1136616.349 | 667772.4262 | | 2984041.612 | 988709.951 | | |
| Q9NS86 | LanC-like protein 2 [OS=Homo sapiens] | -3.48 | 0.917827872 | | 464530.2422 | 2193124.188 | | 2288456.563 | | | 3333946.204 | 2193124.188 | | 2377309.897 | | |
| P32418 | Sodium/calcium exchanger 1 [OS=Homo sapiens] | -3.31 | 0.917827872 | | 198060.1172 | 3364501.281 | | 2684691.469 | | | 1421482.857 | 3364501.281 | | 2788929.318 | | |

| | | | | | | | | | | | | | | | | |
|---|---|---|---|---|---|---|---|---|---|---|---|---|---|---|---|---|
| P84090 | Enhancer of rudimentary homolog [OS=Homo sapiens] | -3.55 | 0.917827872 | | 320805.0703 | 2713230.125 | | 2117557.188 | | | 2302426.73 | 2713230.125 | | 2199775.055 | | Met-loss+Acetyl [N-Term] |
| P19404 | NADH dehydrogenase [ubiquinone] flavoprotein 2, mitochondrial [OS=Hom | -3.43 | 0.917827872 | | 275717.6328 | 4391949.75 | | 2704715.344 | | | 1978832.962 | 4391949.75 | | 2809730.655 | | |
| O15212 | Prefoldin subunit 6 [OS=Homo sapiens] | -3.72 | 0.917827872 | | 277441.1016 | 2216364.438 | | 854509.125 | | | 1991202.345 | 2216364.438 | | 887686.9386 | | |
| Q8IXJ6 | NAD-dependent protein deacetylase sirtuin-2 [OS=Homo sapiens] | -3.28 | 0.917827872 | | 303366.7188 | 4581746 | | 2581315.563 | | | 2177271.206 | 4581746 | | 2681539.661 | | |
| P62316 | Small nuclear ribonucleoprotein Sm D2 [OS=Homo sapiens] | -3.58 | 0.917827872 | | 165950.9063 | 2315041.969 | | 1535779 | | | 1191034.175 | 2315041.969 | | 1595408.31 | | Met-loss+Acetyl [N-Term] |
| Q92544 | Transmembrane 9 superfamily member 4 [OS=Homo sapiens] | 0.71 | 0.917827872 | 64219.69922 | 48047.19531 | | 354333.625 | 314295.4375 | | 434196.7808 | 344836.0298 | | 2590126.315 | 326498.5084 | | |
| Q8TAC9 | Secretory carrier-associated membrane protein 5 [OS=Homo sapiens] | -3.67 | 0.917827872 | | 425192.125 | 2427973 | | 1805849.5 | | | 3051615.465 | 2427973 | | 1875964.77 | | |
| Q86Y82 | Syntaxin-12 [OS=Homo sapiens] | -3.46 | 0.917827872 | | 203012.25 | 4062323.875 | | 2797847.875 | | | 1457024.449 | 4062323.875 | | 2906479.22 | | |
| P25686 | DnaJ homolog subfamily B member 2 [OS=Homo sapiens] | -3.58 | 0.917827872 | | 229424.0078 | 1133947.094 | | 1094620.156 | | | 1646582.355 | 1133947.094 | | 1137120.701 | | |
| Q16775 | Hydroxyacylglutathione hydrolase, mitochondrial [OS=Homo sapiens] | -3.68 | 0.917827872 | | 442929.5391 | 2779054.688 | | 1654988 | | | 3178917.369 | 2779054.688 | | 1719245.808 | | |
| P02649 | Apolipoprotein E [OS=Homo sapiens] | -3.77 | 0.917827872 | | 407032.5625 | 2037924 | | 1505874.625 | | | 2921283.79 | 2037924 | | 1564342.845 | | |
| Q9Y2J2 | Band 4.1-like protein 3 [OS=Homo sapiens] | 0.84 | 0.917827872 | 60110.59766 | 2747077.881 | 38558701.28 | 239818.9297 | 22562465.81 | 82255.66406 | 406414.672 | 19715852.79 | 38558701.28 | 1753040.854 | 23438493.07 | 696062.416 | |
| Q9Y2I8 | WD repeat-containing protein 37 [OS=Homo sapiens] | -3.53 | 0.917827872 | | 390905.375 | 3146150.75 | | 2137242.938 | | | 2805538.526 | 3146150.75 | | 2220225.138 | | |
| Q9NY61 | Protein AATF [OS=Homo sapiens] | 1.34 | 0.917827872 | 849794.2031 | 83888.85352 | | 973875.9102 | 2599554.094 | 501694.1953 | 5745556.454 | 602072.5872 | | 7118888.653 | 2700486.335 | 4245427.688 | |
| P62745 | Rho-related GTP-binding protein RhoB [OS=Homo sapiens] | -2.5 | 0.917827872 | | 141697.1172 | 7267465.125 | | 3747269.094 | | | 1016964.071 | 7267465.125 | | 3892763.38 | | |
| Q9P0V9 | Septin-10 [OS=Homo sapiens] | -3.61 | 0.917827872 | | 364111.4688 | 3190155 | | 1677417.625 | | | 2613237.931 | 3190155 | | 1742546.303 | | |
| Q08188 | Protein-glutamine gamma-glutamyltransferase E [OS=Homo sapiens] | 1.15 | 0.917827872 | 324407.1094 | 95713.93945 | | | 3827298.063 | 175610.7773 | 2193353.819 | 686941.5511 | | | 3975899.614 | 1486050.394 | |
| Q8IZP0 | Abl interactor 1 [OS=Homo sapiens] | -3.41 | 0.917827872 | | 298695.6719 | 3093137.438 | | 1900905.563 | | | 2143746.975 | 3093137.438 | | 1974711.551 | | |
| P46977 | Dolichyl-diphosphooligosaccharide--protein glycosyltransferase subunit ST | 1.3 | 0.917827872 | 1432050.688 | 430421.125 | 1232980.063 | 3367690.594 | 4266306.469 | 637948.2461 | 9682259.586 | 3089144.141 | 1232980.063 | 24617319.42 | 4431953.291 | 5398434.291 | |
| Q07666 | KH domain-containing, RNA-binding, signal transduction-associated proteir | -3.33 | 0.917827872 | | 331026.5859 | 2440022.813 | | 2014313.188 | | | 2375786.826 | 2440022.813 | | 2092522.425 | | |
| P16144 | Integrin beta-4 [OS=Homo sapiens] | 0.54 | 0.917827872 | 1442230.273 | 252745.9922 | 224287.1406 | 2548444.742 | 2821600.516 | 360733.3828 | 9751084.939 | 1813964.872 | 224287.1406 | 18628753.59 | 2931154.098 | 3052591.61 | |
| Q8IXI2 | Mitochondrial Rho GTPase 1 [OS=Homo sapiens] | -3.56 | 0.917913632 | | 202129.5586 | 1998737.531 | | 1423059.672 | | | 1450689.349 | 1998737.531 | | 1478312.456 | | |
| P35606 | Coatomer subunit beta' [OS=Homo sapiens] | 0.76 | 0.917913632 | 140381.375 | 184794.2031 | 369789.1875 | 397417.9844 | 1464248.75 | 185878.9219 | 949134.6401 | 1326273.03 | 369789.1875 | 2905066.6 | 1521100.773 | 1572941.303 | |
| P06730 | Eukaryotic translation initiation factor 4E [OS=Homo sapiens] | -1.62 | 0.917913632 | | 129357.875 | 1580190.5 | | 989107.3594 | 16757.18945 | | 928404.9931 | 1580190.5 | | 1027511.185 | 141802.3903 | |
| Q9UNH7 | Sorting nexin-6 [OS=Homo sapiens] | -3.39 | 0.917913632 | | 354877.1328 | 2831143.281 | | 1996166.281 | | | 2546962.851 | 2831143.281 | | 2073670.934 | | |
| Q13867 | Bleomycin hydrolase [OS=Homo sapiens] | -3.45 | 0.917913632 | | 181920.0313 | 1240274.094 | | 1376729.781 | | | 1305645.021 | 1240274.094 | | 1430183.727 | | |
| Q13555 | Calcium/calmodulin-dependent protein kinase type II subunit gamma [OS=I | -3.06 | 0.917913632 | | 223945.1094 | 5674832.938 | | 2763057.125 | | | 1607260.151 | 5674832.938 | | 2870337.658 | | |
| O60268 | Uncharacterized protein KIAA0513 [OS=Homo sapiens] | -3.55 | 0.917913632 | | 323288.2266 | 2904677.031 | | 1630106.813 | | | 2320248.41 | 2904677.031 | | 1693398.565 | | |
| P32969 | Large ribosomal subunit protein uL6 [OS=Homo sapiens] | 0.77 | 0.917913632 | | 344026.5703 | 4129215.313 | 131749.4688 | 2963378.063 | | | 2469088.068 | 4129215.313 | 963069.1018 | 3078436.407 | | |
| Q9Y4E6 | WD repeat-containing protein 7 [OS=Homo sapiens] | -3.36 | 0.917913632 | | 225323.7227 | 5038938.813 | | 2605335.688 | | | 1617154.496 | 5038938.813 | | 2706492.409 | | |
| P20073 | Annexin A7 [OS=Homo sapiens] | -3.43 | 0.917913632 | | 384288.6953 | 2717337.813 | | 2163042.125 | | | 2758050.436 | 2717337.813 | | 2247026.024 | | |
| Q9UGV2 | Protein NDRG3 [OS=Homo sapiens] | -3.49 | 0.917913632 | | 248961.0781 | 2366085.031 | | 1453639.375 | | | 1786800.44 | 2366085.031 | | 1510079.47 | | |
| Q969T9 | WW domain-binding protein 2 [OS=Homo sapiens] | -3.58 | 0.917913632 | | 342766.6641 | 1845649.188 | | 1312827.813 | | | 2460045.686 | 1845649.188 | | 1363800.652 | | |
| P51530 | DNA replication ATP-dependent helicase/nuclease DNA2 [OS=Homo sapier | -3.36 | 0.917913632 | | 191893.6719 | 1555616.5 | | 1261181.375 | | | 1377226.111 | 1555616.5 | | 1310148.951 | | |
| Q9UI30 | Multifunctional methyltransferase subunit TRM112-like protein [OS=Homo s | -3.06 | 0.917913632 | | 108634.75 | 1131139.469 | | 1071890.813 | | | 779674.5604 | 1131139.469 | | 1113508.851 | | |
| Q9Y2K2 | Serine/threonine-protein kinase SIK3 [OS=Homo sapiens] | -3.53 | 0.917913632 | | 215428.6367 | 2454440.781 | | 1552642.242 | | | 1546137.195 | 2454440.781 | | 1612926.297 | | |
| P48643 | T-complex protein 1 subunit epsilon [OS=Homo sapiens] | 0.55 | 0.917913632 | 25484.44727 | 1398453.023 | 16554590.91 | 305967.1426 | 12169113.31 | 53402.01172 | 172303.2823 | 10036735.45 | 16554590.91 | 2236574.492 | 12641600.45 | 451897.5528 | |
| Q14157 | Ubiquitin-associated protein 2-like [OS=Homo sapiens] | 0.51 | 0.917913632 | | 28809.71484 | 452195.0938 | 33023.01953 | 284004.9375 | 27087.44141 | | 206768.1083 | 452195.0938 | 241393.3814 | 295031.9267 | 229218.8644 | |
| P02671 | Fibrinogen alpha chain [OS=Homo sapiens] | 0.3 | 0.917913632 | 1309514.43 | 1162256.297 | 1231942.875 | 1426132.719 | 3821417.813 | 1394321.766 | 8853777.838 | 8341545.108 | 1231942.875 | 10424818.94 | 3969791.053 | 11799004.82 | |
| O75935 | Dynactin subunit 3 [OS=Homo sapiens] | -3.56 | 0.917913632 | | 279539.9219 | 1547480.375 | | 1360066.625 | | | 2006265.635 | 1547480.375 | | 1412873.594 | | |
| O14910 | Protein lin-7 homolog A [OS=Homo sapiens] | -3.57 | 0.917913632 | | 310030 | 1065266.125 | | 1526180.563 | | | 2225093.757 | 1065266.125 | | 1585437.196 | | |
| Q96PK6 | RNA-binding protein 14 [OS=Homo sapiens] | -3.56 | 0.917913632 | | 48069.83594 | 1448666.969 | | 1376700.422 | | | 344998.5222 | 1448666.969 | | 1430153.227 | | |
| Q14894 | Ketimine reductase mu-crystallin [OS=Homo sapiens] | -3.56 | 0.917913632 | | 228811.9688 | 1423348.656 | | 1385577.063 | | | 1642189.734 | 1423348.656 | | 1439374.519 | | |
| P36776 | Lon protease homolog, mitochondrial [OS=Homo sapiens] | -3.6 | 0.917913632 | | 430323.6719 | 2232093.031 | | 1576931.469 | | | 3088444.717 | 2232093.031 | | 1638158.595 | | |
| Q15349 | Ribosomal protein S6 kinase alpha-2 [OS=Homo sapiens] | -3.42 | 0.917913632 | | 360674.3398 | 3377192.344 | | 1855241.797 | | | 2588569.563 | 3377192.344 | | 1927274.809 | | |
| P11279 | Lysosome-associated membrane glycoprotein 1 [OS=Homo sapiens] | 1.25 | 0.917913632 | 13519888.56 | 2300010.328 | 10474283.5 | 29679272.77 | 32629287.38 | 8459364.938 | 91409523.27 | 16507236.79 | 10474283.5 | 216951087.8 | 33896176.61 | 71584687.37 | |
| Q9UHG3 | Prenylcysteine oxidase 1 [OS=Homo sapiens] | -3.47 | 0.917913632 | | 201755.1563 | 1161461.906 | | 1157723.188 | | | 1448002.253 | 1161461.906 | | 1202673.818 | | |
| Q9UHW9 | Solute carrier family 12 member 6 [OS=Homo sapiens] | -3.62 | 0.917913632 | | 389178.5 | 2220718.5 | | 1782981.75 | | | 2793144.697 | 2220718.5 | | 1852209.14 | | |
| P00390 | Glutathione reductase, mitochondrial [OS=Homo sapiens] | -3.54 | 0.917913632 | | 236556.0391 | 2250371.938 | | 1221942.188 | | | 1697769.137 | 2250371.938 | | 1269386.233 | | |
| Q9UPV7 | PHD finger protein 24 [OS=Homo sapiens] | -3.52 | 0.917913632 | | 224707.375 | 2239649.313 | | 1180322.188 | | | 1612730.952 | 2239649.313 | | 1226150.264 | | |
| Q2NKQ1 | Small G protein signaling modulator 1 [OS=Homo sapiens] | -3.43 | 0.917913632 | | 333334.3438 | 2254264.594 | | 2317651.031 | | | 2392349.667 | 2254264.594 | | 2407637.892 | | |
| Q92932 | Receptor-type tyrosine-protein phosphatase N2 [OS=Homo sapiens] | -2.69 | 0.917913632 | | 53972.47656 | 1321436.875 | | 890818 | | | 387361.9347 | 1321436.875 | | 925405.5693 | | |
| P13584 | Cytochrome P450 4B1 [OS=Homo sapiens] | 0.56 | 0.917913632 | 106443.3203 | 137932.0781 | | 145561.8438 | | 145340.0313 | 719675.5447 | 989942.2826 | | 1064035.517 | | 1229893.825 | |
| Q6NVY1 | 3-hydroxyisobutyryl-CoA hydrolase, mitochondrial [OS=Homo sapiens] | -3.43 | 0.917913632 | | 201424.9063 | 1686155.125 | | 1195671.25 | | | 1445632.04 | 1686155.125 | | 1242095.281 | | |
| Q15700 | Disks large homolog 2 [OS=Homo sapiens] | -3.36 | 0.917913632 | | 378115.082 | 4021128.531 | | 2385084.406 | | | 2713742.245 | 4021128.531 | | 2477689.486 | | |
| O00592 | Podocalyxin [OS=Homo sapiens] | 1.23 | 0.918471863 | 8732860.719 | 1780652.391 | 6726219.188 | 18288302.45 | 22056339.88 | 6600781.129 | 59043876.83 | 12779790.72 | 6726219.188 | 133684782.1 | 22912715.91 | 55857012.55 | |
| Q9NP97 | Dynein light chain roadblock-type 1 [OS=Homo sapiens] | -3.55 | 0.918471863 | | 310306.2813 | 2129635.188 | | 1978271.344 | | | 2227076.635 | 2129635.188 | | 2055081.194 | | Met-loss+Acetyl [N-Term] |
| Q6PCE3 | Glucose 1,6-bisphosphate synthase [OS=Homo sapiens] | -3.47 | 0.919046768 | | 99498.65625 | 2528379.922 | | 1371034.719 | | | 714104.5667 | 2528379.922 | | 1424267.543 | | |
| Q9P2K5 | Myelin expression factor 2 [OS=Homo sapiens] | -3.5 | 0.919140997 | | 183378.0469 | 3476408.281 | | 2001463.078 | | | 1316109.238 | 3476408.281 | | 2079173.388 | | |
| P31025 | Lipocalin-1 [OS=Homo sapiens] | -2.35 | 0.919140997 | 172746.3594 | 122669.5859 | | | 5458813.125 | | 1167958.026 | 880402.9603 | | | 5670761.107 | | |
| Q68D91 | Acyl-coenzyme A thioesterase MBLAC2 [OS=Homo sapiens] | -3.46 | 0.919140997 | | 276175.4688 | 1880454.875 | | 1571529.969 | | | 1982118.863 | 1880454.875 | | 1632547.373 | | |
| A6NDG6 | Glycerol-3-phosphate phosphatase [OS=Homo sapiens] | -3.32 | 0.919140997 | | 207402.7813 | 3617623.063 | | 1988488.781 | | | 1488535.412 | 3617623.063 | | 2065695.341 | | |
| P53680 | AP-2 complex subunit sigma [OS=Homo sapiens] | 0.35 | 0.919140997 | 270502.6953 | 1553295.984 | 16372565.91 | 320548.8672 | 11090360.13 | 221800.5859 | 1828899.869 | 11148047.6 | 16372565.91 | 2343164.739 | 11520962.78 | 1876916.97 | |
| P10398 | Serine/threonine-protein kinase A-Raf [OS=Homo sapiens] | -3.34 | 0.919140997 | | 133020.0625 | 824999.75 | | 1227820.875 | | | 954688.6126 | 824999.75 | | 1275493.171 | | |
| Q8N3J6 | Cell adhesion molecule 2 [OS=Homo sapiens] | -3.52 | 0.919140997 | | 326291.1875 | 2486370.875 | | 1646614.563 | | | 2341800.742 | 2486370.875 | | 1710547.257 | | |
| P49841 | Glycogen synthase kinase-3 beta [OS=Homo sapiens] | -3.18 | 0.919140997 | | 304494.6641 | 3811261.188 | | 2652304.156 | | | 2185366.5 | 3811261.188 | | 2755284.511 | | |
| Q4G0N4 | NAD kinase 2, mitochondrial [OS=Homo sapiens] | -3.46 | 0.919140997 | | 150764.8711 | 1731987.859 | | 1073966.609 | | | 1082043.587 | 1731987.859 | | 1115665.244 | | |
| P08621 | U1 small nuclear ribonucleoprotein 70 kDa [OS=Homo sapiens] | 0.79 | 0.919140997 | | 186892.4688 | 439848.6875 | 263483.0938 | 1035433.781 | 89747.66406 | | 1341332.34 | 439848.6875 | 1926022.388 | 1075636.312 | 759461.085 | |
| Q13642 | Four and a half LIM domains protein 1 [OS=Homo sapiens] | -3.55 | 0.919140997 | | 215073.9531 | 1900377.219 | | 1458396.781 | | | 1543591.621 | 1900377.219 | | 1515021.591 | | |
| P35579 | Myosin-9 [OS=Homo sapiens] | 0.42 | 0.919140997 | 823659.4844 | 842823.7578 | 4222189.875 | 1292095.867 | 6368118.609 | 598111.8047 | 5568856.611 | 6048969.072 | 4222189.875 | 9445029.404 | 6615371.969 | 5061331.065 | |
| P42330 | Aldo-keto reductase family 1 member C3 [OS=Homo sapiens] | 0.65 | 0.919140997 | 202067.3125 | 233514.2188 | 184587.875 | 293337.3125 | 355661.0313 | 278701.9375 | 1366200.366 | 1675937.91 | 184587.875 | 2144252.305 | 369470.1938 | 2358426.574 | |
| Q6VY07 | Phosphofurin acidic cluster sorting protein 1 [OS=Homo sapiens] | -3.44 | 0.919140997 | | 120858.25 | 2513909.625 | | 1186537.594 | | | 867402.953 | 2513909.625 | | 1232606.994 | | |
| Q9UI12 | V-type proton ATPase subunit H [OS=Homo sapiens] | -3.66 | 0.919140997 | 11285.41504 | 1708214.457 | 19986634.88 | | 14780567.28 | 31039.71289 | 76301.9905 | 12259901.7 | 19986634.88 | | 15354448.7 | 262663.7058 | |
| Q6DN90 | IQ motif and SEC7 domain-containing protein 1 [OS=Homo sapiens] | -3.31 | 0.919140997 | | 245116.9375 | 2523794.844 | | 1013388.281 | | | 1759210.938 | 2523794.844 | | 1052734.856 | | |
| Q9HCM2 | Plexin-A4 [OS=Homo sapiens] | -3.34 | 0.919140997 | | 358732.9922 | 2774655.031 | | 1985472.281 | | | 2574636.458 | 2774655.031 | | 2062561.72 | | |
| P17612 | cAMP-dependent protein kinase catalytic subunit alpha [OS=Homo sapiens | -3.58 | 0.919140997 | 25240.03125 | 2147708.625 | 15789943.16 | | 13914227.28 | 24776.31836 | 170650.7575 | 15414163.31 | 15789943.16 | | 14454471.53 | 209661.7201 | |
| Q96DG6 | Carboxymethylenebutenolidase homolog [OS=Homo sapiens] | -3.44 | 0.919140997 | | 199831.2188 | 970800.8125 | | 1169886.969 | | | 1434194.101 | 970800.8125 | | 1215309.88 | | |
| P45985 | Dual specificity mitogen-activated protein kinase kinase 4 [OS=Homo sapiei | -3.48 | 0.919711727 | | 267529.1406 | 2426094.875 | | 1795077.625 | | | 1920063.931 | 2426094.875 | | 1864774.658 | | |
| Q9Y3D6 | Mitochondrial fission 1 protein [OS=Homo sapiens] | -3.38 | 0.919855762 | | 158293.7969 | 2094092.188 | | 1274041.938 | | | 1136078.893 | 2094092.188 | | 1323508.847 | | |
| P54136 | Arginine--tRNA ligase, cytoplasmic [OS=Homo sapiens] | -3.45 | 0.919855762 | | 131126.1406 | 1406414.719 | | 1202570.281 | | | 941095.8837 | 1406414.719 | | 1249262.179 | | |
| Q15435 | Protein phosphatase 1 regulatory subunit 7 [OS=Homo sapiens] | -3.24 | 0.922621141 | | 374045.6914 | 2821388.938 | | 2250685.5 | | | 2684536.118 | 2821388.938 | | 2338072.307 | | |
| O43520 | Phospholipid-transporting ATPase IC [OS=Homo sapiens] | -3.35 | 0.92293427 | | 179820.7813 | 943646.5 | | 1193622.625 | | | 1290578.646 | 943646.5 | | 1239967.114 | | |

| | | | | | | | | | | | | | | | | |
|---|---|---|---|---|---|---|---|---|---|---|---|---|---|---|---|---|
| P51553 | Isocitrate dehydrogenase [NAD] subunit gamma, mitochondrial [OS=Homo | -2.26 | 0.92293427 | 45604.56641 | 3888344.609 | 47298423.03 | 155057.7813 | 29091015.53 | 81824.76758 | 308337.7244 | 27906755.2 | 47298423.03 | 1133449.414 | 30220525.17 | 692416.0915 | |
| O95182 | NADH dehydrogenase [ubiquinone] 1 alpha subcomplex subunit 7 [OS=Hon | -3.11 | 0.92293427 | | 262159.5859 | 3585622.938 | | 1913891.219 | | | 1881526.491 | 3585622.938 | | 1988201.398 | | |
| P53779 | Mitogen-activated protein kinase 10 [OS=Homo sapiens] | -3.4 | 0.92293427 | | 250357.7656 | 2097974.188 | | 1382638.563 | | | 1796824.505 | 2097974.188 | | 1436321.927 | | |
| P05129 | Protein kinase C gamma type [OS=Homo sapiens] | -3.42 | 0.92293427 | | 88570.05469 | 4497718.156 | | 1626737.047 | | | 635669.6955 | 4497718.156 | | 1689897.962 | | |
| P18505 | Gamma-aminobutyric acid receptor subunit beta-1 [OS=Homo sapiens] | -3.35 | 0.92293427 | | 356561.2227 | 2836603.625 | | 2605600.5 | | | 2559049.609 | 2836603.625 | | 2706767.504 | | |
| P23515 | Oligodendrocyte-myelin glycoprotein [OS=Homo sapiens] | -3.46 | 0.92293427 | | 301937.9492 | 1672087.563 | | 1637628.156 | | | 2167016.888 | 1672087.563 | | 1701211.938 | | |
| Q16762 | Thiosulfate sulfurtransferase [OS=Homo sapiens] | -3.26 | 0.92293427 | | 159316.332 | 1778137.5 | | 1112873.875 | | | 1143417.655 | 1778137.5 | | 1156083.153 | | |
| Q8IYB5 | Stromal membrane-associated protein 1 [OS=Homo sapiens] | -3.34 | 0.92293427 | | 123962.7969 | 1402707.875 | | 1716557 | | | 889684.3705 | 1402707.875 | | 1783205.332 | | |
| P46379 | Large proline-rich protein BAG6 [OS=Homo sapiens] | -3.67 | 0.92293427 | | 201118.6875 | 2201637.25 | | 1848273.375 | | | 1443434.3 | 2201637.25 | | 1920035.826 | | |
| Q9UBW8 | COP9 signalosome complex subunit 7a [OS=Homo sapiens] | -3.37 | 0.92293427 | | 202190.0781 | 2319473.75 | | 1605822.75 | | | 1451123.699 | 2319473.75 | | 1668171.631 | | |
| P31431 | Syndecan-4 [OS=Homo sapiens] | 1.17 | 0.92293427 | 2578698.758 | 706761.082 | 825116.7031 | 5336251.07 | 5334851.938 | 1100752.383 | 17434879.22 | 5072443.541 | 825116.7031 | 39007204.92 | 5541986.909 | 9314767.217 | |
| P80723 | Brain acid soluble protein 1 [OS=Homo sapiens] | -3.39 | 0.92293427 | | 271053.5625 | 1474529.75 | | 1694955.5 | | | 1945358.803 | 1474529.75 | | 1760765.116 | | |
| Q13303 | Voltage-gated potassium channel subunit beta-2 [OS=Homo sapiens] | -3.47 | 0.92293427 | | 239932.6289 | 2482728.844 | | 1263680.25 | | | 1722003.014 | 2482728.844 | | 1312744.849 | | |
| O43809 | Cleavage and polyadenylation specificity factor subunit 5 [OS=Homo sapier | -3.32 | 0.92293427 | | 129299.918 | 1401834.813 | | 1185726.313 | | | 927989.034 | 1401834.813 | | 1231764.214 | | |
| Q9Y5B9 | FACT complex subunit SPT16 [OS=Homo sapiens] | 0.82 | 0.92293427 | 291452.9531 | 65922.80469 | 251233.7813 | 683262.0469 | 1180169.781 | 119493.1641 | 1970546.974 | 473129.7653 | 251233.7813 | 4994544.358 | 1225991.94 | 1011172.926 | Met-loss+Acetyl [N-Term] |
| Q14982 | Opioid-binding protein/cell adhesion molecule [OS=Homo sapiens] | -3.44 | 0.92293427 | | 116011.8203 | 4478026 | | 2028348.313 | | | 832619.9951 | 4478026 | | 2107102.489 | | |
| Q9UN36 | Protein NDRG2 [OS=Homo sapiens] | -2.27 | 0.92293427 | | 1072702.141 | 24250432.31 | | 14408401.19 | 47344.92578 | | 7698812.488 | 24250432.31 | | 14967832.6 | 400641.3881 | |
| Q9Y6I3 | Epsin-1 [OS=Homo sapiens] | -3.45 | 0.92293427 | | 245300.8672 | 2301598.281 | | 1786047.688 | | | 1760531.007 | 2301598.281 | | 1855394.118 | | |
| Q9BR01 | Sulfotransferase 4A1 [OS=Homo sapiens] | -3.46 | 0.92293427 | | 211420.0273 | 3469667.063 | | 1939560.656 | | | 1517367.296 | 3469667.063 | | 2014867.496 | | |
| P47736 | Rap1 GTPase-activating protein 1 [OS=Homo sapiens] | -3.39 | 0.92293427 | | 185153.4453 | 2280935.563 | | 1626290.625 | | | 1328851.321 | 2280935.563 | | 1689434.207 | | |
| Q8N608 | Inactive dipeptidyl peptidase 10 [OS=Homo sapiens] | -3.39 | 0.92293427 | | 352529.9863 | 2140577.063 | | 1729845.047 | | | 2530117.316 | 2140577.063 | | 1797009.311 | | |
| P13797 | Plastin-3 [OS=Homo sapiens] | -3.39 | 0.92293427 | | 155793.3672 | 1510132.094 | | 1651100.281 | | | 1118133.241 | 1510132.094 | | 1715207.142 | | |
| P08779 | Keratin, type I cytoskeletal 16 [OS=Homo sapiens] | 1.16 | 0.92293427 | 5038081.025 | 2735026.045 | 5667884.109 | 3491306.309 | 45399965.09 | 8928999.746 | 34063045.9 | 19629356.44 | 5667884.109 | 25520931.98 | 47162698.27 | 75558822.69 | |
| P53801 | Pituitary tumor-transforming gene 1 protein-interacting protein [OS=Homo s | 1.14 | 0.92293427 | 549756.25 | 86785.5625 | 214663.8438 | 1072227.5 | 1317774.5 | 138786.6094 | 3716965.306 | 622862.3465 | 214663.8438 | 7837824.207 | 1368939.403 | 1174437.575 | |
| Q5TF21 | Microtubule cross-linking factor 3 [OS=Homo sapiens] | -3.32 | 0.92293427 | | 332067.7891 | 2946125.719 | | 2167678.094 | | | 2383259.569 | 2946125.719 | | 2251841.993 | | |
| Q8WXF1 | Paraspeckle component 1 [OS=Homo sapiens] | -3.48 | 0.92293427 | | 193696.8672 | 2627901.813 | | 1817681.031 | | | 1390167.693 | 2627901.813 | | 1888255.681 | | |
| Q969X5 | Endoplasmic reticulum-Golgi intermediate compartment protein 1 [OS=Hor | -3.44 | 0.92293427 | | 333300.6133 | 2111157.688 | | 1799953.844 | | | 2392107.582 | 2111157.688 | | 1869840.205 | | |
| P49915 | GMP synthase [glutamine-hydrolyzing] [OS=Homo sapiens] | -3.2 | 0.92293427 | | 182115.9688 | 3808889.219 | | 2433700.531 | | | 1307051.269 | 3808889.219 | | 2528193.217 | | |
| Q14738 | Serine/threonine-protein phosphatase 2A 56 kDa regulatory subunit delta is | -3.3 | 0.92293427 | | 172448.2188 | 1106241.75 | | 1205032.938 | | | 1237665.564 | 1106241.75 | | 1251820.452 | | |
| O75116 | Rho-associated protein kinase 2 [OS=Homo sapiens] | -3.41 | 0.92293427 | | 137399.3242 | 2396987.844 | | 1578801.453 | | | 986118.6933 | 2396987.844 | | 1640101.185 | | |
| P30825 | High affinity cationic amino acid transporter 1 [OS=Homo sapiens] | 0.57 | 0.92293427 | 188976.6875 | 38371.77344 | | 258779.5625 | 349415.4375 | 53543.53906 | 1277693.143 | 275395.2634 | | 1891640.271 | 362982.1039 | 453095.183 | |
| P04271 | Protein S100-B [OS=Homo sapiens] | -2.25 | 0.92293427 | | 91647.58594 | 9823364.75 | | 3125279.844 | | | 657757.2212 | 9823364.75 | | 3246624.308 | | |
| Q8WUW1 | Protein BRICK1 [OS=Homo sapiens] | -3.72 | 0.92293427 | | 273730.1328 | 1755317.375 | | 1790146.688 | | | 1964568.621 | 1755317.375 | | 1859652.268 | | Met-loss+Acetyl [N-Term] |
| O14828 | Secretory carrier-associated membrane protein 3 [OS=Homo sapiens] | 1.13 | 0.92293427 | 1389434.844 | 107672.668 | 1361060.875 | 2574931.648 | 3700449.594 | 41417.33594 | 9394128.96 | 772769.6715 | 1361060.875 | 18822368.95 | 3844126.031 | 350481.0428 | |
| P46976 | Glycogenin-1 [OS=Homo sapiens] | 1.16 | 0.92293427 | 29187880.59 | 2418146.699 | 11314858.67 | 44490774.83 | 45880222.91 | 12973479.31 | 197342621.4 | 17355104.74 | 11314858.67 | 325220974 | 47661602.93 | 109783945.7 | Met-loss+Acetyl [N-Term] |
| Q15691 | Microtubule-associated protein RP/EB family member 1 [OS=Homo sapiens | -3.35 | 0.92293427 | | 333466.5 | 2525956.188 | | 1904374.063 | | | 2393298.156 | 2525956.188 | | 1978314.721 | | Met-loss+Acetyl [N-Term] |
| Q9NZ94 | Neuroligin-3 [OS=Homo sapiens] | -2.85 | 0.92293427 | | 46334.21094 | 1656723.563 | | 944068.8125 | | | 332541.8942 | 1656723.563 | | 980723.9379 | | |
| P52888 | Thimet oligopeptidase [OS=Homo sapiens] | 0 | 0.92293427 | | | 1504305.406 | | 907170.625 | | | | 1504305.406 | | 942393.1136 | | |
| Q01546 | Keratin, type II cytoskeletal 2 oral [OS=Homo sapiens] | 0.99 | 0.92293427 | 596065 | 462178.6875 | 477881.25 | 530365.5625 | 21878543.56 | 459709.6875 | 4030064.097 | 3317069.032 | 477881.25 | 3876893.704 | 22728016.34 | 3890147.133 | |
| P11166 | Solute carrier family 2, facilitated glucose transporter member 1 [OS=Homo | 1.12 | 0.92293427 | 3246929.5 | 1363482.301 | 4177735.922 | 6237189.664 | 9616554.594 | 1338397.102 | 21952864.21 | 9785749.62 | 4177735.922 | 45592932.59 | 9989934.171 | 11325760.13 | |
| Q14011 | Cold-inducible RNA-binding protein [OS=Homo sapiens] | -2.56 | 0.92293427 | | 22822.63086 | 2326888.25 | | 905156.9688 | | | 163798.6435 | 2326888.25 | | 940301.2736 | | |
| Q9Y3B9 | RRP15-like protein [OS=Homo sapiens] | 0.95 | 0.92293427 | 353424.1563 | 86536.10938 | 473821.4063 | 632170.8438 | 1132064 | 343745.1563 | 2389541.414 | 621072.0146 | 473821.4063 | 4621075.231 | 1176018.368 | 2908834.141 | |
| P00338 | L-lactate dehydrogenase A chain [OS=Homo sapiens] | 0.31 | 0.92293427 | 787085.2383 | 1319850.266 | 17116462.48 | 917377.4609 | 14233597.72 | 569972 | 5321573.922 | 9472601.315 | 17116462.48 | 6705893.358 | 14786242.08 | 4823206.911 | Met-loss+Acetyl [N-Term] |
| P20337 | Ras-related protein Rab-3B [OS=Homo sapiens] | -3.46 | 0.92293427 | | 131379.5469 | 2241951.375 | | 2000230.813 | | | 942914.5873 | 2241951.375 | | 2077893.278 | | |
| O15294 | UDP-N-acetylglucosamine--peptide N-acetylglucosaminyltransferase 110 k | 0.65 | 0.92293427 | 1331171.5 | 143452.2266 | 2622280.344 | 1937277.75 | 4309014.094 | 1414338.875 | 9000203.784 | 1029560.538 | 2622280.344 | 14161213.4 | 4476319.114 | 11968393.25 | |
| P30049 | ATP synthase subunit delta, mitochondrial [OS=Homo sapiens] | -3.67 | 0.92293427 | 87236.28906 | 1517108.5 | 10543171 | | 8824342 | 97943.23438 | 589814.5949 | 10888329.04 | 10543171 | | 9166962.536 | 828813.4942 | |
| P30837 | Aldehyde dehydrogenase X, mitochondrial [OS=Homo sapiens] | -3.37 | 0.92293427 | | 153134.6719 | 1368776.875 | | 1150827.156 | | | 1099051.712 | 1368776.875 | | 1195510.036 | | |
| Q14CZ8 | Hepatic and glial cell adhesion molecule [OS=Homo sapiens] | -3.37 | 0.92293427 | | 126163.8359 | 3196481.5 | | 1334157.813 | | | 905481.2878 | 3196481.5 | | 1385958.827 | | |
| P51513 | RNA-binding protein Nova-1 [OS=Homo sapiens] | -3.16 | 0.92293427 | | 226374.2461 | 3520977.938 | | 2298657.344 | | | 1624694.132 | 3520977.938 | | 2387906.742 | | |
| P46459 | Vesicle-fusing ATPase [OS=Homo sapiens] | -2.29 | 0.92293427 | | 19173790.39 | 172839179.7 | | 121487541.9 | 22831.80469 | | 137610816 | 172839179.7 | | 126204508.5 | 193206.8911 | |
| Q96L92 | Sorting nexin-27 [OS=Homo sapiens] | -3.49 | 0.92293427 | | 314578.4492 | 2246756.094 | | 1876960.563 | | | 2257738.101 | 2246756.094 | | 1949836.844 | | Met-loss+Acetyl [N-Term] |
| Q96JJ7 | Protein disulfide-isomerase TMX3 [OS=Homo sapiens] | -3.47 | 0.92293427 | | 249482.3906 | 1385263.625 | | 1412678.5 | | | 1790541.915 | 1385263.625 | | 1467528.217 | | |
| Q96P47 | Arf-GAP with GTPase, ANK repeat and PH domain-containing protein 3 [OS= | 0.16 | 0.925791769 | 1375801.125 | 226043.6484 | 10492899.03 | 1421746.75 | 10150353.73 | 1364621.125 | 9301949.818 | 1622321.423 | 10492899.03 | 10392758.15 | 10544459 | 11547672.59 | |
| Q14103 | Heterogeneous nuclear ribonucleoprotein D0 [OS=Homo sapiens] | 0.12 | 0.925791769 | 385679.7031 | 1741467.096 | 34300462.28 | 258903.418 | 18282963.38 | 186623.9023 | 2607624.881 | 12498556.79 | 34300462.28 | 1892545.637 | 18992831.45 | 1579245.464 | |
| Q9UQ03 | Coronin-2B [OS=Homo sapiens] | -2.78 | 0.928047949 | | 45328.63281 | 3945047.219 | | 1008188.281 | | | 325324.8326 | 3945047.219 | | 1047332.957 | | |
| O00305 | Voltage-dependent L-type calcium channel subunit beta-4 [OS=Homo sapie | -3.39 | 0.928047949 | | 260078.5352 | 1575029.391 | | 1392378.078 | | | 1866590.732 | 1575029.391 | | 1446439.596 | | |
| P35659 | Protein DEK [OS=Homo sapiens] | 0.6 | 0.928047949 | 344231.7969 | 81979.37109 | 291298.8125 | 332221.7031 | 1370430.922 | 323321.8281 | 2327390.814 | 588368.1798 | 291298.8125 | 2428491.441 | 1423640.303 | 2736008.218 | |
| P98164 | Low-density lipoprotein receptor-related protein 2 [OS=Homo sapiens] | 0.42 | 0.928047949 | 120511.2656 | 119180.1406 | 99946.45313 | 149598.2813 | | 113260.5859 | 814790.5427 | 855359.1163 | 99946.45313 | 1093541.277 | | 958431.7139 | |
| P00738 | Haptoglobin [OS=Homo sapiens] | 0.17 | 0.928400742 | 1529456.469 | 1337100.594 | 1142802.25 | 1783786.43 | 12315304.03 | 1734357.641 | 10340831.29 | 9596407.39 | 1142802.25 | 13039214.59 | 12793467.28 | 14676450.35 | |
| P21281 | V-type proton ATPase subunit B, brain isoform [OS=Homo sapiens] | -2.22 | 0.928400742 | 148967.8301 | 10136074.35 | 115998205.6 | 157496.1465 | 76068756.84 | 240474.9121 | 1007188.652 | 72746881.79 | 115998205.6 | 1151273.502 | 79022259.58 | 2034942.52 | |
| Q9P0K1 | Disintegrin and metalloproteinase domain-containing protein 22 [OS=Homo | -3.35 | 0.928400742 | | 240838.1484 | 2662099.188 | | 2008500.813 | | | 1728501.953 | 2662099.188 | | 2086484.375 | | |
| P62328 | Thymosin beta-4 [OS=Homo sapiens] | -2.22 | 0.928400742 | 117109.875 | 4851260.273 | 37461782.41 | | 35958833.53 | 126683.3438 | 791793.3491 | 34817627.15 | 37461782.41 | | 37354998.23 | 1072017.536 | Met-loss+Acetyl [N-Term] |
| P20042 | Eukaryotic translation initiation factor 2 subunit 2 [OS=Homo sapiens] | -1 | 0.928400742 | | 61815.54688 | 334000.6875 | | 378697.5938 | 11087.84473 | | 443651.864 | 334000.6875 | | 393401.1912 | 93827.36225 | |
| Q9Y262 | Eukaryotic translation initiation factor 3 subunit L [OS=Homo sapiens] | -3.3 | 0.930021872 | | 193181.75 | 1767854.938 | | 1283131.219 | | | 1386470.683 | 1767854.938 | | 1332951.036 | | |
| Q96AG4 | Leucine-rich repeat-containing protein 59 [OS=Homo sapiens] | -2.81 | 0.930159365 | | 88232.66016 | 1487689.938 | | 1049625.063 | | | 633248.2058 | 1487689.938 | | 1090378.594 | | |
| P18433 | Receptor-type tyrosine-protein phosphatase alpha [OS=Homo sapiens] | 1.06 | 0.930159365 | 2116374.75 | 275431.5859 | 2187382.281 | 4388460.469 | 5450325.219 | 926522.9531 | 14309053.37 | 1976779.996 | 2187382.281 | 32078996.01 | 5661943.643 | 7840406.039 | |
| P29144 | Tripeptidyl-peptidase 2 [OS=Homo sapiens] | -3.3 | 0.930352742 | | 251280.1484 | 2919439.5 | | 1278424.609 | | | 1803444.471 | 2919439.5 | | 1328061.684 | | |
| P63208 | S-phase kinase-associated protein 1 [OS=Homo sapiens] | 0.61 | 0.932186321 | | 425846.3906 | 5422865.875 | 33802.88672 | 4450500 | 29088.91016 | | 3056311.147 | 5422865.875 | 247094.095 | 4623298.458 | 246155.6576 | |
| P11387 | DNA topoisomerase 1 [OS=Homo sapiens] | 1.06 | 0.932207332 | 7833154.859 | 710730.8691 | 2854891.203 | 16691156.53 | 7810221.656 | 4093080.477 | 52960861.92 | 5100934.811 | 2854891.203 | 122009881.9 | 8113467.193 | 34636392.74 | Met-loss+Acetyl [N-Term] |
| P02786 | Transferrin receptor protein 1 [OS=Homo sapiens] | 1.04 | 0.934193973 | 10704536.77 | 2949003.402 | 3535003.875 | 20820671.89 | 21911007.91 | 4832077.016 | 72374605.61 | 21165077.76 | 3535003.875 | 152196027.6 | 22761741.17 | 40889916.1 | |
| O14775 | Guanine nucleotide-binding protein subunit beta-5 [OS=Homo sapiens] | -3.35 | 0.934193973 | | 130027.9609 | 1786514.938 | | 2263630.344 | | | 933214.2182 | 1786514.938 | | 2351519.757 | | |
| Q07954 | Prolow-density lipoprotein receptor-related protein 1 [OS=Homo sapiens] | 0.92 | 0.934193973 | | 120622.2305 | 4005693.531 | 51129.64844 | 2872942.766 | | | 865709.0343 | 4005693.531 | 373750.1568 | 2984489.801 | | |
| P55290 | Cadherin-13 [OS=Homo sapiens] | -3.29 | 0.934193973 | | 299330.1016 | 3491280.25 | | 2116996.5 | | | 2148300.294 | 3491280.25 | | 2199192.597 | | |
| P19367 | Hexokinase-1 [OS=Homo sapiens] | -2.19 | 0.934193973 | 83803.35156 | 14118513.64 | 125650634.9 | 148169.1484 | 93783426.97 | 237294.0723 | 566604.1091 | 101328957 | 125650634.9 | 1083094.528 | 97424732.8 | 2008025.674 | |
| P30520 | Adenylosuccinate synthetase isozyme 2 [OS=Homo sapiens] | -3.36 | 0.934193973 | | 152640.3125 | 2019852.469 | | 1480323.5 | | | 1095503.681 | 2019852.469 | | 1537799.653 | | |
| Q8N1N4 | Keratin, type II cytoskeletal 78 [OS=Homo sapiens] | 0.89 | 0.934193973 | 931552.6016 | 330998.1523 | 578376.0469 | 710148.1563 | 4277563 | 937223.1641 | 6298334.399 | 2375582.757 | 578376.0469 | 5191077.835 | 4443646.876 | 7930953.173 | |
| Q9BPW8 | Protein NipSnap homolog 1 [OS=Homo sapiens] | -3.63 | 0.934193973 | | 227334.6406 | 2275970.438 | | 1772762.844 | | | 1631586.909 | 2275970.438 | | 1841593.466 | | |
| P00387 | NADH-cytochrome b5 reductase 3 [OS=Homo sapiens] | -3.31 | 0.934716167 | | 288564.0625 | 2439590.75 | | 1758694.938 | | | 2071032.138 | 2439590.75 | | 1826979.349 | | |
| O43707 | Alpha-actinin-4 [OS=Homo sapiens] | 0.12 | 0.934716167 | 621152.1797 | 1178255.551 | 5312297.188 | 766920.875 | 6958998.125 | 403496.6289 | 4199681.407 | 8456372.189 | 5312297.188 | 5606078 | 7229193.417 | 3414461.99 | |
| P31939 | Bifunctional purine biosynthesis protein ATIC [OS=Homo sapiens] | 0.18 | 0.934716167 | | 476197.1016 | 4292542.719 | 29160.11328 | 2697632.203 | | | 3417679.572 | 4292542.719 | 213156.1089 | 2802372.499 | | |
| Q9UKA9 | Polypyrimidine tract-binding protein 2 [OS=Homo sapiens] | 0.5 | 0.934716167 | 79704.51563 | 542079.1719 | 3810864.094 | 176697.9063 | 3835060.531 | 101170.8438 | 538891.4074 | 3890516.985 | 3810864.094 | 1291635.522 | 3983963.474 | 856126.1128 | |
| O14556 | Glyceraldehyde-3-phosphate dehydrogenase, testis-specific [OS=Homo sa | 0 | 0.934716167 | | | 1498561 | | 854844.8125 | | | | 1498561 | | 888035.6598 | | |

| | | | | | | | | | | | | | | | | |
|---|---|---|---|---|---|---|---|---|---|---|---|---|---|---|---|---|
| P20962 | Parathymosin [OS=Homo sapiens] | 0.14 | 0.934716167 | 327021.0313 | 436929 | 3320800 | 1819358.094 | 3318311.5 | 269593.8438 | 2211026.846 | 3135851.337 | 3320800 | 13299238.18 | 3447150.757 | 2281352.225 | |
| P02647 | Apolipoprotein A-I [OS=Homo sapiens] | 0.52 | 0.934716167 | 517809.0703 | 876768.6367 | 556427.3125 | 1207170.648 | 5438522.063 | 1231382.781 | 3500966.746 | 6292592.393 | 556427.3125 | 8824238.634 | 5649682.208 | 10420185.45 | |
| Q5D862 | Filaggrin-2 [OS=Homo sapiens] | 0.88 | 0.934716167 | 733422.375 | 111592.5469 | 511268.6875 | 258847.1719 | 1422806.375 | 394747.625 | 4958753.125 | 800902.7492 | 511268.6875 | 1892134.486 | 1478049.325 | 3340426.324 | |
| Q92900 | Regulator of nonsense transcripts 1 [OS=Homo sapiens] | 0.52 | 0.934716167 | | 62078.73633 | 396653.4688 | 47273.48047 | 624312.0469 | | | 445540.7819 | 396653.4688 | 345562.1401 | 648552.0557 | | |
| Q9NYB9 | Abl interactor 2 [OS=Homo sapiens] | -3.16 | 0.934716167 | | 98212.625 | 1701421.875 | | 1341503.813 | | | 704874.6854 | 1701421.875 | | 1393590.048 | | |
| Q9Y265 | RuvB-like 1 [OS=Homo sapiens] | 0.49 | 0.934716167 | 162410.7578 | 416206.1836 | 4008942.594 | 374529.3242 | 3484593.719 | 33870.90234 | 1098077.834 | 2987123.119 | 4008942.594 | 2737753.885 | 3619889.174 | 286621.7468 | |
| O43488 | Aflatoxin B1 aldehyde reductase member 2 [OS=Homo sapiens] | -3.25 | 0.934716167 | | 241748.6289 | 1671796.188 | | 1328524.813 | | | 1735036.496 | 1671796.188 | | 1380107.116 | | |
| P61247 | Small ribosomal subunit protein eS1 [OS=Homo sapiens] | -0.2 | 0.934716167 | 258648.0078 | 1438654.891 | 11683211.63 | 923417.7695 | 9405112.5 | 92808.99512 | 1748748.962 | 10325265.35 | 11683211.63 | 6750047.119 | 9770282.468 | 785366.6262 | |
| P16070 | CD44 antigen [OS=Homo sapiens] | 1.04 | 0.934716167 | 204439558.8 | 28030106.61 | 136181848.9 | 306469362.7 | 389759249.5 | 116911095.5 | 1382239396 | 201172838.8 | 136181848.9 | 2240245647 | 404892335.1 | 989322990.8 | |
| Q01469 | Fatty acid-binding protein 5 [OS=Homo sapiens] | 0.2 | 0.934716167 | 774549 | 1033228.645 | 4996010.313 | 767338.0156 | 7486350.719 | 1186516.844 | 5236814.972 | 7415510.131 | 4996010.313 | 5609127.236 | 7777021.399 | 10040521.71 | Met-loss+Acetyl [N-Term] |
| Q9BQA1 | Methylosome protein WDR77 [OS=Homo sapiens] | 0.55 | 0.934716167 | 192084.5898 | | 356869.4375 | 245660.7813 | 377936.5625 | 47330.80078 | 1298706.029 | | 356869.4375 | 1795743.924 | 392610.6116 | 400521.8597 | |
| P31937 | 3-hydroxyisobutyrate dehydrogenase, mitochondrial [OS=Homo sapiens] | -3.15 | 0.934716167 | | 277133.0703 | 2283057.875 | | 2038483.375 | | | 1988991.596 | 2283057.875 | | 2117631.063 | | |
| P61619 | Protein transport protein Sec61 subunit alpha isoform 1 [OS=Homo sapiens | 0.44 | 0.934716167 | 617044.2813 | 411146.2656 | 1562971.313 | 1166849.281 | 3476801.25 | 316505.9219 | 4171907.433 | 2950807.95 | 1562971.313 | 8529495.412 | 3611794.149 | 2678330.777 | |
| O75368 | Adapter SH3BGRL [OS=Homo sapiens] | -3.17 | 0.934716167 | | 93609.28125 | 1781825.625 | | 1359497.188 | | | 671836.3619 | 1781825.625 | | 1412282.047 | | |
| P27824 | Calnexin [OS=Homo sapiens] | 0.22 | 0.934716167 | 1635226.879 | 1070700.285 | 12588050.88 | 1532669.332 | 14052284.5 | 1004260.328 | 11055957.21 | 7684445.117 | 12588050.88 | 11203585.8 | 14597889.06 | 8498233.87 | |
| A6NCE7 | Microtubule-associated proteins 1A/1B light chain 3 beta 2 [OS=Homo sapi | -3.27 | 0.934716167 | | 191070.3125 | 1443299.625 | | 1418622.375 | | | 1371316.838 | 1443299.625 | | 1473702.874 | | |
| P06241 | Tyrosine-protein kinase Fyn [OS=Homo sapiens] | -3.28 | 0.935199566 | | 192950.3789 | 2723497.688 | | 1224930.031 | | | 1384810.126 | 2723497.688 | | 1272490.085 | | |
| P60981 | Destrin [OS=Homo sapiens] | 0.17 | 0.935199566 | 597819.75 | 1120697.578 | 15861195.94 | 718618.8828 | 10369567.72 | 580182.6328 | 4041928.164 | 8043277.051 | 15861195.94 | 5252997.592 | 10772184.35 | 4909611.146 | Met-loss+Acetyl [N-Term] |
| Q9ULB1 | Neurexin-1 [OS=Homo sapiens] | -3.2 | 0.935902223 | | 36210.34375 | 3316595.469 | | 1784635.781 | | | 259882.6236 | 3316595.469 | | 1853927.392 | | |
| P62191 | 26S proteasome regulatory subunit 4 [OS=Homo sapiens] | 0.13 | 0.935902223 | 86123.87109 | 1561493.544 | 52274407.03 | 167523.0605 | 3247773.5 | 14418.59277 | 582293.4089 | 11206881.71 | 52274407.03 | 1224568.759 | 3373873.995 | 122012.7591 | |
| P60953 | Cell division control protein 42 homolog [OS=Homo sapiens] | -2.22 | 0.935902223 | | 589573.2656 | 19482275.5 | | 6538912.781 | 45920.13672 | | 4231383.39 | 19482275.5 | | 6792797.524 | 388584.5634 | |
| O00193 | Small acidic protein [OS=Homo sapiens] | 0.99 | 0.936261371 | 5784781.91 | 1119513.746 | 8468832.813 | 6784836.516 | 21789060.13 | 6209868.512 | 39111576.56 | 8034780.656 | 8468832.813 | 49596149.96 | 22635058.55 | 52549038.76 | |
| P49802 | Regulator of G-protein signaling 7 [OS=Homo sapiens] | 0.38 | 0.937020382 | 553051.875 | 910718.0156 | 3336511.563 | 665102.125 | 4004812.438 | 478081.6875 | 3739247.406 | 6536248.01 | 3336511.563 | 4861798.019 | 4160306.296 | 4045614.345 | |
| O15400 | Syntaxin-7 [OS=Homo sapiens] | 0 | 0.938311269 | | | 1300257.438 | | 746837.7813 | | | | 1300257.438 | | 775835.0663 | | |
| Q08211 | ATP-dependent RNA helicase A [OS=Homo sapiens] | 0.12 | 0.938569804 | 700472.5156 | 1526016.887 | 13378274.34 | 1096579.354 | 12124336.81 | 323269.2461 | 4735975.332 | 10952264.77 | 13378274.34 | 8015832.649 | 12595085.43 | 2735563.259 | |
| P80511 | Protein S100-A12 [OS=Homo sapiens] | 0 | 0.938569804 | | | 139029.4063 | | 2457429.375 | | | | 139029.4063 | | 2552843.375 | | Met-loss [N-Term] |
| Q7KZF4 | Staphylococcal nuclease domain-containing protein 1 [OS=Homo sapiens] | 0.4 | 0.938569804 | 38545.44922 | 178628.5039 | 3385844.469 | 56134.61719 | 2054776.828 | 80392.51563 | 260610.2203 | 1282021.639 | 3385844.469 | 410335.737 | 2134557.138 | 680296.1145 | |
| P61601 | Neurocalcin-delta [OS=Homo sapiens] | -3.29 | 0.938569804 | | 254154.8633 | 1431070.844 | | 2142127.406 | | | 1824076.378 | 1431070.844 | | 2225299.255 | | |
| Q13617 | Cullin-2 [OS=Homo sapiens] | 0 | 0.938569804 | | | 1401277.781 | | 1184129.484 | | | | 1401277.781 | | 1230105.386 | | |
| Q9Y3E1 | Hepatoma-derived growth factor-related protein 3 [OS=Homo sapiens] | -3.27 | 0.93867458 | | 161502.3066 | 2953986.625 | | 1844005.781 | | | 1159106.455 | 2953986.625 | | 1915602.536 | | |
| O60763 | General vesicular transport factor p115 [OS=Homo sapiens] | -3.54 | 0.93867458 | | 282443.3125 | 2253824.781 | | 1664970.719 | | | 2027103.349 | 2253824.781 | | 1729616.123 | | |
| P30040 | Endoplasmic reticulum resident protein 29 [OS=Homo sapiens] | -3.25 | 0.93867458 | | 216309.7813 | 1818493.75 | | 1659590.375 | | | 1552461.193 | 1818493.75 | | 1724026.878 | | |
| P59998 | Actin-related protein 2/3 complex subunit 4 [OS=Homo sapiens] | -0.01 | 0.93867458 | 83349.29688 | 2360351.125 | 18958682.38 | 202808.4795 | 15402260.44 | 110105.2598 | 563534.1931 | 16940304.33 | 18958682.38 | 1482499.945 | 16000280.18 | 931730.769 | Met-loss+Acetyl [N-Term] |
| Q02878 | Large ribosomal subunit protein eL6 [OS=Homo sapiens] | -0.12 | 0.93867458 | 885750.375 | 1259411.402 | 6316554.625 | 1114039.313 | 6996518.719 | 620477.9766 | 5988660.272 | 9038829.946 | 6316554.625 | 8143462.363 | 7268170.814 | 5250597.687 | |
| P62917 | Large ribosomal subunit protein uL2 [OS=Homo sapiens] | -0.51 | 0.93867458 | 648847.3125 | 1751342.016 | 9882672.25 | 696629.1992 | 8773437.25 | 212019.2773 | 4386931.389 | 12569429.36 | 9882672.25 | 5092256.262 | 9114081.32 | 1794145.754 | |
| P43003 | Excitatory amino acid transporter 1 [OS=Homo sapiens] | -2.11 | 0.93867458 | | 5025210.938 | 65137788.63 | | 52535980.34 | 197059.6094 | | 36066075.81 | 65137788.63 | | 54575781.81 | 1667554.318 | |
| Q9Y2J0 | Rabphilin-3A [OS=Homo sapiens] | -3.52 | 0.942194548 | | 202879.1367 | 3626637.688 | | 1591077.406 | | | 1456069.092 | 3626637.688 | | 1652853.774 | | |
| O75348 | V-type proton ATPase subunit G 1 [OS=Homo sapiens] | -1.43 | 0.942194548 | | 146754.6719 | 1167242 | | 719235.4375 | 26614.00781 | | 1053262.278 | 1167242 | | 747161.0132 | 225212.5831 | Met-loss+Acetyl [N-Term] |
| O15240 | Neurosecretory protein VGF [OS=Homo sapiens] | -3.37 | 0.942338745 | | 165711.9141 | 1609453.156 | | 1894026.063 | | | 1189318.922 | 1609453.156 | | 1967564.942 | | |
| P84074 | Neuron-specific calcium-binding protein hippocalcin [OS=Homo sapiens] | -3.21 | 0.94253991 | | 287250.9531 | 1797461.5 | | 2394945.5 | | | 2061607.917 | 1797461.5 | | 2487933.454 | | |
| Q9NPG2 | Neuroglobin [OS=Homo sapiens] | -3.29 | 0.94253991 | | 208981.6719 | 1583619 | | 1498747.125 | | | 1499867.153 | 1583619 | | 1556938.607 | | |
| P52597 | Heterogeneous nuclear ribonucleoprotein F [OS=Homo sapiens] | 0.34 | 0.94253991 | 105231.6641 | 65348.07813 | 298593.2813 | 123584.6484 | 405171.375 | 65837.03125 | 711483.3972 | 469004.9371 | 298593.2813 | 903385.4747 | 420902.8634 | 557124.9537 | |
| O00571 | ATP-dependent RNA helicase DDX3X [OS=Homo sapiens] | 0.61 | 0.94253991 | 461153.2227 | 708145.3086 | 8867050.703 | 838191.1797 | 7659721.625 | 217533.8477 | 3117910.036 | 5082378.173 | 8867050.703 | 6127053.371 | 7957123.735 | 1840811.053 | |
| Q12905 | Interleukin enhancer-binding factor 2 [OS=Homo sapiens] | -3.2 | 0.94253991 | 41751.98047 | 705716.457 | 5929954.469 | 140680.625 | 6209943.469 | | 282289.9473 | 5064946.239 | 5929954.469 | 1028354.531 | 6451055.402 | | |
| Q96CX2 | BTB/POZ domain-containing protein KCTD12 [OS=Homo sapiens] | -0.99 | 0.94253991 | 100347.0781 | 101874.0391 | 961160.6875 | | 558767.9375 | 41837.81641 | 678458.1493 | 731152.7538 | 961160.6875 | | 580463.0815 | 354039.2251 | |
| P51148 | Ras-related protein Rab-5C [OS=Homo sapiens] | -0.14 | 0.94253991 | 378434.4355 | 2244854.531 | 21380478.31 | 441622.5234 | 16052592.75 | 320642.0781 | 2558638.792 | 16111382.13 | 21380478.31 | 3228195.233 | 16675862.78 | 2713331.685 | |
| Q9NRV9 | Heme-binding protein 1 [OS=Homo sapiens] | -3.05 | 0.94253991 | | 118482.2656 | 2189151 | | 1199733.25 | | | 850350.4484 | 2189151 | | 1246314.995 | | |
| Q8TCJ2 | Dolichyl-diphosphooligosaccharide--protein glycosyltransferase subunit ST | 1.04 | 0.94253991 | 534855.8008 | 121261.1641 | 379432.5938 | 995128.7422 | 1505474.375 | 180304.2617 | 3616221.653 | 870294.6782 | 379432.5938 | 7274243.707 | 1563927.055 | 1525767.514 | |
| P49593 | Protein phosphatase 1F [OS=Homo sapiens] | 0 | 0.94253991 | | | 1053744.563 | | 576739.5 | | | | 1053744.563 | | 599132.4214 | | |
| O95373 | Importin-7 [OS=Homo sapiens] | -3.16 | 0.942644744 | | 160391.5 | 2382314.75 | | 1506332.594 | | | 1151134.165 | 2382314.75 | | 1564818.595 | | |
| P68400 | Casein kinase II subunit alpha [OS=Homo sapiens] | 0.1 | 0.942644744 | | 602312.3555 | 6134963.281 | 22582.19336 | 4274128.031 | | | 4322812.185 | 6134963.281 | 165072.4886 | 4440078.539 | | |
| Q15181 | Inorganic pyrophosphatase [OS=Homo sapiens] | -3.15 | 0.942644744 | | 159448.0195 | 2225229.438 | | 1272354.922 | | | 1144362.78 | 2225229.438 | | 1321756.33 | | |
| Q15008 | 26S proteasome non-ATPase regulatory subunit 6 [OS=Homo sapiens] | -0.02 | 0.942644744 | | 515981.8203 | 5081325.438 | 90096.35938 | 4875622.281 | | | 3703215.582 | 5081325.438 | 658591.042 | 5064926.857 | | Met-loss [N-Term] |
| O60883 | G-protein coupled receptor 37-like 1 [OS=Homo sapiens] | -3.18 | 0.943727264 | | 246151.1172 | 2773949.125 | | 2305970.875 | | | 1766633.274 | 2773949.125 | | 2395504.234 | | |
| Q15366 | Poly(rC)-binding protein 2 [OS=Homo sapiens] | -0.07 | 0.943727264 | 101260.9531 | 1826133.906 | 17470303.25 | 265307.4844 | 14062033.06 | 172633.0371 | 684636.9634 | 13106212.8 | 17470303.25 | 1939358.413 | 14608016.13 | 1460852.213 | |
| Q6PI78 | Transmembrane protein 65 [OS=Homo sapiens] | -3.17 | 0.943727264 | | 105489.6055 | 1206850.969 | | 1521580.969 | | | 757101.7724 | 1206850.969 | | 1580659.015 | | |
| Q03135 | Caveolin-1 [OS=Homo sapiens] | 0.31 | 0.943727264 | 525889.2813 | 281013.832 | 2004300.594 | 1177838.906 | 2163580.313 | 236228.0547 | 3555597.982 | 2016843.929 | 2004300.594 | 8609827.943 | 2247585.108 | 1999004.839 | |
| Q02156 | Protein kinase C epsilon type [OS=Homo sapiens] | -3 | 0.943727264 | | 116342.8633 | 3697914.688 | | 2348871.664 | | | 834995.8995 | 3697914.688 | | 2440070.721 | | Met-loss [N-Term] |
| Q96PE3 | Inositol polyphosphate-4-phosphatase type I A [OS=Homo sapiens] | -3.29 | 0.943973281 | | 254991.5352 | 2259671.031 | | 1764696.719 | | | 1830081.195 | 2259671.031 | | 1833214.16 | | |
| P50570 | Dynamin-2 [OS=Homo sapiens] | -2.99 | 0.945361677 | | 107719.25 | 1744899.594 | | 1227789.344 | | | 773103.992 | 1744899.594 | | 1275460.416 | | |
| O14735 | CDP-diacylglycerol--inositol 3-phosphatidyltransferase [OS=Homo sapiens | -3.25 | 0.946570607 | | 276730.4141 | 1407033.406 | | 1796709.563 | | | 1986101.721 | 1407033.406 | | 1866469.959 | | |
| Q9C0H9 | SRC kinase signaling inhibitor 1 [OS=Homo sapiens] | -3.21 | 0.946570607 | | 125859.8594 | 2972420.969 | | 1969788.891 | | | 903299.6397 | 2972420.969 | | 2046269.395 | | |
| Q99259 | Glutamate decarboxylase 1 [OS=Homo sapiens] | -3.31 | 0.948048493 | | 128043.9844 | 2588127.063 | | 1812795.266 | | | 918975.1643 | 2588127.063 | | 1883180.217 | | |
| P05387 | Large ribosomal subunit protein P2 [OS=Homo sapiens] | 0.34 | 0.948680899 | 369495.8672 | 2208499.684 | 18570980.84 | 1067817.129 | 15273180.81 | 356229.4453 | 2498204.102 | 15850462.4 | 18570980.84 | 7805585.047 | 15866188.81 | 3014478.47 | |
| O95372 | Acyl-protein thioesterase 2 [OS=Homo sapiens] | -3.27 | 0.949026688 | | 195772.9219 | 3375614.125 | | 1877850.563 | | | 1405067.594 | 3375614.125 | | 1950761.4 | | |
| P50395 | Rab GDP dissociation inhibitor beta [OS=Homo sapiens] | 0.16 | 0.949026688 | 22247.61719 | 1114293.754 | 23169330.47 | 46882.65625 | 11589965.47 | | 150418.7014 | 7997316.63 | 23169330.47 | 342705.2729 | 12039966.18 | | |
| Q02413 | Desmoglein-1 [OS=Homo sapiens] | 0.29 | 0.949026688 | 1786854.469 | 960360.0547 | 766007.1406 | 1868065.66 | 4188504.75 | 2603147.508 | 12081128.8 | 6892530.277 | 766007.1406 | 13655283.28 | 4351130.784 | 22028308.5 | |
| Q07020 | Large ribosomal subunit protein eL18 [OS=Homo sapiens] | 0 | 0.950271117 | 470182.7734 | 910718.7344 | 3573577.063 | 702865.25 | 5241601.313 | 278667.8281 | 3178959.869 | 6536253.169 | 3573577.063 | 5137840.869 | 5445115.665 | 2358137.934 | |
| P62942 | Peptidyl-prolyl cis-trans isomerase FKBP1A [OS=Homo sapiens] | -0.14 | 0.950271117 | 1302576.375 | 1617250.895 | 20579428.38 | 1041902 | 13159185.94 | 885116.5 | 8806868.852 | 11607053.73 | 20579428.38 | 7616149.293 | 13670114.38 | 7490017.088 | Met-loss [N-Term] |
| Q01650 | Large neutral amino acids transporter small subunit 1 [OS=Homo sapiens] | 0.91 | 0.950628616 | 542365.0469 | 221203.0938 | 824563.875 | 876677.1563 | 1349033.875 | 337975.2422 | 3666992.531 | 1587580.63 | 824563.875 | 6408380.159 | 1401412.478 | 2860008.077 | |
| Q7Z4S6 | Kinesin-like protein KIF21A [OS=Homo sapiens] | -2.96 | 0.950628616 | | 144431.9063 | 3316355.281 | | 2315788.938 | | | 1036591.726 | 3316355.281 | | 2405703.5 | | |
| Q9UDW1 | Cytochrome b-c1 complex subunit 9 [OS=Homo sapiens] | -0.04 | 0.950628616 | | 669787.0625 | 6060013.5 | 57781.88281 | 4506232.5 | 91349.73438 | | 4807079.995 | 6060013.5 | 422377.0047 | 4681194.87 | 773018.0959 | |
| P16152 | Carbonyl reductase [NADPH] 1 [OS=Homo sapiens] | -2.14 | 0.950628616 | 42721.42969 | 1227641.383 | 10279584.31 | 45535.54297 | 7173660.563 | 33053.23828 | 288844.5051 | 8810815.651 | 10279584.31 | 332858.0744 | 7452190.5 | 279702.5245 | |
| Q9UMX0 | Ubiquilin-1 [OS=Homo sapiens] | -3.06 | 0.950631187 | | 190875.1914 | 1440357.313 | | 1661645.5 | | | 1369916.449 | 1440357.313 | | 1726161.797 | | |
| Q14847 | LIM and SH3 domain protein 1 [OS=Homo sapiens] | -0.07 | 0.950631187 | | 865302.1172 | 7021011.563 | 73130.28906 | 5398246.5 | | | 6210296.87 | 7021011.563 | 534571.581 | 5607842.876 | | |
| O60939 | Sodium channel subunit beta-2 [OS=Homo sapiens] | 0 | 0.950673042 | | | 1542750.438 | | 758203.75 | | | | 1542750.438 | | 787642.3387 | | |
| Q86Y39 | NADH dehydrogenase [ubiquinone] 1 alpha subcomplex subunit 11 [OS=Hc | 0 | 0.950930709 | | | 1540247.625 | | 617861.9375 | | | | 1540247.625 | | 641851.5095 | | |
| Q9UQ16 | Dynamin-3 [OS=Homo sapiens] | -0.07 | 0.951952263 | 33306.80469 | 1601051.664 | 12843784.97 | 54352.70313 | 8244930.266 | | 225191.1415 | 11490791.41 | 12843784.97 | 397310.2091 | 8565054.126 | | |
| Q7L2H7 | Eukaryotic translation initiation factor 3 subunit M [OS=Homo sapiens] | 0 | 0.953021875 | | | 1099170.375 | | 890944.25 | | | | 1099170.375 | | 925536.7212 | | |
| P07954 | Fumarate hydratase, mitochondrial [OS=Homo sapiens] | -0.37 | 0.953878939 | 30417.94922 | 2113692.164 | 22118633.13 | 267723.5547 | 14802585.94 | 131869.9727 | 205659.257 | 15170026.25 | 22118633.13 | 1957019.53 | 15377322.27 | 1115907.735 | |
| Q9UBB4 | Ataxin-10 [OS=Homo sapiens] | -3.27 | 0.953878939 | | 150220.4922 | 2701050.188 | | 1738043.25 | | | 1078136.565 | 2701050.188 | | 1805525.824 | | |
| P61020 | Ras-related protein Rab-5B [OS=Homo sapiens] | -2.25 | 0.954442347 | | 19088.65234 | 2508766.75 | | 1151218.875 | | | 136999.7779 | 2508766.75 | | 1195916.964 | | |

| | | | | | | | | | | | | | | | | |
|---|---|---|---|---|---|---|---|---|---|---|---|---|---|---|---|---|
| P35052 | Glypican-1 [OS=Homo sapiens] | 0.86 | 0.95473097 | 22632931.08 | 1597299.117 | 3262088.742 | 34806658.48 | 22552296.86 | 3499827.336 | 153023853 | 11463859.28 | 3262088.742 | 254431518 | 23427929.29 | 29616176.57 | |
| O14737 | Programmed cell death protein 5 [OS=Homo sapiens] | -3.19 | 0.95473097 | | 269231.2188 | 2385073.688 | | 1871562.313 | | | 1932279.792 | 2385073.688 | | 1944228.998 | | |
| O75306 | NADH dehydrogenase [ubiquinone] iron-sulfur protein 2, mitochondrial [OS | -0.23 | 0.9548759 | 221900.5059 | 2479771.133 | 23442740.25 | 197867.9629 | 15704374.53 | 154841.2773 | 1500294.87 | 17797384.98 | 23442740.25 | 1446385.501 | 16314124.38 | 1310295.1 | |
| P49327 | Fatty acid synthase [OS=Homo sapiens] | -0.03 | 0.955716038 | 467982.1885 | 975668.2734 | 6987438.938 | 894813.3164 | 6234671.313 | 134906.9648 | 3164081.46 | 7002397.78 | 6987438.938 | 6540952.803 | 6476743.347 | 1141607.316 | |
| Q13217 | DnaJ homolog subfamily C member 3 [OS=Homo sapiens] | -1.23 | 0.956863733 | | 109429.0703 | 1158669.5 | | 787395.5625 | 23985.10352 | | 785375.419 | 1158669.5 | | 817967.5745 | 202966.316 | |
| O75489 | NADH dehydrogenase [ubiquinone] iron-sulfur protein 3, mitochondrial [OS | -0.24 | 0.956863733 | | 874859.5781 | 8050757.375 | 33098.51563 | 5710198 | 52720.83203 | | 6278891.027 | 8050757.375 | 241945.2467 | 5931906.44 | 446133.2862 | |
| O43617 | Trafficking protein particle complex subunit 3 [OS=Homo sapiens] | -3.3 | 0.958366348 | | 261837.6484 | 1936698.938 | | 1674294.438 | | | 1879215.936 | 1936698.938 | | 1739301.852 | | |
| P11217 | Glycogen phosphorylase, muscle form [OS=Homo sapiens] | -3.08 | 0.959800541 | | 75736.82813 | 1724790.938 | | 1584684.781 | | | 543565.2789 | 1724790.938 | | 1646212.944 | | |
| Q99757 | Thioredoxin, mitochondrial [OS=Homo sapiens] | 0 | 0.959800541 | | | 1810892.188 | | 489429.375 | | | | 1810892.188 | | 508432.3278 | | |
| Q15393 | Splicing factor 3B subunit 3 [OS=Homo sapiens] | 0.53 | 0.959800541 | 1293962.313 | 475495.2344 | 2613046.922 | 1591004.121 | 5393601.406 | 340768.0195 | 8748628.183 | 3412642.252 | 2613046.922 | 11630004.47 | 5603017.429 | 2883641.068 | |
| Q13151 | Heterogeneous nuclear ribonucleoprotein A0 [OS=Homo sapiens] | -3.14 | 0.959800541 | | 268696.2617 | 2574800.125 | | 1958741.313 | | | 1928440.391 | 2574800.125 | | 2034792.875 | | |
| Q96QD8 | Sodium-coupled neutral amino acid symporter 2 [OS=Homo sapiens] | 0.87 | 0.959800541 | 674789.5938 | 80427.32813 | 430564.0938 | 915514.1719 | 1316954.438 | 254777.3711 | 4562330.14 | 577229.1251 | 430564.0938 | 6692272.99 | 1368087.5 | 2155972.534 | |
| P13611 | Versican core protein [OS=Homo sapiens] | 0.84 | 0.959800541 | 2473741.262 | 248047.9688 | 2915677.234 | 4133289.984 | 4154904.359 | 706127.1289 | 16725249.5 | 1780247.03 | 2915677.234 | 30213737.56 | 4316225.799 | 5975376.419 | |
| P62906 | Large ribosomal subunit protein uL1 [OS=Homo sapiens] | -0.4 | 0.959800541 | 198597.9844 | 2220266.992 | 12962897.31 | 237717.3086 | 10491099.94 | 153449.75 | 1342743.839 | 15934916.69 | 12962897.31 | 1737678.315 | 10898435.27 | 1298519.743 | |
| P63027 | Vesicle-associated membrane protein 2 [OS=Homo sapiens] | -1.97 | 0.959800541 | | 9854183.578 | 67116647.06 | | 54907717.75 | 100773.6406 | | 70723744.01 | 67116647.06 | | 57039606.07 | 852764.9076 | |
| Q2TAY7 | WD40 repeat-containing protein SMU1 [OS=Homo sapiens] | 0.55 | 0.961346691 | 337752.3281 | 81614.71094 | 1111636.719 | 518183.5938 | 825221.625 | 130444.7266 | 2283582.38 | 585751.0039 | 1111636.719 | 3787845.317 | 857262.3001 | 1103847.043 | |
| Q92747 | Actin-related protein 2/3 complex subunit 1A [OS=Homo sapiens] | 0.28 | 0.961346691 | | 916238.5781 | 10282518.94 | 16819.16406 | 7674144.531 | | | 6575869.238 | 10282518.94 | 122945.598 | 7972106.635 | | Met-loss+Acetyl [N-Term] |
| O75914 | Serine/threonine-protein kinase PAK 3 [OS=Homo sapiens] | 0.28 | 0.961364221 | | 694834.9063 | 8060297.406 | 220180.2031 | 6150165.25 | | | 4986849.052 | 8060297.406 | 1609484.672 | 6388956.189 | | |
| P51858 | Hepatoma-derived growth factor [OS=Homo sapiens] | 0.38 | 0.961697682 | | 103626.9609 | 2590907.563 | 190147.9063 | 2200563.031 | 68113.59375 | | 743733.5219 | 2590907.563 | 1389953.03 | 2286003.746 | 576389.6404 | |
| Q9Y2T3 | Guanine deaminase [OS=Homo sapiens] | -0.11 | 0.961697682 | 39487.35059 | 2110236.629 | 17056260.78 | 51428.82813 | 14128846.5 | 11436.19922 | 266978.5239 | 15145225.78 | 17056260.78 | 375937.116 | 14677423.71 | 96775.20143 | |
| P27797 | Calreticulin [OS=Homo sapiens] | 0.8 | 0.963683325 | 7201764.281 | 2278155.484 | 20547382.73 | 6754993.324 | 32560132.91 | 5355287.43 | 48691957.53 | 16350383.98 | 20547382.73 | 49378000.65 | 33824337.1 | 45317417.95 | |
| P15170 | Eukaryotic peptide chain release factor GTP-binding subunit ERF3A [OS=Ho | 0.41 | 0.963683325 | 43328.51172 | 336242.4434 | 3578694.875 | 165610.125 | 2256230.688 | 56422.52246 | 292949.0566 | 2413221.177 | 3578694.875 | 1210585.484 | 2343832.796 | 477457.6651 | |
| P62072 | Mitochondrial import inner membrane translocase subunit Tim10 [OS=Hom | 0.16 | 0.963683325 | | 353644.168 | 6696532.813 | 15542.77832 | 4102177.125 | 19724.09375 | | 2538113.828 | 6696532.813 | 113615.4073 | 4261451.338 | 166908.875 | |
| Q04695 | Keratin, type I cytoskeletal 17 [OS=Homo sapiens] | 0.05 | 0.963683325 | 258033.4375 | 220249.3438 | 304282.9531 | 548119.9141 | 3575143.969 | 1462411.395 | 1744593.781 | 1580735.541 | 304282.9531 | 4006675.384 | 3713955.196 | 12375191.67 | |
| P09622 | Dihydrolipoyl dehydrogenase, mitochondrial [OS=Homo sapiens] | -0.19 | 0.963683325 | 497303.4688 | 2317964.996 | 25368028.45 | 461318.8789 | 16378575.78 | 390242.2344 | 3362326.012 | 16636097.93 | 25368028.45 | 3372172.674 | 17014502.68 | 3302300.888 | |
| Q13153 | Serine/threonine-protein kinase PAK 1 [OS=Homo sapiens] | 0.17 | 0.963683325 | 167482.4531 | 990435.1992 | 14376875.41 | | 8298118.047 | 90099.21875 | 1132368.15 | 7108380.409 | 14376875.41 | | 8620307.016 | 762436.0049 | |
| P62280 | Small ribosomal subunit protein uS17 [OS=Homo sapiens] | -0.44 | 0.963683325 | 371014.2383 | 1813001.465 | 12616233.09 | 547374.8613 | 11198865.88 | 232884.3672 | 2508469.984 | 13011960.91 | 12616233.09 | 4001229.159 | 11633681.46 | 1970709.946 | Met-loss+Acetyl [N-Term] |
| P02462 | Collagen alpha-1(IV) chain [OS=Homo sapiens] | 0.32 | 0.963683325 | 52255.63672 | 89787.89063 | 611649.7188 | 60246.54688 | 447722.3438 | 58014.69922 | 353306.3766 | 644410.1372 | 611649.7188 | 440393.3339 | 465105.948 | 490930.9549 | |
| Q13526 | Peptidyl-prolyl cis-trans isomerase NIMA-interacting 1 [OS=Homo sapiens] | -3.21 | 0.963683325 | | 147670.2188 | 2947733 | | 1798083.25 | | | 1059833.183 | 2947733 | | 1867896.982 | | |
| P62314 | Small nuclear ribonucleoprotein Sm D1 [OS=Homo sapiens] | 0.48 | 0.963683325 | 76100.30078 | 441233.3477 | 3848149 | 111708.2305 | 3534365.438 | 70605.74219 | 514522.8959 | 3166743.757 | 3848149 | 816570.6185 | 3671593.367 | 597478.6545 | |
| Q9Y639 | Neuroplastin [OS=Homo sapiens] | -1.86 | 0.966143565 | | 831394.1016 | 17408824.47 | | 10967821.03 | 88719.0625 | | 5966938.118 | 17408824.47 | | 11393665.89 | 750756.8712 | |
| P13667 | Protein disulfide-isomerase A4 [OS=Homo sapiens] | 0 | 0.966143565 | | | 992838.8125 | | 868462.75 | | | | 992838.8125 | | 902182.3375 | | |
| O15126 | Secretory carrier-associated membrane protein 1 [OS=Homo sapiens] | 0.02 | 0.966143565 | | 398201.625 | 3509856.813 | 67151.49219 | 2561287.969 | | | 2857903.911 | 3509856.813 | 490867.4614 | 2660734.461 | | Met-loss+Acetyl [N-Term] |
| P48735 | Isocitrate dehydrogenase [NADP], mitochondrial [OS=Homo sapiens] | -0.29 | 0.966143565 | 468395.9727 | 2124000.551 | 24520497.69 | 472830.3389 | 15139160.66 | 447699.0313 | 3166879.103 | 15244009.82 | 24520497.69 | 3456319.741 | 15726965.09 | 3788510.772 | |
| P05388 | Large ribosomal subunit protein uL10 [OS=Homo sapiens] | -0.44 | 0.966143565 | 440553.625 | 2082384.297 | 16728744.47 | 343317.3945 | 14261997.84 | 129410.4023 | 2978633.785 | 14945328.83 | 16728744.47 | 2509599.302 | 14815744.89 | 1095094.403 | |
| P62995 | Transformer-2 protein homolog beta [OS=Homo sapiens] | -0.19 | 0.966143565 | 1348277.875 | 456322.3672 | 3918317.969 | 921631.2188 | 4004724.703 | 614653.3984 | 9115861.955 | 3275038.062 | 3918317.969 | 6736987.697 | 4160215.155 | 5201309.045 | |
| Q9NRW1 | Ras-related protein Rab-6B [OS=Homo sapiens] | -0.05 | 0.966143565 | 40721.05469 | 1102889.176 | 14334061.94 | 44509.62109 | 7969316.594 | | 275319.7394 | 7915465.662 | 14334061.94 | 325358.7374 | 8278739.27 | | Met-loss+Acetyl [N-Term] |
| Q96AE4 | Far upstream element-binding protein 1 [OS=Homo sapiens] | 0.4 | 0.966143565 | 246148.4844 | 366128.4199 | 2767906.969 | 233027.0264 | 2109647.188 | 161901.5625 | 1664238.245 | 2627713.645 | 2767906.969 | 1703393.047 | 2191557.935 | 1370040.52 | |
| Q92688 | Acidic leucine-rich nuclear phosphoprotein 32 family member B [OS=Homo | 0.77 | 0.966143565 | 3966776.484 | 881883.8516 | 7159161.672 | 5742639.734 | 12481181.63 | 3381829.641 | 26819832.55 | 6329304.429 | 7159161.672 | 41977845.86 | 12965785.36 | 28617658.58 | |
| P01040 | Cystatin-A [OS=Homo sapiens] | 0.63 | 0.966143565 | 378860.5859 | 202356.8242 | 271812 | 629342.8047 | 2646204.563 | 745264.6563 | 2561520.044 | 1452320.441 | 271812 | 4600402.684 | 2748948.09 | 6306565.306 | |
| P05556 | Integrin beta-1 [OS=Homo sapiens] | 0.35 | 0.966143565 | 208649.8906 | 118183.4531 | 714839.8125 | 245140.9219 | 722307.75 | 190968 | 1410705.935 | 848205.8629 | 714839.8125 | 1791943.829 | 750352.6137 | 1616006.01 | |
| P60900 | Proteasome subunit alpha type-6 [OS=Homo sapiens] | -0.14 | 0.966143565 | 266154.7813 | 1462576.207 | 11752816.06 | 232809.9668 | 8909194.5 | 263451.2734 | 1799503.121 | 10496949.28 | 11752816.06 | 1701806.373 | 9255109.583 | 2229372.676 | |
| P07355 | Annexin A2 [OS=Homo sapiens] | 0.69 | 0.966143565 | 1917317.881 | 891400.0254 | 4103522.922 | 2822004.488 | 16747034 | 1618203.578 | 12963206.96 | 6397602.268 | 4103522.922 | 20628434.81 | 17397266.93 | 13693533.51 | Met-loss+Acetyl [N-Term] |
| P22001 | Potassium voltage-gated channel subfamily A member 3 [OS=Homo sapien | -2.87 | 0.966143565 | | 87947.28516 | 2642545.938 | | 1108865.688 | | | 631200.0616 | 2642545.938 | | 1151919.34 | | |
| P0DMV9 | Heat shock 70 kDa protein 1B [OS=Homo sapiens] | 0.62 | 0.966143565 | 422292.3828 | 364196.3086 | 2585732.469 | 580595.1797 | 6663949.125 | 266440.1875 | 2855167.424 | 2613846.829 | 2585732.469 | 4244064.766 | 6922688.623 | 2254665.411 | |
| O95202 | Mitochondrial proton/calcium exchanger protein [OS=Homo sapiens] | -3.03 | 0.966143565 | | 206411.0547 | 3607601.313 | | 1702967.563 | | | 1481417.763 | 3607601.313 | | 1769088.261 | | |
| Q9UHV9 | Prefoldin subunit 2 [OS=Homo sapiens] | -3.38 | 0.966143565 | | 265186.6836 | 1965120.156 | | 1750714.906 | | | 1903252.053 | 1965120.156 | | 1818689.479 | | |
| P62879 | Guanine nucleotide-binding protein G(I)/G(S)/G(T) subunit beta-2 [OS=Hom | -0.15 | 0.966143565 | 254328.1602 | 2229162.213 | 46161941.13 | 45992.05078 | 29156614.41 | 76476.04102 | 1719541.975 | 15998757.93 | 46161941.13 | 336195.0789 | 30288671.03 | 647154.1928 | Met-loss+Acetyl [N-Term] |
| P22307 | Sterol carrier protein 2 [OS=Homo sapiens] | -0.11 | 0.966143565 | 449216.0938 | 1109182.313 | 3214719.5 | 383910.4375 | 2550074.531 | 639769.1563 | 3037201.733 | 7960631.676 | 3214719.5 | 2806328.433 | 2649085.642 | 5413843.164 | |
| O43396 | Thioredoxin-like protein 1 [OS=Homo sapiens] | -0.13 | 0.966143565 | | 647039.8984 | 7867510.594 | 39489.40234 | 4507117.125 | 31948.875 | | 4643822.979 | 7867510.594 | 288661.682 | 4682113.842 | 270357.2012 | Met-loss [N-Term] |
| Q9UKU0 | Long-chain-fatty-acid--CoA ligase 6 [OS=Homo sapiens] | 0.08 | 0.966143565 | 1957488.375 | 663221.0508 | 8816348.5 | 1050410.354 | 10672291.84 | 1669356.125 | 13234804.29 | 4759955.55 | 8816348.5 | 7678344.097 | 11086662.27 | 14126395.68 | |
| P62750 | Large ribosomal subunit protein uL23 [OS=Homo sapiens] | -0.35 | 0.966143565 | 61414.84766 | 2255685.672 | 15385976.75 | 169030 | 9199463.469 | 71804.83594 | 415232.8564 | 16189117.52 | 15385976.75 | 1235584.263 | 9556648.753 | 607625.6043 | |
| P81605 | Dermcidin [OS=Homo sapiens] | 0.72 | 0.966143565 | 4747756.5 | 4740882.805 | 15086059.78 | 7291394.586 | 19846282.09 | 11750516.62 | 32100128.36 | 34025445.05 | 15086059.78 | 53299014.42 | 20616848.76 | 99435012.52 | |
| O95741 | Copine-6 [OS=Homo sapiens] | -0.25 | 0.966143565 | | 676201.2734 | 6647765 | 36478.32813 | 4701697 | | | 4853114.962 | 6647765 | 266651.1754 | 4884248.622 | | |
| P11233 | Ras-related protein Ral-A [OS=Homo sapiens] | 0.21 | 0.966143565 | | 1144845.219 | 10700373.34 | 24368.00977 | 8667254 | 20022.82422 | | 8216585.326 | 10700373.34 | 178126.542 | 9003775.319 | 169436.7866 | |
| O75380 | NADH dehydrogenase [ubiquinone] iron-sulfur protein 6, mitochondrial [OS | 0.38 | 0.966143565 | 39163.75391 | 560262.9531 | 6187408.375 | 57401.05859 | 2976420.75 | 46936.70313 | 264790.6495 | 4021022.479 | 6187408.375 | 419593.2361 | 3091985.499 | 397186.9336 | |
| Q01130 | Serine/arginine-rich splicing factor 2 [OS=Homo sapiens] | -0.23 | 0.966143565 | 1744907.406 | 907226.1094 | 4277939 | 886709.1719 | 6454627 | 410349.3281 | 11797519.88 | 6511186.504 | 4277939 | 6481712.707 | 6705239.2 | 3472450.779 | Met-loss+Acetyl [N-Term] |
| P84103 | Serine/arginine-rich splicing factor 3 [OS=Homo sapiens] | -0.15 | 0.966143565 | 1161442.5 | 796151.2422 | 1836478.063 | 1922174.742 | 6851894.188 | 637347.6074 | 7852646.473 | 5713999.156 | 1836478.063 | 14050812.65 | 7117930.982 | 5393351.577 | |
| O15355 | Protein phosphatase 1G [OS=Homo sapiens] | 0.73 | 0.966143565 | 9967543.227 | 1854894.531 | 8547386.984 | 13773245.67 | 29331007.22 | 9019340.516 | 67391707.43 | 13312628.59 | 8547386.984 | 100680385.8 | 30469834.95 | 76323302.74 | |
| P01871 | Immunoglobulin heavy constant mu [OS=Homo sapiens] | 0.78 | 0.966143565 | 488112.6172 | 587573.7734 | 447782.3438 | 712263.2422 | 1981291.953 | 727144.1328 | 3300185.607 | 4217032.98 | 447782.3438 | 5206538.8 | 2058219.084 | 6153226.135 | |
| Q13033 | Striatin-3 [OS=Homo sapiens] | 0.71 | 0.966143565 | 250211424 | 17984384 | 133621948.4 | 238540919 | 203087404.7 | 200815615.6 | 1691708247 | 129074413.9 | 133621948.4 | 1743698785 | 210972628.9 | 1699338327 | |
| Q99613 | Eukaryotic translation initiation factor 3 subunit C [OS=Homo sapiens] | 0.7 | 0.966143565 | 37068549.97 | 2851009.586 | 6928466.227 | 12402995.45 | 38428019.63 | 9443889.836 | 250624734.5 | 20461773.47 | 6928466.227 | 90664059.57 | 39920054.8 | 79915916.44 | |
| P0C0S8 | Histone H2A type 1 [OS=Homo sapiens] | -0.15 | 0.966143565 | 181216.7969 | 1675135.156 | 11309321.81 | 151116.1563 | 23715606.75 | 164047.3594 | 1225227.629 | 12022490.65 | 11309321.81 | 1104636.719 | 24636406.72 | 1388198.644 | |
| O96000 | NADH dehydrogenase [ubiquinone] 1 beta subcomplex subunit 10 [OS=Hor | -2.86 | 0.966143565 | | 102780.4141 | 1859178.25 | | 1338321 | | | 737657.8319 | 1859178.25 | | 1390283.657 | | |
| P84095 | Rho-related GTP-binding protein RhoG [OS=Homo sapiens] | -2.98 | 0.966143565 | | 106664.9063 | 1685611.563 | | 1477151.469 | | | 765536.9382 | 1685611.563 | | 1534504.462 | | |
| Q9UKG1 | DCC-interacting protein 13-alpha [OS=Homo sapiens] | -3 | 0.966143565 | | 152245.1621 | 2362581.547 | | 2161798.594 | | | 1092667.676 | 2362581.547 | | 2245734.211 | | |
| Q9H1E5 | Thioredoxin-related transmembrane protein 4 [OS=Homo sapiens] | -2.94 | 0.966143565 | | 163632.2715 | 1706907.5 | | 1369725.313 | | | 1174393.271 | 1706907.5 | | 1422907.297 | | |
| P31943 | Heterogeneous nuclear ribonucleoprotein H [OS=Homo sapiens] | -0.18 | 0.966143565 | 495823.7109 | 1484513.781 | 19048786.19 | 480960.5352 | 16464827.88 | 487645.2773 | 3352321.199 | 10654395.85 | 19048786.19 | 3515750.272 | 17104103.66 | 4126543.184 | |
| P08708 | Small ribosomal subunit protein eS17 [OS=Homo sapiens] | 0.31 | 0.966143565 | 123207.1484 | 529837.4609 | 4247863.625 | 214992.4453 | 3265578 | 99578.6875 | 833017.7168 | 3802657.892 | 4247863.625 | 1571562.931 | 3392369.786 | 842652.9965 | |
| P01008 | Antithrombin-III [OS=Homo sapiens] | 0.3 | 0.966143565 | 178612.9375 | 62333.03125 | 250858.625 | 183520.5313 | 531231.6875 | 105858.9688 | 1207622.636 | 447365.8635 | 250858.625 | 1341507.9 | 551857.6883 | 895797.8807 | |
| G2XKQ0 | Small ubiquitin-related modifier 5 [OS=Homo sapiens] | 0.33 | 0.966143565 | 121776.7344 | 137447.2578 | 989557.75 | 157803.8203 | 430316.9375 | 52925.125 | 823346.5227 | 986462.714 | 989557.75 | 1153522.553 | 447024.7464 | 447862.0505 | |
| P19338 | Nucleolin [OS=Homo sapiens] | 0.75 | 0.966143565 | 12222653.7 | 2805796.146 | 18958307.63 | 17207475.08 | 28422640.69 | 8315546.504 | 82638768.97 | 20137275.38 | 18958307.63 | 125784094.1 | 29526199.5 | 70367669.58 | |
| Q9H6Z4 | Ran-binding protein 3 [OS=Homo sapiens] | 0.29 | 0.966143565 | 89325.14063 | 73481.34375 | 625779.25 | 100986.0469 | 478765.4375 | 175372.5938 | 603937.5608 | 527377.606 | 625779.25 | 738193.0446 | 497354.3443 | 1484034.841 | |
| Q14689 | Disco-interacting protein 2 homolog A [OS=Homo sapiens] | 0.7 | 0.966143565 | 572335 | 450550.4961 | 1002387.969 | 859574.5625 | 1650555.781 | 911359.9375 | 3869622.834 | 3233613.185 | 1002387.969 | 6283362.732 | 1714641.501 | 7712093.838 | |
| P62273 | Small ribosomal subunit protein uS14 [OS=Homo sapiens] | 0.28 | 0.966143565 | 20802.20898 | 46734.99219 | 487416.1563 | 23422.89648 | 464494.125 | 31819.71875 | 140646.1301 | 335418.3122 | 487416.1563 | 171217.9039 | 482528.9231 | 269264.2575 | Met-loss [N-Term] |
| Q6P587 | Acylpyruvase FAHD1, mitochondrial [OS=Homo sapiens] | 0 | 0.966143565 | | | 987901.3438 | | 585128.7656 | | | | 987901.3438 | | 607847.4149 | | |
| P55795 | Heterogeneous nuclear ribonucleoprotein H2 [OS=Homo sapiens] | 0 | 0.966143565 | | | 1754140.219 | | 1294704.156 | | | | 1754140.219 | | 1344973.313 | | |
| Q9UM19 | Hippocalcin-like protein 4 [OS=Homo sapiens] | -3.02 | 0.966143565 | | 235161.043 | 2517442.625 | | 2001520.281 | | | 1687757.212 | 2517442.625 | | 2079232.812 | | |
| Q13185 | Chromobox protein homolog 3 [OS=Homo sapiens] | 0.27 | 0.967338278 | | 273840.5078 | 3299361.469 | 121264.2891 | 2935602.656 | | | 1965360.785 | 3299361.469 | 886423.9914 | 3049582.572 | | |
| Q9BWD1 | Acetyl-CoA acetyltransferase, cytosolic [OS=Homo sapiens] | 0.74 | 0.967338278 | | 224307.293 | 2825976.813 | 68104.99219 | 1653503.25 | | | 1609859.553 | 2825976.813 | 497837.4052 | 1717703.41 | | |
| Q9UQ80 | Proliferation-associated protein 2G4 [OS=Homo sapiens] | -0.31 | 0.967417127 | 958133.0508 | 918766.2539 | 4840509.125 | 1123440.434 | 5241910.281 | 362575.457 | 6478047.877 | 6594010.435 | 4840509.125 | 8212183.166 | 5445436.63 | 3068179.577 | Met-loss+Acetyl [N-Term] |
| Q9NQ66 | 1-phosphatidylinositol 4,5-bisphosphate phosphodiesterase beta-1 [OS=Hc | -3.09 | 0.967514922 | | 125194.0684 | 3874630.219 | | 1825087.672 | | | 898521.2394 | 3874630.219 | | 1895949.898 | | Met-loss+Acetyl [N-Term] |

| | | | | | | | | | | | | | | | | |
|---|---|---|---|---|---|---|---|---|---|---|---|---|---|---|---|---|
| Q6P2Q9 | Pre-mRNA-processing-splicing factor 8 [OS=Homo sapiens] | 0.21 | 0.967514922 | 61457.09766 | 51769.36719 | 390750.1563 | 65944.875 | 419502.375 | | 415518.5135 | 371550.1587 | 390750.1563 | 482047.2685 | 435790.2895 | | |
| O75475 | PC4 and SFRS1-interacting protein [OS=Homo sapiens] | -0.32 | 0.967787446 | | 542916.332 | 5114884.188 | 46904.28516 | 3363076.375 | | | 3896525.306 | 5114884.188 | 342863.377 | 3493653.706 | | |
| Q9NTX5 | Ethylmalonyl-CoA decarboxylase [OS=Homo sapiens] | 0 | 0.970551984 | | | 742215.4375 | | 511391.8125 | | | | 742215.4375 | | 531247.4954 | | |
| O14979 | Heterogeneous nuclear ribonucleoprotein D-like [OS=Homo sapiens] | 0 | 0.970551984 | | | 841066.9375 | | 268878.8438 | | | | 841066.9375 | | 279318.5359 | | |
| Q9GZT3 | SRA stem-loop-interacting RNA-binding protein, mitochondrial [OS=Homo s | 0.08 | 0.970551984 | 2423574 | 2898137 | 1187761 | 2367404 | 1694936.75 | 1624499.5 | 16386062.87 | 20800008.21 | 1187761 | 17305372.58 | 1760745.638 | 13746810.75 | |
| Q8TBC4 | NEDD8-activating enzyme E1 catalytic subunit [OS=Homo sapiens] | 0.25 | 0.970551984 | 104203.5 | 43012.72266 | 1711884.344 | 72626.82813 | 958214.8281 | | 704531.8616 | 308703.4824 | 1711884.344 | 530891.3561 | 995419.1973 | | |
| P50148 | Guanine nucleotide-binding protein G(q) subunit alpha [OS=Homo sapiens] | -1.6 | 0.970551984 | 360973.5 | 2949524.938 | 24847710.53 | | 17095873.59 | 248404.8027 | 2440583.396 | 21168820.83 | 24847710.53 | | 17759650.83 | 2102046.699 | |
| P63313 | Thymosin beta-10 [OS=Homo sapiens] | -2.23 | 0.970551984 | | 1006270.934 | 10255246.5 | | 8295859.125 | 26707.06055 | | 7222033.906 | 10255246.5 | | 8617960.387 | 226000.0123 | Met-loss+Acetyl [N-Term] |
| P42677 | Small ribosomal subunit protein eS27 [OS=Homo sapiens] | -0.05 | 0.970551984 | | 458858.0625 | 3564263.188 | 113795.1953 | 2944759.656 | 7825.765137 | | 3293236.816 | 3564263.188 | 831826.0224 | 3059095.108 | 66223.05042 | |
| Q02543 | Large ribosomal subunit protein eL20 [OS=Homo sapiens] | 0.32 | 0.970551984 | 384157.1094 | 741045.1563 | 5989615.438 | 597084.3828 | 6295738.703 | 295120.375 | 2597330.449 | 5318501.276 | 5989615.438 | 4364598.398 | 6540181.786 | 2497362.383 | |
| O00170 | AH receptor-interacting protein [OS=Homo sapiens] | 0 | 0.970551984 | | | 389717.8125 | | 354098.5625 | | | | 389717.8125 | | 367847.0595 | | |
| Q9UJC5 | SH3 domain-binding glutamic acid-rich-like protein 2 [OS=Homo sapiens] | 0 | 0.970551984 | | | 389614.2188 | | 187909.8281 | | | | 389614.2188 | | 195205.7564 | | |
| Q9Y4L1 | Hypoxia up-regulated protein 1 [OS=Homo sapiens] | 0.27 | 0.970551984 | 49163.05859 | 836105.3418 | 6884983.563 | 172514.5547 | 6647321.281 | 47460.74219 | 332397.0999 | 6000750.817 | 6884983.563 | 1261055.842 | 6905415.174 | 401621.4476 | |
| A8MWD9 | Putative small nuclear ribonucleoprotein G-like protein 15 [OS=Homo sapie | 0 | 0.970551984 | | | 303078.6563 | | 248798.4531 | | | | 303078.6563 | | 258458.4889 | | |
| P28838 | Cytosol aminopeptidase [OS=Homo sapiens] | -0.6 | 0.970551984 | 383943.2266 | 1253800.453 | 9253265.875 | 577723.6875 | 6483392.313 | 366181.2969 | 2595884.363 | 8998560.011 | 9253265.875 | 4223074.583 | 6735121.376 | 3098692.851 | |
| Q9Y2J8 | Protein-arginine deiminase type-2 [OS=Homo sapiens] | 0 | 0.970551984 | | | 317107.0313 | | 142296.5156 | | | | 317107.0313 | | 147821.4271 | | |
| Q04760 | Lactoylglutathione lyase [OS=Homo sapiens] | -0.06 | 0.970551984 | | 1642440.602 | 11706739 | 44284.375 | 11063791.03 | | | 11787840.94 | 11706739 | 323712.2218 | 11493362.1 | | |
| P05062 | Fructose-bisphosphate aldolase B [OS=Homo sapiens] | 0.12 | 0.970551984 | 718490.1875 | 685178.2813 | 353771.0625 | 722563.7813 | 579962.9063 | 719955.3438 | 4857794.886 | 4917543.192 | 353771.0625 | 5281834.215 | 602480.982 | 6092393.292 | |
| P61160 | Actin-related protein 2 [OS=Homo sapiens] | 0.11 | 0.970551984 | | 1538417.117 | 12491886.5 | 41579.24219 | 10628003.91 | 42592.78516 | | 11041261.56 | 12491886.5 | 303938.1017 | 11040654.77 | 360427.9083 | |
| Q9Y3F4 | Serine-threonine kinase receptor-associated protein [OS=Homo sapiens] | 0.28 | 0.970551984 | 50020.50781 | 215038.1738 | 2516781.875 | 129450.8672 | 1714448.719 | 32014.20313 | 338194.4128 | 1543334.832 | 2516781.875 | 946266.6649 | 1781015.193 | 270910.0197 | |
| P29692 | Elongation factor 1-delta [OS=Homo sapiens] | -0.08 | 0.970551984 | 176253.0391 | 196008.0273 | 1241359.844 | 198600.0156 | 1282978.563 | 43343.44922 | 1191667.091 | 1406754.952 | 1241359.844 | 1451736.698 | 1332792.453 | 366780.1643 | Met-loss+Acetyl [N-Term] |
| O43143 | ATP-dependent RNA helicase DHX15 [OS=Homo sapiens] | -0.72 | 0.970551984 | 49795.08203 | | 1363025.563 | 28040.07227 | 675291.0938 | | 336670.2831 | | 1363025.563 | 204968.775 | 701510.4534 | | |
| Q07955 | Serine/arginine-rich splicing factor 1 [OS=Homo sapiens] | 0.28 | 0.970551984 | 7400711.422 | 1711385.527 | 10252131.92 | 4667487.109 | 14051350.84 | 1343765.639 | 50037062.05 | 12282660.56 | 10252131.92 | 34118639.42 | 14596919.15 | 11371189.66 | |
| O14757 | Serine/threonine-protein kinase Chk1 [OS=Homo sapiens] | 0 | 0.970551984 | | | 520253.6875 | | 237296.7813 | | | | 520253.6875 | | 246510.2445 | | |
| P67870 | Casein kinase II subunit beta [OS=Homo sapiens] | -0.75 | 0.970551984 | | 112107.0313 | 1402973.719 | 12403.04688 | 919849.6875 | 24656.68945 | | 804595.2176 | 1402973.719 | 90664.43552 | 955564.4629 | 208649.3985 | |
| P45880 | Voltage-dependent anion-selective channel protein 2 [OS=Homo sapiens] | -0.07 | 0.970551984 | 1814067.805 | 10806970.34 | 84898260.88 | 2834564.797 | 62672439.9 | 1841978.598 | 12265121.3 | 77561920.56 | 84898260.88 | 20720248.81 | 65105807.16 | 15587158.5 | |
| Q96JE9 | Microtubule-associated protein 6 [OS=Homo sapiens] | -0.25 | 0.970551984 | 78236.71875 | 1554935.379 | 21470250.5 | 60837.80859 | 13638560.28 | 32900.14063 | 528967.4638 | 11159813.58 | 21470250.5 | 444715.3695 | 14168101.27 | 278406.9843 | |
| P02675 | Fibrinogen beta chain [OS=Homo sapiens] | 0.12 | 0.970551984 | 885799.1289 | 631017.1094 | 662242.3672 | 455740.6719 | 6890939.188 | 641553.9922 | 5988989.903 | 4528826.986 | 662242.3672 | 3331396.807 | 7158491.972 | 5428946.771 | |
| P63010 | AP-2 complex subunit beta [OS=Homo sapiens] | -0.09 | 0.970551984 | 31617.05273 | 9422063.232 | 91456907.03 | 43436.02344 | 65512353.91 | 60656.34473 | 213766.534 | 67622404.52 | 91456907.03 | 317510.8976 | 68055985.8 | 513285.0405 | |
| Q9P260 | RAB11-binding protein RELCH [OS=Homo sapiens] | 0 | 0.970551984 | | | 394081.0938 | | 547491.9375 | | | | 394081.0938 | | 568749.271 | | |
| Q08722 | Leukocyte surface antigen CD47 [OS=Homo sapiens] | -0.49 | 0.970551984 | | 574476.8789 | 3015734.25 | 20590.55859 | 2789357.344 | | | 4123036.211 | 3015734.25 | 150513.9334 | 2897659.029 | | |
| Q13114 | TNF receptor-associated factor 3 [OS=Homo sapiens] | 0 | 0.970551984 | | | 1120539.25 | | 285884.25 | | | | 1120539.25 | | 296984.2068 | | |
| Q02750 | Dual specificity mitogen-activated protein kinase kinase 1 [OS=Homo sapie | -0.07 | 0.970551984 | | 4655455.279 | 36556368.69 | 17071.30664 | 30688223.84 | | | 33412329.37 | 36556368.69 | 124788.7229 | 31879747.89 | | |
| P50991 | T-complex protein 1 subunit delta [OS=Homo sapiens] | -0.1 | 0.970551984 | 413600.418 | 2992734.32 | 35461159.03 | 869731.1074 | 26674754.72 | 196723.209 | 2796400.048 | 21478935.75 | 35461159.03 | 6357605.571 | 27710448.8 | 1664707.637 | |
| O43837 | Isocitrate dehydrogenase [NAD] subunit beta, mitochondrial [OS=Homo sap | -1.78 | 0.970551984 | 172710.6563 | 3804518.004 | 38577877.75 | 162007.6523 | 28227328.31 | 198008.3125 | 1167716.633 | 27305129.37 | 38577877.75 | 1184251.942 | 29323303.78 | 1675582.417 | |
| P05141 | ADP/ATP translocase 2 [OS=Homo sapiens] | 0.16 | 0.970551984 | 8057818.137 | 51324351.83 | 490127738.8 | 9941574.141 | 329478862.9 | 8021235.223 | 54479836.21 | 368356271.4 | 490127738.8 | 72671434.42 | 342271456.9 | 67877153.9 | Met-loss+Acetyl [N-Term] |
| P08865 | Small ribosomal subunit protein uS2 [OS=Homo sapiens] | 0.01 | 0.970551984 | 199525.0234 | 1453669.984 | 16267261.84 | 339901.6875 | 12904706.13 | 207205.7344 | 1349011.657 | 10433029.08 | 16267261.84 | 2484630.989 | 13405753.95 | 1753412.676 | Met-loss+Acetyl [N-Term] |
| P22234 | Bifunctional phosphoribosylaminoimidazole carboxylase/phosphoribosylan | 0.04 | 0.970551984 | 37674.16797 | 197029.4727 | 2508344.813 | 105121.0527 | 2152126.656 | 35223.39453 | 254719.3875 | 1414085.893 | 2508344.813 | 768419.325 | 2235686.744 | 298066.7821 | |
| P18124 | Large ribosomal subunit protein uL30 [OS=Homo sapiens] | -0.38 | 0.970551984 | 608006.6816 | 1929738.871 | 11830776.69 | 949787.6758 | 12921262.73 | 439381.8555 | 4110803.181 | 13849788.45 | 11830776.69 | 6942807.227 | 13422953.39 | 3718129.315 | |
| P11177 | Pyruvate dehydrogenase E1 component subunit beta, mitochondrial [OS=H | 0.66 | 0.970551984 | | 1646907.555 | 24752886.5 | 50523.91016 | 15156859.81 | 58670.33203 | | 11819900.39 | 24752886.5 | 369322.2996 | 15745351.44 | 496479.0392 | |
| P68366 | Tubulin alpha-4A chain [OS=Homo sapiens] | 0.17 | 0.970551984 | 436512.4922 | 3964701.582 | 140109254.6 | 391368.5313 | 61099149.41 | 552287.5 | 2951311.22 | 28454771.27 | 140109254.6 | 2860845.994 | 63471430.91 | 4673557.45 | |
| Q9Y4I1 | Unconventional myosin-Va [OS=Homo sapiens] | -0.05 | 0.970551984 | 35250.91016 | 3185769.148 | 26531666.16 | 34370.90234 | 20259183.13 | 35796.64844 | 238335.4624 | 22864351.97 | 26531666.16 | 251246.2051 | 21045781.4 | 302917.7611 | Met-loss+Acetyl [N-Term] |
| P29401 | Transketolase [OS=Homo sapiens] | 0 | 0.970551984 | 357224.0781 | 7354757.086 | 54701730.13 | 801190.4043 | 49545300.88 | 298350.8848 | 2415233.123 | 52785291.98 | 54701730.13 | 5856583.183 | 51468983.97 | 2524699.545 | |
| P04075 | Fructose-bisphosphate aldolase A [OS=Homo sapiens] | 0.01 | 0.970551984 | 872483.2852 | 38104781.22 | 343786694.9 | 3046355.941 | 248393227.7 | 2393583.699 | 5898959.95 | 273479052.9 | 343786694.9 | 22268410.71 | 258037529.9 | 20254941.37 | |
| P55084 | Trifunctional enzyme subunit beta, mitochondrial [OS=Homo sapiens] | -0.44 | 0.970551984 | 299610.793 | 1904313.188 | 10038409.5 | 272428.7031 | 9441447 | 312381.2422 | 2025703.069 | 13667307.63 | 10038409.5 | 1991413.468 | 9808027.718 | 2643426.986 | |
| Q99426 | Tubulin-folding cofactor B [OS=Homo sapiens] | 0.03 | 0.970551984 | 23947.14648 | 107472.8203 | 1073803.125 | 22611.90625 | 1400715.125 | | 161909.4147 | 771335.3594 | 1073803.125 | 165289.6854 | 1455100.343 | | |
| Q9UMS4 | Pre-mRNA-processing factor 19 [OS=Homo sapiens] | -0.76 | 0.970551984 | 39888.90234 | 131897.3945 | 1537584.938 | 66752.47266 | 1426915.969 | | 269693.4616 | 946631.194 | 1537584.938 | 487950.6877 | 1482318.481 | | |
| O43181 | NADH dehydrogenase [ubiquinone] iron-sulfur protein 4, mitochondrial [OS | 0 | 0.970551984 | | | 632048.1875 | | 499857.8438 | | | | 632048.1875 | | 519265.7001 | | |
| P46782 | Small ribosomal subunit protein uS7 [OS=Homo sapiens] | -0.64 | 0.970551984 | 43659.23828 | 252727.4727 | 2398209.313 | 25824.11914 | 1839900.641 | 19187.32422 | 295185.137 | 1813831.957 | 2398209.313 | 188770.4858 | 1911338.006 | 162366.6334 | Met-loss+Acetyl [N-Term] |
| P46781 | Small ribosomal subunit protein uS4 [OS=Homo sapiens] | -0.36 | 0.970551984 | 504880.3281 | 2420981.367 | 14710043.25 | 641922.8477 | 13495867.81 | 289376.2969 | 3413554.031 | 17375449.23 | 14710043.25 | 4692360.935 | 14019868.52 | 2448754.948 | |
| Q92734 | Protein TFG [OS=Homo sapiens] | 0 | 0.970551984 | | 533732.8711 | 3738979.563 | 32768.42578 | 3524331.375 | 47525.57422 | | 3830615.357 | 3738979.563 | 239532.3388 | 3661169.714 | 402170.068 | |
| O75436 | Vacuolar protein sorting-associated protein 26A [OS=Homo sapiens] | 0 | 0.970551984 | | | 850830.625 | | 709762.6875 | | | | 850830.625 | | 737320.4671 | | |
| Q15417 | Calponin-3 [OS=Homo sapiens] | 0 | 0.970551984 | | | 496910.625 | | 302215.5 | | | | 496910.625 | | 313949.5462 | | |
| P55263 | Adenosine kinase [OS=Homo sapiens] | 0 | 0.970551984 | | | 503623.9375 | | 249978.9219 | | | | 503623.9375 | | 259684.7914 | | |
| Q13263 | Transcription intermediary factor 1-beta [OS=Homo sapiens] | 0.12 | 0.970551984 | 81350.63281 | 531111.4102 | 3806855.313 | 82028.28906 | 3615493.875 | 29812.95508 | 550020.9953 | 3811801.06 | 3806855.313 | 599614.643 | 3755871.758 | 252282.6577 | |
| Q9NSE4 | Isoleucine--tRNA ligase, mitochondrial [OS=Homo sapiens] | 0 | 0.970551984 | | | 771616.1875 | | 645323.4375 | | | | 771616.1875 | | 670379.2504 | | |
| Q9Y277 | Voltage-dependent anion-selective channel protein 3 [OS=Homo sapiens] | 0.16 | 0.970551984 | 228726 | 9039130.602 | 92924374.06 | 460329.2988 | 69881702.66 | 663451.25 | 1546442.822 | 64874086.59 | 92924374.06 | 3364938.99 | 72594982.17 | 5614245.356 | Met-loss+Acetyl [N-Term] |
| Q00610 | Clathrin heavy chain 1 [OS=Homo sapiens] | 0.21 | 0.970551984 | 4676797.219 | 47292774.3 | 531942616.6 | 6395976.422 | 349600345 | 3970806.552 | 31620364.49 | 339421529.6 | 531942616.6 | 46753640.27 | 363174190.8 | 33601688.51 | Met-loss+Acetyl [N-Term] |
| P61586 | Transforming protein RhoA [OS=Homo sapiens] | 0.19 | 0.970551984 | 52762.125 | 1145132.648 | 22330060.94 | 95962.33203 | 12213266.41 | 166374.1445 | 356730.8022 | 8218648.216 | 22330060.94 | 701470.4332 | 12687467.87 | 1407888.324 | |
| O60493 | Sorting nexin-3 [OS=Homo sapiens] | -0.48 | 0.970551984 | | 603636.8555 | 3594789.344 | 58488.55078 | 2661378.375 | 20388.49609 | | 4332318.157 | 3594789.344 | 427542.6429 | 2764711.052 | 172531.1687 | Met-loss+Acetyl [N-Term] |
| P06748 | Nucleophosmin [OS=Homo sapiens] | -0.1 | 0.970551984 | 2472522.789 | 3359579.391 | 19820097 | 6295008.164 | 22143371.19 | 2584622 | 16717011.27 | 24111792.82 | 19820097 | 46015577.26 | 23003126.36 | 21871542.27 | |
| P0C0L4 | Complement C4-A [OS=Homo sapiens] | 0 | 0.970551984 | | | 364126.9375 | | 710456.4063 | | | | 364126.9375 | | 738041.1207 | | |
| O14561 | Acyl carrier protein, mitochondrial [OS=Homo sapiens] | -0.74 | 0.970551984 | 81117.48438 | 525175.7656 | 2638980.75 | 62366.98828 | 2117952.688 | 63792.03125 | 548444.6518 | 3769200.778 | 2638980.75 | 455893.4465 | 2200185.911 | 539819.7911 | |
| Q6PML9 | Proton-coupled zinc antiporter SLC30A9, mitochondrial [OS=Homo sapiens | 0 | 0.970551984 | | | 460325.4063 | | 271765.5938 | | | | 460325.4063 | | 282317.369 | | |
| Q14344 | Guanine nucleotide-binding protein subunit alpha-13 [OS=Homo sapiens] | -0.37 | 0.970551984 | | 766437.7656 | 7163884.469 | 17962.27344 | 4673518.516 | | | 5500744.725 | 7163884.469 | 131301.5583 | 4854976.059 | | |
| P61457 | Pterin-4-alpha-carbinolamine dehydratase [OS=Homo sapiens] | -0.05 | 0.970551984 | 29321.58789 | 181690.8203 | 1339670.781 | 37143.26953 | 911240.4375 | 26000.19336 | 198246.6319 | 1303999.967 | 1339670.781 | 271511.7986 | 946620.9437 | 220018.3734 | |
| O94979 | Protein transport protein Sec31A [OS=Homo sapiens] | 0 | 0.970551984 | | | 281934.2188 | | 202569.2969 | | | | 281934.2188 | | 210434.4047 | | |
| P49753 | Acyl-coenzyme A thioesterase 2, mitochondrial [OS=Homo sapiens] | -0.7 | 0.970551984 | | 162982.4688 | 768905.8125 | 62366.79297 | 648064.8125 | 56083.42969 | | 1169729.619 | 768905.8125 | 455892.0188 | 673227.0641 | 474588.2002 | |
| Q12860 | Contactin-1 [OS=Homo sapiens] | -0.06 | 0.970551984 | | 3015542.666 | 42436821.69 | 28132.79102 | 24592386.53 | | | 21642631.87 | 42436821.69 | 205646.5353 | 25547229.02 | | |
| O43390 | Heterogeneous nuclear ribonucleoprotein R [OS=Homo sapiens] | -0.09 | 0.970551984 | 27136.92383 | 884870.1523 | 10097938.03 | 219802.0605 | 7120453.328 | 22538.24805 | 183475.8666 | 6350737.191 | 10097938.03 | 1606720.506 | 7396917.401 | 190722.7614 | |
| P14868 | Aspartate--tRNA ligase, cytoplasmic [OS=Homo sapiens] | -0.38 | 0.970551984 | 641997.8828 | 760416.5742 | 6407790.031 | 875044.0508 | 5981185.797 | 252400.0625 | 4340621.606 | 5457530.47 | 6407790.031 | 6396442.4 | 6213415.813 | 2135855.315 | |
| P62873 | Guanine nucleotide-binding protein G(I)/G(S)/G(T) subunit beta-1 [OS=Hom | -0.07 | 0.970551984 | 294925.793 | 14319834.54 | 181353654.1 | 337623.5078 | 120209142.8 | 774499.1641 | 1994027.244 | 102773842.6 | 181353654.1 | 2467977.833 | 124876473.4 | 6553953.038 | Met-loss+Acetyl [N-Term] |
| P61204 | ADP-ribosylation factor 3 [OS=Homo sapiens] | 0.31 | 0.970551984 | 302139.4063 | 9522786.883 | 99199278.06 | 141482.2852 | 64362769.38 | 199893.9336 | 2042799.314 | 68345300.9 | 99199278.06 | 1034214.548 | 66861766.64 | 1691538.886 | |
| P06899 | Histone H2B type 1-J [OS=Homo sapiens] | 0.46 | 0.970551984 | 327251.75 | 895040.9375 | 5045065.125 | 484759.6875 | 7101494.25 | 533024.1875 | 2212586.762 | 6423733.193 | 5045065.125 | 3543521.513 | 7377222.204 | 4510547.79 | |
| P31930 | Cytochrome b-c1 complex subunit 1, mitochondrial [OS=Homo sapiens] | -0.43 | 0.970551984 | 94270.59375 | 2625152.063 | 42459253.44 | 211858.8633 | 20525477.56 | 222803.0234 | 637374.3388 | 18840787.87 | 42459253.44 | 1548656.91 | 21322415.18 | 1885399.778 | |
| P14174 | Macrophage migration inhibitory factor [OS=Homo sapiens] | 0.05 | 0.970551984 | 130127.8281 | 908832.4375 | 14415429 | 172209.6875 | 7016802.438 | 163204.6406 | 879809.2291 | 6522715.164 | 14415429 | 1258827.308 | 7289242.083 | 1381067.404 | Met-loss [N-Term] |
| P50502 | Hsc70-interacting protein [OS=Homo sapiens] | -0.35 | 0.970551984 | 17413.21289 | 1730675.453 | 14372753.75 | 332663.8848 | 10763322.53 | 126279.001 | 117732.7373 | 12421104.88 | 14372753.75 | 2431723.723 | 11181227.38 | 1068595.914 | |
| P43487 | Ran-specific GTPase-activating protein [OS=Homo sapiens] | -0.01 | 0.970551984 | 42358.42578 | 518560.0156 | 5752446.563 | 136789.4297 | 4233334.891 | | 286390.1939 | 3721719.36 | 5752446.563 | 999910.4698 | 4397701.533 | | |
| P07477 | Serine protease 1 [OS=Homo sapiens] | 0.02 | 0.970551984 | 215112324.5 | 82929313.78 | 204194695.5 | 197921961.9 | 276534459.5 | 146736976.2 | 1454399194 | 595185944.4 | 204194695.5 | 1446780225 | 287271394.3 | 1241715028 | |
| Q8NBM8 | Prenylcysteine oxidase-like [OS=Homo sapiens] | 0 | 0.970551984 | | | 381681.9688 | | 312458.9375 | | | | 381681.9688 | | 324590.7031 | | |
| P30740 | Leukocyte elastase inhibitor [OS=Homo sapiens] | -1.02 | 0.970551984 | 59762.20703 | | 691299.5625 | | 8905879.063 | 40210.73828 | 404059.1628 | | 691299.5625 | | 9251665.417 | 340270.5936 | |
| P04908 | Histone H2A type 1-B/E [OS=Homo sapiens] | 0.29 | 0.970551984 | 10541766.68 | 34204929.85 | 191627555.1 | 12193349.56 | 202227996.1 | 11448672.13 | 71274098.29 | 245489713.4 | 191627555.1 | 89131579.19 | 210079852.3 | 96880749.43 | |

| | | | | | | | | | | | | | | | | |
|---|---|---|---|---|---|---|---|---|---|---|---|---|---|---|---|---|
| Q6P5R6 | Ribosomal protein eL22-like [OS=Homo sapiens] | 0.05 | 0.970551984 | | 245342.6563 | 1712823.5 | 52387.32031 | 1757869.375 | | | 1760830.928 | 1712823.5 | 382943.5519 | 1826121.733 | | |
| P12532 | Creatine kinase U-type, mitochondrial [OS=Homo sapiens] | 0.19 | 0.970551984 | 69087.33594 | 4344770.422 | 83875099.91 | 135884.707 | 45516240.28 | 195202.9531 | 467107.4331 | 31182535.69 | 83875099.91 | 993297.0812 | 47283488.04 | 1651842.955 | |
| Q8N157 | Jouberin [OS=Homo sapiens] | -0.1 | 0.970551984 | 20327.31641 | 2121087.354 | 16606658.34 | 40727.69531 | 14686321 | | 137435.3267 | 15223101.72 | 16606658.34 | 297713.4201 | 15256543.14 | | |
| O76070 | Gamma-synuclein [OS=Homo sapiens] | -0.02 | 0.970551984 | 68251.03125 | 353475.8906 | 2802392.438 | 72509.5625 | 2373596.781 | 17807.97852 | 461453.0808 | 2536906.097 | 2802392.438 | 530034.1618 | 2465755.834 | 150694.3588 | |
| P15924 | Desmoplakin [OS=Homo sapiens] | 0.39 | 0.970551984 | 4316528.287 | 3589201.898 | 5020829.281 | 5118802.223 | 29144654.3 | 7835146.031 | 29184544.76 | 25759799.81 | 5020829.281 | 37417686.05 | 30276246.55 | 66302433.25 | |
| P05166 | Propionyl-CoA carboxylase beta chain, mitochondrial [OS=Homo sapiens] | 0.08 | 0.970551984 | 60099.33594 | 127979.1445 | 1276872.313 | 58783.76953 | 726662.375 | 86322.125 | 406338.5302 | 918509.807 | 1276872.313 | 429700.6482 | 754876.3146 | 730473.5494 | |
| Q92506 | (3R)-3-hydroxyacyl-CoA dehydrogenase [OS=Homo sapiens] | -0.84 | 0.970551984 | | 83681.80469 | 932879.375 | | 459420.7188 | 27443.53516 | | 600586.5921 | 932879.375 | | 477258.5329 | 232232.1946 | |
| Q02790 | Peptidyl-prolyl cis-trans isomerase FKBP4 [OS=Homo sapiens] | 0.07 | 0.970551984 | | 1437754.344 | 11502436.94 | 40607.26563 | 8771976.219 | 12958.03711 | | 10318802.1 | 11502436.94 | 296833.097 | 9112563.561 | 109653.2709 | |
| O00567 | Nucleolar protein 56 [OS=Homo sapiens] | -0.26 | 0.970551984 | 106169.6914 | 445230.6953 | 3627576.844 | 349293.9414 | 3045195.656 | | 717825.5081 | 3195432.83 | 3627576.844 | 2553286.974 | 3163430.712 | | |
| P07996 | Thrombospondin-1 [OS=Homo sapiens] | -0.1 | 0.970551984 | 89915.79688 | 239242.125 | 2277090.5 | 302772.3125 | 1541867.625 | 120293.3711 | 607931.0557 | 1717047.249 | 2277090.5 | 2213220.758 | 1601733.336 | 1017944.423 | |
| P14136 | Glial fibrillary acidic protein [OS=Homo sapiens] | -1.4 | 0.970551984 | | 7780626.039 | 106489328.8 | | 44544859.19 | 64384.64063 | | 55841765.06 | 106489328.8 | | 46274391.37 | 544834.5596 | |
| Q92696 | Geranylgeranyl transferase type-2 subunit alpha [OS=Homo sapiens] | 0 | 0.970551984 | | | 375533.375 | | 242749.7969 | | | | 375533.375 | | 252174.983 | | |
| Q12931 | Heat shock protein 75 kDa, mitochondrial [OS=Homo sapiens] | -0.12 | 0.970551984 | 139537.7344 | 172746.7656 | 921951.5938 | 207998.2344 | 1187161.219 | 99238.82422 | 943430.6887 | 1239808.243 | 921951.5938 | 1520436.285 | 1233254.833 | 839777.0115 | |
| P84098 | Large ribosomal subunit protein eL19 [OS=Homo sapiens] | -0.01 | 0.970551984 | 3227406.961 | 1093755 | 6837473.125 | 538538.75 | 7926110.063 | 378155.6367 | 21820870.07 | 7849909.434 | 6837473.125 | 3936638.494 | 8233855.17 | 3200021.897 | |
| P12814 | Alpha-actinin-1 [OS=Homo sapiens] | 0.25 | 0.970551984 | 403648.1738 | 725612.1094 | 5056997.031 | 267797.3906 | 7773039.875 | 297132.0039 | 2729111.78 | 5207737.878 | 5056997.031 | 1957559.259 | 8074841.764 | 2514385.154 | |
| Q92796 | Disks large homolog 3 [OS=Homo sapiens] | 0 | 0.970551984 | | | 413149.6875 | | 462214.9063 | | | | 413149.6875 | | 480161.2096 | | |
| Q1KMD3 | Heterogeneous nuclear ribonucleoprotein U-like protein 2 [OS=Homo sapiei | 0.09 | 0.970551984 | 34549.33984 | 114553.6465 | 2096676.484 | 34124.65234 | 1495958.5 | | 233592.0647 | 822154.6418 | 2096676.484 | 249446.1541 | 1554041.709 | | |
| Q13367 | AP-3 complex subunit beta-2 [OS=Homo sapiens] | -0.24 | 0.970551984 | 391554.1875 | 603635.6875 | 8047776.188 | 1042418.625 | 5797770.125 | 306957.4688 | 2647342.946 | 4332309.775 | 8047776.188 | 7619925.746 | 6022878.706 | 2597530.027 | |
| P18669 | Phosphoglycerate mutase 1 [OS=Homo sapiens] | 0.01 | 0.970551984 | 283937.9922 | 10615012.13 | 127671588 | 243438.8789 | 89682574.19 | 344679.418 | 1919737.458 | 76184231.22 | 127671588 | 1779502.147 | 93164657.23 | 2916740.034 | |
| P31153 | S-adenosylmethionine synthase isoform type-2 [OS=Homo sapiens] | 0.05 | 0.970551984 | 178278.7188 | 280712.1406 | 2242332.938 | 206797.2734 | 2048618.813 | 72151.14844 | 1205362.945 | 2014678.681 | 2242332.938 | 1511657.438 | 2128160.026 | 610556.1638 | |
| P23528 | Cofilin-1 [OS=Homo sapiens] | 0.25 | 0.970551984 | 4909409.211 | 13488179.29 | 127085966 | 5424749.438 | 86101495.78 | 5145125.934 | 33193080.95 | 96805030.23 | 127085966 | 39654114.87 | 89444537.18 | 43538993.08 | Met-loss+Acetyl [N-Term] |
| Q9H425 | Uncharacterized protein C1orf198 [OS=Homo sapiens] | 0 | 0.970551984 | | | 288881.0938 | | 219841.2344 | | | | 288881.0938 | | 228376.9554 | | |
| O43491 | Band 4.1-like protein 2 [OS=Homo sapiens] | -3.01 | 0.970551984 | | 120386.0313 | 2572803.875 | | 1817530.188 | | | 864013.8261 | 2572803.875 | | 1888098.981 | | |
| P26232 | Catenin alpha-2 [OS=Homo sapiens] | -0.07 | 0.970551984 | 75964.52344 | 1190735.379 | 12019733.19 | 97352.875 | 9489144.969 | 53929.19141 | 513604.8896 | 8545940.255 | 12019733.19 | 711635.0963 | 9857577.644 | 456358.6434 | |
| Q9UPA5 | Protein bassoon [OS=Homo sapiens] | -1.54 | 0.970551984 | | 323278.8359 | 3255537.719 | | 2153338.75 | 30355.02148 | | 2320181.013 | 3255537.719 | | 2236945.899 | 256869.7224 | |
| P63096 | Guanine nucleotide-binding protein G(i) subunit alpha-1 [OS=Homo sapiens | 0.35 | 0.970551984 | 22769.16406 | 1238647.211 | 13298625.75 | 50823.22266 | 10078099.33 | 20691.83984 | 153944.9399 | 8889804.779 | 13298625.75 | 371510.2297 | 10469399.19 | 175098.1187 | |
| Q9H7H0 | Methyltransferase-like protein 17, mitochondrial [OS=Homo sapiens] | -0.88 | 0.970551984 | 400216.4688 | 2161689.25 | 9221970 | 328859.3125 | 8140072 | 390217.3438 | 2705909.627 | 15514502.64 | 9221970 | 2403912.864 | 8456124.554 | 3302090.259 | |
| Q9NZH0 | G-protein coupled receptor family C group 5 member B [OS=Homo sapiens] | 0.22 | 0.970551984 | | 69425.04688 | 741064.3125 | 27222.5332 | 559943.75 | | | 498265.4529 | 741064.3125 | 198992.6855 | 581684.5469 | | |
| P08574 | Cytochrome c1, heme protein, mitochondrial [OS=Homo sapiens] | -0.36 | 0.970551984 | 350670.0625 | 2184947.055 | 16898832.78 | 398367.5547 | 13484087.66 | 426548.668 | 2370920.669 | 15681424.54 | 16898832.78 | 2912007.819 | 14007630.98 | 3609532.544 | Met-loss+Acetyl [N-Term] |
| O60884 | DnaJ homolog subfamily A member 2 [OS=Homo sapiens] | -2.19 | 0.970551984 | | 305666.6875 | 8464326.5 | | 6300634 | 15649.51172 | | 2193778.144 | 8464326.5 | | 6545267.152 | 132429.0195 | |
| Q16795 | NADH dehydrogenase [ubiquinone] 1 alpha subcomplex subunit 9, mitocho | 0.16 | 0.970551984 | 89088.08594 | 1277485.18 | 8550896.063 | 91802.57813 | 7337896.219 | 81791.4375 | 602334.8067 | 9168545.939 | 8550896.063 | 671063.2483 | 7622802.893 | 692134.0463 | |
| Q96GW7 | Brevican core protein [OS=Homo sapiens] | 0.22 | 0.970551984 | | 223376.9375 | 4336377.438 | 99426.46094 | 1975383.234 | | | 1603182.366 | 4336377.438 | 726792.7024 | 2052080.949 | | |
| Q92597 | Protein NDRG1 [OS=Homo sapiens] | -0.76 | 0.970551984 | 22377.2793 | 443908.4688 | 4741789 | 20892.57617 | 1594544.375 | 88668.58594 | 151295.3619 | 3185943.174 | 4741789 | 152721.6372 | 1656455.354 | 750329.7293 | |
| P16949 | Stathmin [OS=Homo sapiens] | -0.28 | 0.970551984 | 23992.29492 | 2925831.383 | 22491989.94 | 303584.5 | 17676763.47 | 133585.875 | 162214.6685 | 20998771.55 | 22491989.94 | 2219157.728 | 18363094.77 | 1130428.013 | Met-loss+Acetyl [N-Term] |
| P62304 | Small nuclear ribonucleoprotein E [OS=Homo sapiens] | 0.14 | 0.970551984 | | 150135.0781 | 1905616.875 | 47357.28516 | 1142470.5 | | | 1077523.546 | 1905616.875 | 346174.7399 | 1186828.918 | | |
| Q13363 | C-terminal-binding protein 1 [OS=Homo sapiens] | -0.26 | 0.970551984 | | 1178195.508 | 10632098.44 | 12602.00488 | 7737596.5 | | | 8455941.259 | 10632098.44 | 92118.78908 | 8038022.238 | | |
| Q01814 | Plasma membrane calcium-transporting ATPase 2 [OS=Homo sapiens] | -0.01 | 0.970551984 | | 737257.0977 | 11328280.38 | 56117.78125 | 6790333.297 | 26956.98047 | | 5291314.276 | 11328280.38 | 410212.6688 | 7053979.882 | 228114.8802 | |
| P11413 | Glucose-6-phosphate 1-dehydrogenase [OS=Homo sapiens] | 0.09 | 0.970551984 | 36273.50781 | 337551.5996 | 2889550.063 | 157456.7422 | 3616979.813 | 33525.61719 | 245249.3629 | 2422617.027 | 2889550.063 | 1150985.463 | 3757415.39 | 283699.8809 | Met-loss+Acetyl [N-Term] |
| P05026 | Sodium/potassium-transporting ATPase subunit beta-1 [OS=Homo sapiens | 0.31 | 0.970551984 | 938250.5801 | 20155105.19 | 212920141.8 | 1048246.504 | 152122734.8 | 1257859.516 | 6343620.203 | 144653739.1 | 212920141.8 | 7662526.677 | 158029166.4 | 10644236.4 | |
| Q15370 | Elongin-B [OS=Homo sapiens] | -0.41 | 0.970551984 | | 556339.2344 | 4956521.625 | 17089.03906 | 2934837.375 | | | 3992861.842 | 4956521.625 | 124918.3443 | 3048787.577 | | |
| Q9ULR3 | Protein phosphatase 1H [OS=Homo sapiens] | 0 | 0.970551984 | | | 783352.5313 | | 167526.75 | | | | 783352.5313 | | 174031.2695 | | |
| P10768 | S-formylglutathione hydrolase [OS=Homo sapiens] | -1.99 | 0.970551984 | | 27615.08789 | 934256.375 | | 1210699.75 | | | 198194.2381 | 934256.375 | | 1257707.289 | | |
| Q969L2 | Protein MAL2 [OS=Homo sapiens] | 0.14 | 0.970551984 | 109894.1484 | 179398 | 1233359 | 219062.6094 | 1361419.625 | 58482.33984 | 743006.9909 | 1287544.334 | 1233359 | 1601315.227 | 1414279.127 | 494888.2151 | |
| P00367 | Glutamate dehydrogenase 1, mitochondrial [OS=Homo sapiens] | -0.07 | 0.970551984 | 503430.9697 | 18349905.66 | 162241898.2 | 744010.4902 | 116645597.6 | 977546.0273 | 3403754.752 | 131697772.9 | 162241898.2 | 5438606.481 | 121174567.3 | 8272172.589 | |
| P49368 | T-complex protein 1 subunit gamma [OS=Homo sapiens] | 0.11 | 0.970551984 | 601636.2969 | 3151053.363 | 39743039.88 | 877275.9805 | 29013619.5 | 272231.3516 | 4067732.276 | 22615195.84 | 39743039.88 | 6412757.475 | 30140124.1 | 2303671.297 | |
| Q9Y266 | Nuclear migration protein nudC [OS=Homo sapiens] | -0.65 | 0.970551984 | 1142459 | 784736.1875 | 6816384.438 | 711485.7383 | 4348275.656 | 159152.7656 | 7724296.843 | 5632072.997 | 6816384.438 | 5200855.362 | 4517105.075 | 1346779.7 | |
| Q13561 | Dynactin subunit 2 [OS=Homo sapiens] | -1.26 | 0.970551984 | | 999605.1387 | 12019253.69 | | 8230056.313 | 39615.58594 | | 7174193.314 | 12019253.69 | | 8549602.665 | 335234.306 | |
| Q969P0 | Immunoglobulin superfamily member 8 [OS=Homo sapiens] | 0.1 | 0.970551984 | 1468055.5 | 1476529.563 | 8487091.313 | 1454160 | 4616037.5 | 1810857.625 | 9925692.269 | 10597092.9 | 8487091.313 | 10629694.21 | 4795263.242 | 15323807.16 | |
| Q96HD9 | N-acyl-aromatic-L-amino acid amidohydrolase (carboxylate-forming) [OS=H | 0.18 | 0.970551984 | 250129.375 | 245084.1406 | 240144.875 | 261397.2969 | | 208459.8438 | 1691153.505 | 1758975.554 | 240144.875 | 1910775.522 | | 1764025.179 | |
| P55209 | Nucleosome assembly protein 1-like 1 [OS=Homo sapiens] | -0.26 | 0.970551984 | 619022.3828 | 1053990.375 | 8599832.75 | 844347.5313 | 6003784.906 | 261879.9141 | 4185281.605 | 7564517.637 | 8599832.75 | 6172055.389 | 6236892.372 | 2216075.546 | |
| Q8WUY3 | Protein prune homolog 2 [OS=Homo sapiens] | 0 | 0.970551984 | | | 495135.2188 | | 490042.2813 | | | | 495135.2188 | | 509069.0312 | | |
| P08133 | Annexin A6 [OS=Homo sapiens] | -0.31 | 0.970551984 | 440023.8945 | 2639806.492 | 16692185.75 | 303794.9844 | 16008561.44 | 317720.1719 | 2975052.216 | 18945963.11 | 16692185.75 | 2220696.338 | 16630121.88 | 2688605.982 | |
| Q9UBG3 | Cornulin [OS=Homo sapiens] | 0 | 0.970551984 | 41100.63281 | | | | 4275002.469 | | 277886.1108 | | | | 4440986.928 | | |
| Q13423 | NAD(P) transhydrogenase, mitochondrial [OS=Homo sapiens] | 0.06 | 0.970551984 | 352452.4336 | 970763.625 | 2426479.422 | 293578.0313 | 4214418.875 | 439059.4219 | 2382971.485 | 6967196.984 | 2426479.422 | 2146011.924 | 4378051.07 | 3715400.823 | |
| P42765 | 3-ketoacyl-CoA thiolase, mitochondrial [OS=Homo sapiens] | -0.75 | 0.970551984 | 67291.90625 | 165227.8789 | 1302931 | 37095.61328 | 1214362.75 | | 454968.3262 | 1185844.988 | 1302931 | 271163.4385 | 1261512.511 | | |
| Q8N335 | Glycerol-3-phosphate dehydrogenase 1-like protein [OS=Homo sapiens] | -0.16 | 0.970551984 | 62010.53906 | 1130169.758 | 10293645 | 62866.53516 | 7717767.656 | 39047.65625 | 419260.3946 | 8111259.143 | 10293645 | 459545.0602 | 8017423.505 | 330428.3815 | |
| Q8NFZ4 | Neuroligin-2 [OS=Homo sapiens] | 0 | 0.970551984 | | | 763438.9375 | | 414780.1563 | | | | 763438.9375 | | 430884.7224 | | |
| P62826 | GTP-binding nuclear protein Ran [OS=Homo sapiens] | -0.17 | 0.970551984 | 219782.6328 | 3147209.227 | 38584333.25 | 599170.5273 | 26548630.88 | 335794.0918 | 1485975.687 | 22587606.36 | 38584333.25 | 4379847.806 | 27579427.97 | 2841550.785 | Met-loss+Acetyl [N-Term] |
| P60983 | Glia maturation factor beta [OS=Homo sapiens] | -0.4 | 0.970551984 | | 759699.3555 | 5705505 | 60449.19922 | 3869155 | | | 5452382.972 | 5705505 | 441874.6926 | 4019381.72 | | |
| O00154 | Cytosolic acyl coenzyme A thioester hydrolase [OS=Homo sapiens] | 0.2 | 0.970551984 | 106880.8438 | 3492732.641 | 49417683.97 | 473679.6484 | 33944561 | 103008.6934 | 722633.691 | 25067437.32 | 49417683.97 | 3462528.069 | 35262518.04 | 871678.3311 | |
| Q9H2X9 | Solute carrier family 12 member 5 [OS=Homo sapiens] | -1.05 | 0.970551984 | 32196.32422 | 2324197.363 | 26759624.63 | 50181.22656 | 16643559.14 | 27396.59961 | 217683.0552 | 16680827.8 | 26759624.63 | 366817.3333 | 17289774.48 | 231835.0175 | |
| P68371 | Tubulin beta-4B chain [OS=Homo sapiens] | 0.29 | 0.970551984 | 6082442.281 | 161850042.3 | 2210311222 | 6306711.838 | 1363634955 | 10473723.5 | 41124092.6 | 1161602163 | 2210311222 | 46101129.38 | 1416580471 | 88630556.56 | |
| Q99798 | Aconitate hydratase, mitochondrial [OS=Homo sapiens] | -0.06 | 0.970551984 | 667637.4688 | 12526093.62 | 124736826.1 | 961865.9531 | 81514795 | 649738.4883 | 4513973.799 | 89900115.16 | 124736826.1 | 7031097.646 | 84679749.7 | 5498205.468 | |
| P06744 | Glucose-6-phosphate isomerase [OS=Homo sapiens] | -0.04 | 0.970551984 | | 5475850.949 | 43902304.5 | 153059.5938 | 35320121.34 | 55925.32031 | | 39300331.45 | 43902304.5 | 1118842.959 | 36691486.92 | 473250.2499 | Met-loss+Acetyl [N-Term] |
| Q13838 | Spliceosome RNA helicase DDX39B [OS=Homo sapiens] | -0.39 | 0.970551984 | 186064.2559 | 1810784.484 | 19113226.98 | 424427.1406 | 14083535.02 | 279265.6563 | 1258001.858 | 12996049.58 | 19113226.98 | 3102499.532 | 14630352.93 | 2363196.864 | |
| Q8NBU5 | Outer mitochondrial transmembrane helix translocase [OS=Homo sapiens] | 0 | 0.970551984 | | | 841542.9375 | | 508529.625 | | | | 841542.9375 | | 528274.1785 | | |
| P55039 | Developmentally-regulated GTP-binding protein 2 [OS=Homo sapiens] | 0 | 0.970551984 | | | 796889.9375 | | 297551.9375 | | | | 796889.9375 | | 309104.9127 | | |
| O95834 | Echinoderm microtubule-associated protein-like 2 [OS=Homo sapiens] | 0 | 0.970551984 | | | 704945.9375 | | 432012.2188 | | | | 704945.9375 | | 448785.8499 | | |
| Q5XKP0 | MICOS complex subunit MIC13 [OS=Homo sapiens] | 0 | 0.970551984 | | | 812662.4375 | | 476725.3438 | | | | 812662.4375 | | 495235.0403 | | |
| P52907 | F-actin-capping protein subunit alpha-1 [OS=Homo sapiens] | -2.63 | 0.970551984 | | 65371.75781 | 2307700.5 | | 1185827.125 | | | 469174.8868 | 2307700.5 | | 1231868.94 | | |
| Q9BTV4 | Transmembrane protein 43 [OS=Homo sapiens] | 0.25 | 0.970551984 | | 98966.95313 | 865333.6875 | 31292.30469 | 1047953.891 | | | 710288.5189 | 865333.6875 | 228742.1123 | 1088642.536 | | |
| P17987 | T-complex protein 1 subunit alpha [OS=Homo sapiens] | 0.35 | 0.970551984 | 1162242.299 | 2811220.209 | 33941029.22 | 1680180.441 | 26511769.44 | 709194.9697 | 7858054.005 | 20176204.03 | 33941029.22 | 12281870.16 | 27541135.33 | 6001337.047 | |
| P36578 | Large ribosomal subunit protein uL4 [OS=Homo sapiens] | 0.23 | 0.970551984 | 19531396.67 | 2838229.457 | 24622898.48 | 17733159.38 | 30930819.13 | 7659671.773 | 132054021.8 | 20370050.14 | 24622898.48 | 129626768.4 | 32131762.36 | 64817538.11 | Met-loss+Acetyl [N-Term] |
| Q9Y3Y2 | Chromatin target of PRMT1 protein [OS=Homo sapiens] | 0 | 0.970551984 | | | 342706.5938 | | 483409.9063 | | | | 342706.5938 | | 502179.1426 | | |
| O60641 | Clathrin coat assembly protein AP180 [OS=Homo sapiens] | -1.49 | 0.970551984 | | 6061922.211 | 54063900.5 | | 41781097.66 | 54237.19141 | | 43506580.86 | 54063900.5 | | 43403321.96 | 458964.9955 | Met-loss+Acetyl [N-Term] |
| P20020 | Plasma membrane calcium-transporting ATPase 1 [OS=Homo sapiens] | -0.31 | 0.970551984 | 35124.07813 | 809685.8867 | 8004231.313 | 87164.15039 | 5653908.844 | | 237477.9364 | 5811137.668 | 8004231.313 | 637157.0286 | 5873431.758 | | |
| P62979 | Ubiquitin-ribosomal protein eS31 fusion protein [OS=Homo sapiens] | 0.33 | 0.970551984 | 16171066.73 | 28618426.34 | 193112831.6 | 43646117.69 | 197134845 | 30027988.25 | 109334444.1 | 205395225.6 | 193112831.6 | 319046655.4 | 204788950.7 | 254102307.6 | |
| P54105 | Methylosome subunit pICln [OS=Homo sapiens] | 0 | 0.970551984 | | | 325628.0625 | 39448.46094 | | | | | 325628.0625 | 288362.4063 | | | |
| O94903 | Pyridoxal phosphate homeostasis protein [OS=Homo sapiens] | 0 | 0.970551984 | | | 651427.9375 | | 445235.2188 | | | | 651427.9375 | | 462522.256 | | |
| P62244 | Small ribosomal subunit protein uS8 [OS=Homo sapiens] | -0.03 | 0.970551984 | 129888.3438 | 678441.5 | 4433551.625 | 118061.4141 | 4663140.75 | 91267.96875 | 878190.0476 | 4869193.129 | 4433551.625 | 863011.4495 | 4844195.358 | 772326.1803 | |
| Q13435 | Splicing factor 3B subunit 2 [OS=Homo sapiens] | -0.03 | 0.970551984 | 27378.19336 | 78713.54688 | 830797.1563 | 44166.74609 | 770156.375 | 27075.06641 | 185107.1177 | 564929.2704 | 830797.1563 | 322852.372 | 800059.0453 | 229114.1449 | |
| P51665 | 26S proteasome non-ATPase regulatory subunit 7 [OS=Homo sapiens] | 0 | 0.970551984 | | | 3456991.594 | | 1479755.203 | | | | 3456991.594 | | 1537209.291 | | |

| | | | | | | | | | | | | | | | | |
|---|---|---|---|---|---|---|---|---|---|---|---|---|---|---|---|---|
| Q9Y6E0 | Serine/threonine-protein kinase 24 [OS=Homo sapiens] | 0.01 | 0.970551984 | | 183097.7734 | 1342224.938 | 42813.01563 | 1261280.125 | | | 1314097.708 | 1342224.938 | 312956.8028 | 1310251.535 | | |
| P61088 | Ubiquitin-conjugating enzyme E2 N [OS=Homo sapiens] | 0.3 | 0.970551984 | | 3477748.512 | 32714700.94 | 101012.8359 | 25895410.13 | | | 24959895.82 | 32714700.94 | 738388.8687 | 26900844.78 | | |
| O43681 | ATPase GET3 [OS=Homo sapiens] | -2.31 | 0.970551984 | | 30946.80078 | 4050232.563 | | 2570905.656 | | | 222106.0323 | 4050232.563 | | 2670725.572 | | |
| Q15233 | Non-POU domain-containing octamer-binding protein [OS=Homo sapiens] | -0.42 | 0.970551984 | 49206.91602 | 1371978.781 | 9508958.266 | 254919.5957 | 8182172.016 | 152634.0332 | 332693.6249 | 9846729.092 | 9508958.266 | 1863424.486 | 8499859.177 | 1291616.998 | |
| P62269 | Small ribosomal subunit protein uS13 [OS=Homo sapiens] | -0.32 | 0.970551984 | 141623.5195 | 2590165.324 | 17153111.63 | 278085.1875 | 13198162.13 | 153505.5859 | 957532.9223 | 18589687.1 | 17153111.63 | 2032761.531 | 13710603.89 | 1298992.236 | Met-loss+Acetyl [N-Term] |
| P63261 | Actin, cytoplasmic 2 [OS=Homo sapiens] | 0 | 0.970551984 | | | 8706575 | | 3449690.75 | | | | 8706575 | | 3583631.036 | | Met-loss+Acetyl [N-Term] |
| P11940 | Polyadenylate-binding protein 1 [OS=Homo sapiens] | -0.36 | 0.970551984 | 277742.5859 | 1448476.43 | 11331637.25 | 309446.4531 | 7328878.313 | 78562.60938 | 1877849.603 | 10395754.8 | 11331637.25 | 2262007.737 | 7613434.851 | 664811.1144 | |
| P22392 | Nucleoside diphosphate kinase B [OS=Homo sapiens] | -0.08 | 0.970551984 | 1584149.781 | 13346196.52 | 94656458.16 | 2140525.418 | 77709560.09 | 1567521.145 | 10710619.07 | 95786016.06 | 94656458.16 | 15646923.75 | 80726769.89 | 13264649.52 | |
| Q4V328 | GRIP1-associated protein 1 [OS=Homo sapiens] | 0 | 0.970551984 | | | 257357.4688 | | 564559.2969 | | | | 257357.4688 | | 586479.3005 | | |
| P21291 | Cysteine and glycine-rich protein 1 [OS=Homo sapiens] | -2.18 | 0.970551984 | | 492883.6875 | 11384121.88 | | 5264479.938 | 23768.81055 | | 3537439.653 | 11384121.88 | | 5468882.593 | 201136.0054 | |
| P17600 | Synapsin-1 [OS=Homo sapiens] | 0 | 0.970551984 | 3382811.477 | 12602226.55 | 157353242 | 3047280.75 | 94704256 | 2035274.242 | 22871577.89 | 90446523.27 | 157353242 | 22275170.92 | 98381314.63 | 17222861.46 | |
| P49419 | Alpha-aminoadipic semialdehyde dehydrogenase [OS=Homo sapiens] | 0.11 | 0.970551984 | | 350667.4375 | 6052206.625 | 43714.0625 | 2632268.313 | 59862.94141 | | 2516749.753 | 6052206.625 | 319543.3219 | 2734470.741 | 506571.117 | |
| Q9H0U4 | Ras-related protein Rab-1B [OS=Homo sapiens] | -0.54 | 0.970551984 | | 670866.1563 | 3603323.875 | 267034.25 | 2893243.594 | 205972.8281 | | 4814824.681 | 3603323.875 | 1951980.814 | 3005578.844 | 1742979.599 | |
| Q96EB6 | NAD-dependent protein deacetylase sirtuin-1 [OS=Homo sapiens] | -0.76 | 0.970551984 | 106143.0313 | | | 58104.11719 | | | 717645.2557 | | | 424732.4903 | | | |
| P63167 | Dynein light chain 1, cytoplasmic [OS=Homo sapiens] | -2.42 | 0.970551984 | | 59681.85938 | 2058661 | | 1433590.625 | | | 428338.3307 | 2058661 | | 1489252.292 | | |
| Q5JWF2 | Guanine nucleotide-binding protein G(s) subunit alpha isoforms XLas [OS=H | 0.19 | 0.970551984 | | 1230243.449 | 12475558.38 | 134942.0938 | 9053637.023 | 96960.54688 | | 8829490.752 | 12475558.38 | 986406.7177 | 9405160.34 | 820497.8136 | |
| P13645 | Keratin, type I cytoskeletal 10 [OS=Homo sapiens] | 0.06 | 0.970551984 | 265730742.6 | 67746490.98 | 196785349.1 | 236342070.9 | 502106710.9 | 180135018.5 | 1796636147 | 486218411.4 | 196785349.1 | 1727625531 | 521601883.5 | 1524335347 | |
| P52209 | 6-phosphogluconate dehydrogenase, decarboxylating [OS=Homo sapiens] | 0.27 | 0.970551984 | 131376.4375 | 421376.9492 | 4771996 | 179472.3047 | 5270495.875 | 138239.0156 | 888251.2208 | 3024233.845 | 4771996 | 1311915.964 | 5475132.11 | 1169803.737 | |
| P05787 | Keratin, type II cytoskeletal 8 [OS=Homo sapiens] | 0.04 | 0.970551984 | 3450894.625 | 2194625.797 | 1204753.422 | 3211653.125 | 2746182.594 | 3437438.203 | 23331895.9 | 15750889.14 | 1204753.422 | 23476708.63 | 2852807.943 | 29088228.37 | |
| Q9BTV5 | Fibronectin type III and SPRY domain-containing protein 1 [OS=Homo sapier | 0 | 0.970551984 | | | 370174.9063 | | 450378.7188 | | | | 370174.9063 | | 467865.4614 | | |
| P27635 | Large ribosomal subunit protein uL16 [OS=Homo sapiens] | -0.27 | 0.970551984 | 511332.5313 | 1258126.293 | 9771921.563 | 595145.9609 | 8247939.375 | 208594.25 | 3457178.119 | 9029606.681 | 9771921.563 | 4350428.821 | 8568180.069 | 1765162.549 | |
| P37837 | Transaldolase [OS=Homo sapiens] | 0.25 | 0.970551984 | 285928.207 | 973670.6211 | 8171098.5 | 277442.707 | 11595160.13 | 209281.0586 | 1933193.53 | 6988060.575 | 8171098.5 | 2028065.094 | 12045362.53 | 1770974.448 | |
| Q09028 | Histone-binding protein RBBP4 [OS=Homo sapiens] | 0 | 0.970551984 | | | 327545.3438 | | 301672.9375 | | | | 327545.3438 | | 313385.9177 | | Met-loss+Acetyl [N-Term] |
| O75533 | Splicing factor 3B subunit 1 [OS=Homo sapiens] | 0 | 0.970551984 | | | 723632.1094 | | 843395.2813 | | | | 723632.1094 | | 876141.5804 | | |
| P52272 | Heterogeneous nuclear ribonucleoprotein M [OS=Homo sapiens] | -0.14 | 0.970551984 | 29608.94922 | 1000631.172 | 8352529.813 | 394553.9004 | 6593236.531 | 51386.2793 | 200189.5149 | 7181557.182 | 8352529.813 | 2884130.571 | 6849230.489 | 434840.0578 | |
| P62330 | ADP-ribosylation factor 6 [OS=Homo sapiens] | -1.25 | 0.970551984 | | 117688.0078 | 3771774.375 | | 1474280.438 | 24130.6875 | | 844650.0384 | 3771774.375 | | 1531521.958 | 204198.2742 | |
| P18085 | ADP-ribosylation factor 4 [OS=Homo sapiens] | -1.09 | 0.970551984 | | 279960.625 | 1234537.5 | 35255.83594 | 909641.9375 | 39137.88672 | | 2009285.033 | 1234537.5 | 257714.9386 | 944960.3791 | 331191.9282 | |
| O00232 | 26S proteasome non-ATPase regulatory subunit 12 [OS=Homo sapiens] | 0.21 | 0.970551984 | | 143717.8672 | 1495727.5 | 32869.23438 | 1161026.125 | | | 1031467.048 | 1495727.5 | 240269.2347 | 1206104.998 | | |
| Q9Y3I0 | RNA-splicing ligase RtcB homolog [OS=Homo sapiens] | -0.08 | 0.970551984 | 549850.2656 | 492558.0703 | 6280384.938 | 311649.875 | 4239384.547 | 314369.5898 | 3717600.956 | 3535102.69 | 6280384.938 | 2278114.425 | 4403986.078 | 2660252.746 | |
| P61956 | Small ubiquitin-related modifier 2 [OS=Homo sapiens] | 0.13 | 0.970551984 | 252086.25 | 421910.4375 | 3900609.25 | 347006.9688 | 4220121.563 | 144029.4844 | 1704384.162 | 3028062.705 | 3900609.25 | 2536569.543 | 4383975.175 | 1218803.738 | |
| Q01844 | RNA-binding protein EWS [OS=Homo sapiens] | -0.04 | 0.970551984 | | 197021.0547 | 1370101.188 | 86304.53125 | 1321269.688 | | | 1414025.477 | 1370101.188 | 630873.34 | 1372570.298 | | |
| P41250 | Glycine--tRNA ligase [OS=Homo sapiens] | 0 | 0.970551984 | 14358.77637 | 268280.1914 | 2407379.625 | 60913.45898 | 2486316.516 | 30108.27734 | 97081.34031 | 1925454.243 | 2407379.625 | 445268.3627 | 2582852.11 | 254781.7285 | |
| P28074 | Proteasome subunit beta type-5 [OS=Homo sapiens] | -0.27 | 0.970551984 | 183720.4063 | 1358789.426 | 7456862.625 | 241591.5234 | 7354184.719 | 147819.4063 | 1242154.82 | 9752068.728 | 7456862.625 | 1765998.252 | 7639723.823 | 1250874.748 | |
| P04114 | Apolipoprotein B-100 [OS=Homo sapiens] | -0.11 | 0.970551984 | 774816.1563 | 532359.8203 | 302431.5781 | 658732.7344 | 949564.4375 | 610402.7461 | 5238621.246 | 3820760.934 | 302431.5781 | 4815238.717 | 986432.9401 | 5165339.251 | |
| Q08AH3 | Acyl-coenzyme A synthetase ACSM2A, mitochondrial [OS=Homo sapiens] | -0.02 | 0.970551984 | 353692.4688 | 673883.1875 | 303542.1875 | 678323.7188 | 318696.4688 | 308930.9688 | 2391355.506 | 4836477.996 | 303542.1875 | 4958445.911 | 331070.4174 | 2614230.144 | |
| Q9H3Z4 | DnaJ homolog subfamily C member 5 [OS=Homo sapiens] | 0.13 | 0.970551984 | 530827.2188 | 3295929.883 | 22738434.19 | 1679670.414 | 23971678 | 172625.9063 | 3588983.946 | 23654978.56 | 22738434.19 | 12278141.93 | 24902420.39 | 1460791.871 | |
| Q92598 | Heat shock protein 105 kDa [OS=Homo sapiens] | 0.24 | 0.970551984 | | 1698209.453 | 20812876.84 | 148384.3242 | 13281112.91 | | | 12188095.51 | 20812876.84 | 1084667.431 | 13796775.38 | | Met-loss+Acetyl [N-Term] |
| Q9H115 | Beta-soluble NSF attachment protein [OS=Homo sapiens] | -0.07 | 0.970551984 | 385116.0234 | 4873829.664 | 56136413.41 | 683373.623 | 36872409.47 | 979300.1621 | 2603813.777 | 34979608.28 | 56136413.41 | 4995359.963 | 38304045.36 | 8287016.397 | |
| Q01082 | Spectrin beta chain, non-erythrocytic 1 [OS=Homo sapiens] | -0.01 | 0.970551984 | 251799.71 | 22985893.87 | 257890208.3 | 312158.3467 | 168664753.8 | 423668.1943 | 1702446.832 | 164970386.6 | 257890208.3 | 2281831.277 | 175213458.3 | 3585157.451 | Met-loss+Acetyl [N-Term] |
| Q14677 | Clathrin interactor 1 [OS=Homo sapiens] | 0 | 0.970551984 | | | 436262.1563 | | 396829.4688 | | | | 436262.1563 | | 412237.068 | | |
| P68871 | Hemoglobin subunit beta [OS=Homo sapiens] | 0.1 | 0.970551984 | 4354420.461 | 6498440.984 | 66177259.2 | 4211501.641 | 38147739.97 | 4281554.953 | 29440738.11 | 46639487.99 | 66177259.2 | 30785453.19 | 39628892.8 | 36231298.1 | |
| Q15021 | Condensin complex subunit 1 [OS=Homo sapiens] | 0.09 | 0.970551984 | 1146403 | 135119.6719 | 283639.9063 | 632433.0625 | 995899.1875 | 340171.625 | 7750962.681 | 969757.5663 | 283639.9063 | 4622992.011 | 1034566.718 | 2878594.269 | |
| Q99832 | T-complex protein 1 subunit eta [OS=Homo sapiens] | -0.24 | 0.970551984 | 376492.3848 | 3229943.066 | 24404932.06 | 531426.9375 | 20713413.34 | 284467.8867 | 2545508.363 | 23181389.39 | 24404932.06 | 3884652.198 | 21517647.9 | 2407219.087 | |
| Q13243 | Serine/arginine-rich splicing factor 5 [OS=Homo sapiens] | 0 | 0.970551984 | | | 510016.8438 | | 367984.9375 | | | | 510016.8438 | | 382272.5973 | | |
| P61019 | Ras-related protein Rab-2A [OS=Homo sapiens] | 0.01 | 0.970551984 | 87344.58594 | 4556231.703 | 52645005.25 | 158287.6328 | 33321314.63 | 186467.0898 | 590546.8025 | 32700199.06 | 52645005.25 | 1157059.15 | 34615073.03 | 1577918.488 | Met-loss+Acetyl [N-Term] |
| Q93050 | V-type proton ATPase 116 kDa subunit a 1 [OS=Homo sapiens] | 0.05 | 0.970551984 | 55739.63574 | 6141709.68 | 71354205.06 | 227185.5859 | 48312762.56 | 125215.0215 | 376862.0952 | 44079217.7 | 71354205.06 | 1660692.982 | 50188590.2 | 1059592.326 | |
| Q9P2R7 | Succinate--CoA ligase [ADP-forming] subunit beta, mitochondrial [OS=Hom | -0.46 | 0.970551984 | 139615.0723 | 3047286.563 | 27085777.94 | 163857.9883 | 18631883.69 | 186437.4844 | 943953.5791 | 21870458.68 | 27085777.94 | 1197777.624 | 19355299.21 | 1577667.961 | |
| P06733 | Alpha-enolase [OS=Homo sapiens] | 0.08 | 0.970551984 | 2748652.207 | 29809466.13 | 299959306 | 3744753.75 | 212696296.5 | 4395484.604 | 18583954.06 | 213943350.6 | 299959306 | 27373595.24 | 220954602.8 | 37195391.56 | |
| Q9Y230 | RuvB-like 2 [OS=Homo sapiens] | -0.03 | 0.970551984 | 193080.6719 | 355511.9063 | 2067997.563 | 329251.2383 | 2154848.391 | 37199.21484 | 1305440.654 | 2551518.637 | 2067997.563 | 2406777.783 | 2238514.154 | 314786.5336 | |
| P63000 | Ras-related C3 botulinum toxin substrate 1 [OS=Homo sapiens] | 0.2 | 0.970551984 | 155195.1719 | 4295941.461 | 44352287.75 | 138398.0996 | 33494102.38 | 190584.3301 | 1049292.426 | 30832088.91 | 44352287.75 | 1011669.609 | 34794569.57 | 1612759.325 | |
| Q9NRW7 | Vacuolar protein sorting-associated protein 45 [OS=Homo sapiens] | -2.83 | 0.970551984 | | 90800.57031 | 1658424.5 | | 1225729.344 | | | 651678.1669 | 1658424.5 | | 1273320.433 | | |
| P04179 | Superoxide dismutase [Mn], mitochondrial [OS=Homo sapiens] | -0.29 | 0.970551984 | 77649.04492 | 3700254.313 | 24179130 | 66446.24414 | 20911452.84 | 82491.89063 | 524994.1333 | 26556826.01 | 24179130 | 485712.2027 | 21723376.63 | 698061.4082 | |
| P61201 | COP9 signalosome complex subunit 2 [OS=Homo sapiens] | -0.46 | 0.970551984 | 59999.99219 | 1524753.25 | 26916382.13 | 77892.75391 | 17528262.06 | 19862.70313 | 405666.8557 | 10943195.62 | 26916382.13 | 569384.4936 | 18208827.54 | 168081.8128 | |
| P16298 | Serine/threonine-protein phosphatase 2B catalytic subunit beta isoform [OS | 0.15 | 0.970551984 | 70490.14844 | 275346.9375 | 2502257.594 | 76827.58594 | 2299191.484 | 18595.82422 | 476592.0099 | 1976172.472 | 2502257.594 | 561598.2735 | 2388461.621 | 157361.2526 | |
| Q6H8Q1 | Actin-binding LIM protein 2 [OS=Homo sapiens] | 0 | 0.970551984 | | | 477985.0938 | | 236784.1406 | | | | 477985.0938 | | 245977.6997 | | |
| P00918 | Carbonic anhydrase 2 [OS=Homo sapiens] | 0 | 0.970551984 | | | 3377631.25 | | 1026460.969 | | | | 3377631.25 | | 1066315.114 | | |
| Q9P0M6 | Core histone macro-H2A.2 [OS=Homo sapiens] | 0.43 | 0.970551984 | | 426507.8984 | 2628753.313 | 49856.66016 | 2207237.813 | 97348.02344 | | 3061058.807 | 2628753.313 | 364444.801 | 2292937.687 | 823776.7108 | |
| P14550 | Aldo-keto reductase family 1 member A1 [OS=Homo sapiens] | -0.38 | 0.970551984 | 716734.8438 | 1414172.609 | 7544983.594 | 532076.2578 | 6092328.781 | 530955.2969 | 4845926.804 | 10149555.35 | 7544983.594 | 3889398.633 | 6328874.118 | 4493040.461 | |
| P07237 | Protein disulfide-isomerase [OS=Homo sapiens] | 0.04 | 0.970551984 | 2515550.16 | 1592245.805 | 14005572.5 | 1722995.43 | 14196674.16 | 913155.9258 | 17007924.28 | 11427591.52 | 14005572.5 | 12594841.38 | 14747884.9 | 7727291.818 | |
| Q02978 | Mitochondrial 2-oxoglutarate/malate carrier protein [OS=Homo sapiens] | 0.06 | 0.970551984 | 101574.6328 | 4681086.852 | 45426189.13 | 146402.6758 | 33920651.59 | 219713.4287 | 686757.7879 | 33596287.87 | 45426189.13 | 1070181.875 | 35237680.31 | 1859255.065 | |
| O14950 | Myosin regulatory light chain 12B [OS=Homo sapiens] | -0.13 | 0.970551984 | 164572.7734 | 586194.5117 | 5205360.625 | 73001.67969 | 4029504.25 | 115858.4141 | 1112695.47 | 4207133.981 | 5205360.625 | 533631.4655 | 4185956.811 | 980415.0088 | |
| P61970 | Nuclear transport factor 2 [OS=Homo sapiens] | 0.17 | 0.970551984 | | 196706.6211 | 1507585.625 | 28897.77148 | 1233909.813 | | | 1411768.779 | 1507585.625 | 211238.4292 | 1281818.522 | | |
| Q96CW1 | AP-2 complex subunit mu [OS=Homo sapiens] | 0.42 | 0.970551984 | 678725.9531 | 8836079.211 | 66761900.38 | 563553.8926 | 53529494.34 | 357004.542 | 4588944.319 | 63416781.23 | 66761900.38 | 4119495.481 | 55607870.73 | 3021037.479 | |
| P14927 | Cytochrome b-c1 complex subunit 7 [OS=Homo sapiens] | 0 | 0.970551984 | | | 572322.0625 | | 219714.3594 | | | | 572322.0625 | | 228245.1542 | | |
| P43034 | Platelet-activating factor acetylhydrolase IB subunit beta [OS=Homo sapien | -0.38 | 0.970551984 | 271089.0098 | 2259558.934 | 19880363.53 | 668294.8086 | 13196495.11 | 148014.7969 | 1832864.009 | 16216916.03 | 19880363.53 | 4885136.063 | 13708872.15 | 1252528.179 | |
| Q9GZP4 | PITH domain-containing protein 1 [OS=Homo sapiens] | -0.1 | 0.970551984 | 30949.4707 | 225500.2656 | 3082557 | | 1342683.125 | 78320.10938 | 209252.935 | 1618421.55 | 3082557 | | 1394815.149 | 662759.0352 | |
| Q15631 | Translin [OS=Homo sapiens] | -1.1 | 0.970551984 | | 154672.9688 | 2593777.031 | | 2554463.813 | 27235.34961 | | 1110092.111 | 2593777.031 | | 2653645.345 | 230470.4906 | |
| Q6PCB7 | Long-chain fatty acid transport protein 1 [OS=Homo sapiens] | 0 | 0.970551984 | | | 404204.0938 | | 200574.6094 | | | | 404204.0938 | | 208362.2699 | | |
| P68402 | Platelet-activating factor acetylhydrolase IB subunit alpha2 [OS=Homo sapi | -0.04 | 0.970551984 | | 1229790.445 | 13400823.19 | 18801.93945 | 8518752.234 | | | 8826239.531 | 13400823.19 | 137439.3925 | 8849507.712 | | Met-loss+Acetyl [N-Term] |
| P63244 | Small ribosomal subunit protein RACK1 [OS=Homo sapiens] | 0.3 | 0.970551984 | 4703124.086 | 1336453.914 | 17509845.22 | 4777648.086 | 18225034 | 2068757.766 | 31798363.47 | 9591766.152 | 17509845.22 | 34923899.84 | 18932652.87 | 17506205.13 | Met-loss+Acetyl [N-Term] |
| Q9Y2V2 | Calcium-regulated heat-stable protein 1 [OS=Homo sapiens] | 0 | 0.970551984 | | | 1374101.875 | | 434134.7813 | | | | 1374101.875 | | 450990.8246 | | |
| Q15149 | Plectin [OS=Homo sapiens] | -0.92 | 0.970551984 | 49696.47266 | 350610.7031 | 4536394.219 | 265823.1914 | 2511337.641 | 106954.7168 | 336003.5737 | 2516342.568 | 4536394.219 | 1943128.155 | 2608844.724 | 905070.3003 | |
| Q14019 | Coactosin-like protein [OS=Homo sapiens] | -0.28 | 0.970551984 | 18107.79492 | 1888620.891 | 14449929 | 61392.73633 | 9995286.813 | 48865.17773 | 122428.8864 | 13554683.59 | 14449929 | 448771.8091 | 10383371.33 | 413506.0371 | |
| P07437 | Tubulin beta chain [OS=Homo sapiens] | -0.05 | 0.970551984 | 490680.5234 | 6620738.387 | 77435870.69 | 474221.3672 | 57424273.41 | 417935.7188 | 3317547.517 | 47517219.78 | 77435870.69 | 3466487.952 | 59653871.42 | 3536648.199 | |
| P30041 | Peroxiredoxin-6 [OS=Homo sapiens] | -0.29 | 0.970551984 | 271356.4648 | 2665144.234 | 23538376.47 | 173342.4375 | 17635882.59 | 287017.2422 | 1834672.303 | 19127812.78 | 23538376.47 | 1267107.543 | 18320626.63 | 2428792.197 | Met-loss [N-Term] |
| P15104 | Glutamine synthetase [OS=Homo sapiens] | 0.64 | 0.970551984 | 44113.10156 | 6276308.328 | 102937388.8 | 72503.26563 | 55842040.16 | 220794.4063 | 298253.7589 | 45045235.86 | 102937388.8 | 529988.1325 | 58010205.18 | 1868402.494 | |
| P14618 | Pyruvate kinase PKM [OS=Homo sapiens] | 0.11 | 0.970551984 | 7472923.922 | 39970761.43 | 403514375.8 | 8874503.523 | 310244084.7 | 6112821.898 | 50525299.08 | 286871243.7 | 403514375.8 | 64871306.27 | 322289854.4 | 51727812.64 | |
| O75643 | U5 small nuclear ribonucleoprotein 200 kDa helicase [OS=Homo sapiens] | -0.16 | 0.970551984 | 2297401.75 | 746038.3125 | 1033412.188 | 1898258.375 | 2605624.25 | 1125909.75 | 15532997.76 | 5354337.294 | 1033412.188 | 13875987.55 | 2706792.176 | 9527653.442 | |
| Q9NQC3 | Reticulon-4 [OS=Homo sapiens] | 0.1 | 0.970551984 | 1259546.313 | 1130368.094 | 5835788.875 | 4473759.391 | 9135873.406 | 875103.1641 | 8515937.645 | 8112682.605 | 5835788.875 | 32702518.49 | 9490589.694 | 7405282.415 | |
| P43304 | Glycerol-3-phosphate dehydrogenase, mitochondrial [OS=Homo sapiens] | 0.18 | 0.970551984 | | 1135692.809 | 15822457.86 | 11249.60645 | 10653799.22 | | | 8150898.229 | 15822457.86 | 82232.95682 | 11067451.64 | | |
| P24539 | ATP synthase F(0) complex subunit B1, mitochondrial [OS=Homo sapiens] | -0.25 | 0.970551984 | 172277.1094 | 2788391.703 | 19235163.13 | 400555.4219 | 15843005.5 | 444016.0313 | 1164785.373 | 20012363.22 | 19235163.13 | 2928000.804 | 16458137.94 | 3757344.555 | |
| P61158 | Actin-related protein 3 [OS=Homo sapiens] | 0.55 | 0.970551984 | 22428.71289 | 1037690.672 | 15105107.91 | 60369.1543 | 12355779.91 | 42709.14844 | 151643.1103 | 7447534.223 | 15105107.91 | 441289.576 | 12835514.7 | 361412.5955 | |

| | | | | | | | | | | | | | | | | |
|---|---|---|---|---|---|---|---|---|---|---|---|---|---|---|---|---|
| P14415 | Sodium/potassium-transporting ATPase subunit beta-2 [OS=Homo sapiens | -0.06 | 0.970551984 | | 7561500.277 | 80553755.03 | 170337.2656 | 53478046.3 | 89822.35156 | | 54269093.5 | 80553755.03 | 1245140.181 | 55554425.12 | 760093.1043 | |
| Q5SSJ5 | Heterochromatin protein 1-binding protein 3 [OS=Homo sapiens] | -0.32 | 0.970551984 | 72924.26563 | 987268.8281 | 6399948.5 | 76172.67969 | 5310644.438 | | 493049.3565 | 7085655.277 | 6399948.5 | 556811.0058 | 5516839.51 | | |
| O14818 | Proteasome subunit alpha type-7 [OS=Homo sapiens] | 0.21 | 0.970551984 | 48238.63281 | 566781.0859 | 9024795.625 | 62598.66016 | 5236645.813 | 37169.14453 | 326146.9507 | 4067803.295 | 9024795.625 | 457586.9336 | 5439967.759 | 314532.0731 | |
| P61106 | Ras-related protein Rab-14 [OS=Homo sapiens] | 0.19 | 0.970551984 | 446682.7734 | 4275235.703 | 58587331.09 | 930191.8789 | 44424962.19 | 610983.8145 | 3020073.663 | 30683483.12 | 58587331.09 | 6799564.855 | 46149839.16 | 5170256.357 | Met-loss+Acetyl [N-Term] |
| Q9P035 | Very-long-chain (3R)-3-hydroxyacyl-CoA dehydratase 3 [OS=Homo sapiens] | -0.24 | 0.970551984 | | 334333.5469 | 2265688.5 | 385557.0918 | 1716752.813 | | | 2399520.975 | 2265688.5 | 2818365.232 | 1783408.748 | | |
| Q14152 | Eukaryotic translation initiation factor 3 subunit A [OS=Homo sapiens] | 0.14 | 0.970551984 | 89435.48828 | 516625.7578 | 3323887.063 | 367029.543 | 3567209.516 | 71312.85156 | 604683.634 | 3707837.139 | 3323887.063 | 2682931.595 | 3705712.674 | 603462.3429 | Met-loss [N-Term] |
| P20700 | Lamin-B1 [OS=Homo sapiens] | -0.02 | 0.970551984 | 11888534.98 | 3303317.809 | 22687233.41 | 10589018.81 | 29701219.94 | 6032395.215 | 80379753.88 | 23708001.91 | 22687233.41 | 77404159.05 | 30854421.83 | 51047227.39 | |
| O95571 | Persulfide dioxygenase ETHE1, mitochondrial [OS=Homo sapiens] | 0 | 0.970551984 | | | 360373.0938 | | 352745.3438 | | | | 360373.0938 | | 366441.2996 | | |
| P18827 | Syndecan-1 [OS=Homo sapiens] | 0.5 | 0.970551984 | 3720613.563 | 321039.5938 | 905629.625 | 4812987.797 | 4145801.641 | 1812892.383 | 25155496.69 | 2304109.911 | 905629.625 | 35182227.9 | 4306769.651 | 15341025.65 | |
| P40925 | Malate dehydrogenase, cytoplasmic [OS=Homo sapiens] | 0.52 | 0.970551984 | 217997.1094 | 4950181.492 | 102229241.1 | 359031.2578 | 54635614.06 | 627773.8809 | 1473903.557 | 35527587.43 | 102229241.1 | 2624465.315 | 56756937.48 | 5312336.958 | |
| Q96F07 | Cytoplasmic FMR1-interacting protein 2 [OS=Homo sapiens] | -0.32 | 0.970551984 | 43501.91016 | 2678871.43 | 25766299.08 | 199582.6367 | 20724279.58 | 106345.4844 | 294121.4234 | 19226333.24 | 25766299.08 | 1458919.512 | 21528936.04 | 899914.8646 | |
| P49821 | NADH dehydrogenase [ubiquinone] flavoprotein 1, mitochondrial [OS=Hom | -0.41 | 0.970551984 | 126973.6328 | 2565811.979 | 25959889.47 | 177513.7617 | 16372824.47 | 203414.8164 | 858483.3514 | 18414902.47 | 25959889.47 | 1297599.305 | 17008528.06 | 1721333.238 | |
| Q9BRX8 | Peroxiredoxin-like 2A [OS=Homo sapiens] | 0 | 0.970551984 | | | 608397.9375 | | 407158.7188 | | | | 608397.9375 | | 422967.3692 | | |
| P51812 | Ribosomal protein S6 kinase alpha-3 [OS=Homo sapiens] | 0 | 0.970551984 | | | 551550 | | 343220.2188 | | | | 551550 | | 356546.345 | | |
| Q9H1Z4 | WD repeat-containing protein 13 [OS=Homo sapiens] | 0 | 0.970551984 | | | 376769.6563 | | 450271.625 | | | | 376769.6563 | | 467754.2096 | | |
| Q9NUB1 | Acetyl-coenzyme A synthetase 2-like, mitochondrial [OS=Homo sapiens] | 0 | 0.970551984 | | | 513518.8438 | | 331065.6563 | | | | 513518.8438 | | 343919.8603 | | |
| Q14974 | Importin subunit beta-1 [OS=Homo sapiens] | -0.06 | 0.970551984 | 138724.1738 | 1402122.555 | 12406352.02 | 150940.8535 | 10164535.72 | 27508.11523 | 937930.1121 | 10063071.77 | 12406352.02 | 1103355.282 | 10559191.62 | 232778.683 | |
| P28482 | Mitogen-activated protein kinase 1 [OS=Homo sapiens] | -0.1 | 0.970551984 | 146472.1641 | 6353560.566 | 56385822.47 | 223144.8594 | 47113711.13 | 114848.75 | 990315.1662 | 45599677.28 | 56385822.47 | 1631155.87 | 48942983.5 | 971871.0476 | |
| P62937 | Peptidyl-prolyl cis-trans isomerase A [OS=Homo sapiens] | 0.39 | 0.970551984 | 2575291.918 | 23128242.85 | 344911527.6 | 3329114.445 | 204316278.3 | 3127757.758 | 17411845.18 | 165992029.1 | 344911527.6 | 24335333.49 | 212249215.7 | 26467656.01 | |
| Q86YZ3 | Hornerin [OS=Homo sapiens] | -1.12 | 0.970551984 | 3753355.269 | 1358026.586 | 2816203.25 | 1635342.879 | 171314.5156 | 9250404.156 | 25376867.14 | 9746593.806 | 2816203.25 | 11954114.22 | 177966.1018 | 78278605.36 | |
| Q9NY33 | Dipeptidyl peptidase 3 [OS=Homo sapiens] | 0 | 0.970551984 | | | 410566.8125 | | 329800.625 | | | | 410566.8125 | | 342605.712 | | |
| Q9HBH5 | Retinol dehydrogenase 14 [OS=Homo sapiens] | 0 | 0.970551984 | | | 331080.2188 | | 314037.875 | | | | 331080.2188 | | 326230.9456 | | |
| P07339 | Cathepsin D [OS=Homo sapiens] | 0 | 0.970551984 | | | 2229886.625 | | 2177409.438 | | | | 2229886.625 | | 2261951.173 | | |
| P35232 | Prohibitin 1 [OS=Homo sapiens] | -0.27 | 0.970551984 | 222774.1738 | 2883512.891 | 25206369.25 | 503367.7871 | 17569306.88 | 454339.0615 | 1506201.84 | 20695050.58 | 25206369.25 | 3679543.965 | 18251465.99 | 3844699.918 | |
| P62318 | Small nuclear ribonucleoprotein Sm D3 [OS=Homo sapiens] | 0.09 | 0.970551984 | 269453.1953 | 365142.1875 | 4443285.063 | 303864.207 | 3112241.938 | 124372.8926 | 1821804.079 | 2620635.428 | 4443285.063 | 2221202.345 | 3233080.182 | 1052466.077 | |
| Q9UPT6 | C-Jun-amino-terminal kinase-interacting protein 3 [OS=Homo sapiens] | 0 | 0.970551984 | | | 264637.0625 | | 152667.6406 | | | | 264637.0625 | | 158595.2292 | | |
| P62258 | 14-3-3 protein epsilon [OS=Homo sapiens] | -0.01 | 0.970551984 | 655021.1172 | 27865486.72 | 222020712.5 | 590105.7461 | 167412719.8 | 787036.3145 | 4428673.193 | 199991357.4 | 222020712.5 | 4313585.597 | 173912811.9 | 6660044.688 | Acetyl [N-Term] |
| Q4J6C6 | Prolyl endopeptidase-like [OS=Homo sapiens] | 0.1 | 0.970551984 | | 433713.0938 | 10746241.88 | | 6558772.172 | 110745.7578 | | 3112770.69 | 10746241.88 | | 6813427.99 | 937150.7801 | |
| P06576 | ATP synthase subunit beta, mitochondrial [OS=Homo sapiens] | 0.06 | 0.970551984 | 5953549.609 | 59809172.81 | 712102817.1 | 6769685.02 | 432965233.1 | 9231478.879 | 40252634.41 | 429252062.7 | 712102817.1 | 49485394.77 | 449775866.7 | 78118456.22 | |
| P46783 | Small ribosomal subunit protein eS10 [OS=Homo sapiens] | 0 | 0.970551984 | 36471.30078 | 403502.5234 | 2934394.563 | 43528.51172 | 2557180.938 | 53188.31055 | 246586.6639 | 2895948.604 | 2934394.563 | 318186.9732 | 2656467.967 | 450089.1746 | |
| P0C0S5 | Histone H2A.Z [OS=Homo sapiens] | -0.5 | 0.970551984 | 327485.0469 | 1546913.781 | 8865104 | 374341.8438 | 8931819.625 | 321398.5156 | 2214164.109 | 11102242.35 | 8865104 | 2736383.43 | 9278613.168 | 2719732.797 | |
| Q9P0J0 | NADH dehydrogenase [ubiquinone] 1 alpha subcomplex subunit 13 [OS=Hc | -0.33 | 0.970551984 | 67440.23438 | 1445573.875 | 8956386.25 | 62046.98828 | 8043085.25 | 126002 | 455971.1898 | 10374923.08 | 8956386.25 | 453554.2939 | 8355372.124 | 1066251.881 | |
| Q9BZL4 | Protein phosphatase 1 regulatory subunit 12C [OS=Homo sapiens] | 0 | 0.970551984 | | | 199814.1875 | | 197340.9219 | | | | 199814.1875 | | 205003.0288 | | |
| P09493 | Tropomyosin alpha-1 chain [OS=Homo sapiens] | -0.84 | 0.970551984 | 54040.43359 | 211276.3594 | 3120956.313 | 27908.09961 | 1528896.688 | 33521.01953 | 365373.5938 | 1516336.187 | 3120956.313 | 204004.0744 | 1588258.78 | 283660.9747 | |
| P63241 | Eukaryotic translation initiation factor 5A-1 [OS=Homo sapiens] | -0.83 | 0.970551984 | 345854.6211 | 1283222.676 | 15225063.84 | 855574.4297 | 9208264.672 | 567899.0547 | 2338362.915 | 9209724.105 | 15225063.84 | 6254122.353 | 9565791.678 | 4805665.27 | Met-loss+Acetyl [N-Term] |
| O95433 | Activator of 90 kDa heat shock protein ATPase homolog 1 [OS=Homo sapien | -0.07 | 0.970551984 | | 378894.4453 | 2263453 | 57322.11328 | 2413055 | | | 2719335.757 | 2263453 | 419016.1575 | 2506746.087 | | |
| P53396 | ATP-citrate synthase [OS=Homo sapiens] | -0.23 | 0.970551984 | 136895.6523 | 2746087.84 | 23414778.67 | 261371.1289 | 18559691.88 | 76286.10547 | 925567.2679 | 19708747.24 | 23414778.67 | 1910584.238 | 19280304.42 | 645546.9236 | |
| O75367 | Core histone macro-H2A.1 [OS=Homo sapiens] | 0.26 | 0.970551984 | 1134957.461 | 3080477.557 | 26718386.19 | 1409928.84 | 19338125.64 | 1580413.77 | 7673578.074 | 22108671.35 | 26718386.19 | 10306370.98 | 20088962.24 | 13373749.26 | |
| Q00341 | Vigilin [OS=Homo sapiens] | -1.75 | 0.970551984 | | 1468240.625 | 31004058 | | 19398974 | 169042.0313 | | 10537602.97 | 31004058 | | 20152173.15 | 1430464.467 | |
| Q9Y2I2 | Netrin-G1 [OS=Homo sapiens] | 0 | 0.970551984 | | | 301980.625 | | 158505.7969 | | | | 301980.625 | | 164660.0621 | | |
| P62910 | Large ribosomal subunit protein eL32 [OS=Homo sapiens] | 0.24 | 0.970551984 | 180770.8047 | 739145.082 | 2069111.688 | 227647.0625 | 4906023 | 122668.2305 | 1222212.225 | 5304864.391 | 2069111.688 | 1664066.308 | 5096507.937 | 1038040.916 | Met-loss [N-Term] |
| P39019 | Small ribosomal subunit protein eS19 [OS=Homo sapiens] | -0.33 | 0.970551984 | 290869.9258 | 1147397.391 | 8162289.172 | 436530.4336 | 8454603.75 | 202135.3828 | 1966605.059 | 8234902.333 | 8162289.172 | 3190972.811 | 8782868.55 | 1710506.438 | |
| P08559 | Pyruvate dehydrogenase E1 component subunit alpha, somatic form, mitoc | -0.33 | 0.970551984 | 29315.13281 | 3541651.797 | 44920968.25 | 145953.7949 | 21896302.94 | 25598.16992 | 198202.9883 | 25418531.44 | 44920968.25 | 1066900.622 | 22746465.25 | 216616.3778 | |
| P08243 | Asparagine synthetase [glutamine-hydrolyzing] [OS=Homo sapiens] | 0.31 | 0.970551984 | 2289433934 | 1389614464 | 3007608448 | 2624729600 | 3800225280 | 2447785675 | 15479126437 | 9973300867 | 3007608448 | 19186384601 | 3947775683 | 20713608356 | |
| Q02153 | Guanylate cyclase soluble subunit beta-1 [OS=Homo sapiens] | 0.12 | 0.970551984 | | 33840.77734 | 440303.0313 | 29312.9082 | 254823.4375 | | | 242876.181 | 440303.0313 | 214273.0171 | 264717.4038 | | |
| Q13200 | 26S proteasome non-ATPase regulatory subunit 2 [OS=Homo sapiens] | 0.15 | 0.970551984 | | 1388743.508 | 9994378.813 | 179004.0117 | 10571736.28 | 43609.69141 | | 9967049.991 | 9994378.813 | 1308492.812 | 10982202.46 | 369033.1542 | |
| Q13162 | Peroxiredoxin-4 [OS=Homo sapiens] | 0.01 | 0.970551984 | 183092.3047 | 219746.0352 | 627166.5625 | 183309.1836 | 820797.7188 | 197596.8438 | 1237908.153 | 1577123.281 | 627166.5625 | 1339962.98 | 852666.628 | 1672100.493 | |
| P29508 | Serpin B3 [OS=Homo sapiens] | 0.4 | 0.970551984 | 323899.4648 | 329158.623 | 254663.4688 | 239503.8262 | 3093778.484 | 727697.0469 | 2189921.576 | 2362380.405 | 254663.4688 | 1750737.494 | 3213899.853 | 6157904.995 | |
| P35908 | Keratin, type II cytoskeletal 2 epidermal [OS=Homo sapiens] | 0.27 | 0.970551984 | 96767310.72 | 29363847.7 | 92376335.09 | 91029037.75 | 328439132.4 | 78766393.63 | 654254929.7 | 210745134.9 | 92376335.09 | 665408782.7 | 341191357.1 | 666535573.8 | |
| Q9UQB3 | Catenin delta-2 [OS=Homo sapiens] | 0 | 0.970551984 | | | 588055.7969 | | 272890.0313 | | | | 588055.7969 | | 283485.4647 | | |
| P35637 | RNA-binding protein FUS [OS=Homo sapiens] | -0.03 | 0.970551984 | 105248.0508 | 494111.4375 | 6123946.938 | 95398.05859 | 6019853.188 | 80579.17969 | 711594.1897 | 3546251.249 | 6123946.938 | 697345.678 | 6253584.532 | 681875.6998 | |
| Q9BQ69 | ADP-ribose glycohydrolase MACROD1 [OS=Homo sapiens] | 0 | 0.970551984 | | | 707136 | | 394247.7188 | | | | 707136 | | 409555.077 | | |
| Q15907 | Ras-related protein Rab-11B [OS=Homo sapiens] | -0.24 | 0.970551984 | 168739.9609 | 3013675.805 | 27447190.03 | 394817.1016 | 21653990.91 | 200191.6875 | 1140870.305 | 21629233.36 | 27447190.03 | 2886054.532 | 22494745.03 | 1694058.534 | Met-loss+Acetyl [N-Term] |
| Q16555 | Dihydropyrimidinase-related protein 2 [OS=Homo sapiens] | 0.1 | 0.970551984 | 492615.9102 | 54231991.91 | 622793746 | 455616.1113 | 431168567.7 | 2208118.436 | 3330632.889 | 389224483.5 | 622793746 | 3330486.288 | 447909442.6 | 18685500.51 | |
| Q8WY54 | Protein phosphatase 1E [OS=Homo sapiens] | 0 | 0.970551984 | | | 397946.25 | | 512902.7188 | | | | 397946.25 | | 532817.0653 | | |
| P35754 | Glutaredoxin-1 [OS=Homo sapiens] | 0.02 | 0.970551984 | 166375.6563 | 117305.0469 | 1739243.813 | 156055.8125 | 403124.6563 | 171582.75 | 1124884.969 | 841901.5173 | 1739243.813 | 1140744.874 | 418776.6772 | 1451964.492 | Met-loss+Acetyl [N-Term] |
| P18077 | Large ribosomal subunit protein eL33 [OS=Homo sapiens] | -0.5 | 0.970551984 | | 618104.6094 | 3381383.313 | 126783.7852 | 2294799.719 | 41330.75391 | | 4436153.621 | 3381383.313 | 926770.6903 | 2383899.338 | 349748.3699 | |
| Q9HCC0 | Methylcrotonoyl-CoA carboxylase beta chain, mitochondrial [OS=Homo sap | 0 | 0.970551984 | | | 378308.6875 | | 236048.9219 | | | | 378308.6875 | | 245213.9348 | | |
| P07195 | L-lactate dehydrogenase B chain [OS=Homo sapiens] | -0.08 | 0.970551984 | 2582372.949 | 14364748.66 | 132005608.5 | 3059873.25 | 92182325.25 | 1927491.086 | 17459720.85 | 103096192.5 | 132005608.5 | 22367220.23 | 95761465.5 | 16310780.75 | |
| Q00839 | Heterogeneous nuclear ribonucleoprotein U [OS=Homo sapiens] | 0.03 | 0.970551984 | 6734399.273 | 5840679.977 | 45742505.69 | 8508250.992 | 44725946.88 | 3691144.428 | 45532048.9 | 41918719.32 | 45742505.69 | 62194054.51 | 46462510.11 | 31235136.66 | Met-loss+Acetyl [N-Term] |
| P60709 | Actin, cytoplasmic 1 [OS=Homo sapiens] | 0.22 | 0.970551984 | 28052918.48 | 259751194.3 | 2106560374 | 31184412.33 | 1889162732 | 26773255.46 | 189669011.9 | 1864241398 | 2106560374 | 227953435.1 | 1962512784 | 226560165.7 | Met-loss+Acetyl [N-Term] |
| P14866 | Heterogeneous nuclear ribonucleoprotein L [OS=Homo sapiens] | -0.39 | 0.970551984 | 43245.27734 | 1757063.82 | 12224534.56 | 169569.0313 | 9906403.031 | 80405.37891 | 292386.2994 | 12610494.91 | 12224534.56 | 1239524.502 | 10291036.48 | 680404.9659 | |
| Q9C0C9 | (E3-independent) E2 ubiquitin-conjugating enzyme [OS=Homo sapiens] | -0.04 | 0.970551984 | 476954.0625 | 261070.1367 | 3050884.75 | 380371.5469 | 2717221.344 | 116020.7266 | 3224741.334 | 1873707.484 | 3050884.75 | 2780459.667 | 2822722.222 | 981788.5268 | |
| Q3MHD2 | Protein LSM12 [OS=Homo sapiens] | 0 | 0.970551984 | | | 843674.125 | | 483399.75 | | | | 843674.125 | | 502168.5921 | | |
| Q7Z7K6 | Centromere protein V [OS=Homo sapiens] | -2.47 | 0.970551984 | | 36450.76563 | 2148526.188 | | 1302362.406 | | | 261608.138 | 2148526.188 | | 1352928.908 | | |
| Q9NY47 | Voltage-dependent calcium channel subunit alpha-2/delta-2 [OS=Homo sa | 0.29 | 0.970551984 | 524093216 | 1147523.512 | 11929317.63 | 7171585.5 | 12980648.84 | 4061514.75 | 3543454578 | 8235807.507 | 11929317.63 | 52423227.75 | 13484645.27 | 34369277.81 | |
| Q01484 | Ankyrin-2 [OS=Homo sapiens] | 0.08 | 0.970551984 | 101064.9766 | 4824225.637 | 75282098.75 | 98877.94531 | 40390558.34 | 100952.8516 | 683311.9433 | 34623598.83 | 75282098.75 | 722783.1344 | 41958792.52 | 854281.4232 | |
| Q9NVJ2 | ADP-ribosylation factor-like protein 8B [OS=Homo sapiens] | -0.62 | 0.970551984 | | 562802.9844 | 6726157.875 | 17965.63281 | 3638986.875 | 20404.875 | | 4039252.352 | 6726157.875 | 131326.1148 | 3780276.915 | 172669.77 | |
| Q9NPJ3 | Acyl-coenzyme A thioesterase 13 [OS=Homo sapiens] | 0 | 0.970551984 | | | 156611.6406 | | 246229.1875 | | | | 156611.6406 | | 255789.467 | | |
| P25788 | Proteasome subunit alpha type-3 [OS=Homo sapiens] | -0.24 | 0.970551984 | 360382.043 | 1355185.643 | 10880079.61 | 333289.8516 | 7750600.063 | 116333.9902 | 2436584.487 | 9726204.278 | 10880079.61 | 2436299.448 | 8051530.687 | 984439.421 | Met-loss+Acetyl [N-Term] |
| Q99873 | Protein arginine N-methyltransferase 1 [OS=Homo sapiens] | 0.07 | 0.970551984 | 549176.6543 | 501343.3477 | 5948063.906 | 835626.5703 | 5013852.688 | 296389.5664 | 3713046.593 | 3598154.865 | 5948063.906 | 6108306.456 | 5208524.301 | 2508102.512 | |
| P12109 | Collagen alpha-1(VI) chain [OS=Homo sapiens] | 0 | 0.970551984 | | 256484.6875 | | | | 131275.5156 | | 1840797.59 | | | | 1110877.331 | |
| P53999 | Activated RNA polymerase II transcriptional coactivator p15 [OS=Homo sap | -0.02 | 0.970551984 | 314957.75 | 417502.8594 | 1758011.313 | 962705.7891 | 2239975.156 | 468250.9219 | 2129465.612 | 2996429.396 | 1758011.313 | 7037236.722 | 2326946.115 | 3962424.615 | |
| Q96DR5 | BPI fold-containing family A member 2 [OS=Homo sapiens] | -1.16 | 0.970551984 | | 33858.91016 | 255646.0781 | | 37915376.5 | | | 243006.3207 | 255646.0781 | | 39387507.41 | | |
| P26641 | Elongation factor 1-gamma [OS=Homo sapiens] | -0.04 | 0.970551984 | 980357.4219 | 2714306.617 | 22187932.25 | 1102920.77 | 19031937.72 | 609400.7656 | 6628309.409 | 19480652.54 | 22187932.25 | 8062187.461 | 19770886.04 | 5156860.31 | Met-loss+Acetyl [N-Term] |
| O60313 | Dynamin-like 120 kDa protein, mitochondrial [OS=Homo sapiens] | -1.66 | 0.970551984 | | 2029825.176 | 18800270.38 | | 11842606.81 | 23273.45508 | | 14568110.59 | 18800270.38 | | 12302416.77 | 196944.2172 | |
| P21399 | Cytoplasmic aconitate hydratase [OS=Homo sapiens] | -0.72 | 0.970551984 | 45022.05078 | 435129.1367 | 2678794.844 | 75389.39453 | 1587657.219 | 15243.30664 | 304399.2692 | 3122933.669 | 2678794.844 | 551085.3073 | 1649300.791 | 128991.6381 | |
| P51149 | Ras-related protein Rab-7a [OS=Homo sapiens] | -0.05 | 0.970551984 | 166489.9609 | 2725197.453 | 26249014.19 | 217866.9375 | 19762359.66 | 212766.8047 | 1125657.796 | 19558816.37 | 26249014.19 | 1592575.043 | 20529667.89 | 1800471.467 | |
| Q15717 | ELAV-like protein 1 [OS=Homo sapiens] | -0.08 | 0.970551984 | | 691554.0547 | 5905136.906 | 113355.4844 | 4922036.438 | 82447.10938 | | 4963302.291 | 5905136.906 | 828611.8006 | 5113143.124 | 697682.4611 | |
| O76003 | Glutaredoxin-3 [OS=Homo sapiens] | 0 | 0.970551984 | | | 673794.5625 | | 447723.7188 | | | | 673794.5625 | | 465107.3764 | | |
| P62857 | Small ribosomal subunit protein eS28 [OS=Homo sapiens] | -0.36 | 0.970551984 | | 509422.0469 | 3646254.688 | 39741.74609 | 2662614.875 | | | 3656135.909 | 3646254.688 | 290506.2774 | 2765995.562 | | |
| O15498 | Synaptobrevin homolog YKT6 [OS=Homo sapiens] | -0.25 | 0.970551984 | | 910302.3906 | 7010741.625 | 89977.09766 | 5218264.563 | | | 6533265.058 | 7010741.625 | 657719.2565 | 5420872.825 | | |

| | | | | | | | | | | | | | | | | |
|---|---|---|---|---|---|---|---|---|---|---|---|---|---|---|---|---|
| Q14254 | Flotillin-2 [OS=Homo sapiens] | -0.15 | 0.970551984 | 123318.7383 | 1060393.664 | 13829491.28 | 74075.38281 | 8144501.148 | 112826.3477 | 833772.1886 | 7610474.217 | 13829491.28 | 541480.0763 | 8460725.672 | 954757.1105 | |
| Q13813 | Spectrin alpha chain, non-erythrocytic 1 [OS=Homo sapiens] | -0.1 | 0.970551984 | 1630560.247 | 29217089.62 | 307787654.7 | 1886795.519 | 203491107.3 | 2581769.527 | 11024405.58 | 209691848.2 | 307787654.7 | 13792195.77 | 211392006 | 21847404.13 | |
| P62256 | Ubiquitin-conjugating enzyme E2 H [OS=Homo sapiens] | 0.28 | 0.970551984 | 172680.5391 | 50632.52734 | 278105 | 277055.4453 | 1409743.031 | 261644.6641 | 1167513.007 | 363391.0282 | 278105 | 2025234.268 | 1464478.774 | 2214084.818 | |
| P04843 | Dolichyl-diphosphooligosaccharide--protein glycosyltransferase subunit 1 [ | 0.02 | 0.970551984 | 466487.3203 | 397365.8867 | 3929273.625 | 999952.0469 | 4418037.156 | 239183.4922 | 3153974.485 | 2851905.795 | 3929273.625 | 7309501.35 | 4589575.188 | 2024014.289 | |
| P22352 | Glutathione peroxidase 3 [OS=Homo sapiens] | 0.14 | 0.970551984 | 289706.875 | 289334.9688 | 718335.9375 | 295494.5 | 929678.125 | 369897.2188 | 1958741.539 | 2076564.953 | 718335.9375 | 2160021.026 | 965774.5067 | 3130137.659 | |
| P50583 | Bis(5'-nucleosyl)-tetraphosphatase [asymmetrical] [OS=Homo sapiens] | 0 | 0.970551984 | | | 440261.9375 | | 122146.5938 | | | | 440261.9375 | | 126889.1492 | | |
| O43920 | NADH dehydrogenase [ubiquinone] iron-sulfur protein 5 [OS=Homo sapiens | -0.78 | 0.970551984 | 111754.375 | 407365.5938 | 2219625 | 121023.7734 | 1637461.75 | 100557.1094 | 755584.1969 | 2923673.965 | 2219625 | 884665.8578 | 1701039.071 | 850932.5807 | Met-loss [N-Term] |
| Q02218 | 2-oxoglutarate dehydrogenase complex component E1 [OS=Homo sapiens] | -0.4 | 0.970551984 | 132092.3672 | 2641192.303 | 24454728.5 | 331478.8203 | 17211933.83 | 295106.6914 | 893091.7039 | 18955909.11 | 24454728.5 | 2423061.078 | 17880217.31 | 2497246.59 | |
| Q01581 | Hydroxymethylglutaryl-CoA synthase, cytoplasmic [OS=Homo sapiens] | 0 | 0.970551984 | | | 268053.3438 | | 158370.6719 | | | | 268053.3438 | | 164519.6906 | | |
| P02686 | Myelin basic protein [OS=Homo sapiens] | 0.04 | 0.970551984 | 73939.96875 | 30853977.46 | 431308164.8 | 86878.75781 | 227220971.3 | 569497.3594 | 499916.6423 | 221439836.8 | 431308164.8 | 635070.8511 | 236043223.5 | 4819190.416 | |
| P62424 | Large ribosomal subunit protein eL8 [OS=Homo sapiens] | -0.39 | 0.970551984 | 253482.4453 | 1749761.953 | 11181045.66 | 488483.9727 | 9940587.25 | 269461.166 | 1713823.999 | 12558089.21 | 11181045.66 | 3570745.486 | 10326547.96 | 2280229.482 | |
| Q9Y2Q0 | Phospholipid-transporting ATPase IA [OS=Homo sapiens] | -1.51 | 0.970551984 | | 2205215.395 | 16417391.22 | | 11907626.78 | 19129.97656 | | 15826890.97 | 16417391.22 | | 12369961.25 | 161881.3471 | |
| P62249 | Small ribosomal subunit protein uS9 [OS=Homo sapiens] | -0.32 | 0.970551984 | 647277.1953 | 2114197.367 | 13907741.42 | 917404.25 | 12814171.16 | 574688.7773 | 4376315.646 | 15173652.1 | 13907741.42 | 6706089.182 | 13311703.8 | 4863121.14 | |
| Q99497 | Parkinson disease protein 7 [OS=Homo sapiens] | -0.57 | 0.970551984 | 104034.1172 | 3315718.125 | 27390107.84 | 254514.2344 | 18934183.99 | 96481.43359 | 703386.6449 | 23796999.32 | 27390107.84 | 1860461.355 | 19669336.85 | 816443.4697 | |
| O94905 | Erlin-2 [OS=Homo sapiens] | 0 | 0.970551984 | 160419.0742 | 885369.9189 | 6857234.25 | 137791.4414 | 6594856.469 | 25579.59961 | 1084611.832 | 6354324.03 | 6857234.25 | 1007235.027 | 6850913.323 | 216459.2324 | |
| Q9H0E2 | Toll-interacting protein [OS=Homo sapiens] | 0.52 | 0.970551984 | 4184004.5 | 883796.6055 | 6178498.078 | 3836473.75 | 8834615.742 | 3944708.5 | 28288536.18 | 6343032.316 | 6178498.078 | 28044054.85 | 9177635.174 | 33380842.04 | Met-loss+Acetyl [N-Term] |
| O94826 | Mitochondrial import receptor subunit TOM70 [OS=Homo sapiens] | -1.51 | 0.970551984 | 41736.67188 | 2176252.703 | 19173402.63 | | 15421204.53 | 36027.71484 | 282186.4442 | 15619024.94 | 19173402.63 | | 16019959.81 | 304873.0871 | |
| Q03252 | Lamin-B2 [OS=Homo sapiens] | -0.33 | 0.970551984 | | 514908.7578 | 3228869.625 | 31091.19336 | 2869773.188 | | | 3695514.183 | 3228869.625 | 227272.0182 | 2981197.158 | | |
| P16615 | Sarcoplasmic/endoplasmic reticulum calcium ATPase 2 [OS=Homo sapien: | -0.09 | 0.970551984 | 1278944.84 | 3331235.877 | 24425793.02 | 3246818.813 | 24444104.58 | 1197515.908 | 8647093.322 | 23908370.65 | 24425793.02 | 23733764.6 | 25393189.76 | 10133597.8 | |
| Q9H0Q0 | CYFIP-related Rac1 interactor A [OS=Homo sapiens] | 0 | 0.970551984 | | | 582135.3125 | | 274542.3438 | | | | 582135.3125 | | 285201.9312 | | |
| Q96DH6 | RNA-binding protein Musashi homolog 2 [OS=Homo sapiens] | 0 | 0.970551984 | | | 668616.9063 | | 186114.25 | | | | 668616.9063 | | 193340.4618 | | |
| P05089 | Arginase-1 [OS=Homo sapiens] | 0.02 | 0.970551984 | 89124.67969 | 64330.46094 | | 83841.86719 | | 150321.625 | 602582.2213 | 461701.4708 | | 612871.6305 | | 1272048.979 | |
| Q9NRG7 | Epimerase family protein SDR39U1 [OS=Homo sapiens] | 0 | 0.970551984 | | | 374969.7813 | | 326462.4688 | | | | 374969.7813 | | 339137.946 | | |
| Q969Z3 | Mitochondrial amidoxime reducing component 2 [OS=Homo sapiens] | -0.85 | 0.970551984 | 103045.7734 | 250412.1875 | 1407231.5 | 120606.8438 | 961300 | 67809.6875 | 696704.339 | 1797215.092 | 1407231.5 | 881618.1635 | 998624.1564 | 573817.9303 | |
| Q9NZD2 | Glycolipid transfer protein [OS=Homo sapiens] | 0 | 0.970551984 | | | 309628.0625 | | 150601.7813 | | | | 309628.0625 | | 156449.1592 | | |
| P62266 | Small ribosomal subunit protein uS12 [OS=Homo sapiens] | 0.03 | 0.970551984 | 188203.9844 | 421448.3125 | 6435081.5 | 177398.1563 | 3534256.75 | 101049.3359 | 1272468.808 | 3024746.021 | 6435081.5 | 1296754.246 | 3671480.459 | 855097.8916 | |
| A6NKH3 | Putative ribosomal protein eL43-like [OS=Homo sapiens] | 0 | 0.970551984 | | | 465186.2813 | | 312496.4375 | | | | 465186.2813 | | 324629.6591 | | |
| Q8TDB8 | Solute carrier family 2, facilitated glucose transporter member 14 [OS=Hom | 0 | 0.970551984 | | | 380307.6875 | | 207740.7813 | | | | 380307.6875 | | 215806.681 | | |
| P63173 | Large ribosomal subunit protein eL38 [OS=Homo sapiens] | 0.34 | 0.970551984 | 51431.21875 | | 5121205.5 | 60172.44531 | 1942518.25 | 59454.71094 | 347732.3918 | | 5121205.5 | 439851.6625 | 2017939.924 | 503116.5964 | |
| P02533 | Keratin, type I cytoskeletal 14 [OS=Homo sapiens] | 0 | 0.970551984 | 28565618.87 | 14228706.7 | 30027083.81 | 23657118.73 | 145498008 | 42209276.17 | 193135438.3 | 102119815.7 | 30027083.81 | 172930033.8 | 151147223 | 357182585.3 | |
| P53597 | Succinate--CoA ligase [ADP/GDP-forming] subunit alpha, mitochondrial [OS | -0.47 | 0.970551984 | 110340.2578 | 2242421.172 | 17776393.91 | 106108.0313 | 11660134.19 | 115422.1484 | 746023.1878 | 16093917.84 | 17776393.91 | 775633.9917 | 12112859.32 | 976723.2496 | |
| O43295 | SLIT-ROBO Rho GTPase-activating protein 3 [OS=Homo sapiens] | 0 | 0.970551984 | | | 340348.3125 | | 266649.75 | | | | 340348.3125 | | 277002.8936 | | |
| P41091 | Eukaryotic translation initiation factor 2 subunit 3 [OS=Homo sapiens] | -0.55 | 0.970551984 | | 463292.3418 | 4012157.688 | 216920.3945 | 2815484.813 | 83941.26172 | | 3325061.759 | 4012157.688 | 1585655.954 | 2924800.942 | 710326.2505 | |
| P07814 | Bifunctional glutamate/proline--tRNA ligase [OS=Homo sapiens] | 0.05 | 0.970551984 | 31059.02148 | 142875.2578 | 1298275.344 | 175945.3164 | 672547.9375 | | 209993.6204 | 1025419.618 | 1298275.344 | 1286134.202 | 698660.7893 | | |
| Q96CS3 | FAS-associated factor 2 [OS=Homo sapiens] | 0 | 0.970551984 | | | 453806.4375 | | 335915.875 | | | | 453806.4375 | | 348958.3973 | | |
| Q4KMP7 | TBC1 domain family member 10B [OS=Homo sapiens] | 0 | 0.970551984 | | | 492693.4063 | | 300284.9688 | | | | 492693.4063 | | 311944.0587 | | |
| P61353 | Large ribosomal subunit protein eL27 [OS=Homo sapiens] | -0.2 | 0.970551984 | 1246746.506 | 2042624.078 | 11387070.19 | 1140492.943 | 10765279.69 | 1036864.133 | 8429396.678 | 14659968.66 | 11387070.19 | 8336834.486 | 11183260.53 | 8774133.205 | |
| Q9UHD1 | Cysteine and histidine-rich domain-containing protein 1 [OS=Homo sapiens | 0.54 | 0.970551984 | | 155307.5313 | 2965050.188 | 116920.7344 | 1561508.438 | | | 1114646.383 | 2965050.188 | 854673.25 | 1622136.738 | | |
| Q99729 | Heterogeneous nuclear ribonucleoprotein A/B [OS=Homo sapiens] | 0 | 0.970551984 | | | 390802.4688 | | 144876.2656 | | | | 390802.4688 | | 150501.3404 | | |
| P49748 | Very long-chain specific acyl-CoA dehydrogenase, mitochondrial [OS=Hom | 0 | 0.970551984 | | | 652409.125 | | 511502.6875 | | | | 652409.125 | | 531362.6753 | | |
| Q9Y3A5 | Ribosome maturation protein SBDS [OS=Homo sapiens] | 0 | 0.970551984 | | | 377747.6875 | | 168982.7656 | | | | 377747.6875 | | 175543.8175 | | |
| Q53HC9 | EARP and GARP complex-interacting protein 1 [OS=Homo sapiens] | 0 | 0.970551984 | | | 397362.3438 | | 272663.3438 | | | | 397362.3438 | | 283249.9757 | | |
| Q14151 | Scaffold attachment factor B2 [OS=Homo sapiens] | 0 | 0.970551984 | | | 563673.4375 | | 377080.625 | | | | 563673.4375 | | 391721.4408 | | |
| P07737 | Profilin-1 [OS=Homo sapiens] | 0.47 | 0.970551984 | 340789.4844 | 498088.2461 | 8624424.75 | 636032.0586 | 9371182.25 | 313261.8516 | 2304116.943 | 3574792.912 | 8624424.75 | 4649300.139 | 9735034.816 | 2650878.863 | |
| Q13247 | Serine/arginine-rich splicing factor 6 [OS=Homo sapiens] | 0.3 | 0.970551984 | 5366912.563 | 1313943.297 | 5377527.281 | 5261590.5 | 10805146.38 | 2147230.25 | 36286313.79 | 9430206.839 | 5377527.281 | 38461447.21 | 11224675.11 | 18170253.59 | |
| Q9NV96 | Cell cycle control protein 50A [OS=Homo sapiens] | 0 | 0.970551984 | | | 405203 | | 348180.0625 | | | | 405203 | | 361698.7633 | | |
| Q9UL25 | Ras-related protein Rab-21 [OS=Homo sapiens] | -0.29 | 0.970551984 | | 553164.4453 | 6073365.063 | 35066.23438 | 4270621.344 | 39801.32031 | | 3970076.295 | 6073365.063 | 256328.9793 | 4436435.698 | 336806.0242 | |
| Q99747 | Gamma-soluble NSF attachment protein [OS=Homo sapiens] | 0.01 | 0.970551984 | | 2358435.617 | 26439290.5 | 336232.1797 | 18584231.13 | 189220.3711 | | 16926556.68 | 26439290.5 | 2457807.431 | 19305796.45 | 1601217.256 | |
| P26640 | Valine--tRNA ligase [OS=Homo sapiens] | 0.02 | 0.970551984 | 25972.10352 | 115326.8555 | 782597.5 | 44267.79297 | 791245.1875 | | 175600.3823 | 827703.9837 | 782597.5 | 323591.0097 | 821966.6679 | | |
| Q13596 | Sorting nexin-1 [OS=Homo sapiens] | -0.03 | 0.970551984 | | 96851.21875 | 953894.7813 | 44056.69922 | 660474.0156 | | | 695103.8356 | 953894.7813 | 322047.9456 | 686118.0762 | | |
| P17858 | ATP-dependent 6-phosphofructokinase, liver type [OS=Homo sapiens] | 0.21 | 0.970551984 | | 315995.2266 | 5557767.625 | 53615.84766 | 3763214.125 | 36471.82813 | | 2267906.35 | 5557767.625 | 391923.9048 | 3909327.505 | 308631.2546 | |
| Q9Y6T7 | Diacylglycerol kinase beta [OS=Homo sapiens] | 0 | 0.970551984 | | | 232195.6406 | | 342039.8438 | | | | 232195.6406 | | 355320.1398 | | |
| Q92945 | Far upstream element-binding protein 2 [OS=Homo sapiens] | -0.1 | 0.970551984 | | 238243.0078 | 2197813.281 | 38137.14063 | 2070752.781 | | | 1709876.558 | 2197813.281 | 278776.8491 | 2151153.385 | | |
| P04406 | Glyceraldehyde-3-phosphate dehydrogenase [OS=Homo sapiens] | 0.01 | 0.970551984 | 9519070.973 | 71263132.29 | 566498802.2 | 10904757.28 | 403506042.3 | 8949365.34 | 64359534.89 | 511457441.9 | 566498802.2 | 79712160.53 | 419172871.2 | 75731160.05 | |
| P11498 | Pyruvate carboxylase, mitochondrial [OS=Homo sapiens] | -0.8 | 0.970551984 | 441750.9258 | 1565962.824 | 13237812.69 | 427601.4688 | 6937282.844 | 325977.9414 | 2986728.873 | 11238957.85 | 13237812.69 | 3125703.4 | 7206635.002 | 2758484.732 | |
| P36404 | ADP-ribosylation factor-like protein 2 [OS=Homo sapiens] | 0 | 0.970551984 | | | 302905.5313 | | 245107.4531 | | | | 302905.5313 | | 254624.1793 | | |
| Q9Y6R1 | Electrogenic sodium bicarbonate cotransporter 1 [OS=Homo sapiens] | -0.3 | 0.970551984 | 221001.7734 | 3672253.961 | 37244815.94 | 90719.11328 | 24410295.69 | 86495.22656 | 1494218.437 | 26355866.73 | 37244815.94 | 663143.2808 | 25358068.18 | 731938.3663 | |
| O43776 | Asparagine--tRNA ligase, cytoplasmic [OS=Homo sapiens] | -0.76 | 0.970551984 | 50918.69531 | 39491.04688 | 4027351.719 | 27883.10156 | 2346872.969 | | 344267.1619 | 283428.3193 | 4027351.719 | 203821.3424 | 2437994.423 | | |
| P04040 | Catalase [OS=Homo sapiens] | 0.33 | 0.970551984 | 371095.6875 | 344905.7813 | 1012172.5 | 431692.9375 | 2247111.938 | 434103.5625 | 2509020.672 | 2475398.189 | 1012172.5 | 3155611.431 | 2334359.995 | 3673463.438 | |
| P23246 | Splicing factor, proline- and glutamine-rich [OS=Homo sapiens] | -0.28 | 0.970551984 | 290567.7969 | 3231172.023 | 22010811 | 361214.9219 | 18866874.19 | 176387.335 | 1964562.331 | 23190209.64 | 22010811 | 2640427.576 | 19599413.63 | 1492621.766 | |
| Q01518 | Adenylyl cyclase-associated protein 1 [OS=Homo sapiens] | -0.02 | 0.970551984 | 68653.22656 | 1300091.057 | 8767840.453 | 74084.24121 | 8012574.219 | 49215.57031 | 464172.3696 | 9330788.934 | 8767840.453 | 541544.8299 | 8323676.448 | 416471.1229 | Met-loss+Acetyl [N-Term] |
| P21579 | Synaptotagmin-1 [OS=Homo sapiens] | 0.3 | 0.970551984 | 146111.4082 | 7724363.051 | 92966010.97 | 405243.7422 | 62649343.59 | 99090.11914 | 987876.0543 | 55437964.06 | 92966010.97 | 2962271.731 | 65081814.1 | 838518.6421 | |
| Q08AM6 | Protein VAC14 homolog [OS=Homo sapiens] | 0 | 0.970551984 | | | 295849.875 | | 235632.3281 | | | | 295849.875 | | 244781.166 | | |
| Q15111 | Inactive phospholipase C-like protein 1 [OS=Homo sapiens] | 0 | 0.970551984 | | | 192981.9063 | | 144356.3281 | | | | 192981.9063 | | 149961.2154 | | |
| Q5VYK3 | Proteasome adapter and scaffold protein ECM29 [OS=Homo sapiens] | 0.1 | 0.970551984 | 9333390 | 4702706.5 | 14237259 | 9273219 | 19139078 | 7435730 | 63104124.45 | 33751452.67 | 14237259 | 67785857.34 | 19882186.23 | 62922502.03 | |
| P80404 | 4-aminobutyrate aminotransferase, mitochondrial [OS=Homo sapiens] | 0.23 | 0.970551984 | 26040.56445 | 4987702.539 | 66679850.63 | 81981.70703 | 37539613.61 | 94949.14844 | 176063.2546 | 35796877.01 | 66679850.63 | 599274.1353 | 38997154.87 | 803476.9935 | |
| Q9Y4G6 | Talin-2 [OS=Homo sapiens] | -0.28 | 0.970551984 | 187677.4688 | 2611197.164 | 77653051.81 | 180889.8828 | 43919856.41 | 309970.9063 | 1268908.976 | 18740633.19 | 77653051.81 | 1322278.25 | 45625121.77 | 2623030.284 | |
| P07910 | Heterogeneous nuclear ribonucleoproteins C1/C2 [OS=Homo sapiens] | 0.42 | 0.970551984 | 2476229.539 | 1748476.563 | 14147935 | 3944135.469 | 14400904.75 | 1871981.996 | 16742073.03 | 12548863.93 | 14147935 | 28831046.07 | 14960045.1 | 15841052.72 | Met-loss+Acetyl [N-Term] |
| P14854 | Cytochrome c oxidase subunit 6B1 [OS=Homo sapiens] | -0.91 | 0.970551984 | 17631.55469 | 271506.875 | 9089565.625 | 19752.44336 | 3636597.5 | 24407.86914 | 119208.9714 | 1948612.239 | 9089565.625 | 144387.4352 | 3777794.768 | 206543.8357 | |
| P62805 | Histone H4 [OS=Homo sapiens] | -0.09 | 0.970551984 | 11125613.23 | 72485043.57 | 499753023.3 | 10397427.05 | 430573029.7 | 13037090.63 | 75221552.04 | 520227132.5 | 499753023.3 | 76003651.69 | 447290781.8 | 110322236.2 | |
| Q6IAA8 | Ragulator complex protein LAMTOR1 [OS=Homo sapiens] | 0 | 0.970551984 | | | 684087.75 | | 421926.125 | | | | 684087.75 | | 438308.1459 | | |
| P50452 | Serpin B8 [OS=Homo sapiens] | -0.75 | 0.970551984 | 53087.14063 | 168076.6563 | 741310.8125 | 58376.41797 | 692803.5625 | | 358928.2703 | 1206290.741 | 741310.8125 | 426722.9686 | 719702.8744 | | |
| Q16720 | Plasma membrane calcium-transporting ATPase 3 [OS=Homo sapiens] | -0.06 | 0.970551984 | 78172.49219 | 3646250.898 | 31174370.28 | 108224.1602 | 24106548.17 | 21647.39648 | 528533.2206 | 26169242.04 | 31174370.28 | 791102.5806 | 25042527.13 | 183184.2131 | Met-loss+Acetyl [N-Term] |
| P39656 | Dolichyl-diphosphooligosaccharide--protein glycosyltransferase 48 kDa sul | 0.5 | 0.970551984 | | 407909.3359 | 2350846.938 | 152916.9023 | 2056048.469 | 77716.04688 | | 2927576.417 | 2350846.938 | 1117799.906 | 2135878.152 | 657647.3483 | |
| O15144 | Actin-related protein 2/3 complex subunit 2 [OS=Homo sapiens] | -0.36 | 0.970551984 | 476424.9375 | 3364148.555 | 23862340.38 | 323078.623 | 20978534.25 | 307909.6992 | 3221163.859 | 24144585.83 | 23862340.38 | 2361656.88 | 21793062.59 | 2605587.975 | |
| P62070 | Ras-related protein R-Ras2 [OS=Homo sapiens] | -0.02 | 0.970551984 | | 146604.0469 | 808939.7969 | 24217.83594 | 1077511.5 | | | 1052181.239 | 808939.7969 | 177028.7936 | 1119347.771 | | |
| Q969M1 | Mitochondrial import receptor subunit TOM40B [OS=Homo sapiens] | 0 | 0.970551984 | | | 628783.875 | | 327445.3438 | | | | 628783.875 | | 340158.9828 | | |
| P82979 | SAP domain-containing ribonucleoprotein [OS=Homo sapiens] | 0 | 0.970551984 | | | 548931.8125 | | 327356.5625 | | | | 548931.8125 | | 340066.7545 | | |
| Q00325 | Solute carrier family 25 member 3 [OS=Homo sapiens] | -0.05 | 0.970551984 | 2551515.043 | 8262249.641 | 64419023.22 | 3497766.277 | 58679099.03 | 1741370.02 | 17251086.99 | 59298390.77 | 64419023.22 | 25568153.4 | 60957417.84 | 14735790.38 | |
| Q14137 | Ribosome biogenesis protein BOP1 [OS=Homo sapiens] | 0.32 | 0.970551984 | 552619.7031 | 32296.21484 | | 319023.2188 | 279060.1875 | 127372.4258 | 3736325.442 | 231790.8138 | | 2332012.475 | 289895.1881 | 1077848.674 | |
| P14923 | Junction plakoglobin [OS=Homo sapiens] | 0.15 | 0.970551984 | 2378867.211 | 1976272.758 | 3549480.281 | 2385989.578 | 14731912.28 | 3506821.031 | 16083795.12 | 14183763.43 | 3549480.281 | 17441230.4 | 15303904.58 | 29675358.5 | |
| P25786 | Proteasome subunit alpha type-1 [OS=Homo sapiens] | -0.59 | 0.970551984 | 124710.1289 | 1430839.926 | 15456680.94 | 205458.9844 | 9935073.938 | 138427.832 | 843179.5408 | 10269177.13 | 15456680.94 | 1501874.743 | 10320820.59 | 1171401.536 | |

| | | | | | | | | | | | | | | | | |
|---|---|---|---|---|---|---|---|---|---|---|---|---|---|---|---|---|
| Q01064 | Dual specificity calcium/calmodulin-dependent 3',5'-cyclic nucleotide phos | -2.8 | 0.970551984 | | 83631.30078 | 3279581.094 | | 1684757.328 | | | 600224.1242 | 3279581.094 | | 1750170.983 | | |
| P09382 | Galectin-1 [OS=Homo sapiens] | 0.26 | 0.970551984 | 282583.9766 | 323056.6641 | 850846.5625 | 313437.4844 | | 345228.4219 | 1910582.803 | 2318586.479 | 850846.5625 | 2291181.584 | | 2921385.805 | |
| Q13085 | Acetyl-CoA carboxylase 1 [OS=Homo sapiens] | 0.45 | 0.970551984 | 12087799 | 7976283 | 18251048 | 15288314 | 22245354 | 14612248 | 81727000.84 | 57246000.4 | 18251048 | 111755310.8 | 23109068.84 | 123651504.9 | |
| P15880 | Small ribosomal subunit protein uS5 [OS=Homo sapiens] | -0.45 | 0.970551984 | 225973.8027 | 1553606.945 | 10412518.88 | 442218.8594 | 9227899.313 | 257257.2813 | 1527834.899 | 11150279.37 | 10412518.88 | 3232554.361 | 9586188.668 | 2176957.985 | |
| P60866 | Small ribosomal subunit protein uS10 [OS=Homo sapiens] | -0.2 | 0.970551984 | 238542.7422 | 1189004.012 | 9379447.563 | 260259.4844 | 7725406.234 | 185582.7734 | 1612814.946 | 8533514.186 | 9379447.563 | 1902458.281 | 8025358.664 | 1570435.241 | |
| P08727 | Keratin, type I cytoskeletal 19 [OS=Homo sapiens] | 0.06 | 0.970551984 | 3190241.266 | 1702258.242 | 1466352.5 | 2620643.125 | 3284113.563 | 2440796.875 | 21569588.53 | 12217153.78 | 1466352.5 | 19156513.07 | 3411625.024 | 20654467.86 | |
| P31150 | Rab GDP dissociation inhibitor alpha [OS=Homo sapiens] | 0.35 | 0.970551984 | 187637.2324 | 15737413.94 | 156592875.5 | 343880.8516 | 107611860.1 | 772910.627 | 1268636.933 | 112947848.6 | 156592875.5 | 2513718.089 | 111790079.1 | 6540510.547 | |
| Q99719 | Septin-5 [OS=Homo sapiens] | 0.09 | 0.970551984 | | 2622238.152 | 40815198.94 | | 23658997.77 | 76288.80469 | | 18819874.66 | 40815198.94 | | 24577599.8 | 645569.7648 | |
| P62701 | Small ribosomal subunit protein eS4, X isoform [OS=Homo sapiens] | -0.29 | 0.970551984 | 612686.2109 | 3125437.699 | 20279491.47 | 696940.707 | 17275799.28 | 469773.0352 | 4142442.018 | 22431351.52 | 20279491.47 | 5094533.338 | 17946562.45 | 3975305.015 | |
| P13591 | Neural cell adhesion molecule 1 [OS=Homo sapiens] | -1.09 | 0.970551984 | | 10135910.76 | 86521613.3 | | 65479728.53 | 29537.32617 | | 72745707.66 | 86521613.3 | | 68022093.69 | 249950.2357 | |
| Q9NTJ5 | Phosphatidylinositol-3-phosphatase SAC1 [OS=Homo sapiens] | -0.35 | 0.970551984 | | 444979.8945 | 4381688.469 | 105658.5664 | 1732044.281 | | | 3193632.826 | 4381688.469 | 772348.4703 | 1799293.934 | | |
| P08195 | Amino acid transporter heavy chain SLC3A2 [OS=Homo sapiens] | 0.44 | 0.970551984 | 2253446.227 | 2069405.02 | 7417770.844 | 5312603.555 | 16219034.91 | 1417412.34 | 15235809.4 | 14852176.2 | 7417770.844 | 38834345.08 | 16848767.35 | 11994401.47 | |
| Q12756 | Kinesin-like protein KIF1A [OS=Homo sapiens] | -0.51 | 0.970551984 | 1353807.625 | 279539.8281 | 1626914.656 | 881901.8125 | 977709.375 | 1049084.75 | 9153249.232 | 2006264.962 | 1626914.656 | 6446571.622 | 1015670.654 | 8877546.294 | |
| P11586 | C-1-tetrahydrofolate synthase, cytoplasmic [OS=Homo sapiens] | -0.04 | 0.970551984 | 21262.89648 | 183774.1367 | 1787569.25 | 25010.41797 | 1276312.75 | 29942.58203 | 143760.8914 | 1318951.986 | 1787569.25 | 182822.4508 | 1325867.828 | 253379.5846 | |
| O75390 | Citrate synthase, mitochondrial [OS=Homo sapiens] | -1.28 | 0.970551984 | 1273971.969 | 9666046.176 | 80188215.38 | 585003.8281 | 60318057.81 | 1202895.859 | 8613471.168 | 69373476.75 | 80188215.38 | 4276291.333 | 62660012.07 | 10179123.93 | |
| O00231 | 26S proteasome non-ATPase regulatory subunit 11 [OS=Homo sapiens] | -0.35 | 0.970551984 | | 933993.707 | 7387937.469 | 38170.72656 | 5385122.203 | | | 6703298.281 | 7387937.469 | 279022.3573 | 5594209.006 | | Met-loss+Acetyl [N-Term] |
| O75821 | Eukaryotic translation initiation factor 3 subunit G [OS=Homo sapiens] | 0 | 0.970551984 | | | 307398.5313 | | 179723.2344 | | | | 307398.5313 | | 186701.3038 | | |
| P09104 | Gamma-enolase [OS=Homo sapiens] | 0.14 | 0.970551984 | 160794.2031 | 18296218.39 | 218509507.3 | 164357.4219 | 148326857.6 | 625347.4063 | 1087148.121 | 131312457.8 | 218509507.3 | 1201428.409 | 154085907.7 | 5291803.688 | |
| P60660 | Myosin light polypeptide 6 [OS=Homo sapiens] | -0.04 | 0.970551984 | 295396.3594 | 1064882.641 | 7575285.375 | 309234.125 | 7870270.563 | 298620.2969 | 1997208.798 | 7642691.724 | 7575285.375 | 2260455.65 | 8175847.603 | 2526979.36 | Met-loss+Acetyl [N-Term] |
| Q9UPP2 | IQ motif and SEC7 domain-containing protein 3 [OS=Homo sapiens] | 0 | 0.970551984 | | | 866436.3906 | | 198076.2188 | | | | 866436.3906 | | 205766.8749 | | |
| Q16629 | Serine/arginine-rich splicing factor 7 [OS=Homo sapiens] | -0.01 | 0.970551984 | 2357430.25 | 1142023.602 | 7691039.563 | 1829631.93 | 8897728.641 | 791779.7188 | 15938857.36 | 8196334.502 | 7691039.563 | 13374338.4 | 9243198.541 | 6700184.24 | |
| P61266 | Syntaxin-1B [OS=Homo sapiens] | -1.58 | 0.970551984 | | 13596728.57 | 117224588 | | 77103417.3 | 52875.7207 | | 97584091.41 | 117224588 | | 80097092.53 | 447443.9824 | |
| P48739 | Phosphatidylinositol transfer protein beta isoform [OS=Homo sapiens] | 0 | 0.970551984 | | | 336203.5938 | | 369505.1563 | | | | 336203.5938 | | 383851.8412 | | |
| P30048 | Thioredoxin-dependent peroxide reductase, mitochondrial [OS=Homo sapie | -0.48 | 0.970551984 | 23402.41016 | 1772756 | 9876741.5 | 63367.25391 | 8071096.25 | 56992.51367 | 158226.3897 | 12723118.11 | 9876741.5 | 463205.2401 | 8384470.7 | 482281.0345 | |
| Q96CM8 | Medium-chain acyl-CoA ligase ACSF2, mitochondrial [OS=Homo sapiens] | 0 | 0.970551984 | | | 277476.7813 | | 138389.2969 | | | | 277476.7813 | | 143762.5037 | | |
| O60636 | Tetraspanin-2 [OS=Homo sapiens] | 0 | 0.970551984 | | | 1021326.313 | | 468197.0938 | | | | 1021326.313 | | 486375.6661 | | |
| P13639 | Elongation factor 2 [OS=Homo sapiens] | 0.08 | 0.970551984 | 6095033.75 | 6016727.605 | 35075717.8 | 10638336.64 | 39228158.88 | 3603484.652 | 41209224.97 | 43182217.95 | 35075717.8 | 77764665.09 | 40751260.86 | 30493343.67 | Met-loss [N-Term] |
| P50990 | T-complex protein 1 subunit theta [OS=Homo sapiens] | 0.04 | 0.970551984 | 921259.8086 | 3288419.266 | 25144108.59 | 1521837.289 | 22949181.59 | 619346.0195 | 6228743.641 | 23601074.66 | 25144108.59 | 11124405.17 | 23840223.77 | 5241018.86 | |
| P01023 | Alpha-2-macroglobulin [OS=Homo sapiens] | 0.04 | 0.970551984 | 2231472.852 | 2079314.977 | 1581475.438 | 2666320.227 | 2368129.063 | 1983990.086 | 15087244.88 | 14923300.24 | 1581475.438 | 19490405.92 | 2460075.822 | 16788885.58 | |
| O60814 | Histone H2B type 1-K [OS=Homo sapiens] | -0.09 | 0.970551984 | 8239769.75 | 26205591.75 | 208261826.5 | 8115253.506 | 196144466.1 | 7898909.262 | 55710032.02 | 188078245.9 | 208261826.5 | 59321301.1 | 203760118.6 | 66842009.33 | |
| O95563 | Mitochondrial pyruvate carrier 2 [OS=Homo sapiens] | -1.12 | 0.970551984 | 120380.8594 | 661911.4063 | 5718858.125 | 41655.32422 | 3133734.063 | 141704.5039 | 813908.8511 | 4750556.196 | 5718858.125 | 304494.2501 | 3255406.777 | 1199129.33 | |
| Q99536 | Synaptic vesicle membrane protein VAT-1 homolog [OS=Homo sapiens] | -1.37 | 0.970551984 | | 2500069.941 | 24686853.88 | | 19908521.88 | 45699.66797 | | 17943070.08 | 24686853.88 | | 20681505.11 | 386718.9167 | |
| P47756 | F-actin-capping protein subunit beta [OS=Homo sapiens] | -0.29 | 0.970551984 | 155304.7383 | 2047349.51 | 18048682.59 | 148450.1172 | 15070336.44 | 35983.17578 | 1050033.218 | 14693883.21 | 18048682.59 | 1085148.368 | 15655468.65 | 304496.1895 | Met-loss+Acetyl [N-Term] |
| Q9UQ35 | Serine/arginine repetitive matrix protein 2 [OS=Homo sapiens] | 0.44 | 0.970551984 | 3523558.914 | 529627.9785 | 3709031.18 | 3171039.609 | 7298903.031 | 1597982.379 | 23823187.53 | 3801154.431 | 3709031.18 | 23179829.85 | 7582295.727 | 13522418.04 | |
| P13647 | Keratin, type II cytoskeletal 5 [OS=Homo sapiens] | 0.21 | 0.970551984 | 22101249.86 | 13644590.86 | 28357667.8 | 21895352.87 | 191628719.8 | 34311078.75 | 149429095.1 | 97927600.34 | 28357667.8 | 160051786.3 | 199069040.6 | 290346599.8 | |
| P47755 | F-actin-capping protein subunit alpha-2 [OS=Homo sapiens] | 0.23 | 0.970551984 | 86784.83594 | 1425506.793 | 18143531.13 | 163635.5039 | 13284965.16 | | 586762.2683 | 10230901.09 | 18143531.13 | 1196151.296 | 13800777.2 | | Met-loss+Acetyl [N-Term] |
| Q16531 | DNA damage-binding protein 1 [OS=Homo sapiens] | -0.06 | 0.970551984 | 1196898.281 | 1470554.25 | 12762105.16 | 2245017.762 | 12925428.78 | 680209.8711 | 8092367.092 | 10554207.92 | 12762105.16 | 16410747.31 | 13427281.2 | 5756059.861 | |
| Q9UL26 | Ras-related protein Rab-22A [OS=Homo sapiens] | 0 | 0.970551984 | | | 753244.6563 | | 342851.0938 | | | | 753244.6563 | | 356162.888 | | |
| P55884 | Eukaryotic translation initiation factor 3 subunit B [OS=Homo sapiens] | -0.03 | 0.970551984 | 754933.8594 | 636260.1582 | 4446563.188 | 606767.0234 | 3698627.781 | 354511.4063 | 5104194.747 | 4566456.49 | 4446563.188 | 4435377.067 | 3842233.484 | 2999940.111 | |
| Q6P1M0 | Long-chain fatty acid transport protein 4 [OS=Homo sapiens] | 0 | 0.970551984 | | | 313452.7188 | | 228214.25 | | | | 313452.7188 | | 237075.068 | | |
| Q9P2S2 | Neurexin-2 [OS=Homo sapiens] | 0 | 0.970551984 | | | 461827.625 | | 224247.9688 | | | | 461827.625 | | 232954.7889 | | |
| P40227 | T-complex protein 1 subunit zeta [OS=Homo sapiens] | -0.22 | 0.970551984 | 74752.35352 | 3275568.078 | 24315811.16 | 445746.2109 | 19533247.19 | 72465.67383 | 505409.2692 | 23508841.34 | 24315811.16 | 3258338.778 | 20291659.73 | 613217.7349 | |
| P46777 | Large ribosomal subunit protein uL18 [OS=Homo sapiens] | 0.21 | 0.970551984 | 353633.4336 | 748527.1797 | 8614512.875 | 552123.5039 | 6511616.969 | 253520.6152 | 2390956.362 | 5372199.962 | 8614512.875 | 4035941.033 | 6764441.904 | 2145337.637 | |
| P15311 | Ezrin [OS=Homo sapiens] | -0.09 | 0.970551984 | 169886.7695 | 249880.9648 | 1173039.25 | 210422.2734 | 1692788.438 | 99946.98438 | 1148624.01 | 1793402.493 | 1173039.25 | 1538155.651 | 1758513.914 | 845769.5918 | |
| Q53H96 | Pyrroline-5-carboxylate reductase 3 [OS=Homo sapiens] | 0 | 0.970551984 | | | 313740.1563 | | 207345.1563 | | | | 313740.1563 | | 215395.6951 | | |
| Q71U36 | Tubulin alpha-1A chain [OS=Homo sapiens] | 0.08 | 0.970551984 | 6431259.492 | 256411282.4 | 2737523059 | 6565927.475 | 1816206457 | 12137065.21 | 43482485.93 | 1840270759 | 2737523059 | 47995957.29 | 1886723854 | 102706057.2 | |
| P32119 | Peroxiredoxin-2 [OS=Homo sapiens] | -0.71 | 0.970551984 | 163022.9375 | 1432257.875 | 10475650.75 | 134042.918 | 6029495 | 169285.3047 | 1102216.851 | 10279353.79 | 10475650.75 | 979833.8759 | 6263600.705 | 1432523.092 | |
| Q04837 | Single-stranded DNA-binding protein, mitochondrial [OS=Homo sapiens] | -0.54 | 0.970551984 | 64333.07031 | 187822.1406 | 1193797.25 | 64438.07813 | 894530.4375 | 53715.625 | 434963.2958 | 1348004.62 | 1193797.25 | 471032.8065 | 929262.1487 | 454551.4056 | |
| P07900 | Heat shock protein HSP 90-alpha [OS=Homo sapiens] | -0.01 | 0.970551984 | 13614537.2 | 25387104.26 | 160806369.8 | 14138927.79 | 158182372.4 | 7342669.898 | 92049453.6 | 182203938.9 | 160806369.8 | 103353467.9 | 164324080.1 | 62135010.49 | |
| P00167 | Cytochrome b5 [OS=Homo sapiens] | -0.01 | 0.970551984 | 596602.8125 | 713856.125 | 2859087.75 | 410079.0625 | 1921754.688 | 574804.8594 | 4033700.309 | 5123364.858 | 2859087.75 | 2997617.206 | 1996370.18 | 4864103.448 | |
| Q9ULF5 | Zinc transporter ZIP10 [OS=Homo sapiens] | 0.14 | 0.970551984 | | 52119.71875 | 423566.0938 | 56649.38281 | 395737.6875 | | | 374064.6414 | 423566.0938 | 414098.5975 | 411102.8964 | | |
| P02679 | Fibrinogen gamma chain [OS=Homo sapiens] | 0.07 | 0.970551984 | 1581715.336 | 1485725.422 | 4676048.25 | 1759419.969 | 9357869.281 | 1721095.787 | 10694159.51 | 10663091.83 | 4676048.25 | 12861099.37 | 9721204.948 | 14564226.13 | |
| Q16836 | Hydroxyacyl-coenzyme A dehydrogenase, mitochondrial [OS=Homo sapien: | -0.86 | 0.970551984 | 155501.7188 | 356569.6406 | 1662506.688 | 181787.4043 | 1344555.313 | 131325.543 | 1051365.025 | 2559110.025 | 1662506.688 | 1328838.999 | 1396760.028 | 1111300.672 | |
| P55786 | Puromycin-sensitive aminopeptidase [OS=Homo sapiens] | -0.09 | 0.970551984 | 79128.66406 | 3586641.076 | 35638719.72 | 118377.0469 | 27217077.44 | 44628.08984 | 534998.0087 | 25741420.72 | 35638719.72 | 865318.6787 | 28273828.14 | 377651.0274 | |
| P00558 | Phosphoglycerate kinase 1 [OS=Homo sapiens] | 0.03 | 0.970551984 | 874036.5791 | 19171729.12 | 182232131.9 | 1325901.779 | 140762178.9 | 1373604.57 | 5909461.949 | 137596022.2 | 182232131.9 | 9692145.614 | 146227517.1 | 11623692.14 | Met-loss+Acetyl [N-Term] |
| P00966 | Argininosuccinate synthase [OS=Homo sapiens] | -0.67 | 0.970551984 | 812414.4668 | 2045458.988 | 11301478.91 | 923220.0059 | 7895286.844 | 691060.2402 | 5492827.753 | 14680314.89 | 11301478.91 | 6748601.496 | 8201835.186 | 5847877.662 | |
| P63104 | 14-3-3 protein zeta/delta [OS=Homo sapiens] | 0.2 | 0.970551984 | 419131.4961 | 30221261.16 | 237703135 | 746048.1719 | 205428058.8 | 477974.2227 | 2833796.304 | 216898814.7 | 237703135 | 5453501.632 | 213404163 | 4044704.957 | Acetyl [N-Term] |
| P60174 | Triosephosphate isomerase [OS=Homo sapiens] | 0.62 | 0.970551984 | 720449.2227 | 14071957.13 | 159187837.7 | 1155329.469 | 105438231.3 | 1100128.398 | 4871040.148 | 100994819.7 | 159187837.7 | 8445287.289 | 109532055.3 | 9309486.947 | |
| P40429 | Large ribosomal subunit protein uL13 [OS=Homo sapiens] | -0.18 | 0.970551984 | 272831.874 | 1431361.246 | 8778911 | 372254.8438 | 8816132.875 | 203701.5615 | 1844647.715 | 10272918.66 | 8778911 | 2721127.769 | 9158434.677 | 1723759.727 | |
| P09429 | High mobility group protein B1 [OS=Homo sapiens] | 0.18 | 0.970551984 | 1911286.057 | 2544117.703 | 24814868.81 | 2059515.508 | 22738918.03 | 1221104.777 | 12922425.1 | 18259202.07 | 24814868.81 | 15054753.31 | 23621796.36 | 10333211.11 | |
| Q9NP72 | Ras-related protein Rab-18 [OS=Homo sapiens] | 0.18 | 0.970551984 | | 294838.0781 | 3614687.313 | 74151.90625 | 3109509.469 | | | 2116060.919 | 3614687.313 | 542039.4513 | 3230241.621 | | |
| P51659 | Peroxisomal multifunctional enzyme type 2 [OS=Homo sapiens] | 0.14 | 0.970551984 | 205540.375 | 270829.4375 | 2095416.688 | 258943.418 | 1606330.156 | 175051.4531 | 1389682.142 | 1943750.252 | 2095416.688 | 1892838.031 | 1668698.738 | 1481317.29 | |
| P69905 | Hemoglobin subunit alpha [OS=Homo sapiens] | -0.04 | 0.970551984 | 12629811.29 | 35980010.19 | 181269049.3 | 14637785.09 | 146459981 | 13926976.63 | 85391608.34 | 258229513.4 | 181269049.3 | 107000040.8 | 152146546.3 | 117852613.7 | Met-loss [N-Term] |
| P68104 | Elongation factor 1-alpha 1 [OS=Homo sapiens] | 0.17 | 0.970551984 | 35037774.23 | 34489737.67 | 234532724.7 | 45312035.52 | 252014417.4 | 20648731.72 | 236894425.9 | 247533786.9 | 234532724.7 | 331224268.1 | 261799318.6 | 174733329.9 | |
| Q13885 | Tubulin beta-2A chain [OS=Homo sapiens] | 0.07 | 0.970551984 | 1743446.283 | 37871235.24 | 639198757.8 | 1180445.5 | 333551965.4 | 2617148.088 | 11787641.06 | 271802887.1 | 639198757.8 | 8628881.757 | 346502704.7 | 22146783.96 | |
| Q16658 | Fascin [OS=Homo sapiens] | 0.41 | 0.970551984 | 2004949.117 | 1670066.605 | 16970508.69 | 2675026.23 | 15387705.17 | 1017563.762 | 13555691.83 | 11986113.53 | 16970508.69 | 19554045.52 | 15985159.78 | 8610809.949 | |
| P27144 | Adenylate kinase 4, mitochondrial [OS=Homo sapiens] | -0.69 | 0.970551984 | 24489.41992 | 105138.7188 | 450665.2813 | | 450176.2188 | 34140.26953 | 165575.7878 | 754583.4489 | 450665.2813 | | 467655.099 | 288901.181 | |
| P78371 | T-complex protein 1 subunit beta [OS=Homo sapiens] | 0.11 | 0.970551984 | 76268.82813 | 2589946.129 | 28530118.09 | 220943.0117 | 22746364.13 | | 515662.3287 | 18588113.93 | 28530118.09 | 1615060.69 | 23629531.56 | | Met-loss+Acetyl [N-Term] |
| Q02338 | D-beta-hydroxybutyrate dehydrogenase, mitochondrial [OS=Homo sapiens] | -2.86 | 0.970551984 | | 73685.36719 | 2654765.281 | | 1292448.125 | | | 528841.8878 | 2654765.281 | | 1342629.687 | | |
| Q99943 | 1-acyl-sn-glycerol-3-phosphate acyltransferase alpha [OS=Homo sapiens] | 0 | 0.970551984 | | | 322612.4531 | | 343581.2813 | | | | 322612.4531 | | 356921.4263 | | |
| O95822 | Malonyl-CoA decarboxylase, mitochondrial [OS=Homo sapiens] | 0 | 0.970551984 | | | 375206 | | 227436.9844 | | | | 375206 | | 236267.6237 | | |
| Q02252 | Methylmalonate-semialdehyde/malonate-semialdehyde dehydrogenase [ac | -0.53 | 0.970551984 | 817607.0195 | 1962797.227 | 10942251.16 | 892384.4922 | 8989910.297 | 880968.2305 | 5527935.199 | 14087049.17 | 10942251.16 | 6523198.458 | 9338959.312 | 7454913.676 | |
| P04899 | Guanine nucleotide-binding protein G(i) subunit alpha-2 [OS=Homo sapiens | 0.4 | 0.970551984 | 603619.3359 | 10943488.05 | 139770427.1 | 968533.1719 | 87801321.67 | 952107.8945 | 4081139.832 | 78541711.86 | 139770427.1 | 7079834.027 | 91210361.79 | 8056910.475 | |
| P04632 | Calpain small subunit 1 [OS=Homo sapiens] | -0.51 | 0.970551984 | 31107.93555 | 186468.5 | 1013142.625 | 20142.23828 | 955533.0625 | 41415.95703 | 210324.334 | 1338289.505 | 1013142.625 | 147236.7784 | 992633.3074 | 350469.3743 | |
| Q15334 | Lethal(2) giant larvae protein homolog 1 [OS=Homo sapiens] | 0 | 0.970551984 | | | 1122197.438 | | 345550.4688 | | | | 1122197.438 | | 358967.071 | | |
| O95140 | Mitofusin-2 [OS=Homo sapiens] | -0.06 | 0.970551984 | 26617.52148 | 101607.3086 | 1352264.313 | 23600.67578 | 803740 | | 179964.1275 | 729238.422 | 1352264.313 | 172517.4442 | 834946.6134 | | |
| P0DTE7 | Alpha-amylase 1B [OS=Homo sapiens] | 0.35 | 0.970551984 | 31534.50391 | 93275.78125 | 4345502.781 | | 2193593379 | 857987.4961 | 213208.4118 | 669442.8233 | 4345502.781 | | 2278763485 | 7260446.515 | |
| P30086 | Phosphatidylethanolamine-binding protein 1 [OS=Homo sapiens] | 0 | 0.970551984 | 102437.9844 | 3314475.422 | 31589388.94 | 139151.9063 | 21854477.84 | 114680.1719 | 692595.0072 | 23788080.4 | 31589388.94 | 1017179.824 | 22703016.23 | 970444.5088 | |
| P31689 | DnaJ homolog subfamily A member 1 [OS=Homo sapiens] | 0.04 | 0.970551984 | 22726.06836 | 607729.9297 | 6542837.875 | 37068.42578 | 3949811.813 | 95864.68359 | 153653.5649 | 4361694.263 | 6542837.875 | 270964.7019 | 4103170.175 | 811224.4188 | Met-loss [N-Term] |
| P61962 | DDB1- and CUL4-associated factor 7 [OS=Homo sapiens] | 0.26 | 0.970551984 | | 263338.1367 | 2369985.156 | 11291.05371 | 1990924.75 | 22184.81055 | | 1889984.982 | 2369985.156 | 82535.9302 | 2068225.891 | 187731.9088 | |
| P22033 | Methylmalonyl-CoA mutase, mitochondrial [OS=Homo sapiens] | 0 | 0.970551984 | | | 596007.5625 | | 383532.25 | | | | 596007.5625 | | 398423.5614 | | |
| O00139 | Kinesin-like protein KIF2A [OS=Homo sapiens] | 0.3 | 0.970551984 | 179979.6406 | 956085.1641 | 7587738.063 | 296508.8125 | 5548176.758 | 129368.3516 | 1216863.073 | 6861849.27 | 7587738.063 | 2167435.5 | 5763594.439 | 1094738.561 | |

| | | | | | | | | | | | | | | | | |
|---|---|---|---|---|---|---|---|---|---|---|---|---|---|---|---|---|
| P00325 | All-trans-retinol dehydrogenase [NAD(+)] ADH1B [OS=Homo sapiens] | -0.66 | 0.970594536 | 215126.0781 | 243178.8594 | 2165077.938 | 125740.625 | 1756297.125 | | 1454492.184 | 1745301.299 | 2165077.938 | 919145.3441 | 1824488.437 | | |
| P08237 | ATP-dependent 6-phosphofructokinase, muscle type [OS=Homo sapiens] | -0.11 | 0.970594536 | 630000.168 | 6359886.176 | 61660698.16 | 1002514.191 | 47053962.84 | 215917.2207 | 4259503.675 | 45645076.35 | 61660698.16 | 7328230.247 | 48880915.38 | 1827130.861 | |
| P62136 | Serine/threonine-protein phosphatase PP1-alpha catalytic subunit [OS=Hor | -0.11 | 0.970793328 | | 1288193.408 | 18178747.38 | 31710.76758 | 12587284.78 | 11856.7168 | | 9245399.187 | 18178747.38 | 231801.014 | 13076008.16 | 100333.6978 | |
| Q9Y697 | Cysteine desulfurase [OS=Homo sapiens] | -0.13 | 0.970793328 | | 148667.2813 | 1332863.844 | 50072.15625 | 985238.7813 | | | 1066989.128 | 1332863.844 | 366020.0455 | 1023492.403 | | |
| P62861 | Ubiquitin-like FUBI-ribosomal protein eS30 fusion protein [OS=Homo sapier | -0.56 | 0.971269559 | 119301.3672 | 209692.5039 | 1544500.313 | 74693.57813 | 741486.375 | 56951.51563 | 806610.2801 | 1504968.813 | 1544500.313 | 545998.9925 | 770275.8824 | 481934.1016 | |
| Q5T749 | Keratinocyte proline-rich protein [OS=Homo sapiens] | -0.12 | 0.971327473 | 534737.9219 | 277579.0625 | 526267.0938 | 394480.5938 | 1078073.281 | 520428.5547 | 3615424.66 | 1992192.494 | 526267.0938 | 2883594.71 | 1119931.365 | 4403961.25 | |
| P49755 | Transmembrane emp24 domain-containing protein 10 [OS=Homo sapiens] | -0.14 | 0.971881815 | 216134.5938 | 592291.5 | 3986014.609 | 492557.6328 | 4197138.531 | 200368.3672 | 1461310.874 | 4250892.232 | 3986014.609 | 3600523.338 | 4360099.787 | 1695553.63 | |
| P46821 | Microtubule-associated protein 1B [OS=Homo sapiens] | -0.12 | 0.971881815 | 51715.10547 | 4918484.305 | 56553705.19 | 96635.98438 | 35170129.45 | 34181.11328 | 349651.7826 | 35300095.86 | 56553705.19 | 706394.7321 | 36535671.34 | 289246.8083 | |
| Q9HB71 | Calcyclin-binding protein [OS=Homo sapiens] | -0.18 | 0.971881815 | | 373408.6563 | 4454742.625 | 63283.55469 | 2537157.844 | 47023.34766 | | 2679964.099 | 4454742.625 | 462593.4112 | 2635667.441 | 397920.1354 | |
| Q12906 | Interleukin enhancer-binding factor 3 [OS=Homo sapiens] | -0.16 | 0.971881815 | 43766.4375 | 765779.3281 | 3583645.5 | 118359.9141 | 131135688.4 | 35019.42188 | 295909.9233 | 5496019.101 | 3583645.5 | 865193.4403 | 136227261.2 | 296340.7283 | |
| P62851 | Small ribosomal subunit protein eS25 [OS=Homo sapiens] | -0.55 | 0.971881815 | 295172.1797 | 1277927.273 | 6133058.625 | 248121.2891 | 6789868.063 | 258437.9688 | 1995693.093 | 9171718.858 | 6133058.625 | 1813729.871 | 7053496.584 | 2186949.178 | |
| P34932 | Heat shock 70 kDa protein 4 [OS=Homo sapiens] | -0.13 | 0.971881815 | 156398.4336 | 4422234.957 | 35545111.81 | 297346.5098 | 25715844.58 | 151148.8691 | 1057427.817 | 31738500.77 | 35545111.81 | 2173558.943 | 26714307.29 | 1279049.27 | |
| P53621 | Coatomer subunit alpha [OS=Homo sapiens] | -0.17 | 0.971881815 | | 250492.6152 | 2847205.438 | 63854.27148 | 2761013.406 | | | 1797792.324 | 2847205.438 | 466765.2665 | 2868214.588 | | |
| P20618 | Proteasome subunit beta type-1 [OS=Homo sapiens] | -0.15 | 0.971881815 | 96646.72656 | 1181608.547 | 10111716.19 | 137831.1406 | 8751664 | 115697.7813 | 653439.6463 | 8480436.733 | 10111716.19 | 1007525.222 | 9091462.685 | 979055.7047 | |
| P22626 | Heterogeneous nuclear ribonucleoproteins A2/B1 [OS=Homo sapiens] | -0.12 | 0.971881815 | 1682532.523 | 11242397.7 | 300996921.7 | 2085434.215 | 75395587.56 | 1584387.023 | 11375796.12 | 80686994.61 | 300996921.7 | 15244215.22 | 78322953.32 | 13407371.66 | |
| Q86VP6 | Cullin-associated NEDD8-dissociated protein 1 [OS=Homo sapiens] | -0.13 | 0.971881815 | 178745.998 | 5198369.859 | 48172008.88 | 303522.5898 | 37782995.53 | 141959.6289 | 1208522.274 | 37308842.11 | 48172008.88 | 2218705.174 | 39249986.52 | 1201288.244 | Met-loss+Acetyl [N-Term] |
| P09651 | Heterogeneous nuclear ribonucleoprotein A1 [OS=Homo sapiens] | -0.12 | 0.971881815 | 1998129.379 | 2438578.033 | 28212174.19 | 2396091.008 | 21357206.91 | 1407100.947 | 13509582.8 | 17501740.98 | 28212174.19 | 17515070.36 | 22186437.88 | 11907144.59 | |
| P45974 | Ubiquitin carboxyl-terminal hydrolase 5 [OS=Homo sapiens] | -0.13 | 0.972564997 | | 1387225.117 | 17540323.5 | 42999.61719 | 12703797.94 | 100886.8672 | | 9956152.46 | 17540323.5 | 314320.8325 | 13197045.15 | 853723.0514 | Met-loss+Acetyl [N-Term] |
| P60842 | Eukaryotic initiation factor 4A-I [OS=Homo sapiens] | -0.16 | 0.972679046 | 92938.40625 | 862060.8906 | 6217049.594 | 281577.2021 | 5381451.063 | 96242.41406 | 628367.2657 | 6187034.498 | 6217049.594 | 2058287.64 | 5590395.327 | 814420.8429 | |
| Q9BXY0 | Protein MAK16 homolog [OS=Homo sapiens] | -0.31 | 0.972679046 | 1217418.281 | 111498.9219 | 576356.0469 | 906440.2188 | 1406506.938 | 444435.3281 | 8231105.175 | 800230.8 | 576356.0469 | 6625943.737 | 1461117.033 | 3760892.721 | |
| Q06033 | Inter-alpha-trypsin inhibitor heavy chain H3 [OS=Homo sapiens] | -0.17 | 0.972679046 | 108130.7227 | 312899.6328 | 3728173.313 | 133293.7773 | 2358759.875 | | 731084.2662 | 2245689.189 | 3728173.313 | 974357.7689 | 2450342.859 | | |
| P23396 | Small ribosomal subunit protein uS3 [OS=Homo sapiens] | -0.15 | 0.973381193 | 764160.293 | 3341266.652 | 24084865.56 | 1089926.441 | 22450951.56 | 546442.4199 | 5166575.727 | 23980361.79 | 24084865.56 | 7967200.847 | 23322649.08 | 4624095.317 | |
| P0DP23 | Calmodulin-1 [OS=Homo sapiens] | -0.15 | 0.973381193 | 243891.0898 | 10467290.9 | 103214123.6 | 473150.0313 | 61857448.5 | 544916.6719 | 1648975.74 | 75124031.95 | 103214123.6 | 3458656.645 | 64259172.29 | 4611184.159 | Met-loss+Acetyl [N-Term] |
| Q96IX5 | ATP synthase membrane subunit K, mitochondrial [OS=Homo sapiens] | -0.26 | 0.973381193 | 140353.9688 | 1078084.195 | 7237665.563 | 142974.7969 | 6134713.063 | 111739.7109 | 948949.3433 | 7737439.641 | 7237665.563 | 1045124.588 | 6372904.043 | 945561.7926 | |
| Q92841 | Probable ATP-dependent RNA helicase DDX17 [OS=Homo sapiens] | -0.3 | 0.973381193 | 414426.0596 | 1025868.063 | 10618608.63 | 414248.1055 | 8638752.313 | 93431.14844 | 2801982.306 | 7362683.034 | 10618608.63 | 3028092.293 | 8974167.003 | 790631.4009 | |
| P36957 | Dihydrolipoyllysine-residue succinyltransferase component of 2-oxoglutara | -0.17 | 0.973381193 | 139353.4063 | 901116.209 | 8890843.406 | 186905.7813 | 5242512 | 162601.8906 | 942184.425 | 6467335.582 | 8890843.406 | 1366253.576 | 5446061.712 | 1375966.824 | |
| P12036 | Neurofilament heavy polypeptide [OS=Homo sapiens] | -0.14 | 0.973381193 | | 1243690.766 | 14290693.09 | 30899.22852 | 9164073.219 | 104019.0391 | | 8926002.508 | 14290693.09 | 225868.7837 | 9519884.414 | 880228.0605 | |
| P40926 | Malate dehydrogenase, mitochondrial [OS=Homo sapiens] | -0.15 | 0.973381193 | 1170418.141 | 28238574.73 | 277611316.4 | 1060805.316 | 187033093.2 | 1894355.908 | 7913331.812 | 202669020.2 | 277611316.4 | 7754329.736 | 194294980.7 | 16030384.84 | |
| P26368 | Splicing factor U2AF 65 kDa subunit [OS=Homo sapiens] | -0.64 | 0.973381193 | | 122222.25 | 1158725.703 | 54203.24414 | 1232936.313 | | | 877192.4181 | 1158725.703 | 396217.6861 | 1280807.225 | | |
| P10599 | Thioredoxin [OS=Homo sapiens] | -0.13 | 0.973381193 | 339696.8398 | 772564.1641 | 4000655.984 | 234125.0039 | 9020020.75 | 280512.8809 | 2296729.447 | 5544714.053 | 4000655.984 | 1711419.1 | 9370238.856 | 2373751.106 | |
| P53041 | Serine/threonine-protein phosphatase 5 [OS=Homo sapiens] | -0.16 | 0.973381193 | | 228745.5977 | 3186970.203 | 40596.86328 | 2794540.531 | 34881.05078 | | 1641713.386 | 3186970.203 | 296757.0574 | 2903043.463 | 295169.8069 | |
| P62241 | Small ribosomal subunit protein eS8 [OS=Homo sapiens] | -0.44 | 0.973761878 | 205810.1406 | 1506694.984 | 11098930.13 | 251451.7266 | 7968856.75 | | 1391506.058 | 10813590.95 | 11098930.13 | 1838074.876 | 8278261.572 | | |
| P17844 | Probable ATP-dependent RNA helicase DDX5 [OS=Homo sapiens] | -0.18 | 0.974137818 | 70018.85938 | 702059.5107 | 6739212.594 | 253907.2734 | 4485327.219 | 67983.69531 | 473405.5703 | 5038700.236 | 6739212.594 | 1856024.56 | 4659477.905 | 575290.4161 | |
| P10515 | Dihydrolipoyllysine-residue acetyltransferase component of pyruvate dehyd | -0.16 | 0.974265056 | 512908.4688 | 3776252.5 | 34466774.81 | 248104.7383 | 27306266.38 | 146250.918 | 3467833.214 | 27102267.07 | 34466774.81 | 1813608.888 | 28366480 | 1237601.914 | |
| P43243 | Matrin-3 [OS=Homo sapiens] | -0.16 | 0.974314026 | | 2422296.234 | 18099402.38 | 413624.6406 | 16476273.25 | 58746.14453 | | 17384886.07 | 18099402.38 | 3023534.857 | 17115993.42 | 497120.5784 | |
| P11142 | Heat shock cognate 71 kDa protein [OS=Homo sapiens] | -0.17 | 0.974977753 | 6971668.359 | 68524969.08 | 572662474 | 10772288.05 | 445926074.5 | 5747630.871 | 47136252.51 | 491805570 | 572662474 | 78743830.07 | 463239935.4 | 48637499.63 | |
| P13637 | Sodium/potassium-transporting ATPase subunit alpha-3 [OS=Homo sapien | -0.17 | 0.974977753 | 4452127.301 | 134603302.6 | 1377737037 | 8190229.781 | 887373573.2 | 6375358.708 | 30101345.3 | 966051569.8 | 1377737037 | 59869366.56 | 921827406.6 | 53949446.96 | |
| Q7L014 | Probable ATP-dependent RNA helicase DDX46 [OS=Homo sapiens] | -0.2 | 0.974977753 | 838465.5156 | 70354.22656 | 558288.7109 | 754112.4141 | 955620.25 | 373344.5469 | 5668961.893 | 504934.2008 | 558288.7109 | 5512450.048 | 992723.8802 | 3159309.578 | |
| Q16643 | Drebrin [OS=Homo sapiens] | -0.34 | 0.974977753 | 383003.7695 | 939574.8828 | 11213105.06 | 343252.9336 | 6142365.75 | 491971.9365 | 2589532.585 | 6743354.532 | 11213105.06 | 2509128.102 | 6380853.86 | 4163156.163 | Met-loss+Acetyl [N-Term] |
| O75891 | Cytosolic 10-formyltetrahydrofolate dehydrogenase [OS=Homo sapiens] | -0.64 | 0.974977753 | 39994.69141 | 735575.5195 | 6562907.469 | 50276.59766 | 3330907.25 | 42391.76172 | 270408.7136 | 5279245.544 | 6562907.469 | 367514.4819 | 3460235.559 | 358726.8113 | |
| O43242 | 26S proteasome non-ATPase regulatory subunit 3 [OS=Homo sapiens] | -0.19 | 0.974977753 | 124722.6426 | 910438.125 | 8380023.578 | 180492.6016 | 5788878.359 | 143232.1094 | 843264.1472 | 6534239.227 | 8380023.578 | 1319374.183 | 6013641.702 | 1212056.206 | |
| P50213 | Isocitrate dehydrogenase [NAD] subunit alpha, mitochondrial [OS=Homo sa | -0.17 | 0.974977753 | 113910.4258 | 7003162.234 | 92517191.39 | 195302.9316 | 57513019.09 | 443333.416 | 770161.5045 | 50261886.15 | 92517191.39 | 1427635.502 | 59746062.8 | 3751568.14 | |
| P08670 | Vimentin [OS=Homo sapiens] | -0.18 | 0.975584489 | 3223875.426 | 7528271.52 | 54132604 | 1788134.348 | 42767439.88 | 1557919.5 | 21796992.96 | 54030609.8 | 54132604 | 13070997.22 | 44427960.64 | 13183398.66 | |
| P23526 | Adenosylhomocysteinase [OS=Homo sapiens] | -0.26 | 0.975751049 | 1132518.555 | 2600045.578 | 16516347.94 | 1749596.617 | 15436646.5 | 635147.7305 | 7657088.348 | 18660597.95 | 16516347.94 | 12789292.12 | 16036001.34 | 5374735.817 | Met-loss+Acetyl [N-Term] |
| P36542 | ATP synthase subunit gamma, mitochondrial [OS=Homo sapiens] | -0.19 | 0.975751049 | 756861.6719 | 3232714.914 | 28124302.38 | 740615.125 | 17831614.69 | 766711.1563 | 5117228.91 | 23201283.01 | 28124302.38 | 5413786.864 | 18523958.36 | 6488049.496 | |
| P61981 | 14-3-3 protein gamma [OS=Homo sapiens] | -0.23 | 0.975751049 | 219777.8633 | 36394415.99 | 194905207.2 | 563568.6172 | 176312753.6 | 180498.1992 | 1485943.439 | 261203715.1 | 194905207.2 | 4119603.116 | 183158405.1 | 1527408.648 | Met-loss+Acetyl [N-Term] |
| Q14240 | Eukaryotic initiation factor 4A-II [OS=Homo sapiens] | -0.26 | 0.975751049 | 814557.7344 | 5538056.703 | 39333053.72 | 899707.1172 | 34193863.81 | 784676.9297 | 5507318.631 | 39746783.84 | 39333053.72 | 6576725.762 | 35521500.47 | 6640079.145 | |
| Q9NR31 | GTP-binding protein SAR1a [OS=Homo sapiens] | -0.2 | 0.975751049 | 57452.88672 | 698335.4922 | 6153934.125 | 52281.42188 | 4294941.938 | 55618.84766 | 388445.5823 | 5011972.853 | 6153934.125 | 382169.45 | 4461700.582 | 470656.8224 | |
| P07305 | Histone H1.0 [OS=Homo sapiens] | -0.84 | 0.975751049 | 494777.0137 | 2568804.969 | 18968639.09 | 641578.4063 | 13931371.25 | 722308.1914 | 3345244.359 | 18436383.25 | 18968639.09 | 4689843.119 | 14472281.14 | 6112303.518 | Met-loss+Acetyl [N-Term] |
| P84243 | Histone H3.3 [OS=Homo sapiens] | -0.26 | 0.975751049 | 13876864.06 | 48285834.77 | 246703076.3 | 12844156.98 | 271716866.7 | 11829864.06 | 93823075.73 | 346548751.6 | 246703076.3 | 93888885.05 | 282266750 | 100106465.1 | |
| O00264 | Membrane-associated progesterone receptor component 1 [OS=Homo sapi | -0.23 | 0.975751049 | | 2414797.211 | 19421056.13 | 72877.51563 | 14267730 | | | 17331065.37 | 19421056.13 | 532723.8446 | 14821699.61 | | |
| Q9UEY8 | Gamma-adducin [OS=Homo sapiens] | -0.21 | 0.975751049 | 79492.04688 | 416541.5781 | 2879166.25 | 80778.67969 | 2641015.188 | 102880.1758 | 537454.8817 | 2989530.255 | 2879166.25 | 590480.1836 | 2743557.228 | 870590.7919 | |
| P46439 | Glutathione S-transferase Mu 5 [OS=Homo sapiens] | -0.25 | 0.975751049 | 330503.5156 | 4543288 | 30431237.63 | 324737.5156 | 27735586.81 | 329568.1563 | 2234572.324 | 32607301.75 | 30431237.63 | 2373783.139 | 28812469.55 | 2788865.784 | |
| O75746 | Electrogenic aspartate/glutamate antiporter SLC25A12, mitochondrial [OS= | -0.25 | 0.975751049 | 124260.8008 | 6909531.555 | 59864727.06 | 120281.1719 | 46806488.5 | 111042.8438 | 840141.5816 | 49589896.21 | 59864727.06 | 879237.5504 | 48623832.41 | 939664.7755 | |
| O00408 | cGMP-dependent 3',5'-cyclic phosphodiesterase [OS=Homo sapiens] | -0.22 | 0.975751049 | 51661.52344 | 655182.2852 | 4329961.156 | 124847.6094 | 4399627.75 | 9199.181641 | 349289.5083 | 4702261.111 | 4329961.156 | 912617.5321 | 4570451.004 | 77845.1511 | |
| P04424 | Argininosuccinate lyase [OS=Homo sapiens] | -0.62 | 0.975751049 | | 190331.7734 | 1595931.938 | 26733.92969 | 347963.6875 | 34541.875 | | 1366016.323 | 1595931.938 | 195421.0662 | 361473.9872 | 292299.6397 | |
| P62495 | Eukaryotic peptide chain release factor subunit 1 [OS=Homo sapiens] | -0.59 | 0.975751049 | | 124153.4922 | 1015723.75 | 135409.9531 | 906770.25 | | | 891052.9959 | 1015723.75 | 989826.7004 | 941977.1933 | | |
| P31948 | Stress-induced-phosphoprotein 1 [OS=Homo sapiens] | -0.25 | 0.975751049 | 256034.8047 | 3897254.277 | 28510775.5 | 207619.2773 | 21013892.64 | 218931.1602 | 1731080.795 | 27970700.13 | 28510775.5 | 1517666.165 | 21829793.83 | 1852635.366 | |
| P62913 | Large ribosomal subunit protein uL5 [OS=Homo sapiens] | -0.24 | 0.975751049 | 81074 | 730226.5313 | 4456625.25 | 242697.0078 | 4401114.875 | 82655.10938 | 548150.649 | 5240855.709 | 4456625.25 | 1774079.179 | 4571995.869 | 699442.595 | |
| P16435 | NADPH--cytochrome P450 reductase [OS=Homo sapiens] | -0.74 | 0.975751049 | 808629.752 | 625223.8633 | 5446227.938 | 46180.34961 | 3737939.188 | 16894.24023 | 5467238.86 | 4487248.701 | 5446227.938 | 337571.5154 | 3883071.224 | 142962.1389 | |
| P08758 | Annexin A5 [OS=Homo sapiens] | -0.21 | 0.975751049 | 488306.9453 | 4231234.125 | 30864596.69 | 502720.625 | 25671944.47 | 493092.8311 | 3301499.482 | 30367682.59 | 30864596.69 | 3674813.306 | 26668702.68 | 4172641.376 | |
| P61604 | 10 kDa heat shock protein, mitochondrial [OS=Homo sapiens] | -0.22 | 0.975751049 | 515292.5742 | 5577373.516 | 53954568.06 | 722047.0957 | 39565585.13 | 676193.6953 | 3483952.426 | 40028961.67 | 53954568.06 | 5278057.32 | 41101788.27 | 5722074.25 | |
| P35222 | Catenin beta-1 [OS=Homo sapiens] | -0.22 | 0.975751049 | 81259.85938 | 717287.3359 | 7774861.375 | 112435.3438 | 7257812.828 | 13649.89551 | 549407.2656 | 5147990.752 | 7774861.375 | 821885.7089 | 7539610.125 | 115507.9028 | |
| P54289 | Voltage-dependent calcium channel subunit alpha-2/delta-1 [OS=Homo sa | -1.1 | 0.975751049 | 110950.1484 | 1178638.684 | 11136526.38 | 47811.94531 | 7046509.102 | | 750146.7285 | 8459121.943 | 11136526.38 | 349498.2383 | 7320102.161 | | |
| Q86UY8 | 5'-nucleotidase domain-containing protein 3 [OS=Homo sapiens] | -0.47 | 0.975751049 | 47290.28125 | 61218.34375 | 962372.1563 | | 723049.5625 | 40588.49219 | 319735.0366 | 439365.7209 | 962372.1563 | | 751123.2283 | 343467.216 | |
| P61978 | Heterogeneous nuclear ribonucleoprotein K [OS=Homo sapiens] | -0.2 | 0.975751049 | 746612.2305 | 4862697.531 | 49542282.34 | 987723.9609 | 37748879.22 | 589845.4219 | 5047931.256 | 34899712.66 | 49542282.34 | 7220115.853 | 39214545.58 | 4991379.428 | |
| P24534 | Elongation factor 1-beta [OS=Homo sapiens] | -0.55 | 0.975751049 | 101729.9531 | 435149.8438 | 3201528 | 247855.7266 | 2685459.688 | 57352.99219 | 687807.9264 | 3123082.284 | 3201528 | 1811788.649 | 2789727.364 | 485331.4694 | |
| P11021 | Endoplasmic reticulum chaperone BiP [OS=Homo sapiens] | -0.2 | 0.975751049 | 1503204.012 | 14465710.55 | 98642216.8 | 2125583.221 | 83984751.55 | 1792232.221 | 10163335.4 | 103820798.7 | 98642216.8 | 15537698.5 | 87245606.65 | 15166195.59 | |
| Q9Y6C9 | Mitochondrial carrier homolog 2 [OS=Homo sapiens] | -0.2 | 0.975751049 | 56535.54688 | 597914.375 | 12519881.63 | 45615.08594 | 5932133.25 | 110550.7578 | 382243.342 | 4291247.759 | 12519881.63 | 333439.5217 | 6162458.715 | 935500.6546 | |
| O15145 | Actin-related protein 2/3 complex subunit 3 [OS=Homo sapiens] | -0.33 | 0.975751049 | 49353.87109 | 1298612.656 | 8595567.109 | 85528.55469 | 8070222.406 | 44934.28516 | 333687.2051 | 9320178.414 | 8595567.109 | 625201.0663 | 8383562.928 | 380242.1079 | Met-loss [N-Term] |
| P11216 | Glycogen phosphorylase, brain form [OS=Homo sapiens] | -0.2 | 0.975751049 | | 6135500.414 | 66299462.13 | 84653.83789 | 46614767.16 | 71822.03125 | | 44034653.63 | 66299462.13 | 618807.0161 | 48424667.15 | 607771.1142 | |
| P51991 | Heterogeneous nuclear ribonucleoprotein A3 [OS=Homo sapiens] | -0.21 | 0.975751049 | 3287857.59 | 5015849.068 | 65936906.81 | 3361563.07 | 36957530.19 | 4259429.277 | 22229583.73 | 35998885.42 | 65936906.81 | 24572528.13 | 38392471.04 | 36044066.6 | |
| Q13148 | TAR DNA-binding protein 43 [OS=Homo sapiens] | -0.19 | 0.975751049 | 23160.34766 | 552050.6719 | 4451907.531 | 56437.72656 | 3817582.875 | 43831.95703 | 156589.7772 | 3962082.712 | 4451907.531 | 412551.4216 | 3965807.217 | 370914.0064 | |
| P62753 | Small ribosomal subunit protein eS6 [OS=Homo sapiens] | -0.35 | 0.975751049 | 348438.582 | 847687.8984 | 6304945.125 | 613353.4844 | 4452217.625 | 242687.6797 | 2355833.373 | 6083879.142 | 6304945.125 | 4483523.121 | 4625082.773 | 2053667.362 | |
| Q99623 | Prohibitin-2 [OS=Homo sapiens] | -0.21 | 0.975751049 | 521212.2539 | 4423644.961 | 39071200.38 | 625096.4844 | 29831251.25 | 652441.2705 | 3523976.062 | 31748620.4 | 39071200.38 | 4569362.712 | 30989501.83 | 5521076.904 | |
| P02765 | Alpha-2-HS-glycoprotein [OS=Homo sapiens] | -0.13 | 0.975751049 | 89417.35156 | 95788 | | 76653.43555 | 769025.0313 | | 604561.0096 | 687473.0857 | | 560325.2599 | 798883.7752 | | |
| P27348 | 14-3-3 protein theta [OS=Homo sapiens] | -0.18 | 0.975751049 | 108297.1465 | 6239174.922 | 52288226.06 | 152832.8984 | 41556554.31 | 106696.5977 | 732209.4769 | 44778728.4 | 52288226.06 | 1117185.85 | 43170060.33 | 902886.0491 | |
| Q96A72 | Protein mago nashi homolog 2 [OS=Homo sapiens] | -0.17 | 0.975751049 | | 91229.46875 | 545196.1875 | 69946.10156 | 561235.6875 | | | 654756.3827 | 545196.1875 | 511295.642 | 583026.6462 | | |
| P38159 | RNA-binding motif protein, X chromosome [OS=Homo sapiens] | -0.22 | 0.975751049 | 89177.71875 | 894427.7344 | 8565531.656 | 116173.6406 | 5989049.344 | 71153.42969 | 602940.8245 | 6419332.218 | 8565531.656 | 849212.1053 | 6221584.675 | 602113.2859 | |
| Q99460 | 26S proteasome non-ATPase regulatory subunit 1 [OS=Homo sapiens] | -1.67 | 0.975751049 | | 636110.5977 | 12596018 | | 8492384.484 | 83225.5625 | | 4565383.09 | 12596018 | | 8822116.188 | 704269.8733 | |
| P60228 | Eukaryotic translation initiation factor 3 subunit E [OS=Homo sapiens] | -0.18 | 0.975751049 | | 132391.7695 | 1205743.594 | 45583.91016 | 1081397 | | | 950179.3369 | 1205743.594 | 333211.6314 | 1123384.133 | | |
| O75083 | WD repeat-containing protein 1 [OS=Homo sapiens] | -0.26 | 0.975751049 | 2681939.039 | 3823155.555 | 48299826.16 | 3791713.047 | 37754462.66 | 1566866.172 | 18132898.65 | 27438891.58 | 48299826.16 | 27716860.74 | 39220345.8 | 13259107.02 | |

| | | | | | | | | | | | | | | | | |
|---|---|---|---|---|---|---|---|---|---|---|---|---|---|---|---|---|
| Q9Y570 | Protein phosphatase methylesterase 1 [OS=Homo sapiens] | -1.19 | 0.975751049 | | 340603.8359 | 7212250.125 | | 3176535.906 | 48527.03906 | | 2444523.01 | 7212250.125 | | 3299870.477 | 410644.646 | |
| P61313 | Large ribosomal subunit protein eL15 [OS=Homo sapiens] | -0.2 | 0.975751049 | 60949.96875 | 302875 | 2353792.688 | 226836.1016 | 2441759.313 | 71387.14063 | 412089.7566 | 2173742.127 | 2353792.688 | 1658138.303 | 2536564.895 | 604090.9905 | |
| Q96S97 | Myeloid-associated differentiation marker [OS=Homo sapiens] | -0.23 | 0.975751049 | | 140695.375 | 847781.8125 | 259172.4219 | 830925.75 | | | 1009774.54 | 847781.8125 | 1894512.015 | 863187.8978 | | |
| P78352 | Disks large homolog 4 [OS=Homo sapiens] | -0.32 | 0.975751049 | 51961.24609 | 1250240.488 | 15578819.69 | 46645.73828 | 10981435.52 | 103089.7422 | 351315.9677 | 8973010.046 | 15578819.69 | 340973.4377 | 11407808.99 | 872364.181 | |
| Q04637 | Eukaryotic translation initiation factor 4 gamma 1 [OS=Homo sapiens] | -0.59 | 0.975751049 | 121949.7109 | 552090.6875 | 3354306.938 | 179222.6055 | 2525253.875 | 82627.75 | 824516.0371 | 3962369.906 | 3354306.938 | 1310090.7 | 2623301.28 | 699211.075 | |
| P00505 | Aspartate aminotransferase, mitochondrial [OS=Homo sapiens] | -0.19 | 0.975751049 | 521222.1777 | 8233006.652 | 88028845.78 | 578111.2344 | 56058671.19 | 816456.3242 | 3524043.158 | 59088513.05 | 88028845.78 | 4225907.493 | 58235247.29 | 6909002.171 | |
| P10253 | Lysosomal alpha-glucosidase [OS=Homo sapiens] | -0.16 | 0.975751049 | | 184041.4063 | 1032130.375 | 8814.916016 | 1135367.875 | 15488.02832 | | 1320870.787 | 1032130.375 | 64435.73041 | 1179450.522 | 131062.5175 | |
| P68036 | Ubiquitin-conjugating enzyme E2 L3 [OS=Homo sapiens] | -0.32 | 0.975751049 | 82594.71875 | 1249493.098 | 9581955.125 | 117264.8359 | 8369097.281 | 89630.08594 | 558432.4035 | 8967646.004 | 9581955.125 | 857188.5814 | 8694042.143 | 758466.1175 | |
| P08238 | Heat shock protein HSP 90-beta [OS=Homo sapiens] | -0.22 | 0.975751049 | 38453246.26 | 63265283.29 | 392658575.2 | 51162098.25 | 397103757.8 | 21392647.39 | 259986825.5 | 454056661.8 | 392658575.2 | 373987360.1 | 412522006.8 | 181028479.9 | |
| P07197 | Neurofilament medium polypeptide [OS=Homo sapiens] | -0.18 | 0.975751049 | 161322.2969 | 6386745.73 | 63031223.44 | 157972.75 | 45079629.59 | 435323.4219 | 1090718.624 | 45837848.11 | 63031223.44 | 1154757.404 | 46829925.18 | 3683786.11 | |
| P18621 | Large ribosomal subunit protein uL22 [OS=Homo sapiens] | -0.22 | 0.975751049 | 29082.15234 | 456115.2695 | 5922388.063 | 93368.59766 | 3838795.75 | | 196627.7806 | 3273551.716 | 5922388.063 | 682510.6191 | 3987843.719 | | |
| P24666 | Low molecular weight phosphotyrosine protein phosphatase [OS=Homo sap | -0.14 | 0.975751049 | 21827.16211 | 397519.7813 | 3615667.625 | 29053.39063 | 2194279.531 | 22458.42773 | 147575.9562 | 2853010.3 | 3615667.625 | 212375.9821 | 2279476.278 | 190047.3073 | |
| Q9HAV7 | GrpE protein homolog 1, mitochondrial [OS=Homo sapiens] | -0.15 | 0.975751049 | 38931.96484 | 207293.4219 | 1688880.875 | 38445.37891 | 1154212.094 | 28964.89453 | 263223.497 | 1487750.536 | 1688880.875 | 281030.0253 | 1199026.4 | 245106.2148 | |
| O75340 | Programmed cell death protein 6 [OS=Homo sapiens] | -0.14 | 0.975751049 | 26947.54102 | 239485.625 | 2174418 | 24674.76563 | 1496882 | 36172.42578 | 182195.4276 | 1718794.855 | 2174418 | 180368.8819 | 1555001.066 | 306097.6575 | |
| P17980 | 26S proteasome regulatory subunit 6A [OS=Homo sapiens] | -0.25 | 0.975751049 | | 292614.3125 | 3231012.875 | 22289.38086 | 2402525.906 | | | 2100100.893 | 3231012.875 | 162932.0726 | 2495808.183 | | |
| P13798 | Acylamino-acid-releasing enzyme [OS=Homo sapiens] | -0.15 | 0.975751049 | | 72154.89063 | 423980.7188 | 40024.33203 | 449849.2188 | 34240.36719 | | 517857.6158 | 423980.7188 | 292571.9387 | 467315.4027 | 289748.2256 | |
| Q9Y2A7 | Nck-associated protein 1 [OS=Homo sapiens] | -1.58 | 0.975751049 | | 976173.1523 | 9708242.75 | | 8234518.625 | 16764.45313 | | 7006021.31 | 9708242.75 | | 8554238.235 | 141863.8568 | |
| P13804 | Electron transfer flavoprotein subunit alpha, mitochondrial [OS=Homo sapi | -0.4 | 0.975751049 | 108388.9688 | 1291154.816 | 11631178.38 | 128386.9414 | 7377500.906 | 171103.4414 | 732830.2978 | 9266653.294 | 11631178.38 | 938489.525 | 7663945.302 | 1447908.496 | |
| Q14204 | Cytoplasmic dynein 1 heavy chain 1 [OS=Homo sapiens] | -0.24 | 0.975751049 | 441396.2637 | 12170763.07 | 110016837.9 | 827087.875 | 86049519.56 | 110959.584 | 2984330.962 | 87349898.13 | 110016837.9 | 6045889.858 | 89390542.89 | 938960.2162 | |
| P23284 | Peptidyl-prolyl cis-trans isomerase B [OS=Homo sapiens] | -0.27 | 0.975751049 | 133688.1406 | 494911.3164 | 3501423.313 | 337447.9688 | 2484419.813 | 258518.3164 | 903880.9118 | 3551992.001 | 3501423.313 | 2466694.669 | 2580881.764 | 2187629.095 | |
| P62277 | Small ribosomal subunit protein uS15 [OS=Homo sapiens] | -0.65 | 0.975751049 | 194085.4844 | 1240785.164 | 10024021.19 | 298588.1953 | 7049313.625 | 198195.1055 | 1312234.307 | 8905148.927 | 10024021.19 | 2182635.481 | 7323015.575 | 1677163.093 | |
| O95197 | Reticulon-3 [OS=Homo sapiens] | -0.27 | 0.976412699 | 360742.8125 | 4003601.953 | 26836258.88 | 533463.1836 | 22185961 | 175392.9688 | 2439023.692 | 28733960.29 | 26836258.88 | 3899536.856 | 23047369.8 | 1484207.258 | |
| O75061 | Putative tyrosine-protein phosphatase auxilin [OS=Homo sapiens] | -0.75 | 0.976412699 | 67921.1875 | 928076.1484 | 7725114.344 | 36203.03906 | 5860168.469 | 35650.89453 | 459222.9692 | 6660827.802 | 7725114.344 | 264638.8532 | 6087699.775 | 301684.365 | |
| P28331 | NADH-ubiquinone oxidoreductase 75 kDa subunit, mitochondrial [OS=Hom | -0.3 | 0.977036887 | 187930.2539 | 5425652.961 | 53594746.34 | 428722.4766 | 32665438.19 | 335040.1641 | 1270618.085 | 38940059.12 | 53594746.34 | 3133897.801 | 33933731.04 | 2835170.912 | |
| O60506 | Heterogeneous nuclear ribonucleoprotein Q [OS=Homo sapiens] | -0.23 | 0.977036887 | 262349 | 152902.1094 | 307331.5 | 178562.4688 | 1219120.328 | 105156.0078 | 1773771.796 | 1097382.605 | 307331.5 | 1305265.198 | 1266454.81 | 889849.2972 | |
| Q9HCJ6 | Synaptic vesicle membrane protein VAT-1 homolog-like [OS=Homo sapiens | -0.3 | 0.977036887 | 51013.9375 | 7592541.43 | 67939503.88 | 58011.13281 | 46944232.75 | 45471.06641 | 344911.1053 | 54491876.69 | 67939503.88 | 424052.7883 | 48766924.82 | 384784.4486 | |
| Q9BR76 | Coronin-1B [OS=Homo sapiens] | -0.44 | 0.977036887 | 31497.10938 | 229386.7656 | 1664998.688 | 21467.95508 | 1436897.125 | 19134.23047 | 212955.5831 | 1646315.066 | 1664998.688 | 156927.5718 | 1492687.173 | 161917.3444 | |
| O95299 | NADH dehydrogenase [ubiquinone] 1 alpha subcomplex subunit 10, mitoch | -0.57 | 0.977036887 | | 393334.8125 | 2929679.5 | 95832.75781 | 1824589.125 | | | 2822974.665 | 2929679.5 | 700523.2649 | 1895431.994 | | |
| P17302 | Gap junction alpha-1 protein [OS=Homo sapiens] | -0.55 | 0.977036887 | 28959.14453 | 600827.0547 | 4532916.875 | 26875.47266 | 1303826.813 | | 195796.1106 | 4312152.141 | 4532916.875 | 196455.7243 | 1354450.172 | | Met-loss [N-Term] |
| P05091 | Aldehyde dehydrogenase, mitochondrial [OS=Homo sapiens] | -0.52 | 0.977036887 | 170348.4141 | 1231394.072 | 9371057.188 | 242405.4375 | 5892347.938 | 216323.4844 | 1151745.242 | 8837748.805 | 9371057.188 | 1771947.843 | 6121128.668 | 1830568.738 | |
| P38646 | Stress-70 protein, mitochondrial [OS=Homo sapiens] | -0.31 | 0.977036887 | 764060.4375 | 5081811.336 | 41367012.7 | 430119.6758 | 28335038.28 | 525666.0527 | 5165900.593 | 36472298.41 | 41367012.7 | 3144111.121 | 29435195.78 | 4448281.913 | |
| P05023 | Sodium/potassium-transporting ATPase subunit alpha-1 [OS=Homo sapien | -0.33 | 0.977036887 | 4394472.422 | 8370467.172 | 47436695.16 | 7490651.898 | 43791903.02 | 4018702.082 | 29711534.03 | 60075070.94 | 47436695.16 | 54755555.86 | 45492200.36 | 34006989.21 | |
| P50993 | Sodium/potassium-transporting ATPase subunit alpha-2 [OS=Homo sapien | -0.28 | 0.977036887 | 113641.1328 | 27484334.26 | 283248786.8 | 195858.6875 | 179908145.2 | 229578.2627 | 768340.7837 | 197255815.8 | 283248786.8 | 1431697.995 | 186893394 | 1942733.087 | |
| Q99714 | 3-hydroxyacyl-CoA dehydrogenase type-2 [OS=Homo sapiens] | -0.49 | 0.977036887 | | 157254.7969 | 1037488.375 | 52380.0625 | 639567.875 | 55680.33594 | | 1128621.961 | 1037488.375 | 382890.4983 | 664400.2181 | 471177.1475 | |
| Q15382 | GTP-binding protein Rheb [OS=Homo sapiens] | -0.29 | 0.977036887 | 30768.00781 | 496737.0859 | 3059335.531 | 48583.84375 | 1907857.031 | | 208026.0435 | 3565095.599 | 3059335.531 | 355140.7016 | 1981932.922 | | |
| P06753 | Tropomyosin alpha-3 chain [OS=Homo sapiens] | -0.34 | 0.977036887 | 815566.5859 | 5949361.355 | 43970257.25 | 1193125.004 | 38592305.94 | 849956.8984 | 5514139.593 | 42698728.53 | 43970257.25 | 8721567.053 | 40090719.82 | 7192490.134 | |
| Q13347 | Eukaryotic translation initiation factor 3 subunit I [OS=Homo sapiens] | -0.16 | 0.977036887 | 127009.8066 | 145534.125 | 1213284.063 | 144930.7188 | 1670949.563 | 112138.9395 | 858727.9268 | 1044502.38 | 1213284.063 | 1059422.087 | 1735827.106 | 948940.137 | |
| Q86V81 | THO complex subunit 4 [OS=Homo sapiens] | -0.24 | 0.977036887 | 87674.66406 | 324126.3125 | 1942305.125 | 122549.8594 | 1895237.25 | 54775.12109 | 592778.4987 | 2326263.375 | 1942305.125 | 895821.3199 | 1968823.156 | 463517.0545 | |
| A0A0B4J2D5 | Putative glutamine amidotransferase-like class 1 domain-containing protei | -1.37 | 0.977036887 | | 523096.5156 | 6791720.188 | | 4016506.688 | 48376.42969 | | 3754277.944 | 6791720.188 | | 4172454.595 | 409370.162 | |
| Q6UXD5 | Seizure 6-like protein 2 [OS=Homo sapiens] | -0.18 | 0.977036887 | 58059.24609 | 84133.8125 | 681173.5938 | 92196.96875 | 253512.5625 | 17894.23047 | 392545.2479 | 603830.6645 | 681173.5938 | 673946.1853 | 263355.6318 | 151424.2385 | |
| P30044 | Peroxiredoxin-5, mitochondrial [OS=Homo sapiens] | -0.33 | 0.977036887 | 1466467.5 | 4401043.563 | 33576551 | 1581021.75 | 21325923.81 | 1720740.313 | 9914955.619 | 31586409.55 | 33576551 | 11557034.81 | 22153940.16 | 14561218.04 | |
| P22087 | rRNA 2'-O-methyltransferase fibrillarin [OS=Homo sapiens] | -0.23 | 0.977036887 | 104997.9922 | 351125.1211 | 4608002.781 | 83046.99219 | 3308983.938 | 56748.86328 | 709903.5148 | 2520034.56 | 4608002.781 | 607061.212 | 3437461.035 | 480219.2205 | |
| O14936 | Peripheral plasma membrane protein CASK [OS=Homo sapiens] | -0.36 | 0.977036887 | | 363093.1914 | 3280860.875 | 13949.61719 | 2312017.219 | 15962.31348 | | 2605929.727 | 3280860.875 | 101969.6354 | 2401785.337 | 135076.0049 | |
| P10809 | 60 kDa heat shock protein, mitochondrial [OS=Homo sapiens] | -0.28 | 0.977036887 | 3468394.48 | 20275368.55 | 184836673 | 3188355.051 | 125770809 | 2494946.689 | 23450214.44 | 145516872.5 | 184836673 | 23306403.16 | 130654080.9 | 21112693.46 | |
| P61764 | Syntaxin-binding protein 1 [OS=Homo sapiens] | -0.81 | 0.977036887 | 444393125.1 | 30277819.52 | 698906806.4 | 233922370 | 1196855830 | 230911467.6 | 3004593087 | 217304735.6 | 698906806.4 | 1709937876 | 1243325854 | 1954014910 | |
| O43865 | S-adenosylhomocysteine hydrolase-like protein 1 [OS=Homo sapiens] | -0.34 | 0.977036887 | 180966.1797 | 7466047.736 | 67382972.41 | 246138.1406 | 45101557.11 | 130890.9492 | 1223533.178 | 53584028.01 | 67382972.41 | 1799233.35 | 46852704.07 | 1107623.06 | |
| Q16799 | Reticulon-1 [OS=Homo sapiens] | -0.3 | 0.977036887 | | 8236960.406 | 38419578.25 | 80438.27344 | 39878607.81 | 40937.32813 | | 59116889.25 | 38419578.25 | 587991.8643 | 41426964.62 | 346419.1292 | |
| P26373 | Large ribosomal subunit protein eL13 [OS=Homo sapiens] | -0.4 | 0.977036887 | 361605.1563 | 934869.25 | 4059510.938 | 239276.7656 | 4383765.375 | 276934.3672 | 2444854.097 | 6709582.077 | 4059510.938 | 1749077.715 | 4553972.745 | 2343469.072 | |
| Q8WUM4 | Programmed cell death 6-interacting protein [OS=Homo sapiens] | -0.29 | 0.977036887 | 13856.89258 | 798220.3906 | 6064492.406 | 26524.20703 | 4394802.5 | 16163.87207 | 93688.04622 | 5728849.491 | 6064492.406 | 193888.0246 | 4565438.405 | 136781.6305 | |
| P48047 | ATP synthase subunit O, mitochondrial [OS=Homo sapiens] | -0.32 | 0.977036887 | 586873.9375 | 3551394.125 | 31645853.44 | 686773.0625 | 20574595.5 | 570441.9375 | 3967922.265 | 25488452.39 | 31645853.44 | 5020209.362 | 21373440.21 | 4827183.608 | |
| P62854 | Small ribosomal subunit protein eS26 [OS=Homo sapiens] | -0.38 | 0.977036887 | 444421.4063 | 900317.7656 | 6382452.125 | 218363.5625 | 5869514.375 | 205803.7891 | 3004784.299 | 6461605.132 | 6382452.125 | 1596205.298 | 6097408.552 | 1741549.16 | |
| Q9UBQ7 | Glyoxylate reductase/hydroxypyruvate reductase [OS=Homo sapiens] | -0.26 | 0.977036887 | 40449.73047 | 732429.625 | 4278808 | 46721.14453 | 4202789.375 | 55527.96484 | 273485.2851 | 5256667.373 | 4278808 | 341524.6462 | 4365970.034 | 469887.7555 | |
| Q05193 | Dynamin-1 [OS=Homo sapiens] | -0.28 | 0.977036887 | 520108.582 | 22066652.38 | 190322101.8 | 483302.6563 | 135889201 | 902703.3086 | 3516514.009 | 158372965.3 | 190322101.8 | 3532870.83 | 141165337.3 | 7638839.867 | |
| P20674 | Cytochrome c oxidase subunit 5A, mitochondrial [OS=Homo sapiens] | -0.32 | 0.977036887 | 483437.4688 | 5401147.344 | 37692332.75 | 448407.1875 | 29758988.41 | 498378.8496 | 3268576.39 | 38764181.64 | 37692332.75 | 3277790.123 | 30914433.26 | 4217372.628 | |
| P36543 | V-type proton ATPase subunit E 1 [OS=Homo sapiens] | -0.32 | 0.977036887 | 596085.3301 | 8438369.914 | 68205015.94 | 487203.8252 | 50529788.63 | 135054.3379 | 4030201.551 | 60562410.77 | 68205015.94 | 3561387.798 | 52491696.18 | 1142854.414 | Met-loss+Acetyl [N-Term] |
| P56385 | ATP synthase subunit e, mitochondrial [OS=Homo sapiens] | -0.27 | 0.977036887 | 102057.1484 | 865132.6641 | 6360491.781 | 103581.2578 | 4951922.375 | 136185 | 690020.1316 | 6209080.7 | 6360491.781 | 757163.6522 | 5144189.435 | 1152422.282 | Met-loss [N-Term] |
| Q9UNM6 | 26S proteasome non-ATPase regulatory subunit 13 [OS=Homo sapiens] | -0.7 | 0.977036887 | 26546.70898 | 539633.9609 | 4826616.313 | 28349.16797 | 3073380.563 | | 179485.3561 | 3872967.64 | 4826616.313 | 207228.2188 | 3192709.946 | | |
| P83881 | Large ribosomal subunit protein eL42 [OS=Homo sapiens] | -0.47 | 0.977036887 | 28786.78125 | 102681.5078 | 1529640.25 | 34421.70313 | 891381.125 | | 194630.7426 | 736947.9791 | 1529640.25 | 251617.5513 | 925990.5586 | | |
| P02545 | Prelamin-A/C [OS=Homo sapiens] | -0.31 | 0.977036887 | 1777007.344 | 5321804.957 | 28621524.03 | 2191295.521 | 27779603.19 | 1520928.813 | 12014551.26 | 38194739.17 | 28621524.03 | 16018045.69 | 28858194.94 | 12870376.72 | |
| Q92973 | Transportin-1 [OS=Homo sapiens] | -0.4 | 0.977036887 | 63311.48047 | 126471.8594 | 2072648.578 | 62761.6875 | 1120478.781 | 63202.88281 | 428056.2092 | 907691.9804 | 2072648.578 | 458778.6394 | 1163983.333 | 534834.3097 | |
| P05783 | Keratin, type I cytoskeletal 18 [OS=Homo sapiens] | -0.53 | 0.977036887 | 858963.1563 | 310940.5625 | 609051.75 | 766839.1094 | 574096.875 | 546269.3828 | 5807548.802 | 2231628.888 | 609051.75 | 5605480.305 | 596387.1918 | 4622631.046 | |
| Q15363 | Transmembrane emp24 domain-containing protein 2 [OS=Homo sapiens] | -0.2 | 0.977036887 | 41046.125 | 144179.4375 | 911426.3125 | 61308.98438 | 869121.8125 | 60918.88281 | 277517.5772 | 1034779.751 | 911426.3125 | 448159.5947 | 902866.9893 | 515506.6856 | |
| Q93008 | Probable ubiquitin carboxyl-terminal hydrolase FAF-X [OS=Homo sapiens] | -0.37 | 0.977036887 | 43245.51172 | 1307344.324 | 13210405.05 | 58734.20313 | 9211269.063 | 34605.99609 | 292387.8841 | 9382845.833 | 13210405.05 | 429338.3251 | 9568912.719 | 292842.244 | |
| P16403 | Histone H1.2 [OS=Homo sapiens] | -0.35 | 0.977036887 | 2668675.641 | 10917009.75 | 52764142.05 | 3012912.543 | 57678458.53 | 2739629.883 | 18043223.28 | 78351676.37 | 52764142.05 | 22023944.42 | 59917925.72 | 23183247.22 | Met-loss+Acetyl [N-Term] |
| P51114 | RNA-binding protein FXR1 [OS=Homo sapiens] | -0.36 | 0.977036887 | | 371501.3672 | 1901438.719 | 39467.97266 | 1960998.75 | | | 2666275.434 | 1901438.719 | 288505.0341 | 2037137.962 | | |
| P20340 | Ras-related protein Rab-6A [OS=Homo sapiens] | -0.75 | 0.977036887 | 525525.0625 | 145572.1797 | 952300.5625 | 433986.7344 | 890834.5625 | 310848.3281 | 3553135.457 | 1044775.5 | 952300.5625 | 3172378.746 | 925422.7749 | 2630455.188 | |
| O00487 | 26S proteasome non-ATPase regulatory subunit 14 [OS=Homo sapiens] | -0.41 | 0.977036887 | 42575.76563 | 204448.5625 | 1664374.063 | 29595.11133 | 1127625.375 | 20442.02734 | 287859.6536 | 1467332.903 | 1664374.063 | 216335.8802 | 1171407.405 | 172984.1599 | |
| P35580 | Myosin-10 [OS=Homo sapiens] | -0.29 | 0.977036887 | 458047.7539 | 3601901.113 | 31123776.88 | 758738.2344 | 22593803.86 | 396312.2148 | 3096913.604 | 25850942.42 | 31123776.88 | 5546264.109 | 23471047.87 | 3353666.169 | |
| P02794 | Ferritin heavy chain [OS=Homo sapiens] | -0.3 | 0.977036887 | 90548.39844 | 883891.125 | 7866415.594 | 75569.38281 | 5003427 | 101498.5957 | 612208.1477 | 6343710.685 | 7866415.594 | 552400.995 | 5197693.818 | 858899.6096 | |
| P60891 | Ribose-phosphate pyrophosphokinase 1 [OS=Homo sapiens] | -1.17 | 0.977036887 | 72414.35156 | 2059659.078 | 21076589.63 | 44145.24707 | 14409488.97 | | 489601.7688 | 14782229.32 | 21076589.63 | 322695.2173 | 14968962.62 | | |
| P35527 | Keratin, type I cytoskeletal 9 [OS=Homo sapiens] | -0.82 | 0.977036887 | 284823009 | 175077393.2 | 295226572.6 | 177801706 | 305645310.9 | 322400402.7 | 1925721158 | 1256535221 | 295226572.6 | 1299704135 | 317512525.4 | 2728210948 | |
| P25705 | ATP synthase subunit alpha, mitochondrial [OS=Homo sapiens] | -0.31 | 0.977036887 | 9513975.648 | 74358944.36 | 570676283.6 | 9694797.719 | 395897752.6 | 10428447.89 | 64325084.81 | 533676169.5 | 570676283.6 | 70867535.33 | 411269176.4 | 88247426.06 | |
| P12235 | ADP/ATP translocase 1 [OS=Homo sapiens] | -0.95 | 0.977036887 | | 432929.6719 | 18936623.75 | | 9886863 | 79168.40625 | | 3107148.05 | 18936623.75 | | 10270737.77 | 669937.4779 | |
| P38606 | V-type proton ATPase catalytic subunit A [OS=Homo sapiens] | -0.28 | 0.977036887 | 327764.2793 | 15955429.92 | 140303507.3 | 328608.5762 | 99275226.03 | 713338.2422 | 2216052.032 | 114512555.2 | 140303507.3 | 2402080.018 | 103129760.6 | 6036398.173 | |
| P25398 | Small ribosomal subunit protein eS12 [OS=Homo sapiens] | -0.39 | 0.977036887 | 121914.6563 | 944229.3086 | 10528569.81 | 168275.1133 | 6719105.438 | 182534.5371 | 824279.0283 | 6776759.473 | 10528569.81 | 1230066.153 | 6979986.476 | 1544640.51 | Met-loss+Acetyl [N-Term] |
| Q9UPY8 | Microtubule-associated protein RP/EB family member 3 [OS=Homo sapiens | -0.29 | 0.977036887 | | 5245412.133 | 33539569.78 | 75451.03125 | 29204752.17 | 95856.64063 | | 37646465.79 | 33539569.78 | 551535.8626 | 30338677.83 | 811156.3577 | Met-loss+Acetyl [N-Term] |
| P30101 | Protein disulfide-isomerase A3 [OS=Homo sapiens] | -0.32 | 0.977036887 | 822348.8672 | 3724374.867 | 23792550.03 | 908972.1641 | 20322626.56 | 428460.8164 | 5559995.377 | 26729939.89 | 23792550.03 | 6644451.882 | 21111688.14 | 3625713.493 | |
| P09012 | U1 small nuclear ribonucleoprotein A [OS=Homo sapiens] | -0.24 | 0.977036887 | | 152193.75 | 1282359.469 | 40204.92188 | 1303683.5 | | | 1092298.69 | 1282359.469 | 293892.0238 | 1354301.296 | | |
| Q5T750 | Protein KPLCE [OS=Homo sapiens] | -0.36 | 0.977036887 | 686925.5 | 297853.5625 | 480103.5625 | 494442.9375 | 1039280.75 | 692765.375 | 4644382.399 | 2137703.133 | 480103.5625 | 3614304.636 | 1079632.646 | 5862306.823 | |
| Q8TD22 | Sideroflexin-5 [OS=Homo sapiens] | -0.35 | 0.977117324 | | 1705560.125 | 14411238.66 | 81754.50781 | 8792596.188 | 43293.5625 | | 12240851.48 | 14411238.66 | 597613.3426 | 9133984.136 | 366358.0138 | |
| P22314 | Ubiquitin-like modifier-activating enzyme 1 [OS=Homo sapiens] | -0.35 | 0.977120188 | 31999.02344 | 6709409.453 | 53479339.63 | 142979.8555 | 39517172.81 | 102103.0645 | 216349.0819 | 48153614.44 | 53479339.63 | 1045161.565 | 41051496.27 | 864014.7343 | |
| P61026 | Ras-related protein Rab-10 [OS=Homo sapiens] | -0.38 | 0.97719056 | 718183.5684 | 9457488.184 | 58604113.88 | 1416099.781 | 51405340.8 | 890846.0703 | 4855721.798 | 67876650.36 | 58604113.88 | 10351479.65 | 53401243.2 | 7538501.756 | |

| | | | | | | | | | | | | | | | | |
|---|---|---|---|---|---|---|---|---|---|---|---|---|---|---|---|---|
| Q15286 | Ras-related protein Rab-35 [OS=Homo sapiens] | -0.49 | 0.97719056 | 174690.8594 | 995343.6484 | 8557029.5 | 345707.6875 | 5788977.219 | 113535.0742 | 1181105.014 | 7143608.482 | 8557029.5 | 2527071.989 | 6013744.4 | 960754.4838 | |
| O96008 | Mitochondrial import receptor subunit TOM40 homolog [OS=Homo sapiens] | -0.49 | 0.97719056 | | 180891.7813 | 1260385.5 | 50845.35547 | 892691.1875 | | | 1298265.242 | 1260385.5 | 371672.0173 | 927351.4866 | | |
| P04181 | Ornithine aminotransferase, mitochondrial [OS=Homo sapiens] | -0.38 | 0.97719056 | 19422.86133 | 531790.3438 | 4511502.875 | 51023.51953 | 3396666.141 | | 131320.2018 | 3816673.785 | 4511502.875 | 372974.3702 | 3528547.653 | | |
| P32189 | Glycerol kinase [OS=Homo sapiens] | -0.28 | 0.97719056 | 27823.04883 | 242778.5391 | 1984559.313 | 40540.55664 | 1433552.125 | | 188114.8367 | 1742428.189 | 1984559.313 | 296345.4642 | 1489212.297 | | |
| P46778 | Large ribosomal subunit protein eL21 [OS=Homo sapiens] | -0.31 | 0.97719056 | 21445.96484 | 432415.8574 | 2615079.719 | 113620.3574 | 2445100.625 | 12352.6123 | 144998.6376 | 3103460.39 | 2615079.719 | 830547.9833 | 2540035.94 | 104530.0559 | |
| P31040 | Succinate dehydrogenase [ubiquinone] flavoprotein subunit, mitochondrial | -0.38 | 0.97719056 | 607883.0898 | 4326671.883 | 34225202.94 | 630170.7656 | 25344195.8 | 792058.3867 | 4109967.563 | 31052641.98 | 34225202.94 | 4606454.956 | 26328228.59 | 6702542.379 | |
| O94811 | Tubulin polymerization-promoting protein [OS=Homo sapiens] | -1.33 | 0.97719056 | | 198974.6719 | 10575358.16 | | 4647609.25 | 54080.83594 | | 1428046.641 | 10575358.16 | | 4828060.82 | 457641.8871 | |
| P61254 | Large ribosomal subunit protein uL24 [OS=Homo sapiens] | -0.66 | 0.97719056 | 120414.6328 | 795113.3984 | 5772916.344 | 195983.7656 | 5079451.531 | 117821.4961 | 814137.1972 | 5706550.523 | 5772916.344 | 1432612.298 | 5276670.135 | 997026.9666 | |
| O14683 | Tumor protein p53-inducible protein 11 [OS=Homo sapiens] | -0.24 | 0.97719056 | 21466.3457 | 213556.6094 | 1895411.719 | 35702.48438 | 1224555.188 | 26793.39063 | 145136.435 | 1532701.604 | 1895411.719 | 260979.8725 | 1272100.688 | 226730.5531 | |
| P23588 | Eukaryotic translation initiation factor 4B [OS=Homo sapiens] | -0.3 | 0.97719056 | 131876.3438 | 259258.5469 | 2069886.5 | 229417.7813 | 1674055.563 | 149585.3359 | 891631.1445 | 1860705.654 | 2069886.5 | 1677010 | 1739053.702 | 1265818.367 | |
| P14678 | Small nuclear ribonucleoprotein-associated proteins B and B' [OS=Homo sa | -0.29 | 0.97719056 | 169704.6602 | 535471 | 4373398.172 | 226601.7422 | 2711618.25 | 119386.3594 | 1147392.747 | 3843089.956 | 4373398.172 | 1656425.171 | 2816901.578 | 1010269.125 | |
| P62760 | Visinin-like protein 1 [OS=Homo sapiens] | -0.4 | 0.97719056 | | 8915201.051 | 71784543.84 | | 60732713.61 | 138971.25 | | 63984640.83 | 71784543.84 | | 63090767.6 | 1176000.038 | |
| Q9NZ45 | CDGSH iron-sulfur domain-containing protein 1 [OS=Homo sapiens] | -0.67 | 0.97719056 | 176805.8438 | 771133.8594 | 4703210.813 | 137902.75 | 3352952.906 | 99407.38281 | 1195404.667 | 5534448.718 | 4703210.813 | 1008048.676 | 3483137.176 | 841203.3851 | |
| P35268 | Large ribosomal subunit protein eL22 [OS=Homo sapiens] | -0.48 | 0.97719056 | 157175.1719 | 327007.75 | 1485629.375 | 131493.9844 | 2656825.875 | 76273.17188 | 1062679.434 | 2346943.531 | 1485629.375 | 961201.5489 | 2759981.794 | 645437.4771 | |
| P05937 | Calbindin [OS=Homo sapiens] | -0.38 | 0.97719056 | | 1625157.813 | 10366688.06 | 25786.36523 | 9272414.844 | | | 11663801.9 | 10366688.06 | 188494.5104 | 9632432.592 | | |
| Q08209 | Protein phosphatase 3 catalytic subunit alpha [OS=Homo sapiens] | -0.42 | 0.97719056 | | 4957424.641 | 36007444.22 | 459190.875 | 30209470.05 | | | 35579571.71 | 36007444.22 | 3356617.281 | 31382405.65 | | |
| Q9Y2X3 | Nucleolar protein 58 [OS=Homo sapiens] | -0.31 | 0.97719056 | | 187354.5039 | 1335517.406 | 88061.14844 | 1053239.25 | | | 1344648.379 | 1335517.406 | 643713.9515 | 1094133.109 | | |
| Q96KP4 | Cytosolic non-specific dipeptidase [OS=Homo sapiens] | -0.9 | 0.97719056 | | 291417.207 | 2755026.984 | | 1259904.125 | 75271.22656 | | 2091509.234 | 2755026.984 | | 1308822.11 | 636958.8333 | |
| Q14165 | Malectin [OS=Homo sapiens] | -0.55 | 0.97719056 | | 225630.5391 | 1243661.563 | 48387.76172 | 1050331.5 | 25163.93164 | | 1619356.526 | 1243661.563 | 353707.371 | 1091112.46 | 212941.7743 | |
| P55072 | Transitional endoplasmic reticulum ATPase [OS=Homo sapiens] | -0.41 | 0.97719056 | 871994.502 | 7854904.492 | 78920787.38 | 759947.7285 | 55393408.97 | 673990.4609 | 5895655.23 | 56374863.54 | 78920787.38 | 5555105.332 | 57544155.1 | 5703430.079 | Met-loss+Acetyl [N-Term] |
| Q96IU4 | Putative protein-lysine deacylase ABHD14B [OS=Homo sapiens] | -0.42 | 0.97719056 | | 84982.77344 | 467802.5938 | 82814.75781 | 314921.9375 | 41397.50781 | | 609923.6803 | 467802.5938 | 605363.6131 | 327149.3334 | 350313.2536 | |
| P62263 | Small ribosomal subunit protein uS11 [OS=Homo sapiens] | -0.56 | 0.97719056 | 217255.4531 | 646065.0664 | 4063529.594 | 155913.1719 | 3314082.438 | 121985.8867 | 1468889.134 | 4636826.583 | 4063529.594 | 1139702.193 | 3442757.493 | 1032266.799 | |
| P55809 | Succinyl-CoA:3-ketoacid coenzyme A transferase 1, mitochondrial [OS=Hor | -0.53 | 0.97719056 | 78229.25781 | 1120865.203 | 6298213.5 | 235644.375 | 4571118.75 | 139726.6406 | 528917.0195 | 8044480.1 | 6298213.5 | 1722525.478 | 4748600.443 | 1182392.291 | |
| P50914 | Large ribosomal subunit protein eL14 [OS=Homo sapiens] | -0.27 | 0.97719056 | 231100.4531 | 364004.2344 | 4503339.5 | 287209.4453 | 4143041 | 189053.0391 | 1562496.773 | 2612468.307 | 4503339.5 | 2099458.504 | 4303901.824 | 1599801.261 | |
| P30050 | Large ribosomal subunit protein uL11 [OS=Homo sapiens] | -0.47 | 0.97719056 | 278664.1719 | 528432.6641 | 6951669.625 | 211840.5508 | 5058508.25 | 111856.9531 | 1884080.552 | 3792575.627 | 6951669.625 | 1548523.048 | 5254913.694 | 946553.9173 | |
| P61421 | V-type proton ATPase subunit d 1 [OS=Homo sapiens] | -0.39 | 0.97719056 | 45845.13672 | 5850377.594 | 53117348.5 | 24561.78711 | 36635751.94 | 49547.95508 | 309964.248 | 41988319.38 | 53117348.5 | 179543.0257 | 38058199.18 | 419283.8233 | |
| P12236 | ADP/ATP translocase 3 [OS=Homo sapiens] | -0.39 | 0.97719056 | 414174.4531 | 3851338.832 | 25179480 | 561100.9844 | 22759591.19 | 241312.7109 | 2800281.166 | 27641163.72 | 25179480 | 4101565.086 | 23643272.18 | 2042032.126 | |
| O75874 | Isocitrate dehydrogenase [NADP] cytoplasmic [OS=Homo sapiens] | -1.62 | 0.97719056 | | 931332.9883 | 8452059.344 | | 7666333.906 | 26830.01563 | | 6684202.23 | 8452059.344 | | 7963992.749 | 227040.4806 | |
| P49411 | Elongation factor Tu, mitochondrial [OS=Homo sapiens] | -0.37 | 0.97719056 | 285153.4297 | 3669675.656 | 27920245.88 | 457639.6406 | 22103104.94 | 195200.5703 | 1927955.171 | 26337362.16 | 27920245.88 | 3345277.987 | 22961296.7 | 1651822.791 | |
| P62820 | Ras-related protein Rab-1A [OS=Homo sapiens] | -0.36 | 0.97719056 | 2402975.168 | 4490136.434 | 40153254.81 | 2718870.242 | 30149883.56 | 1126616.734 | 16246791.79 | 32225831.51 | 40153254.81 | 19874538.75 | 31320505.61 | 9533636.072 | Met-loss+Acetyl [N-Term] |
| P62829 | Large ribosomal subunit protein uL14 [OS=Homo sapiens] | -0.37 | 0.97719056 | 144040.8438 | 827392.8594 | 7614528.688 | 160661.1563 | 4913957.438 | 114052.5313 | 973876.7297 | 5938221.094 | 7614528.688 | 1174409.255 | 5104750.443 | 965133.2994 | |
| P60468 | Protein transport protein Sec61 subunit beta [OS=Homo sapiens] | -0.21 | 0.97719056 | 40591.96094 | 127092.2422 | 622155.0625 | 37662.89453 | 757953.0625 | 44229.68359 | 274446.9217 | 912144.4847 | 622155.0625 | 275310.1804 | 787381.9178 | 374279.6411 | |
| P26440 | Isovaleryl-CoA dehydrogenase, mitochondrial [OS=Homo sapiens] | -0.54 | 0.97719056 | 108617.5781 | 454523.4375 | 4276828.969 | 122117.5703 | 2159521.75 | 109961.7656 | 734375.9521 | 3262127.094 | 4276828.969 | 892661.3509 | 2243368.965 | 930516.4954 | |
| P55010 | Eukaryotic translation initiation factor 5 [OS=Homo sapiens] | -0.28 | 0.97719056 | | 78324.10938 | 1404633.406 | 85562.38867 | 704992.5469 | | | 562134.2669 | 1404633.406 | 625448.3877 | 732365.1174 | | |
| O43175 | D-3-phosphoglycerate dehydrogenase [OS=Homo sapiens] | -0.47 | 0.97719056 | 71161.11328 | 1118634.203 | 10105915.38 | 52445.42578 | 6510750.75 | 40514.24219 | 481128.4805 | 8028468.152 | 10105915.38 | 383368.2942 | 6763542.053 | 342838.899 | |
| P14625 | Endoplasmin [OS=Homo sapiens] | -0.39 | 0.97719056 | 2551021 | 5937608.68 | 32626518.63 | 3059333.121 | 33499936.97 | 1900439.607 | 17247746.71 | 42614379.26 | 32626518.63 | 22363271.97 | 34800630.7 | 16081866.21 | |
| P21283 | V-type proton ATPase subunit C 1 [OS=Homo sapiens] | -0.38 | 0.97719056 | 62227.05859 | 4263937.586 | 28863173.53 | 102602.5557 | 23387354.38 | 98591.41016 | 420724.3081 | 30602396.22 | 28863173.53 | 750009.4844 | 24295409.38 | 834298.4758 | |
| P21796 | Voltage-dependent anion-selective channel protein 1 [OS=Homo sapiens] | -0.39 | 0.97719056 | 3178214.385 | 21802040.79 | 199580960.3 | 2700294.203 | 139009591.9 | 3334976.799 | 21488273.4 | 156473840.7 | 199580960.3 | 19738750.66 | 144406882.8 | 28221181.29 | Met-loss+Acetyl [N-Term] |
| P49207 | Large ribosomal subunit protein eL34 [OS=Homo sapiens] | -0.36 | 0.97719056 | 269041.2695 | 690685.6094 | 4705599.875 | 339890.6758 | 3589642.875 | 280433.6172 | 1819019.001 | 4957069.436 | 4705599.875 | 2484550.495 | 3729017.048 | 2373080.363 | |
| P51608 | Methyl-CpG-binding protein 2 [OS=Homo sapiens] | -1.06 | 0.97719056 | | 380117.582 | 5934136.719 | | 3252987.094 | 10117.32617 | | 2728114.242 | 5934136.719 | | 3379290.016 | 85614.65741 | |
| O60749 | Sorting nexin-2 [OS=Homo sapiens] | -0.29 | 0.97719056 | 26397.42578 | 511158.4414 | 2619962.094 | 37423.63672 | 2094498.25 | | 178476.0351 | 3668598.06 | 2619962.094 | 273561.2412 | 2175820.813 | | |
| P46926 | Glucosamine-6-phosphate isomerase 1 [OS=Homo sapiens] | -0.3 | 0.97719056 | 65331.35742 | 379656.2969 | 2517270.188 | 49477.63965 | 2153837.375 | 176033.9473 | 441712.8299 | 2724803.586 | 2517270.188 | 361674.2172 | 2237463.884 | 1489631.335 | |
| P30153 | Serine/threonine-protein phosphatase 2A 65 kDa regulatory subunit A alpha | -0.42 | 0.97719056 | 90929.14063 | 7131047.42 | 63090485.44 | 62759.04688 | 49476013.39 | 64144.91797 | 614782.3895 | 51179721.61 | 63090485.44 | 458759.3368 | 51397006.28 | 542805.9828 | Met-loss+Acetyl [N-Term] |
| P01112 | GTPase HRas [OS=Homo sapiens] | -0.36 | 0.97719056 | 23966.93164 | 941474.8047 | 13564639.58 | 35539.15625 | 8082386.891 | 33958.62109 | 162043.1844 | 6756990.323 | 13564639.58 | 259785.9681 | 8396199.719 | 287364.0388 | |
| O14617 | AP-3 complex subunit delta-1 [OS=Homo sapiens] | -0.38 | 0.97719056 | 46102.96094 | 481551.8555 | 6135438.813 | 32738.9375 | 3681922.594 | | 311707.4273 | 3456110.786 | 6135438.813 | 239316.7838 | 3824879.688 | | |
| Q9Y224 | RNA transcription, translation and transport factor protein [OS=Homo sapie | -0.61 | 0.97728572 | | 278095.75 | 2244166.25 | 21551.56055 | 1259664.875 | | | 1995900.774 | 2244166.25 | 157538.7154 | 1308573.57 | | |
| Q92581 | Sodium/hydrogen exchanger 6 [OS=Homo sapiens] | -0.26 | 0.97728572 | | 62410.79688 | 456630.1563 | 56005.875 | 359417.7813 | | | 447923.9895 | 456630.1563 | 409394.6507 | 373372.8062 | | |
| P99999 | Cytochrome c [OS=Homo sapiens] | -0.43 | 0.97728572 | 1933429.688 | 5630315.016 | 33680121.63 | 1815924.375 | 28555344.13 | 1816091.938 | 13072140.73 | 40408924.26 | 33680121.63 | 13274138.21 | 29664055.38 | 15368100.86 | |
| Q9NYF8 | Bcl-2-associated transcription factor 1 [OS=Homo sapiens] | -0.23 | 0.97728572 | 89230.03125 | 48662.96875 | 366868.5313 | | 286342.8125 | 45551.42969 | 603294.5154 | 349255.4525 | 366868.5313 | | 297460.5738 | 385464.4973 | |
| P83731 | Large ribosomal subunit protein eL24 [OS=Homo sapiens] | -0.41 | 0.97728572 | 348067.9922 | 643135.375 | 3606112.656 | 389539.4141 | 3308601.375 | 203567.332 | 2353327.772 | 4615800.108 | 3606112.656 | 2847475.418 | 3437063.619 | 1722623.853 | |
| O43426 | Synaptojanin-1 [OS=Homo sapiens] | -0.71 | 0.97728572 | | 3798010.449 | 24068620.39 | 601353.3545 | 83982346.64 | 242303.0303 | | 27258424.47 | 24068620.39 | 4395803.949 | 87243108.36 | 2050412.389 | |
| Q9Y512 | Sorting and assembly machinery component 50 homolog [OS=Homo sapien | -0.61 | 0.97728572 | 43776.01953 | 723915.6523 | 6736604.469 | 104975.1328 | 3795318.156 | 101827.5762 | 295974.7085 | 5195562.359 | 6736604.469 | 767352.6719 | 3942678.032 | 861683.5022 | |
| P38117 | Electron transfer flavoprotein subunit beta [OS=Homo sapiens] | -0.52 | 0.97728572 | 275376.0234 | 1008163.477 | 7592313.094 | 306475.4531 | 4936836.438 | 335557.7383 | 1861849.002 | 7235616.738 | 7592313.094 | 2240290.167 | 5128517.76 | 2839550.719 | |
| P61224 | Ras-related protein Rap-1b [OS=Homo sapiens] | -0.51 | 0.97728572 | | 389478.4297 | 3187960.531 | 24447.32031 | 2650156.844 | | | 2795297.301 | 3187960.531 | 178706.2903 | 2753053.825 | | |
| P05165 | Propionyl-CoA carboxylase alpha chain, mitochondrial [OS=Homo sapiens] | -0.3 | 0.97728572 | | 194150.0313 | 1571908.828 | 34544.69922 | 1103811.938 | 46094.01563 | | 1393420.064 | 1571908.828 | 252516.6345 | 1146669.37 | 390055.9584 | |
| Q63HM9 | PI-PLC X domain-containing protein 3 [OS=Homo sapiens] | -1.91 | 0.97728572 | | 484383 | 8091308.125 | | 3960673.938 | 46957.66016 | | 3476429.988 | 8091308.125 | | 4114454.041 | 397364.2758 | |
| Q14247 | Src substrate cortactin [OS=Homo sapiens] | -0.39 | 0.97728572 | 22355.34961 | 970764.7109 | 6053031.531 | 38241.52734 | 5040072.656 | 21801.03906 | 151147.0927 | 6967204.778 | 6053031.531 | 279539.9005 | 5235762.306 | 184484.3646 | |
| Q86Y46 | Keratin, type II cytoskeletal 73 [OS=Homo sapiens] | -0.34 | 0.97728572 | 307248.4375 | 256541.875 | 803744.5313 | 224027.7813 | 1335184.906 | 201627.0313 | 2077342.063 | 1841208.026 | 803744.5313 | 1637609.898 | 1387025.799 | 1706204.674 | |
| P26639 | Threonine--tRNA ligase 1, cytoplasmic [OS=Homo sapiens] | -0.34 | 0.97728572 | | 210964.4531 | 1054380.813 | 59967.63281 | 679734.375 | 16585.53711 | | 1514097.628 | 1054380.813 | 438354.5135 | 706126.2528 | 140349.8369 | |
| Q16864 | V-type proton ATPase subunit F [OS=Homo sapiens] | -0.4 | 0.978430385 | 85247.95313 | 920704.3203 | 5844351.344 | 59858.95313 | 5096639.25 | 41876.74219 | 576371.2266 | 6607919.991 | 5844351.344 | 437560.081 | 5294525.197 | 354368.6222 | |
| P19021 | Peptidyl-glycine alpha-amidating monooxygenase [OS=Homo sapiens] | -1.49 | 0.978430385 | | 375603.5 | 4861347.938 | | 4182925.5 | 60016.0957 | | 2695716.553 | 4861347.938 | | 4345334.91 | 507867.1366 | |
| P09543 | 2',3'-cyclic-nucleotide 3'-phosphodiesterase [OS=Homo sapiens] | -0.45 | 0.979117894 | 130298.6797 | 8034599.84 | 81385107.31 | 195097.6406 | 44583859.06 | 605436.6875 | 880964.3761 | 57664541.95 | 81385107.31 | 1426134.855 | 46314905.49 | 5123315.558 | |
| Q9NZ01 | Very-long-chain enoyl-CoA reductase [OS=Homo sapiens] | -0.32 | 0.979354808 | 65824.51563 | 266687.5 | 2980070.563 | 48632.34766 | 2671576.875 | | 445047.1293 | 1914023.453 | 2980070.563 | 355495.2579 | 2775305.527 | | |
| O14949 | Cytochrome b-c1 complex subunit 8 [OS=Homo sapiens] | -0.44 | 0.979354808 | 117280.3125 | 536789.25 | 4374263.344 | 79989 | 3238766.844 | 107853.1719 | 792945.6966 | 3852551.072 | 4374263.344 | 584707.742 | 3364517.64 | 912673.1909 | |
| P21912 | Succinate dehydrogenase [ubiquinone] iron-sulfur subunit, mitochondrial [( | -1.17 | 0.979993919 | 123130.9375 | 1479572.234 | 7816039.656 | 184550.125 | 5077254.406 | 154615.4063 | 832502.4459 | 10618930.24 | 7816039.656 | 1349034.078 | 5274387.702 | 1308383.738 | |
| O43148 | mRNA cap guanine-N7 methyltransferase [OS=Homo sapiens] | -0.4 | 0.980306615 | 83652.59375 | 230016.7656 | 1731337.625 | 90616.21094 | 1207640.969 | 39714.32031 | 565584.8182 | 1650836.594 | 1731337.625 | 662391.0799 | 1254529.745 | 336069.8143 | |
| P11766 | Alcohol dehydrogenase class-3 [OS=Homo sapiens] | -0.38 | 0.980630457 | 132006.8516 | 157176.5156 | 3369503.938 | 144899.8711 | 2537902.063 | 30199.98633 | 892513.5229 | 1128060.135 | 3369503.938 | 1059196.595 | 2636440.556 | 255557.7866 | |
| Q96P63 | Serpin B12 [OS=Homo sapiens] | -0.33 | 0.980630457 | 472776.625 | 243802.5469 | 329656.4375 | 349190.7656 | 370133.7813 | 546143.4375 | 3196497.198 | 1749777.521 | 329656.4375 | 2552532.775 | 384504.8737 | 4621565.274 | |
| P62081 | Small ribosomal subunit protein eS7 [OS=Homo sapiens] | -0.43 | 0.980630457 | 163257.4531 | 748561.8535 | 5308959.25 | 86516.90625 | 4121583.609 | 69000.75781 | 1103802.438 | 5372448.817 | 5308959.25 | 632425.7698 | 4281611.313 | 583896.9843 | |
| Q9UIJ7 | GTP:AMP phosphotransferase AK3, mitochondrial [OS=Homo sapiens] | -0.46 | 0.980630457 | 47421.4375 | 365457.7031 | 2532596.844 | 31935.73828 | 1871472.781 | 33860.66016 | 320621.7991 | 2622899.892 | 2532596.844 | 233445.5165 | 1944135.99 | 286535.0755 | |
| P15121 | Aldo-keto reductase family 1 member B1 [OS=Homo sapiens] | -0.32 | 0.980630457 | 35245.39844 | 554751.5938 | 8337441.188 | 29992.34375 | 4915252.969 | 59107.27734 | 238298.197 | 3981467.302 | 8337441.188 | 219239.5903 | 5106096.275 | 500176.55 | |
| O43747 | AP-1 complex subunit gamma-1 [OS=Homo sapiens] | -0.5 | 0.981344014 | 53308.80469 | 540893.0898 | 5736155.406 | 34862.23047 | 3437182.125 | 34476.55469 | 360426.9665 | 3882004.442 | 5736155.406 | 254837.7409 | 3570636.742 | 291746.8873 | |
| Q9Y3B3 | Transmembrane emp24 domain-containing protein 7 [OS=Homo sapiens] | -0.35 | 0.981344014 | 302923.4609 | 245500.7344 | 2317931.25 | 245781 | 2854255.75 | 99054.48438 | 2048100.398 | 1761965.459 | 2317931.25 | 1796622.705 | 2965077.229 | 838217.0942 | |
| Q16891 | MICOS complex subunit MIC60 [OS=Homo sapiens] | -0.47 | 0.981344014 | 148668.9297 | 4321757.031 | 34884864.98 | 145868.7109 | 24515959.73 | 122153.0781 | 1005167.752 | 31017367.96 | 34884864.98 | 1066278.671 | 25467834.82 | 1033681.603 | |
| P46776 | Large ribosomal subunit protein uL15 [OS=Homo sapiens] | -0.43 | 0.981533221 | 261294.0391 | 381431.3359 | 2031653.969 | 217681.8516 | 2294434.531 | 85651.65625 | 1766639.084 | 2737543.09 | 2031653.969 | 1591222.092 | 2383519.971 | 724799.9206 | |
| P49773 | Adenosine 5'-monophosphoramidase HINT1 [OS=Homo sapiens] | -0.44 | 0.982106128 | 45212.27344 | 762142.125 | 8274949.5 | 62201.5 | 3881380.5 | 67032.35156 | 305685.3865 | 5469914.795 | 8274949.5 | 454683.7517 | 4032081.897 | 567239.972 | |
| P30626 | Sorcin [OS=Homo sapiens] | -0.45 | 0.983254406 | 65586.70313 | 337731 | 2352009.125 | 80270.875 | 1714559.813 | 70229.89844 | 443439.2516 | 2423904.588 | 2352009.125 | 586768.2065 | 1781130.601 | 594298.1962 | |
| P04264 | Keratin, type II cytoskeletal 1 [OS=Homo sapiens] | -0.49 | 0.984678404 | 551595411.5 | 263011616.7 | 559592110.2 | 362523130.9 | 756412689.2 | 547968372.6 | 3729400088 | 1887641539 | 559592110.2 | 2649990390 | 785781736.8 | 4637008207 | |
| Q5TEC6 | Histone H3-7 [OS=Homo sapiens] | -0.64 | 0.987609541 | 686428.4375 | 4758455.125 | 21492495 | 1016882.313 | 21044900.75 | 972939.5156 | 4641021.702 | 34151562.07 | 21492495 | 7433259.084 | 21862005.89 | 8233191.448 | |
| P51572 | B-cell receptor-associated protein 31 [OS=Homo sapiens] | -0.29 | 0.989202662 | 53396.1875 | 193874.5 | 767646.5625 | 86932.66406 | 1092120.125 | 52450.45703 | 361017.7718 | 1391442.568 | 767646.5625 | 635464.8978 | 1134523.602 | 443845.3237 | |
| P24752 | Acetyl-CoA acetyltransferase, mitochondrial [OS=Homo sapiens] | -0.72 | 0.989202662 | 204354.9531 | 944471.5625 | 4773826.688 | 128892.7422 | 3788690.031 | 179103.4219 | 1381667.368 | 6778498.136 | 4773826.688 | 942186.8538 | 3935792.558 | 1515605.788 | |
| P22695 | Cytochrome b-c1 complex subunit 2, mitochondrial [OS=Homo sapiens] | -0.51 | 0.989747725 | | 2263591.805 | 13045934 | 30044.42578 | 11537387.5 | | | 16245860.05 | 13045934 | 219620.3023 | 11985346.78 | | |

| | | | | | | | | | | | | | | | |
|---|---|---|---|---|---|---|---|---|---|---|---|---|---|---|---|
| Q13509 | Tubulin beta-3 chain [OS=Homo sapiens] | -0.52 | 0.992901458 | 71846.97656 | 16999076.36 | 234964744.1 | 46204.79297 | 136060320.6 | 631866.249 | 485765.6811 | 122002834.2 | 234964744.1 | 337750.1927 | 141343100.9 | 5346967.323 |
| O75534 | Cold shock domain-containing protein E1 [OS=Homo sapiens] | -0.29 | 0.993815505 | 47617.46484 | 125197.1289 | 1403314.625 | 123835.2852 | 1205780.953 | 64642.26563 | 321947.1625 | 898543.2051 | 1403314.625 | 905217.592 | 1252597.511 | 547014.6293 |
| P54727 | UV excision repair protein RAD23 homolog B [OS=Homo sapiens] | -0.59 | 0.993822058 | | 362939.7188 | 2820949.625 | 20900.7168 | 2612738.281 | | | 2604828.25 | 2820949.625 | 152781.144 | 2714182.422 | |
| P18206 | Vinculin [OS=Homo sapiens] | -0.33 | 0.993822058 | 28572.48828 | 180826.7422 | 1879547.406 | 107783.2109 | 1087157.906 | 55771.33203 | 193181.8831 | 1297798.455 | 1879547.406 | 787879.3071 | 1129368.716 | 471947.173 |
| P30038 | Delta-1-pyrroline-5-carboxylate dehydrogenase, mitochondrial [OS=Homo s | -0.31 | 0.993822058 | 75207.57813 | 109053.7891 | 1588151.469 | 167425.3789 | 722267.5313 | 116098.5625 | 508487.0952 | 782682.0152 | 1588151.469 | 1223854.721 | 750310.8333 | 982447.1887 |
| P62899 | Large ribosomal subunit protein eL31 [OS=Homo sapiens] | -0.5 | 0.993822058 | 145831.7656 | 712757.5703 | 6270754 | 98441.46875 | 3750627.125 | 100253.3828 | 985985.3587 | 5115480.501 | 6270754 | 719592.5554 | 3896251.793 | 848362.3912 |
| O94856 | Neurofascin [OS=Homo sapiens] | -0.54 | 0.993822058 | | 1845584.992 | 22945332.09 | | 12654582.03 | 108496.9844 | | 13245813.77 | 22945332.09 | | 13145918.35 | 918121.2496 |
| P62841 | Small ribosomal subunit protein uS19 [OS=Homo sapiens] | -0.61 | 0.993822058 | | 534637.75 | 2722034 | 62156.5625 | 2420378 | 63543.03125 | | 3837109.698 | 2722034 | 454355.2652 | 2514353.416 | 537712.7078 |
| Q9UNF0 | Protein kinase C and casein kinase substrate in neurons protein 2 [OS=Hom | -0.44 | 0.993822058 | | 180955.0547 | 2095547.5 | 18611.67773 | 1683671.438 | | | 1298719.357 | 2095547.5 | 136048.6075 | 1749042.93 | |
| P57088 | Transmembrane protein 33 [OS=Homo sapiens] | -0.36 | 0.993951145 | 50198.73438 | 289908.6406 | 1753455.125 | 42272.47266 | 1539808.063 | 27000.3125 | 339399.4231 | 2080682.212 | 1753455.125 | 309005.5137 | 1599593.808 | 228481.5637 |
| P30084 | Enoyl-CoA hydratase, mitochondrial [OS=Homo sapiens] | -0.37 | 0.997763353 | 37436.42969 | 616395.4141 | 5282066.938 | 117175.9219 | 2657448.438 | 27193.38867 | 253112.0116 | 4423886.68 | 5282066.938 | 856538.6328 | 2760628.528 | 230115.4095 |
| P42766 | Large ribosomal subunit protein uL29 [OS=Homo sapiens] | -0.54 | 0.997763353 | 98570.5 | 757794.5313 | 4173396 | 146790.2422 | 4412839.531 | 112530.5039 | 666446.5 | 5438711.996 | 4173396 | 1073014.928 | 4584175.756 | 952253.6268 |
| Q9BWM7 | Sideroflexin-3 [OS=Homo sapiens] | -0.58 | 0.99954882 | 14050205 | 3863605.469 | 13061539.56 | 7275818.5 | 8371366.531 | 8976767 | 94995053.77 | 27729201.71 | 13061539.56 | 53185155.54 | 8696399.501 | 75963037.89 |
| P05198 | Eukaryotic translation initiation factor 2 subunit 1 [OS=Homo sapiens] | -0.34 | 0.99954882 | 82412.35938 | 207956.0391 | 1360686.063 | 112640.8438 | 1308547.063 | 29032.30664 | 557199.4508 | 1492506.158 | 1360686.063 | 823387.8835 | 1359353.694 | 245676.6684 |
| P22676 | Calretinin [OS=Homo sapiens] | -0.58 | 0.99954882 | 160989.75 | 7226515.953 | 64660924.25 | 99727.49219 | 51718108.69 | 226243.4063 | 1088470.236 | 51864901.88 | 64660924.25 | 728993.1963 | 53726154.85 | 1914512.925 |
| Q06830 | Peroxiredoxin-1 [OS=Homo sapiens] | -0.57 | 0.99954882 | 3419559.234 | 8297946 | 50021745.94 | 3226322.609 | 44406871.31 | 3203746.43 | 23120033.72 | 59554584.51 | 50021745.94 | 23583940.39 | 46131045.88 | 27110686 |
| P25787 | Proteasome subunit alpha type-2 [OS=Homo sapiens] | -0.4 | 0.999743859 | 48385.10156 | 669656.832 | 5830871.813 | 43754.28516 | 3814683.75 | 37707.40234 | 327137.2428 | 4806145.328 | 5830871.813 | 319837.3436 | 3962795.529 | 319086.9088 |
| P83111 | Serine beta-lactamase-like protein LACTB, mitochondrial [OS=Homo sapier | -0.29 | 0.999743859 | 44331.69922 | 66461.30469 | 525565.25 | 35288.14063 | 374794.2188 | | 299731.7228 | 476994.5944 | 525565.25 | 257951.0811 | 389346.2608 | |
