## Supplemental table 3 for "AquIRE reveals multiple mechanisms of clinically induced RNA damage and the conservation and dynamics of glycoRNAs"

**For all parameters captured: n=8 independent experiments**

| | Total nucleoli analyzed | Total nuclei analyzed |
|---|---|---|
| Control | 663 | 305 |
| Cisplatin | 619 | 304 |
| Oxaliplatin | 361 | 285 |

**Related to Figure 5G**

| Nucleoli per nucleus | Mean | SD |
|---|---|---|
| Control | 2.19 | 0.22 |
| Cisplatin | 2.09 | 0.22 |
| Oxaliplatin | 1.27 | 0.23 |

Statistics: One-way ANOVA $p<0.0001$
Holm-Sidak's multiple comparison results:
Control vs. Cisplatin $p=0.373$
Control vs. Oxaliplatin $p<0.0001$
Cisplatin vs. Oxaliplatin. $p<0.0001$

| Nucleolar area | Mean | SD |
|---|---|---|
| Control | 0.39 | 0.03 |
| Cisplatin | 0.35 | 0.03 |
| Oxaliplatin | 0.25 | 0.03 |

Statistics: One-way ANOVA $p<0.0001$
Holm-Sidak's multiple comparison results:
Control vs. Cisplatin $p=0.041$
Control vs. Oxaliplatin $p<0.0001$
Cisplatin vs. Oxaliplatin $p<0.0001$

**Related to Supplemental Figure 6F**

| Nucleolar circularity | Mean | SD |
|---|---|---|
| Control | 0.91 | 0.02 |
| Cisplatin | 0.93 | 0.02 |
| Oxaliplatin | 0.96 | 0.03 |

Statistics: One-way ANOVA $p=0.0016$
Holm-Sidak's multiple comparison results:
Control vs. Cisplatin $p=0.016$
Control vs. Oxaliplatin $p=0.001$
Cisplatin vs. Oxaliplatin $p=0.042$

| Nucleolar roundness | Mean | SD |
|---|---|---|
| Control | 0.69 | 0.03 |
| Cisplatin | 0.72 | 0.04 |
| Oxaliplatin | 0.78 | 0.04 |

Statistics: One-way ANOVA $p<0.0001$
Holm-Sidak's multiple comparison results:
Control vs. Cisplatin $p=0.080$
Control vs. Oxaliplatin $p<0.0001$
Cisplatin vs. Oxaliplatin $p=0.0038$

| **Nucleolar optical thickness** | Mean | SD |
|---|---|---|
| Control | 1 | 0 |
| Cisplatin | 1.14 | 0.1 |
| Oxaliplatin | 1.32 | 0.16 |

<u>Statistics</u>: One-way ANOVA $p<0.0001$

Holm-Sidak's multiple comparison results:

Control vs. Cisplatin $p=0.022$

Control vs. Oxaliplatin $p<0.0001$

Cisplatin vs. Oxaliplatin. $p=0.009$
