## Supplemental table 4 for "AquIRE reveals multiple mechanisms of clinically induced RNA damage and the conservation and dynamics of glycoRNAs"

| **Tumour ID** | **Primary tumour** | **Metastatic site** | **Site sampled** | **Date sampled** | **Sample weight** | **Age (years)** | **Mucinous** | **MMR status** | ***KRAS*** |
|---|---|---|---|---|---|---|---|---|---|
| **MCRC-Om-001** | Caecum | Omentum | Metastasis | 23/05/2024 | 4000mg | 66 | Yes | MSS | WT |
| **MCRC-PN-003** | Transverse colon | Left pararenal nodule | Metastasis | 18/07/2024 | 1800mg | 60 | Yes | Unknown | WT |
| **MCRC-Ca-005** | Caecum | N/A | Primary | 22/08/2024 | 1100mg | 52 | Yes | MSS | WT |
| **MCRC-LM-007** | Caecum | Liver metastasis | Metastasis | 11/10/2024 | 1000mg | 56 | No | MSS | Gly12Val |
| **MCRC-OM-008** | Caecum | Omentum | Metastasis | 06/11/2024 | 1800mg | 81 | Yes | MSS | WT |
